# Spatially patterned, spectral single-molecule microscopy

**DOI:** 10.64898/2026.04.08.715690

**Authors:** Joseph S. Beckwith, Brendan Cullinane, Daniel F. Heraghty, Sina Krokowski, Caroline L. Jones, Shengbo Yang, Rebecca C. Gregory, R. Andres Floto, Yuxuan Zhang, Ana Mafalda Santos, Simon J. Davis, Michele Vendruscolo, David Klenerman, Viv Lindo, Praveen Kallamvalliillam Sankaran, Steven F. Lee

## Abstract

Multicolour and spectrally resolved single-molecule microscopy can reveal molecular interactions, nanoscale environments and dynamics, but usually depends on experimentally complex detection architectures based on beam splitting, spectral dispersion or engineered point spread functions. Here we show that spatially patterned detectors offer a conceptually simpler route to spectral single-molecule imaging. By replacing a conventional monochrome camera with a commercially available colour CMOS detector and fitting the raw detector response directly, we recover both molecular position and spectral fingerprint from a single image without optical splitting, channel registration or demosaicing. We term this approach spatial-spectral single-molecule microscopy, or S^3^M. We show that S^3^M retains single-molecule sensitivity across the visible spectrum (400–785 nm), enables robust spectral multiplexing, and supports applications spanning multicolour single-molecule tracking, single-molecule Förster resonance energy transfer, multicolour localisation microscopy and spectral PAINT. Although spatial patterning necessarily trades photon efficiency for spectral information, current low-noise detectors already provide sufficient performance (<15 nm localisation precision at above 1000 detected photons) for a broad range of experiments. Spatially patterned detection therefore establishes a widely accessible strategy for simplified spectral microscopy and single-molecule spectroscopy, and points towards a new class of detector-informed photonic measurement schemes for nanoscale imaging.

## Introduction

Multicolour and spectrally-resolved single-molecule microscopy and spectroscopy have enabled transformative insights across both biology and chemistry. The single-molecule Förster resonance energy transfer (FRET) community routinely splits emission into multiple spectral channels(<u>1</u>) to deduce the molecular dynamics of biomolecules at, in some cases, sub-millisecond timescales.(2) These experiments are extremely elegant, but their use of point detectors inherently decreases throughput. To increase throughput, the single-molecule imaging community has developed various strategies for multicolour imaging. In effect, these may be split into four broad classes: splitting the emission spectrum using dichroic mirrors and projecting different spectral ranges onto one detector;(3, 4) using dichroic mirrors and projecting different spectral ranges onto multiple detectors;(5, 6) using spectrally dispersive elements such as gratings or prisms to generate a spectral image as well as an image of the molecular positions;(7) and using point spread function (PSF) engineering to encode spectral information into an image.(8, 9) Despite the power of extracting spectral information from single molecules, all of these different methodologies are often prohibitively complex from either a hardware or a software perspective— multiple groups have worked on simplifying the optical designs or providing open software for these methods.(10– 15) A complementary method, that has seen much research activity in recent years, is that of sequential DNA-PAINT. These experiments do not multiplex using spectral information, but instead use separate imaging rounds with different DNA imager strands(16, 17) to produce up to 30 different “channels” of single-molecule imaging data. Instead of the technical complexity of the hardware or software, what these experiments are is low-throughput: the duration of these experiments is linear with the number of channels and prohibits dynamic processes from being studied. In any of the strategies for effective multicolour, multichannel or spectrally-resolved single-molecule microscopy there are technical challenges or throughput trade-offs that prevent their broad use in understanding biological and chemical systems. Here, we highlight that these challenges can be circumvented by carefully considering recent innovations in detector technology.

Parallel to the development of sophisticated strategies for spectral sensing in the context of single molecules, detection technology has been changing rapidly. Work from both the Huang(<u>18</u>, <u>19</u>) and Bewersdorf(<u>20</u>) labs showed that scientific CMOS (sCMOS) detectors, sufficiently characterised and with analysis algorithms to account for pixel-to-pixel variabilities, could collect high-quality single-molecule localisation microscopy data (<10 nm localisation precision at 1,000 photons). This work, as well as technological developments in commercial CMOS detectors, has enabled high-quality localisation microscopy data to be generated using industrial-grade CMOS detectors, demonstrating localisation precisions of ∼10 nm.(<u>21</u>, <u>22</u>)

In this work, we take advantage of the changing detector landscape to massively simplify multicolour single-molecule microscopy. This we term spatial-spectral single-molecule microscopy (S^3^M). We demonstrate S^3^M with a commercially available Bayer-patterned detector, but our conceptual methodology is general—with simulations, we demonstrate that detectors with arbitrary spatial patterns of pixels, and pixels with different spectral responses, are also compatible with S^3^M. We highlight S^3^M’s utility by demonstrating optically simplified six-colour multicolour imaging, three-colour single-molecule tracking, two-colour single-molecule FRET, multiplexed two-colour single-molecule localisation microscopy, and spectral PAINT.

## Results & Discussion

### Conceptual Framework

In Fig. 1a we highlight the key conceptual advancement of S^3^M. In “conventional” single-molecule imaging, the use of an unpatterned detector generates approximately identical PSFs from three dyes across the visible spectrum. However, using a spatially patterned detector (specifically, a commercially available Bayer-pattern(23) detector) where the quantum efficiencies (QEs) of individual pixels differ in sensitivity as a function of wavelength, these three dyes produce extremely different PSFs—the PSF is dependent on the dyes’ relative QE per pixel, which, combined with the spatial position, gives rise to a **spectral fingerprint**. Thus, assuming we could build an analysis routine to extract these relative pixel QEs, we used a forward model to generate these spectral fingerprints for the large numbers of chromophores used in single-molecule and single-particle microscopy(24, 25) (Fig. 1b). This highlights that, assuming a spatially-patterned detector with pixel QEs as in Fig. 1a, chromophores can be easily distinguished in terms of their spectral fingerprint—motivating the case for using these detectors to simplify multicolour single-molecule microscopy. We thus simulated (see methods) for the three dyes we highlight in Fig. 1a, ATTO 488, ATTO 565, and ATTO 647N, the yield of acceptable fits to raw images that would be generated by such a detector of 0.1–10 root-mean squared (RMS) e^*−*^ read noise and a peak QE of 0–1 (Fig. 1c). In doing so, we find a large area of read noise–QE space that would yield >80% acceptable fits(26) from dyes emitting 1,000 photons(27) across the visible spectrum. Overlaying as scatter points the peak QEs and read noise values of spatially-patterned CMOS detectors that have been made available since 2018, we observe that almost all of these detectors should yield high-quality single-molecule data at relatively low photon fluxes. Thus, this motivated us to apply this concept experimentally and detect single molecules using S^3^M.

**Fig 1.**
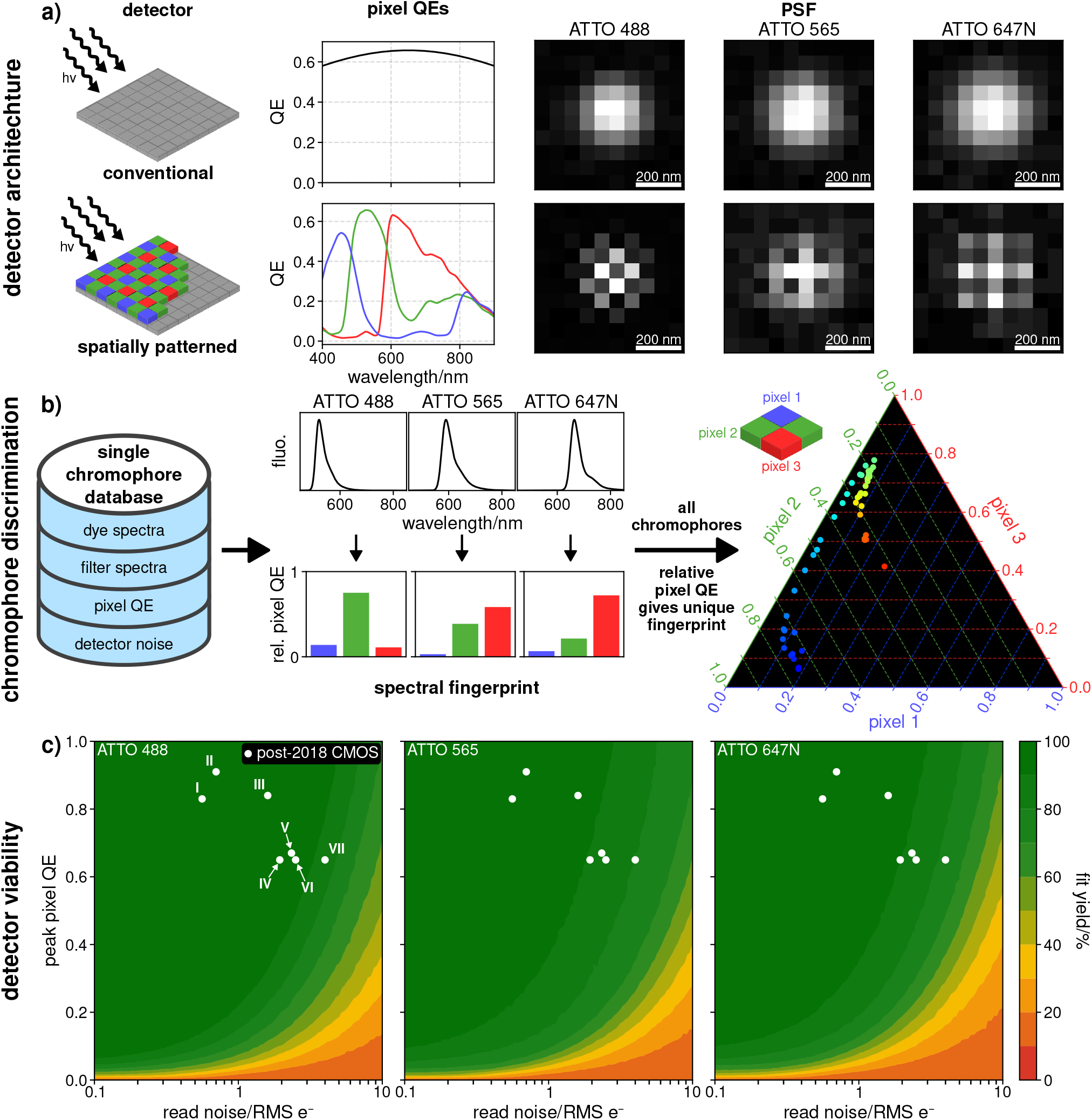
Concept of spatial-spectral single-molecule microscopy (S^3^M). **a)** A conventional detector produces approximately identical PSFs for different dye spectra, in contrast to a spatially patterned camera. Images simulated as 4,000 photons impinging on the detector. **b)** A forward model, using freely available spectral data(24) from a range of different chromophores, as well as the pixel QEs, highlights that different pixel QEs enable spectral multiplexing. **c)** Simulations of different camera QEs and read noise values, with post-2018 Bayer detectors overlaid as scatter points, highlight that current detectors should be single-molecule sensitive. Different detectors are I: ZWO ASI676MC, II: ZWO ASI585MC, III: ZWO ASI183MC, IV: Thorlabs CS505CU, V: Ximea MC050CG-SY, VI: Thorlabs CS126CU, VII: Thorlabs CS165CU. Detailed descriptions of the detectors may be found in section S19. Fit yield is defined as the percentage of fits yielding localisation coordinates within the 12*×*12 pixel ROI of the simulated image.

A detailed overview of the extraction of the relative QE of a particular PSF in the channels of the spatially patterned detector is given in Supplementary Note S3. This is distinct from the work of the Huang lab,(28, 29) which relies on a specialised detector that simultaneously gives a subsampled conventional camera image. The key conceptual difference is that S^3^M measures both the spatial and spectral information simultaneously. Furthermore, our method both outperforms and is more general than demosaicing,(30) the typical method of extracting colour from spatially patterned detectors (see SI Note S4), and is insensitive to shift variance within the pattern (see SI Note S31). S^3^M does not require specific spatial arrangements of the pixels, or specific pixel QEs (section S16). Therefore, it can work on both simple commercial hardware and more specialised detectors—the fitting routine only requires the spatial pattern of the detector. As the pattern and pixel QEs are manufacturer-provided, the calibration process is equivalent to standard CMOS detectors.(20) To remove barriers to S^3^M, we provide user accessible code (see code availability).

The original Bayer detector patent dates back to 1976,(23) and was introduced to capture colour information with minimal optical complexity. The relevance to single-molecule microscopy reflects these recent technological developments. Whilst these spatially patterned detectors provide spectral information, they will always result in losses in sensitivity and localisation precision. This is due to a fundamental trade-off: in order to spectrally distinguish dyes, the relative pixel QEs of some pixels must be lower than others, resulting in a net loss of detected photons. Until now, given this loss of detected photons and (until recently) high read noise of these detectors, this technology was not suitable for single-molecule microscopy.

### Single-Molecule Detection

Replacing a standard detector on a single-molecule microscope with a Bayer-patterned detector, we demonstrate that S^3^M is single-molecule sensitive. We explored and calibrated a range of detectors suitable for S^3^M, including a Ximea MC050CG-SY, a Thorlabs CS505CU and a ZWO ASI 585MC. We find all three detectors suitable for single-molecule microscopy (SI Notes S5 and S6), and use the Ximea MC050CG-SY for the majority of this work. Fig. 2b&c highlights the single-molecule sensitivity of the Bayer-patterned detector, showing single-step photobleaching traces of single ATTO 565 and ATTO 633 molecules. Crucially, we are able to detect single dye molecules across the visible spectrum—we have tested 12 spectrally distinct fluorophores (specifically: ATTO 488, ATTO 514, ATTO 520, ATTO Rho6G, ATTO 565, ATTO 594, ATTO 620, ATTO 633, ATTO 647N, ATTO 655, LD 655, ATTO 700) and find that they are all compatible with S^3^M, giving single-step photobleaching traces that are characteristic of single molecules (see SI Fig. S21–S32). We also characterised, using Tetraspeck™ beads, both the localisation precision and spectral precision expected as a function of number of photons (Fig. 2d, full analysis in S22). Overlaying the ranges of photons expected for fluorescent proteins(27) and organic fluorophores(31, 32) typically used in imaging, we find that in the best case we may achieve <10 nm localisation precision and <5 % error in spectral precision. Whilst more than sufficient for super-resolution microscopy, S^3^M always gives less precise localisations than super-resolution on an unpatterned detector due to the losses of the Bayer pattern. At 1,000 photons, S^3^M has a localisation precision of 14 nm, approximately 40 % higher than the 10 nm precision achieved by Bewersdorf on a typical sCMOS detector (specifically, an ORCA-Flash 4.0, Hamamatsu Photonics).(20) Additionally, to accurately assess both the localisation and spectral fingerprint of a PSF, each pixel type in the spatial pattern must be sampled, decreasing the overall signal-to-noise ratio (SNR). For a full discussion of the limits and trade-offs inherent in the S^3^M concept, see SI Note S10.

**Fig 2.**
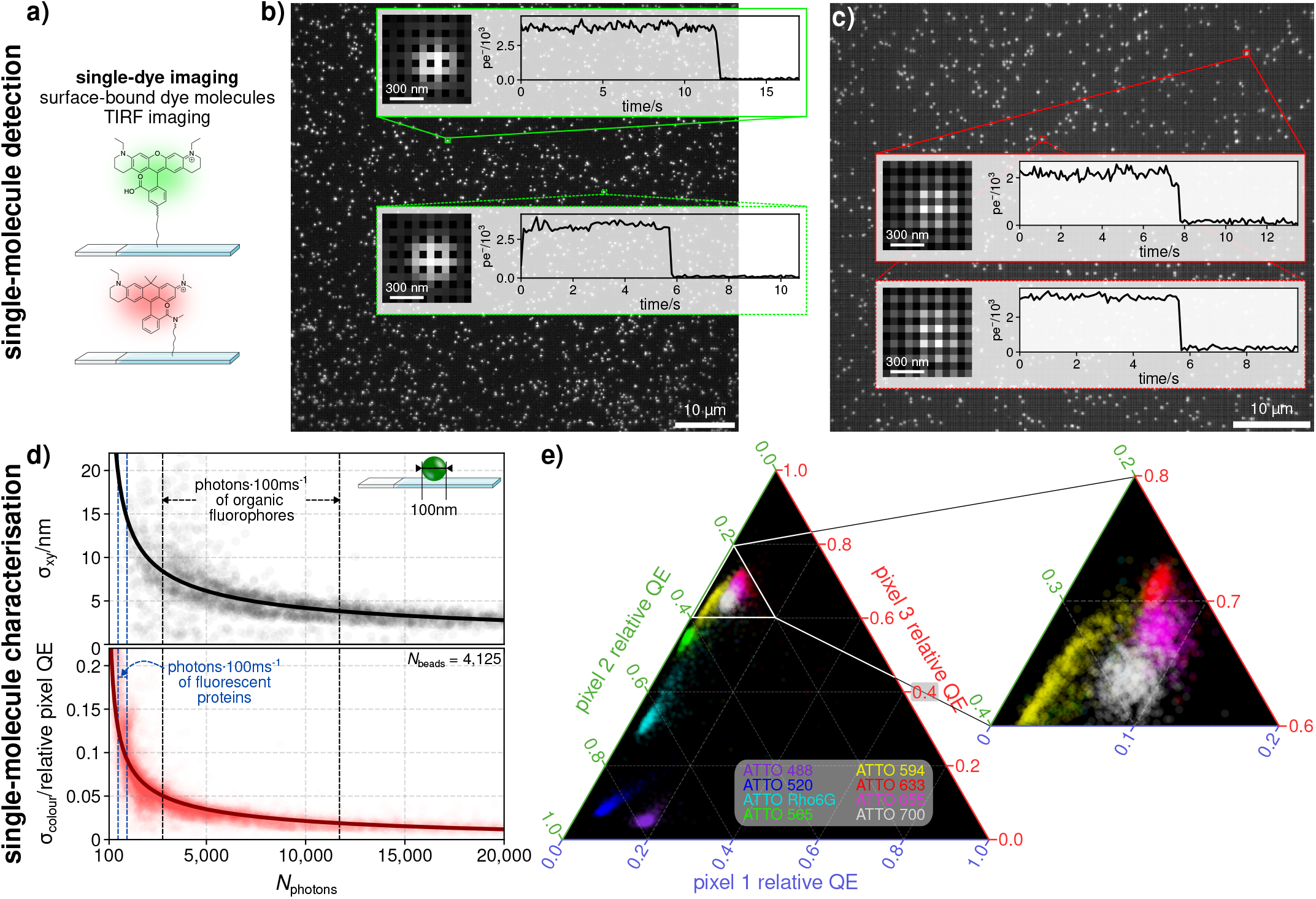
Detector is single-molecule sensitive. **a)** Schematic of single-dye imaging experiments. **b)** Exemplar field-of-view of single ATTO 565 molecules, with insets highlighting point spread functions and photobleaching traces. pe^*−*^=photoelectrons. **c)** Exemplar field-of-view of single ATTO 633 molecules, with insets highlighting point spread functions and photobleaching traces. **d)** Spatial localisation precision (black) and spectral precision (red) measured using Tetraspeck™ beads on the Ximea MC050CG-SY. Typical photon ranges (25th–75th percentile) of fluorescent proteins (550–1,000 photons)(27) and organic fluorophores (3,000–12,000 photons)(31, 32) are indicated. **e)** Analysis of different single dye molecules highlights that different spectra lead to different relative pixel QEs, enabling multiplexing. 1,000 of each molecule is shown.

The multiplexing potential of S^3^M becomes especially clear when fitted spectral fingerprints are projected into a pixel QE ternary colour map (Fig. 2e). In this representation, each molecule is positioned according to the relative fractions of photons detected by the three pixel classes shown in Fig. 1a, such that spectrally distinct dyes occupy distinct regions of ternary space. As shown in Fig. 2e, the 8 fluorophores examined here form separable clusters in this map, indicating that their effective detector responses are sufficiently different to enable reliable colour assignment from a single image. More generally, this style of representation is useful for any spatially patterned detector, regardless of the specific pixel arrangement or spectral design. This is due to the representation’s ability to quantify how well different emitters can be distinguished from one another—it captures the spectral heterogeneity of the emitters, the photon budgets, and the relationship between emission properties and detector filter design.

### Spectral Multiplexing

Next, we have quantitated the number of photons needed to confidently assign an extracted spectral fingerprint to a specific molecule (Fig. 3a–c). In the case of highly spectrally distinct fluorophores, *i*.*e*. ATTO 565 and ATTO 633, even at low numbers of photons detected (500 photons per molecule) the error rate is only approximately 5%, and at readily achievable photon numbers of 2,000 photons per molecule, this error rate is below 1% (Fig. 3c). Achieving below 5% error is possible for all dye pairs tested (SI note S23). The most challenging pair tested, ATTO 520 and ATTO Rho6G, only have a ∼ 20 nm shift between their emission spectra peaks. We also note that previous investigations of Rhodamines have shown pervasive spectral diffusion at the single-molecule level.(33) In fact, our data support this, with S^3^M showing a subpopulation of higher-wavelength ATTO Rho6G molecules (S33, ATTO Rho6G). Thus while these molecules are difficult to distinguish at <5,000 photons, S^3^M allows access to small spectral shifts relating to underlying chromophore photophysics. The low fluorophore misidentification rates are essential to enable higher-order multiplexing experiments, and thus to enable simplified multicolour imaging and single-molecule spectroscopy. As an exemplar of this, we measured 6 different quantum dots (QDs) simultaneously (Qdot 525, Qdot 585, Qdot 605, Qdot 655, Qdot 705, Qdot 800, Thermo Fisher). From the raw image (Fig. 3e), it is difficult to visually distinguish the 6 different spectral fingerprints. However, our analysis (Fig. 3f), coupled with control experiments on each individual QD sample (Fig. 3g), shows that we are capable of differentiating all species of QDs in one single image. This is highlighted by the confusion matrix in Fig. 3g, where a synthetic data set comprised of QDs from each of our control experiments is sorted without prior knowledge of their identity. In these data, we are able to correctly identify QDs 88%–100% of the time, depending on the QD spectral fingerprint. A more complete description is provided in Note S24.

**Fig 3.**
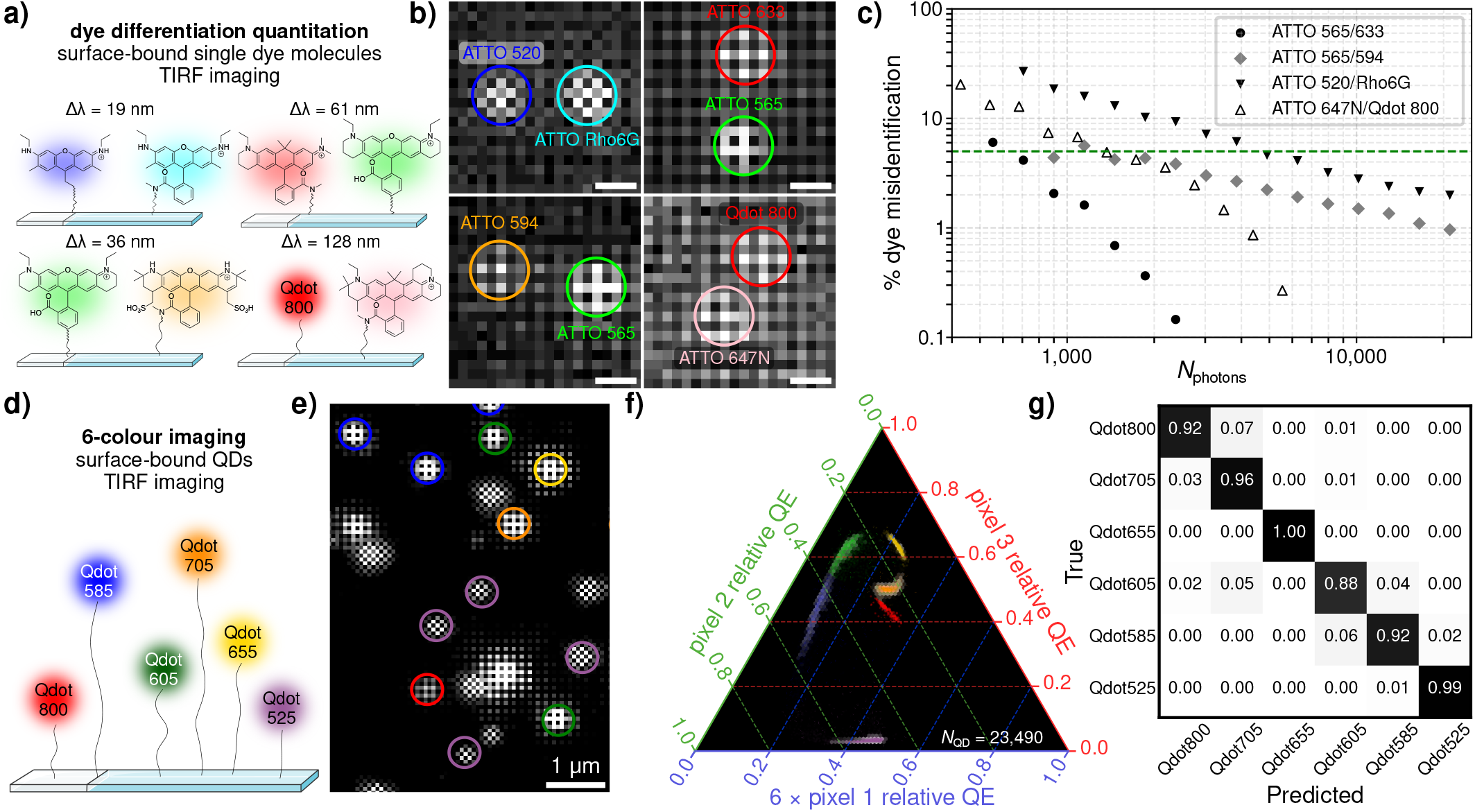
Colour differentiation at the single-molecule level. **a)** Schematic of the experiment, with two spectrally distinct dyes bound to the surface. Distance in peak emission wavelength is shown above the pairs. **b)** Representative PSFs of dye mixtures, scale bar is 300 nm. **c)** Dye misidentification rate by the number of photons detected per molecule. Misidentification rate of 5% is shown in green. **d)** Schematic of the experiment, with six spectrally distinct QDs bound to the surface. **e)** Maximum intensity projection image of a mixture of six spectrally distinct QDs, circled with their assigned identity (Qdot 525 in violet, Qdot 585 in blue, Qdot 605 in green, Qdot 655 in yellow, Qdot 705 in orange, and Qdot 800 in red). **f)** Ternary plot of the colour of all detected QDs. Coloured points show the identity of a QD determined by post processing. Pixel 1 axis has been multiplied by 6 for easier visual identification of the different clusters. **g)** Confusion matrix of a synthetic data set comprised of 4,500 of each QD.

These data, taken together, show the capability of S^3^M to enable multiplexing. We then turned to benchmarking the performance of S^3^M in two classes of single-molecule biophysics experiments on dynamic systems, as well as the field of super-resolution microscopy on static samples.

### Single-Molecule Tracking

To demonstrate S^3^M’s compatibility with single-molecule biophysics experiments, we tracked three His-tagged proteins (ICAM1, CD58 and UCHT1) diffusing in a supported lipid bilayer (Fig. 4a). These are all His-tagged proteins relevant in immunological signalling.(34–36) By simultaneously linking tracks and using the spectral fingerprint of localisations, we separated out three populations of diffusing molecules (Fig. 4b). Analysing these tracks using a mean-squared displacement analysis,(37) we find diffusion coefficients on the order of 0.1 µm^2^/s, Fig. 4c, consistent with previous work.(38, 39) These data highlight the compatibility of S^3^M with the tracking of multiple biomolecules simultaneously, opening the door to multiplexed experiments that interrogate biological assembly.

**Fig 4.**
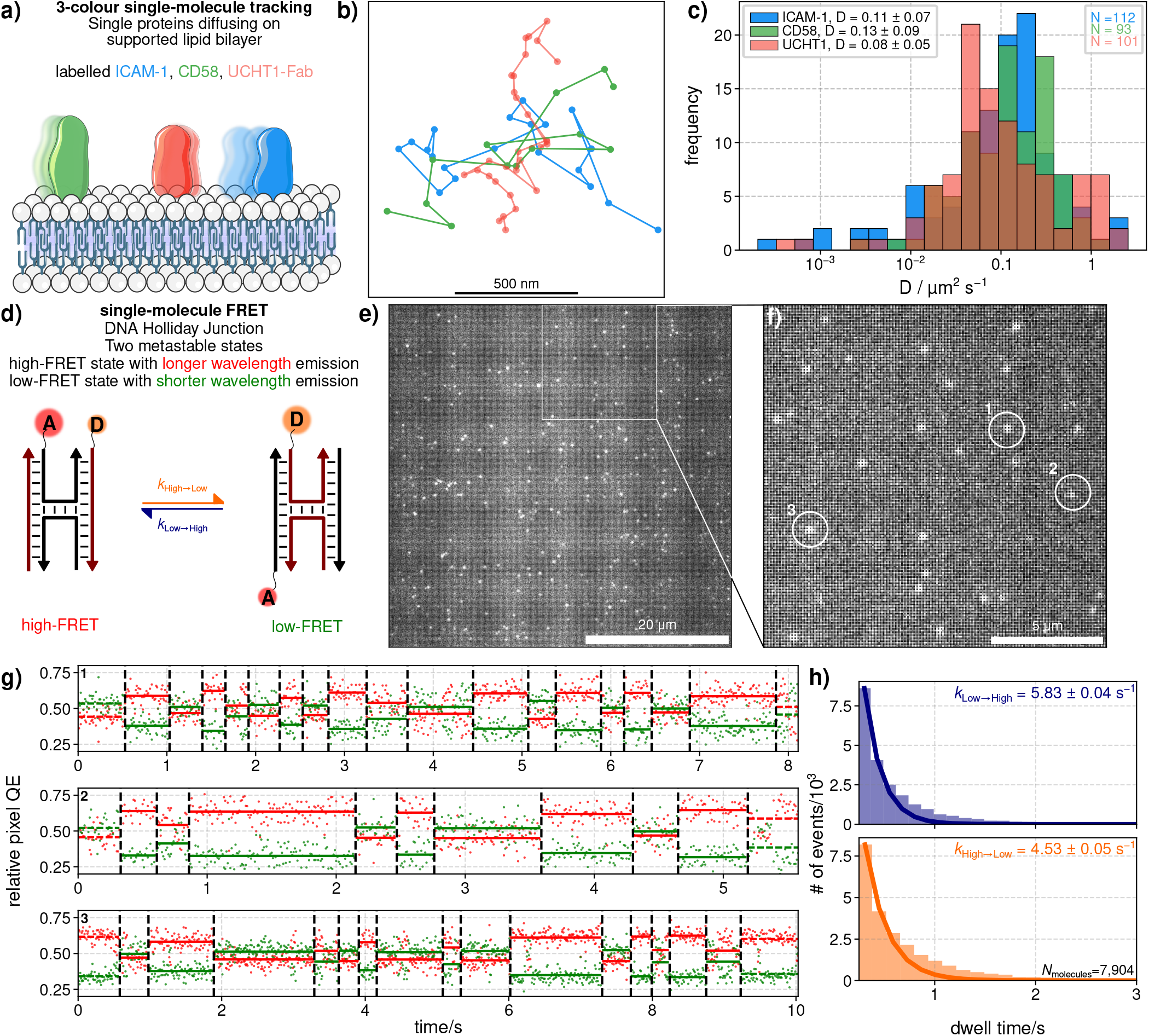
Single-molecule biophysics utility. **a)** Schematic of the single-molecule tracking experiment, with three his-tagged proteins labelled with AF488 (ICAM-1), AF555 (CD58) and JF646 (UCHT1-Fab) diffusing on a supported lipid bilayer. **b)** Single-molecule tracks of three exemplar molecules colour-coded by their identity. **c)** Extracted diffusion coefficients by molecule identity. **d)** Schematic of the single-molecule FRET experiment, where relative pixel QE gives access to Holliday Junction dynamics. **e)** Raw image of Holliday Junctions, with a zoom-in (**f)** highlighting the 3 molecules in **g). g)** Relative pixel QE traces of red and green pixels of representative molecules. Detected change points (CPs) are shown as dashed vertical black lines, with solid horizontal coloured lines showing the mean pixel QEs between CPs. The first and last CPs (dashed lines) are omitted from further analysis to avoid use of incomplete observations. **h)** Histograms showing the dwell time in the high (orange) and low (blue) states, with fits to monoexponential decays.

### Single-Molecule FRET

S^3^M is able to detect small spectral fingerprint shifts (<0.05 pixel QE, quantified in S26). We highlight this by using single-molecule FRET to study a well-characterised(40, 41) Cy3-Cy5 labelled DNA Holliday Junction (HJ, see Fig. 4d). Raw images of these HJs attached to a passivated surface are shown in Fig. 4e. This HJ transitions between two metastable states: one where the Cy3 and Cy5 are held in close proximity, leading to a higher level of energy transfer (high-FRET); and another where the Cy3 and Cy5 dyes are held at a greater distance, leading to a lower level of energy transfer (low-FRET). When the HJ is imaged, it is difficult to detect different sub-populations by eye (Fig. 4f). Examining individual molecules over time however, shows distinct and repeated anti-correlated changes in the spectral fingerprint that correspond to transitions between the high-FRET and low-FRET states (Fig. 4h). The simplified optical layout of S^3^M enables a large number of molecules to be interrogated in one field-of-view (FOV), using only one detector. We observe two populations of molecules with different energy transfer efficiencies, as expected (Fig. S80), and analysis of transitions from the high to the low state, and vice versa, gives rates of k_High*→*Low_ and k_Low*→*High_ (Fig. 4i) that are in agreement with previous work.(40)

This highlights that S^3^M enables not only multiplexing, but also spectroscopic insight into nanoscale dynamics. It is particularly encouraging that technically demanding experiments, such as three- or four-colour FRET,(42–44) could in principle benefit from S^3^M, since they would not require complex optical splitting and registration, only careful consideration of the multiple FRET labels in use.

### Single-Molecule Localisation Microscopy

The obvious advantage of S^3^M is its enabling nature for multicolour super-resolution. We have demonstrated this by using DNA-PAINT to image two cellular substructures (vimentin and mitochondria) labelled with different dyes (ATTO 655 and Cy3B, respectively) in BSC-1 cells. Using one acquisition, we super-resolved the two structures simultaneously. We observe thin vimentin structures, resolving distances of ∼180 nm between two ∼40 nm wide filaments (Fig. 5c), and achieve a resolution of 60± 1 nm for both the Cy3B and ATTO 655 channels using a Fourier ring correlation (FRC)(46) analysis (Fig. 5e). Crucially, we have demonstrated the compatibility of S^3^M with state-of-the-art single-molecule localisation microscopy methods using a widely available commercial two-colour standard (MASSIVE Photonics). S^3^M is also compatible with other SMLM modalities, *e*.*g*. dSTORM imaging, see SI note S13, and DNA-PAINT with spectrally overlapping dyes, see SI note S15.

**Fig 5.**
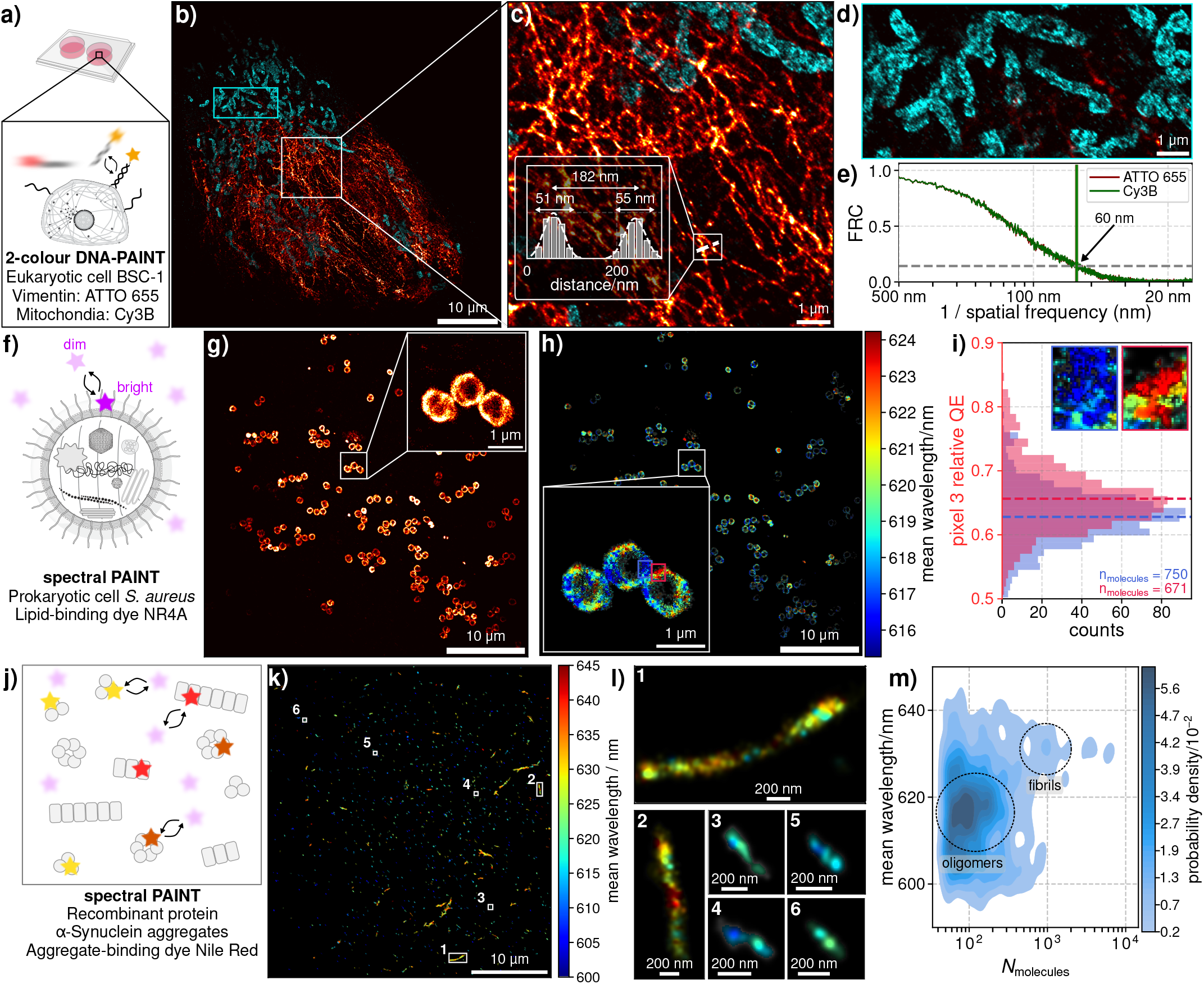
Colour differentiation below the diffraction limit. **a)** Schematic of the 2-colour DNA-PAINT experiment. **b)** Two-colour DNA-PAINT image of ATTO 655 labelled vimentin (orange) and Cy3B labelled mitochondria (cyan) in a BSC-1 cell. **c)** Zoom-in of cell, line plot demonstrating resolution of individual vimentin filaments below the diffraction limit. **d)** Zoom-in of mitochondria from **b). e)** FRC of ATTO 655 (red) and Cy3B (green) showing a resolution of 60 nm for both. **f)** Schematic of the spectral PAINT imaging of *Staphylococcus aureus*. **g)** PAINT images of the lipid bilayer of *Staphylococcus aureus* using Nile Red 4A. Inset shows a zoom-in of exemplar cells. **h)** Spectrally resolved PAINT image of the cells shown in **g. i)** Histogram of the quantum efficiency of pixel 3 for the red and blue areas highlighted in **g)**. Insets show zoom-ins of those areas. **j)** Schematic of the spectral PAINT imaging of *α*-Synuclein aggregates using Nile Red. **k)** Spectrally resolved PAINT image of *α*-Synuclein aggregates. **l)** Zoom-ins of individual aggregates, showing long hydrophilic (longer wavelength) fibrils and smaller more hydrophobic (shorter wavelength) oligomers. **m)** Analysis of the dataset highlights the aggregates with higher numbers of localisations have a greater mean wavelength. Clusters defining oligomers and fibrils are highlighted that correspond to wavelength regions found previously.(45)

By lowering the technical barriers to multicolour, we hope S^3^M will enable these experiments to become routine, both in standard commercial samples and in highly multiplexed biological measurements. This offers a parallel approach to reducing experimental time, multiplexing using separable spectral fingerprints as channel numbers. To give a few examples of approaches that could benefit from this, FLASH-PAINT used DNA “eraser” strands to sequentially image 13 different biomolecules relating to the Golgi apparatus,(16) and SUM-PAINT used toehold-mediated strand displacement to sequentially image 30 different proteins in neurons.(17) These methods undeniably generate high-quality data of biological structures, but do so at the expense of time—Jungmann’s lab estimate that in order to generate 13-plex data, all of the state-of-the-art methods take 10+ hours.(47) Thus, enabling simultaneous, simple, detector-based multiplexing of these experiments will enable higher-throughput deep understanding of biological systems.

### Prokaryote-based Spectral PAINT

We have performed spectral PAINT (sPAINT)(45, 48) in prokaryotic cells (*Staphylococcus aureus*, Fig. 5f). In these experiments, we utilised the transient binding of the environmentally sensitive dye NR4A,(49) which has been shown to have a polarity and hydrophobicity sensitive emission peak. The emission peak shifts from ∼580 nm to ∼660 nm, dependent on the local environment.(49) Thus by observing single NR4A molecules as they bind transiently to the lipid bilayer of fixed *Staphylococcus aureus* using S^3^M, we both super-resolved these cells and gained insight into the local lipid environment. As Fig. 5g demonstrates, we are able to super-resolve membrane lipids in these bacteria, with an FRC-derived resolution of 109± 1 nm (Fig. S86). Using a forward model (see SI Note S17) we were also able to provide maps of the bacterial membrane’s nanoscopic environment (Fig.5h). The lipids in these bacteria appear relatively homogeneous, with very minor spectral shifts from pixel-to-pixel (the mean emission peaks at ∼619 nm with a standard deviation of 1 nm, Fig. S74). We do not find a correlation between the mean emission wavelength and the number of photons per analysis pixel (Fig. S74), and coupled with our forward model simulations (SI Note S17) this gives confidence in our ability to unpick small shifts in emission wavelength. Fig. 5i highlights this, showing that the molecules in the areas highlighted in Fig. 5h differ by very small amounts in their relative pixel 3 QE. Nonetheless, with the large amount of localisations we are able to easily acquire using S^3^M, unpicking these small variations in membrane environment becomes possible. This will enable future experiments probing the local environments of prokaryotes and eukaryotes using the increasing range of probes(49–51) whose spectra shift depending on their local environment.

### Spectral PAINT on Recombinant Protein Aggregates

We then used Nile Red based sPAINT to look at larger variations in the different aggregation states of a recombinant protein, *α*-Synuclein (Fig. 5j). sPAINT on *α*-Synuclein aggregates gives a multi-modal dataset: we observe a range of aggregate shapes, sizes, and local environments (Fig. 5k). As we have previously demonstrated,(45, 48) we find oligomeric *α*-Synuclein to give emission at shorter wavelengths than fibrillar, implying that it is more “hydrophobic”(52) (Fig. 5l, section S30). This agrees with previous work that found oligomers are more able to bind to and disrupt lipid membranes,(53) highlighting that S^3^M may, with its spectral fingerprinting ability, probe the local environment of single protein aggregates.

Thus despite the inherent loss of detected photons the Bayer pattern produces, we still achieve sufficient sensitivity to detect and localise Nile Red and NR4A, (median photon values per localisation of 870 and 1,740 photons, respectively— leading to median localisation precisions of 30 nm and 14 nm, and median spectral fingerprint precisions of ±13% and ±12%) and extract reliable spectral information (SI Note S17). This is particularly true when the same nanoscopic area is repeatedly bound to by a dye molecule, which enables more precise spectral information to be extracted. Overall, these data demonstrate clearly that S^3^M allows one access to multi-modal single-molecule spectroscopy datasets with a minimally complex optical footprint.

Taken together, the data in Fig. 5 serve to highlight the compatibility of S^3^M with SMLM, and its enabling potential for multichannel and complex SMLM experiments. Despite the inherent loss of sensitivity the Bayer pattern (or any spatial patterning of the pixels) produces, we can generate multi-colour SMLM datasets with a resolution comparable to the state-of-the-art, as well as more complex experiments where the SMLM probe’s emission wavelength encodes local environment information.

## Conclusions

Here, we have introduced S^3^M, an easy to implement optical method that uses spatially patterned detectors to enable multicolour microscopy and simplified single-molecule spectroscopy. The key conceptual advance is that detector patterning, despite the inevitable loss in photon efficiency, encodes additional spectral information that can be extracted directly from raw single-molecule images during localisation fitting, without optical splitting or demosaicing. This is applied to widely available Bayer-patterned detectors, but the principle of S^3^M is applicable to any detector with spectrally distinct pixels, in any pattern. We show that S^3^M is sufficiently sensitive to detect single dye molecules reliably, enables straightforward multiplexing and single-molecule FRET, and is compatible with spectrally multiplexed localisation microscopy on par with other cutting-edge spectral-sensing methods with minimal experimental complexity (see Table S1) for comparisons. More broadly, S^3^M establishes a practical trade-off of detector area for spectral information. For full details of this trade-off, see SI Section S10. S^3^M can be readily incorporated into routine single-molecule applications that require spectral sensing such as DNA sequencing.(54) By combining conceptual simplicity with widely available hardware, we hope this approach will make experiments such as imaging-based single-molecule FRET, sPAINT, and multicolour super-resolution microscopy routine. Further improvements in the quality of spatially patterned detectors are expected to accelerate the ability of S^3^M further.

## ACKNOWLEDGEMENTS

James Manton (Laboratory of Molecular Biology, Cambridge) and Arnulf Rosspeintner (Departement de Chimie Physique, Université de Genéve) are both warmly acknowledged for incisive discussions. Rohan Ranasinghe (Yusuf Hamied Department of Chemistry, Cambridge) is thanked for useful advice on the purchase of DNA oligonucleotides. Gonzalo Angulo (Instytut Chemii Fizycznej, Polskiej Akademii Nauk, Warszawa) is thanked for stimulating discussions on the topic of single-molecule FRET. Andrey Klymchenko (Université de Strasbourg) is thanked for the gift of the NR4A. J.S.B. and S.F.L. acknowledge funding from the University of Cambridge Technology Investment Fund, and the Isaac Newton Trust. C.L.J. and S.Y. acknowledge funding from the Centre for Doctoral Training in Connected Electronic and Photonic Systems (CEPS) (EP/S022139/1).

## AUTHOR CONTRIBUTIONS

J.S.B. co-conceptualised the project, performed imaging experiments, wrote all code relating to the work, co-supervised the project, acquired funding, and wrote the manuscript. B.C. performed imaging experiments and analysis, and wrote the manuscript. D.F.H. performed mammalian cell culture, prepared supported lipid bilayers, prepared samples for imaging and assisted in imaging for the dSTORM experiment. S.K. performed all bacterial cell work relating to the *S. aureus* experiments. C.L.J. performed calculations and initial experiments relating to the spatial patterned detector concept. S.Y. provided technical assistance in the setup of the ZWO camera. R.C.G. purified and provided the *α*-Synuclein monomer. R.A.F., M.V., D.K., V.L., and P.K.S. provided supervision and acquired funding. Y.Z., A.M.S. and S.J.D. provided the proteins for the supported lipid bilayer experiment. S.F.L. co-conceptualised the project, acquired funding, co-supervised the project, and wrote the manuscript.

## CODE AVAILABILITY

Code relating to this work is available at github.com/JBeckwith/pyS3M for academic use.

## DATA AVAILABILITY

Data are available at the EBI (10.6019/S-BIAD3210).

## Methods

### Optical Setup

Experiments were performed on a wide-field fluorescence microscope (Eclipse Ti2, Nikon). Five lasers (Cobolt C-FLEX combiner with 405-, 488-, 515-, 561- and two 638-nm lasers, free space) are expanded, spatially filtered, combined with dichroics, and circularly polarised with a quarter-wave plate, then focused using an achromatic doublet lens (250mm, PAC067A, Newport) to a spot in the back focal plane of an oil-immersion objective (Plan Apo, 100× 1.49 NA oil, Nikon). A schematic of the setup is shown in Fig. S34. The fluorescence passes through dichroics and optical filters, and is demagnified 2 ×(2.5 ×) by a 4f system of a 100 mm and 50 mm (40 mm) achromatic lenses (AC254-050-A-ML, AC254-100-A-ML, Thorlabs; APAC17, Newport) before being focused onto the Ximea MC050CG-SY (ZWO ASI 585MC), resulting in a total system magnification of 50× (40×) and a virtual pixel size of 69 nm ×69 nm (72 nm× 72 nm). The dichroics and optical filters for each experiment are listed in S6, S7, and S8.

### Simulation of the S^3^M pipeline

Simulations were performed to characterise the localisation and colour precision of the Bayer-SMLM pipeline as a function of experimental parameters. Unless stated otherwise, all simulations used camera calibration values drawn from a Ximea MC050CG-SY sCMOS sensor (camera characterisation details in table S2), a NA of 1.49, a pixel size of 69 nm, a uniform background of 5 photons per pixel, and a Gaussian pre-smoothing kernel with *σ* = 1.5 pixels applied before fitting. For each condition, a two-dimensional Gaussian point-spread function (for full details of image simulation, see S1) was placed at a position drawn uniformly within the central pixel of a simulated sensor patch and fitted by Levenberg–Marquardt optimisation (for full details of fitting see S3); the process was repeated 20,000 times (bootstraps) to obtain statistical estimates of spatial precision (*σ*_xy_, the root-mean-square localisation error averaged over x and y) and colour precision (*σ*_colour_, the standard deviation of the Euclidean distance in RGB colour space from the known spectral fingerprint of the dye).

To compare fit yield depending on read noise and peak pixel QE, the values of read noise and peak pixel QE (assuming normalised spectra shown in Fig. 1a) were scanned in logarithmic space over 300 read levels between 0.1–10 RMS e^*−*^, and over 60 pixel QE levels in linear space between 0 and 1. An ATTO 488, ATTO 565 or ATTO 647N molecule emitting 1,000 photons was simulated on a 12× 12 pixel Bayer grid. The fraction of bootstraps returning a converged, physically plausible fit was reported as the fit yield and is plotted in Fig. 1c.

### Biotin-BSA/Neutravidin Surface Preparation

Glass coverslips (VWR Collection, 631-0124) were cleaned with argon plasma for 60 minutes (Expanded Plasma Cleaner, PDC-002, Harrick Plasma). An imaging chamber was created on the coverslips using Frame-Seal slide chambers (9 × 9 mm^2^, SLF0201, Bio-Rad). Glass coverslips were then washed three times with filtered HEPES buffer (0.02 µm syringe filter, Whatman, 6809-1102). Washed coverslips were then incubated in 1 mg/ml biotinylated-BSA in PBS (9048-45-8, Sigma-Aldrich; GATTAquant) for 5 minutes before removing the excess biotinylated-BSA solution and washing three times with filtered HEPES. Biotinylated-BSA coated coverslips were then incubated in 1 mg/ml Neutravidin in PBS (31000, Thermo Scientific; GATTAquant) for 5 minutes before removing the excess Neutravidin solution and washing three times with filtered HEPES, leaving the final biotinylated-BSA/neutravidin surface.

### Biotinylated-Dye Experiments

Biotinylated-dyes were diluted to a concentration of 5–50 pM in a filtered solution of 100 mM NaCl 20 mM HEPES before incubation on the biotinylated-BSA/neutravidin coated coverglass for 10 minutes. Excess dye solution was removed before washing four times with filtered 100 mM NaCl 20 mM HEPES solution. The imaging chamber was then filled with an imaging solution consisting of 100 mM NaCl, 20 mM HEPES, 3 mM Trolox (53188-07-1, Sigma-Aldrich),(55) and either 30 mM Sodium Sulphite or 2.5 mM Protocatechuic acid (99-50-3, Sigma-Aldrich) and 0.25 U/mL Protocatechuate-3,4-dioxygenase (9029-47-4, MP Biomedicals),(56) before sealing the chamber with an argon plasma cleaned glass slide and clear nail polish. After waiting 10 minutes for the nail polish to dry, samples were imaged immediately with TIRF illumination.

### Quantum Dot Experiments

Quantum Dots were diluted to a concentration of 20 pM–1 nM in a filtered solution of 100 mM NaCl 20 mM HEPES before incubation on the biotinylated-BSA/neutravidin coated coverglass for 10 minutes. Excess Quantum Dot solution was removed before washing four times with filtered 100 mM NaCl 20 mM HEPES solution. The imaging chamber was then filled with an imaging solution consisting of 100 mM NaCl, 20 mM HEPES, 3 mM Trolox, and 30 mM Sodium Sulphite,(55) before sealing the chamber with an argon plasma cleaned glass slide and clear nail polish. After waiting 10 minutes for the nail polish to dry, samples were imaged immediately with TIRF illumination.

### Holliday Junction Experiments

The DNA sequences that make up Holliday Junction 7 (hereafter referred to as J7) from McKinney *et al*.(40) were purchased from ATDBio, purified by high-performance liquid chromatography. The specific DNA sequences are listed in Supplementary Table S5. Holliday Junction DNA was annealed by mixing the four strands at ~100 µM concentration in 20 mM HEPES pH 8 with ∼100 mM NaCl. DNA samples were stored at -20°C before use.

Holliday Junctions were diluted to a concentration of 5– 20 pM in filtered solution of 20 mM HEPES before incubation on the biotinylated-BSA/neutravidin coated coverglass for 10 minutes. Excess Holliday Junction solution was removed before washing two times with filtered 20 mM HEPES solution, followed by two more washes with imaging buffer (50 mM MgCl_2_, 20 mM HEPES, 3 mM Trolox, 2.5 mM Protocatechuic acid, and 0.25 U/mL Protocatechuate-3,4-dioxygenase). The imaging chamber was then filled with imaging buffer and sealed with an argon plasma cleaned glass slide and clear nail polish. After waiting 10 minutes for the nail polish to dry, samples were imaged immediately with TIRF illumination.

### Immobilised TetraSpeck Bead Imaging

0.1 µm TetraSpeck Beads (TetraSpeck™ Microspheres, 0.1 µm, fluorescent blue/green/orange/dark red, Invitrogen, T7279) were diluted to single-particle concentrations in an aqueous solution of 1% polyvinyl alcohol (average mol. wt. 85,000– 124,000, 87-89% hydrolysed, Sigma-Aldrich, 363081). The sample was then spun-coat at 3,000 rpm for 45 s onto an argon plasma cleaned coverslip, and sealed with an argon plasma cleaned glass slide and clear nail polish. For both the glass slides and coverslips, the argon plasma cleaning was performed for 1 h. After waiting 10 minutes for the nail polish to dry, samples were imaged immediately with TIRF illumination.

### Imaging of Diffusion on Supported Lipid Bilayers

Supported Lipid bilayers, consisting of 98 mol % POPC (850457C-25mg, Avanti Polar Lipids) and 2 mol % DGS-NTA(Ni) (790404C-5 mg, Avanti Polar Lipids) were prepared by vesicle fusion as described previously.(57) Double Hexahistidine tagged ICAM-1, CD58 and UCHT1-Fab-Halotag were labelled with Alexa Fluor 488, Alexa Fluor 555 and Janelia Fluor 646-Halotag ligand respectively. These proteins were diluted to a concentration of 2 nM, added to the bilayer (to a total volume of 10 µL) and incubated at room temperature for 1 hour. The bilayer was then washed 10 times with 5 µL of PBS and imaged immediately with TIRF illumination.

### PAINT imaging of *Staphylococcus aureus*

#### Sample Preparation

A single colony of *Staphylococcus aureus* JE2 grown on tryptic soy agar was used to inoculate 5 mL of tryptic soy broth. Cultures were grown for 16 h at 37 ° C with shaking at 700 rpm to stationary phase. Bacteria were harvested by centrifugation (6,000 g for 3 min), washed once in sterile PBS (0.22 µm filtered), and resuspended in 3 % paraformaldehyde (PFA) in PBS for 15 min at room temperature. Bacteria were subsequently washed three times with PBS by centrifugation and resuspension. Samples were stored at 4 ° C until imaging.

#### Imaging

Glass coverslips were cleaned with argon plasma for 60 minutes. An imaging chamber was created on the coverslips using Frame-Seal slide chambers. Glass coverslips were then washed three times with filtered PBS buffer. Washed coverslips were then incubated in a 0.02 µm filtered 0.01% solution of poly-L-lysine (PLL) (mol. wt. 150,000– 300,000, Sigma-Aldrich, P4832) for 20 minutes before removing excess solution and washing three times with PBS. A 1 in 10 dilution of the *Staphylococcus aureus* colony was prepared in PBS and deposited onto the PLL-coated coverslips for 20 minutes, before removing excess solution and washing three times with filtered PBS. An imaging buffer containing 100 mM NaCl, 20 mM HEPES, 3 mM Trolox, 30 mM Sodium Sulphite, and 500 pM Nile Red 4A(49) was added to the imaging chamber and immediately imaged with HILO illumination.

### Two-colour PAINT imaging of BSC-1 cells

Single-domain antibody labelled BSC-1 cells, along with DNA imagers, were purchased from MASSIVE-Photonics (MASSIVE-Cells 2-PLEX). 300 µL of DNA imager solution was added to the slide (250 pM ATTO 565, 750 pM Cy3B) and the cells were immediately imaged with HILO illumination.

### PAINT imaging of *α*-Synuclein aggregates

*α*-Synuclein aggregates, generated in accordance with Horne *et al*.(58) were prepared as described in Bruggeman *et al*.(59) Glass coverslips were cleaned with argon plasma for 60 minutes. An imaging chamber was created on the coverslips using Frame-Seal slide chambers. Glass coverslips were then washed three times with filtered PBS buffer. Washed coverslips were then incubated in a 0.02 µm filtered 0.01% solution of poly-L-lysine (PLL) (mol. wt. 150,000–300,000, Sigma-Aldrich, P4832) for 20 minutes before removing excess solution and washing three times with PBS. A solution of 10 µM *α*-Synuclein aggregates (by monomer concentration) and 2.5% v/v 60 nm gold nanoparticles was prepared in PBS and deposited onto the PLL-coated coverslips for 20 minutes, before removing excess solution and washing three times with filtered PBS. An imaging buffer containing 100 mM NaCl, 20 mM HEPES, 3 mM Trolox, 30 mM Sodium Sulphite, and 1 nM Nile Red was added to the imaging chamber and immediately imaged with HILO illumination.

### Data Analysis

#### Single-molecule localisation and drift correction

Raw image frames were converted to photoelectron images by subtracting the per-pixel offset map and dividing by the gain map, both obtained from camera calibration (see S18). Candidate single-molecule puncta were identified as discussed in SI Note S2, implemented following Hekrdla *et al*.(60) Each candidate was extracted as a 12 ×12 pixel region of interest from the raw Bayer image, and fitted by Levenberg– Marquardt weighted least squares to a multi-channel Gaussian point-spread function model in which the background and per-channel amplitudes (*A*_R_, *A*_G_, *A*_B_) enter as their square roots to enforce positivity. For full details of this see SI Note S3. Localisations were retained if the reduced chi-squared 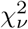 was below 3, the fitted PSF width *σ*_*xy*_ lay within 75–160 nm, the per-axis localisation uncertainty *δx, δy <* 1 pixel, and the total detected photon count exceeded 100 photons.

Drift was corrected using the Adaptive Intersection Maximisation (AIM) algorithm(61) The temporal segmentation was chosen to match the localisation density of each dataset (typically 10–50 frames). The intersection distance and search-region radius were fixed at 20 nm and 60 nm respectively across all datasets. For super-resolution experiments (*i*.*e*. all data presented in Fig. 5), repeated detections of the same blinking emitter were linked across frames: localisations separated by less than the mean per-axis localisation precision in the image plane and by no more than two consecutive dark frames were consolidated into a single event, with photon counts summed and spectral fractions, PSF widths, and 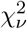 averaged over contributing frames; events whose first or last frame coincided with the start or end of the acquisition were discarded as their on-times were indeterminate. For single-molecule experiments (*i*.*e*. data presented in Fig. 2, Fig. 3a–e), localisations from the same physical emitter were grouped into single-molecule identities by HDBSCAN(62) applied to the two-dimensional localisation coordinates, with a minimum cluster size of 10 localisations; localisations assigned noise labels (− 1) were discarded, and photon-weighted means of the position, relative pixel QE fractions, and PSF width were taken as the representative single-molecule observables.

#### Mixture Identification

Single chromophores were identified in mixtures by their spectral fingerprint using a Gaussian mixture model (GMM). The misidentification rate of the two extracted Gaussian distributions is then estimated. First, the GMM determines the means, variances, and weights of the distributions. Next, a synthetic data set using those same parameters are generated with known identities. Bayes’ decision rule is then used to assign identities to the synthetic data, and the true positive rate and misidentification rate are generated. For the photon-dependent rate of misidentification, the spectral fingerprint is recorded as a function of cumulative detected photons. Using the high cumulative photon data, the means of the two Gaussians are determined *via* a GMM. These means are then used for fitting the low cumulative photon data, where the populations and standard deviations of the underlying Gaussians change.

#### Quantum dot identification

The six QDs were identified using a GMM. The means and covariances for the mixture were generated by measuring the spectral print of each QD in separate experiments under identical imaging conditions to the mixture experiment. This method is benchmarked in S24.

#### Single-particle tracking and diffusion analysis

Raw image frames were converted to photoelectron images by the same camera-calibration procedure described above. Candidate single-molecule puncta were identified by the same manner as the analysis discussed above, and each candidate was extracted as a 20× 20 pixel ROI from the raw image. Because continuous illumination induces motion blur that elongates the effective PSF in the direction of molecular diffusion, localisation was performed with a rotated, elliptical Gaussian model rather than the symmetric model used for static emitters. The eleven-parameter model adds an independent minor-axis width *σ*_minor_ and an in-plane rotation angle *θ* relative to the isotropic model, allowing the fit to absorb anisotropy introduced by sub-frame motion. Fitting proceeded in the same two-stage Levenberg-Marquardt weighted least-squares procedure described above. Localisations were retained if 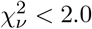, the per-axis localisation uncertainty *δx, δy <* 1.5 pixels, the per-channel colour-fraction uncertainty was below 0.15, and the total detected photon count exceeded 100 photons.

Prior to trajectory linking, stationary emitters were identified and removed. DBSCAN(63) was applied to the two-dimensional localisation positions (*x*_*c*_, *y*_*c*_) across all frames using a neighbourhood radius 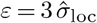, where 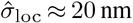 is the per-axis localisation precision, and a minimum cluster size of ten observations. Localisations that belonged to a dense spatial cluster—indicative of a molecule immobilised on the coverslip surface—were discarded before linking.

Validated localisations were linked into single-particle trajectories using a spectral-assisted Linear Assignment Problem (LAP) framework following Jaqaman *et al*.(64) The cost of linking localisation *i* in frame *t* to localisation *j* in frame *t* +Δ*t* was

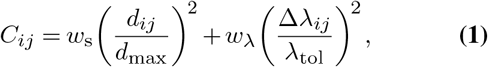

where *d*_*ij*_ is the Euclidean separation, *d*_max_ = 690 nm is the maximum permitted linking distance, Δ*λ*_*ij*_ is the Euclidean distance in normalised spectral-fraction space (*A*_R_, *A*_G_, *A*_B_), and *λ*_tol_ = 0.25 is the spectral tolerance. Spatial and spectral weights were set to *w*_s_ = *w*_*λ*_ = 1. To prevent systematic underestimation of the diffusion coefficient that arises from applying a fixed spatial cutoff across variable-length dark periods, the maximum permitted linking distance was scaled with the elapsed gap *g* (in frames) according to

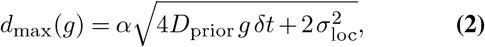

where *D*_prior_ = 0.1 µm^2^ s^*−*1^ is a prior estimate of the diffusion coefficient, *δt* is the camera frame interval, *σ*_loc_ is the per-axis localisation precision, and *α* = 4 covers more than 99 % of Brownian displacements in two dimensions. Gap-closing was permitted for up to five consecutive dark frames. Trajectories containing fewer than ten localisations were discarded.

Molecular species were assigned by *k*-means clustering (*k* = 3) applied to the time-averaged, normalised spectral fractions ⟨ *A*_R_⟩, ⟨ *A*_G_⟩, ⟨ *A*_B_⟩ of each trajectory. Clusters were ordered by increasing mean *A*_R_ to yield a consistent spectral ordering across acquisitions.

Diffusion coefficients were estimated from the mean-squared displacement (MSD) of each trajectory. The MSD as a function of lag time was computed analytically *via* the Fast Fourier Transform(37) and fitted using the covariance-based *Dσ*^2^ ordinary-least-squares estimator with motion-blur correction parameter 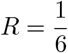, appropriate for uniform exposure throughout the acquisition frame.(37) The camera frame time was Δ*t* = 0.05 s and the pixel size was 69 nm. Molecules whose estimated diffusion coefficient lay outside the range 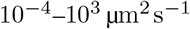 were excluded from further analysis.

#### FRET single-molecule analysis

FRET data were processed using a dedicated pipeline that separates spot detection, photobleaching identification, and spectral fitting. Candidate puncta were identified on the variance-aware demosaiced sum of the first 50 frames of each acquisition; summing increases the signal-to-noise ratio for detection whilst the variance map is scaled accordingly 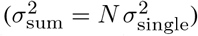, preserving the statistical validity of the puncta detection pipeline (SI Note S2). A 12 × 12 pixel ROI was extracted for each detected punctum.

Before fitting the PSF in every frame, the photobleaching transition was located using the PELT change-point algorithm(65) applied to the one-dimensional total-intensity trace (sum of photoelectrons within the ROI) across the full acquisition. The noise level 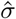 was estimated from the last 100 frames of each trace, which correspond to the post-bleach background; the penalty was set to 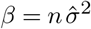, where *n* is the trace length, and a minimum segment length of 5 frames was imposed. Puncta for which no change point was detected (i.e. no bleaching step) were discarded. These puncta were then fit independently at every frame from the start of the acquisition up to the identified bleaching transition, yielding per-frame position, PSF width, photon count, and relative pixel QEs (*A*_R_, *A*_G_, *A*_B_).

Localisations in which all three relative pixel QEs lay within 0.01 of the fitter’s initial value of 1/3 were subsequently discarded as unconverged fits. The FRET ratio at each frame was computed as *A*_R_*/A*_G_; no additional spectral correction was applied beyond the per-pixel quantum efficiency weighting implicit in the fitting model.

FRET state transitions were then identified by a second application of PELT, now operating jointly on the two-dimensional signal [*A*_R_(*t*), *A*_G_(*t*)]. Joint detection exploits the constraint *A*_R_ + *A*_G_ + *A*_B_ = 1, under which a FRET transition shifts *A*_R_ and *A*_G_ in anti-correlated directions; a coherent change in both channels is detected with greater sensitivity than either channel alone.(66) An *l*_2_ cost function was used. The penalty was set to 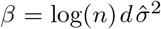, where *n* is the number of pre-bleach frames, *d* = 2 is the signal dimension, and 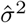 is the mean per-channel variance of the trace. This Bayesian Information Criterion (BIC) scaling is necessary because spectral fractions are bounded on [0, 1] and their variance is orders of magnitude smaller than that of photon-count traces; applying an unscaled penalty would suppress all but the largest transitions. A minimum segment length of 25 frames was imposed. Puncta for which no FRET transition was detected were excluded from further analysis; all change-point searches were parallelised across puncta. After changepoint analysis, the mean relative pixel QE between each changepoint is determined. These values are used to calculate an apparent FRET efficiency based on the photophysical properties of the donor and acceptor (See S25). These FRET efficiencies are sorted into two populations via a Gaussian mixture model to identify dwell times in the high-FRET and low-FRET states. A histogram of the dwell times is fit to a monoexponential decay to extract the rate of transition as done in previous kinetic assessment of Holliday Junctions.(40)

#### Analysis of NR4A and Nile Red Data

Localisation data, post single-molecule localisation and drift correction, were rendered into a super-resolved image (8 × oversampling, effective pixel size *≈*8.6 nm) using Gaussian blurring, and aggregates were segmented by percentile (in the case of *α*-Synuclein aggregates) or Li (in the case of *S. aureus*) intensity thresholding of the rendered image. This was followed by connected-component labelling. Segments smaller than 0.01 µm^2^ or containing fewer than 20 localisations were discarded. Fiducial markers were identified by their characteristic spectral fingerprint (low *A*_R_ *<* 0.45, high *A*_G_ *>* 0.45) combined with a high temporal density (present in *>*20% of acquisition frames) and removed prior to analysis.

The mean wavelength *µ* of Nile Red was extracted from each localisation’s relative pixel QEs and PSF width using a forward–inverse modelling approach (see Note S17). The single free parameter *µ* was recovered by minimising the weighted residuals

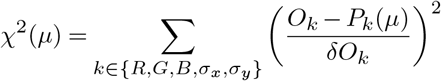

where, for each observable *k, O*_*k*_ is the measured value (the fitted Bayer relative pixel QEs *A*_pixel1_, *A*_pixel2_, *A*_pixel3_ and PSF widths *σ*_*x*_, *σ*_*y*_), *P*_*k*_(*µ*) is the value predicted by the forward model at central wavelength *µ*, and *δO*_*k*_ is the corresponding fit uncertainty returned by the PSF fitting step. This minimisation used the Trust Region Reflective algorithm with bounds *µ* ∈ [500, 750] nm. To obtain spatially resolved wavelength maps, localisations within each aggregate were first binned onto a regular 50 nm grid; each grid pixel containing at least 10 localisations was fitted independently. Grid pixels with too few localisations were assigned a cluster-level wavelength, obtained from a single fit to the photon-weighted average observables of all localisations in that cluster.

#### Imaging Conditions

Full imaging conditions, including dichroic mirrors used, emission filters used, camera frame rates, excitation wavelengths, and power densities at the sample plane are contained in Tables S6, S7, and S8.

