## Supplementary Information for "Spatially patterned, spectral single-molecule microscopy"

[1]Yusuf Hamied Department of Chemistry, University of Cambridge, Lensfield Road, Cambridge, CB2 1EW, United Kingdom
[2]Analytical Sciences, Biopharmaceutical Development, BioPharmaceuticals R&D, AstraZeneca, Cambridge, United Kingdom
[3]Victor Phillip Dahdaleh Heart & Lung Research Institute, University of Cambridge, Cambridge, United Kingdom
[4]Cambridge Centre for AI in Medicine, University of Cambridge, Cambridge, United Kingdom
[5]Cambridge Centre for Lung Infection, Royal Papworth Hospital, Cambridge, United Kingdom
[6]Department of Medicine, University of Cambridge, Cambridge, United Kingdom
[7]Cambridge University Hospitals Trust, Cambridge, United Kingdom
[8]Radcliffe Department of Medicine, John Radcliffe Hospital, University of Oxford, Oxford OX3 9DS, UK
[9]Medical Research Council Human Immunology Unit, John Radcliffe Hospital, University of Oxford, Oxford OX3 9DS, UK
[10]Centre for Misfolding Diseases, Yusuf Hamied Department of Chemistry, University of Cambridge, Cambridge, United Kingdom
[11]UK Dementia Research Institute, Cambridge, United Kingdom
[12]Present address: Department of Chemistry, University of Warwick, Coventry CV4 7AL, UK
[†]These authors contributed equally.

### Contents


**S1 Image Simulation** 4

**S2 Puncta Detection** 5

**S3 Puncta Fitting** 6

**S4 Fitting raw data *versus* Demosaicing** 9

**S5 Thorlabs CS505CU Single-Molecule Data** 15

**S6 ZWO ASI 585MC Single-Molecule Data** 17

**S7 Single-step photobleaching of single dyes** 18

**S8 Single-molecule spectral fingerprint *vs.* predicted value** 31

**S9 Effect of Background on $S^3M$** 32

**S10 Limitations of $S^3M$** 34

**S11 Effect of Motion Blur on Extraction of Spectral Signature** 41

**S12 Effect of Z Defocus on Extraction of Spectral Signature** 44

**S13 dSTORM on HeLa cells** 50

**S14 Raw Data and Fits of ATTO 655 and Cy3B DNA-PAINT on BSC-1** 51

**S15 ATTO 594 and Cy3B DNA-PAINT on BSC-1 Cells** 54

**S16 Alternative Pattern Simulations** 55

**S17 Nile Red/NR4A Forward Model** 59

**S18 Camera Calibration** 62

**S19 Timeline of commercially available Bayer detectors** 62

**S20 Additional *S. aureus* imaging** 64

**S21 DNA Sequences** 67

**S22 Calibrating Precision** 67

**S23 Quantifying Dye Pair Accuracy** 68

**S24 Quantum Dot Multiplexing** 70

**S25 FRET Efficiency Calculation** 71

**S26 Resolving Spectral Shifts** 72

**S27 Example Localisations of Holliday Junctions** 74

**S28 Example Localisations of $\alpha$-Synuclein** 75

**S29 Fourier Ring Correlation for *S. aureus*** 75

**S30 Nile Red and NR4A Characterisation** 77

**S31 Shift-Invariance** **78**

**S32 Zoom-Ins of Experimental PSFs** **87**

**S33 Imaging Conditions** **89**

**S34 Optical path for experiments** **92**

#### Supplementary Note S1: Image Simulation

Simulation of microscopy images follows the approach discussed in Appendix A of Fazel *et al.*(1) First, for each dye molecule being simulated in an image, a defined number of photons expected to impinge on the detector, $N_{\mathrm{photons}}$, was sampled from the observed emission spectrum. This observed emission spectrum is the dye emission spectrum multiplied by any spectral filters, dichroic mirrors, and wavelength-varying transmissive optical elements in the optical path before the detector. The photons were sampled from this effective emission spectrum, $\mathrm{Spectrum}_{\mathrm{eff}}$ by first converting it to a probability density function *via*

$$\mathrm{PDF}_{\mathrm{Spectrum}} = \mathrm{Spectrum}_{\mathrm{eff}} / \int_{\lambda_{\min}}^{\lambda_{\max}} \mathrm{Spectrum}_{\mathrm{eff}} \tag{S1}$$

where $\lambda_{\min}$ and $\lambda_{\max}$ are the wavelength limits of the camera QE. This $\mathrm{PDF}_{\mathrm{Spectrum}}$ was then converted into a normalised cumulative distribution function (CDF), from which wavelengths were randomly sampled *via* inverse transform sampling: uniform random variates on [0, 1] were mapped to wavelengths by interpolating the inverse of the CDF. With a specified number of $N_{\mathrm{photons}}$, we then calculated the mean wavelength impinging on our detector, $\bar{\lambda}$. This $\bar{\lambda}$ defines the width of our point spread function, $\sigma$, *via*

$$\sigma = \frac{\bar{\lambda}}{\sqrt{2} \cdot \pi \cdot \mathrm{NA}} \tag{S2}$$

where NA is the Numerical Aperture of the objective used (for calculations here, assumed 1.49). The pixellated PSF, $g(x, y)$, was then simulated at coordinates $(x_0, y_0)$ on a pixellated detector (pixels here simulated as 69×69 nm$^2$) using

$$g(x,y) = g_x(x) \cdot g_y(y) \tag{S3}$$

where

$$g_x(x) = \frac{1}{\sigma\sqrt{2\pi}} \cdot \exp\left(-\frac{1}{2}(\frac{x - x_0}{\sigma})^2\right) \tag{S4}$$

and

$$g_y(y) = \frac{1}{\sigma\sqrt{2\pi}} \cdot \exp\left(-\frac{1}{2}(\frac{y - y_0}{\sigma})^2\right). \tag{S5}$$

This pixellated PSF has a number of background photons per pixel, $b_{\mathrm{photons}}$ added,

$$g_{\mathrm{b}}(x,y) = g(x,y) + b_{\mathrm{photons}} \tag{S6}$$

and then the number of photons impinging on the detector per pixel, $N_{\mathrm{ph,n}}$ was generated by

$$N_{\mathrm{ph},n} = \mathrm{Poisson}(g_{\mathrm{b}}(x_n, y_n)). \tag{S7}$$

where $n$ indicates pixel index. These photons per pixel were then converted to photoelectrons per pixel ($N_{\mathrm{pe,n}}$) using the equation

$$N_{\mathrm{pe},n} = \mathrm{Binomial}(N_{\mathrm{ph},n}, \mathrm{QE}_{\mathrm{d},n}) \tag{S8}$$

where $\mathrm{QE}_{\mathrm{d},n}$ is the combined quantum efficiency of the pixel and the dye at pixel index $n$. This pixel-dependent quantum efficiency was calculated using

$$\mathrm{QE}_{\mathrm{d},n} = \int \mathrm{PDF}_{\mathrm{Spectrum}} \cdot \mathrm{QE}_n \mathrm{d}\lambda \tag{S9}$$

where, as above, $\mathrm{PDF}_{\mathrm{Spectrum}}$ is the normalised effective emission spectrum (*i.e.* $\int \mathrm{PDF}_{\mathrm{Spectrum}} \mathrm{d}\lambda = 1$) and $\mathrm{QE}_n$ is the Quantum Efficiency of the detector at pixel index $n$. The photoelectrons per pixel were then converted into the final image per pixel $w_n$ by

$$w_n = \mathrm{Normal}(\gamma_n N_{\mathrm{pe},n} + \mu_n, \sigma^2_{\mathrm{ro},n}) \tag{S10}$$

where $\gamma$ is the gain, $\mu$ is the offset, and $\sigma_{\mathrm{ro}}$ is the readout noise standard deviation. Given we simulate an image generated by a CMOS camera, each of these terms is noted as pixel-dependent by the subscript $n$. This is schematically shown in Fig. S1.

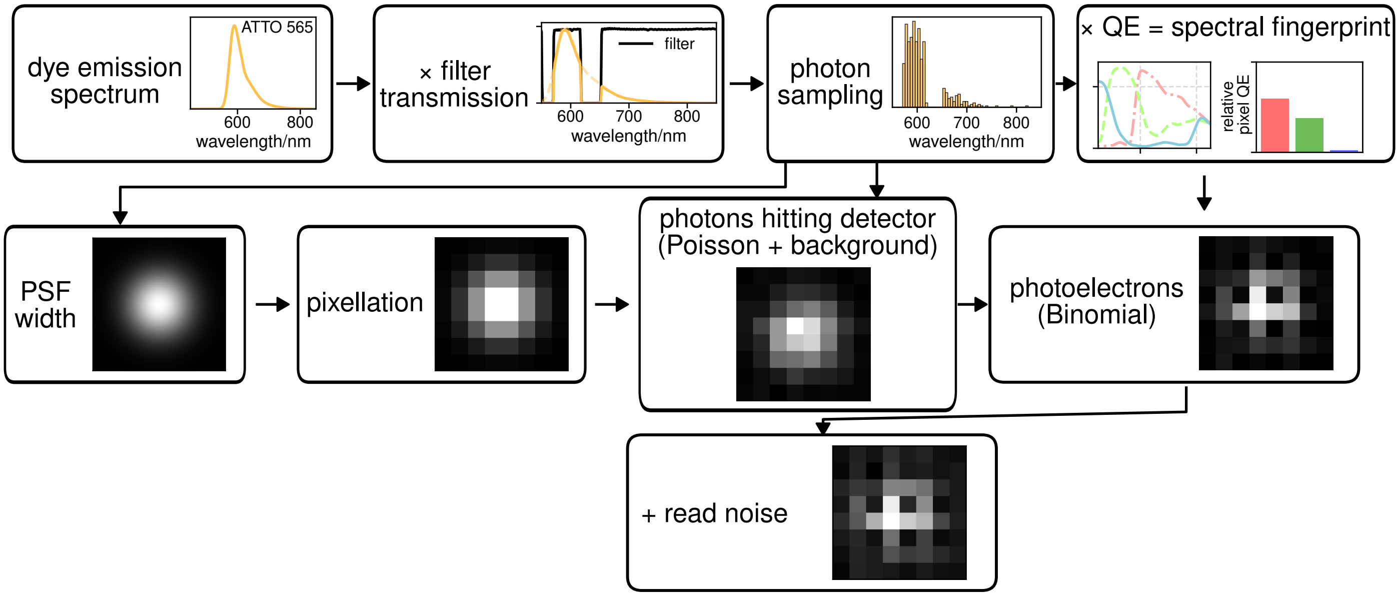


**Fig. S1. Flow diagram of the simulation pipeline**.

#### Supplementary Note S2: Puncta Detection

Before fitting 2D Gaussians to puncta locations, as described in Section S3, initial guess locations of puncta in images were detected using the approach of Hekrdla *et al.* (2) In brief, this first involved applying inverse square-root variance weighting to the raw data to produce a whitened image:

$$I_{\text{whitened}}(x,y) = I_{\text{raw}}(x,y) \cdot w \tag{S11}$$

where $w = 1/\sqrt{\text{Var}}$ is the inverse square root of the variance. The whitened image is then used for match filtering.(2) A schematic of this routine is shown in Fig. S2, and an example of this puncta detection routine on raw data is shown in Fig. S3.

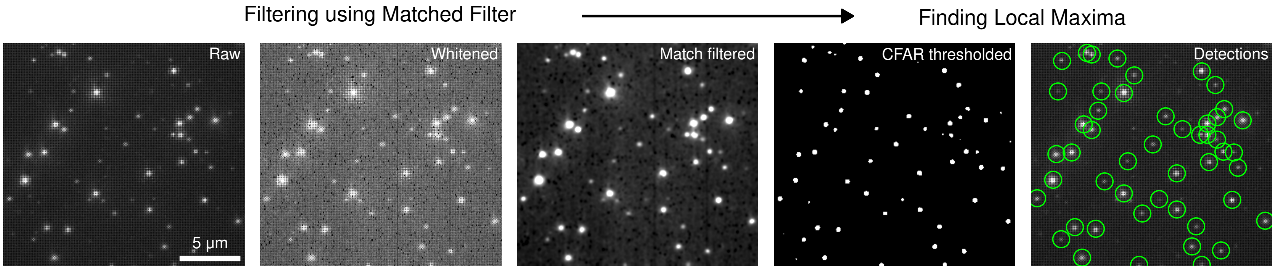


**Fig. S2. An example of the puncta detection routine, based on Fig. 1 of Hekrdla *et al.* (2)**

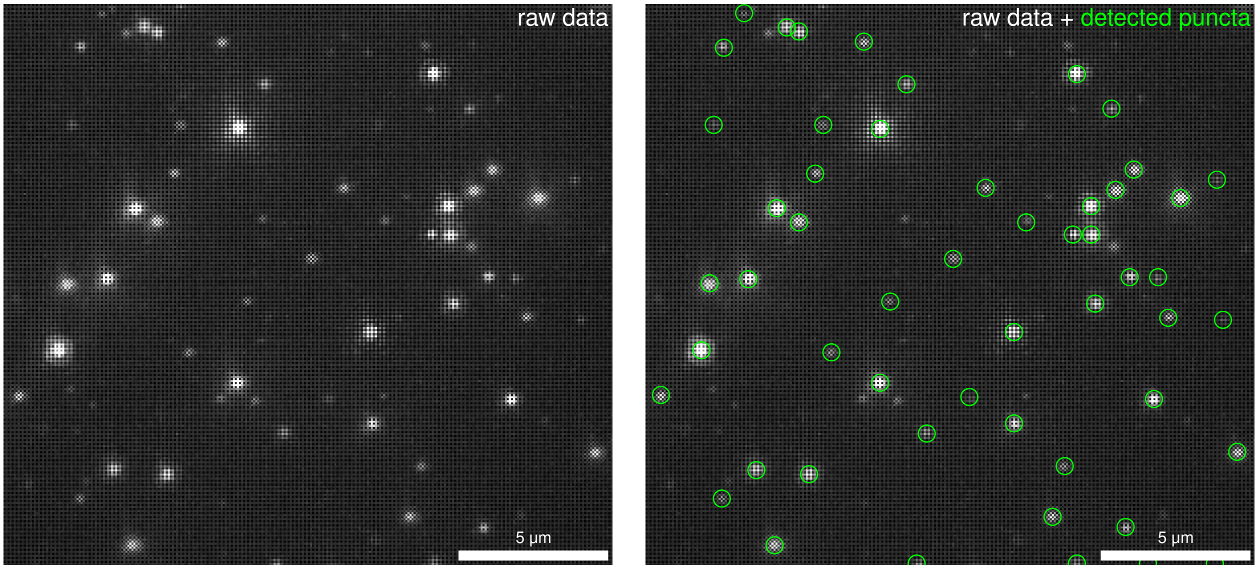


**Fig. S3. Example FOV and detected puncta**.

#### Supplementary Note S3: Puncta Fitting

Post puncta detection, these preliminary puncta locations are used to define N×N pixel boxes for fitting. Here, we used N=12, but such a parameter is tuneable dependent on pixel size. These areas are then used to compare a model to the data, where the model is

$$I_{\mathrm{model},x,y}(\boldsymbol{\theta}) = A_{p(x,y)} \exp\left[-\frac{(x-x_c)^2+(y-y_c)^2}{2\sigma^2}\right] + b_{p(x,y)} \tag{S12}$$

and where the parameter vector is

$$\boldsymbol{\theta} = \{x_c, y_c, \sigma, A_{\mathrm{pixel\,1}}, A_{\mathrm{pixel\,2}}, A_{\mathrm{pixel\,3}}, b^2_{\mathrm{pixel\,1}}, b^2_{\mathrm{pixel\,2}}, b^2_{\mathrm{pixel\,3}}\}$$

in the case of circular 2D Gaussian fitting, applied to the vast majority of datasets, and

$$\boldsymbol{\theta} = \{x_c, y_c, \sigma, \sigma_{\mathrm{minor}}, \theta_{\mathrm{rot}}, A_{\mathrm{pixel\,1}}, A_{\mathrm{pixel\,2}}, A_{\mathrm{pixel\,3}}, b^2_{\mathrm{pixel\,1}}, b^2_{\mathrm{pixel\,2}}, b^2_{\mathrm{pixel\,3}}\}$$

in the case of elliptical 2D Gaussian fitting, applied to the single-particle tracking datasets. Further,

$$p(x,y) \in \{\mathrm{pixel\,1}, \mathrm{pixel\,2}, \mathrm{pixel\,3}\}$$

maps each pixel coordinate to its corresponding pixel type. In the case of a Bayer detector, pixel 1 maps to blue pixels, pixel 2 to green, and pixel 3 to red. However, this notation is general and can be extended to the case where more pixel variants are present (Section S16) Here $A$ refers to the number of photons in a particular pixel type, $x_c$ and $y_c$ refer to the x and y centre positions of the PSF, $\sigma$ is the PSF width, and $b$ values are backgrounds per pixel type. These are squared in the fitting algorithm to avoid negative values perturbing the fit. The $\sigma_{\mathrm{minor}}$ and $\theta_{\mathrm{rot}}$ in the elliptical 2D Gaussian model refer to the width of the minor axis of the Gaussian, and its rotational angle respectively. This model is compared to the data by minimising the $\chi^2$ equation

$$\chi^2 = \sum_n w_n (d_n - I_{\mathrm{model}}(\boldsymbol{\theta}))^2 \tag{S13}$$

using the numpy leastsq algorithm(3) in our implementation. Here $d_n$ refers to data at pixel $n$, with the $\chi^2$ summed over all $n$ pixels. The fitting weights are given by, in accordance with Lin *et al.*,(4) first calculating the noise variance estimate of a pixel $\sigma_n^2$

$$\sigma_n^2 = \sigma_{\mathrm{ro},n}^2 + \max(f(d)_n, 0) + 1 \tag{S14}$$

where, as before, $\sigma^2_{\mathrm{ro},n}$ is the read noise of a single pixel, and $f(d)$ represents the image converted to photoelectrons and then smoothed by a 1.5 pixel radius Gaussian filter, which was found to achieve the most reasonable weighting. This is then converted to the weights factor by

$$w_n = \sigma_n^{-2}. \quad \textbf{(S15)}$$

The parameters $A_{\text{pixel}\,1}, A_{\text{pixel}\,2}$ and $A_{\text{pixel}\,3}$ are then summed to give the parameter of total photons detected, and then normalised by photons detected to give the relative pixel QE parameters used to determine the spectral fingerprint of an emitter. The entirety of the analysis routine is schematically depicted in Fig. S4, with optional postprocessing steps detailed in Fig. S5. Exemplar fits from raw data are shown in Fig. S6. Fits that reported fewer than 200 photons were excluded from further postprocessing in this work.

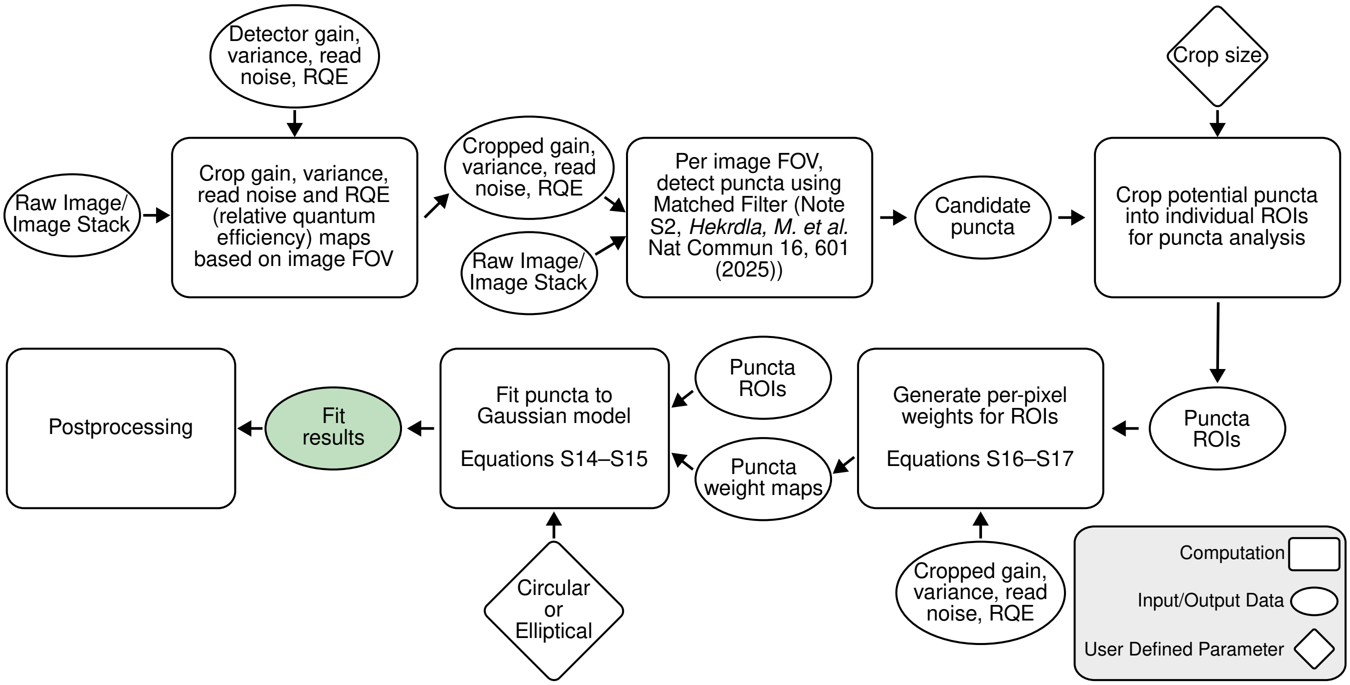


**Fig. S4. Flow diagram of the analysis pipeline.**

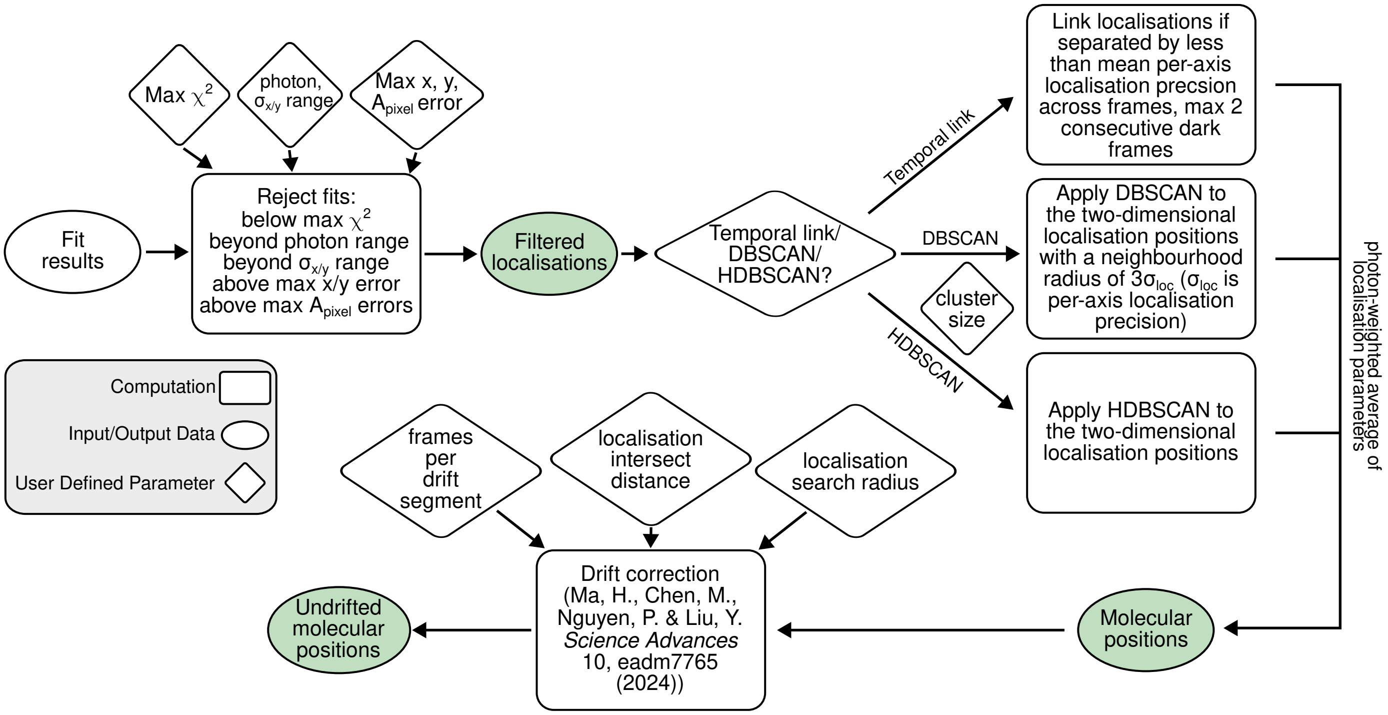


**Fig. S5. Flow diagram of the postprocessing pipeline.**

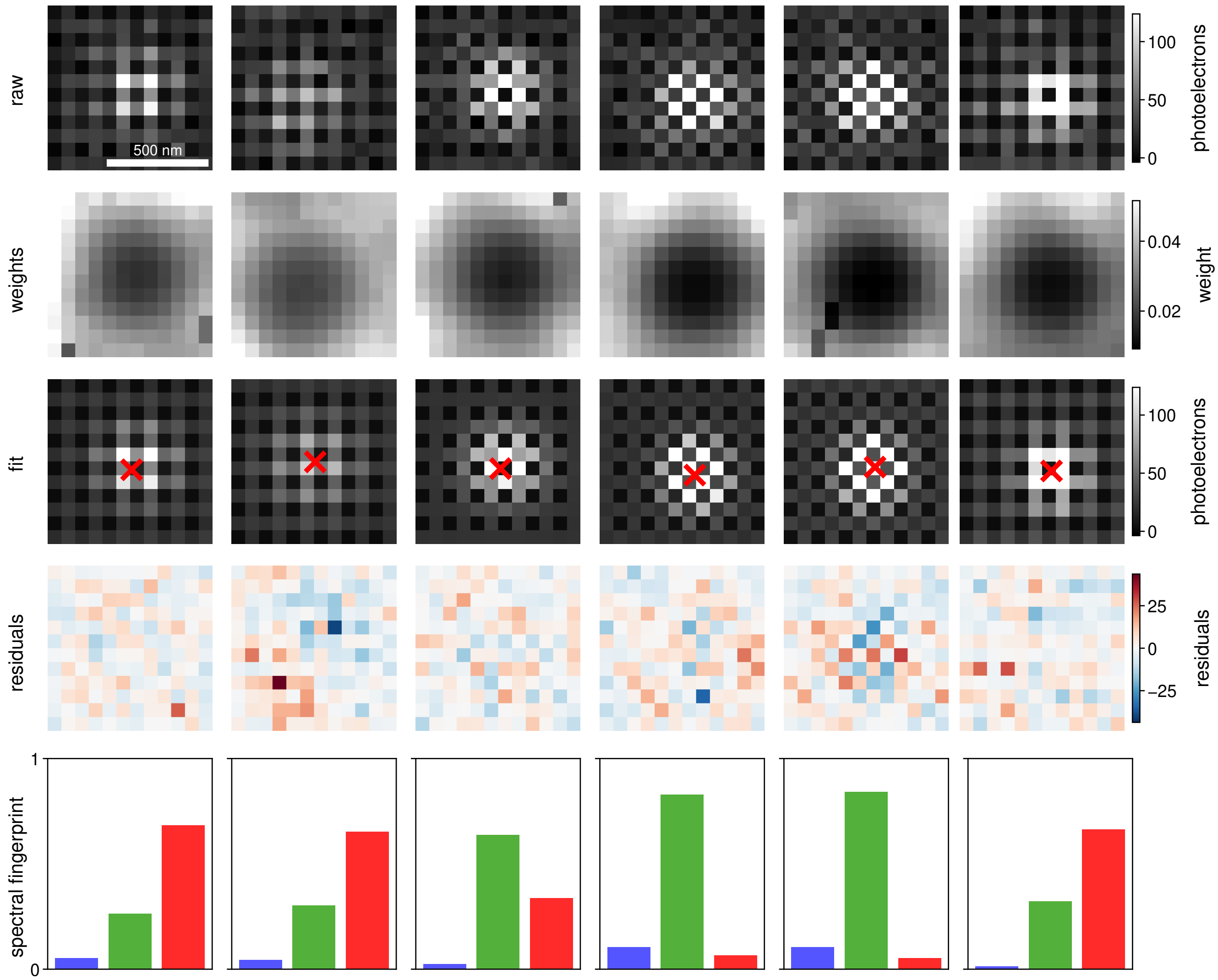


**Fig. S6. Exemplar fits for a range of Qdots (Fig. 3)d–g.**

#### Supplementary Note S4: Fitting raw data *versus* Demosaicing

Herein, we tested our strategy of directly fitting the image from a spatially-patterned camera versus that of demosaicing.(5) We tested three specific demosaicing algorithms: bilinear interpolation,(6) the demosaicing approach of Malvar *et al.*,(7) and the demosaicing approach of Menon *et al.*(8) We used the versions of these algorithms that are implemented in the Python *Colour* package.(9) An example of the direct fitting pipeline is shown in Fig. S7, and is explained in detail in section S3. Examples of the demosaicing analysis strategies with the three different demosaicing algorithms tested are shown in Fig. S8–S10. In detail, the demosaicing analysis strategy corresponds to comparing the three demosaiced puncta images to the model

$$I_{\text{model},x,y}(\boldsymbol{\theta}) = A \cdot \exp\left[-\frac{(x-x_c)^2+(y-y_c)^2}{2\sigma^2}\right] + b \tag{S16}$$

and where the parameter vector is

$$\boldsymbol{\theta} = \{x_c, y_c, \sigma, A, b^2\}$$

Here $A$ refers to the number of photons in an individual demosaiced image, $x_c$ and $y_c$ refer to the x and y centre positions of the PSF, $\sigma$ is the PSF width, and $b$ values are backgrounds per demosaiced image. These are squared in the fitting algorithm to avoid negative values perturbing the fit. This model is compared to the data of an individual demosaiced image by minimising the $\chi^2$ equation

$$\chi^2 = \sum_n w_n (d_n - I_{\text{model}}(\boldsymbol{\theta}))^2 \tag{S17}$$

using the numpy leastsq algorithm(3) in our implementation. Here $d_n$ refers to demosaiced data at pixel $n$, with the $\chi^2$ summed over all $n$ pixels. The fitting weights are given by, in accordance with Lin *et al.*,(4) first calculating the noise variance estimate of a pixel $\sigma_n^2$

$$\sigma_n^2 = \sigma_{\text{ro},n}^2 + \max(f(d)_n, 0) + 1 \tag{S18}$$

where, as before, $\sigma_{\text{ro},n}^2$ is the read noise of a single pixel, and $f(d)$ represents the demosaiced image converted to photoelectrons and then smoothed by a 1.5 pixel radius Gaussian filter, which was found to achieve the most reasonable weighting. This is then converted to the weights factor by

$$w_n = \sigma_n^{-2}. \tag{S19}$$

The $A$ parameters from the three separate demosaiced images are then summed to give the parameter of total photons detected, and then these separate $A$ values are further normalised by this number of total photons detected to give the relative pixel QE parameters used to determine the spectral fingerprint of an emitter.

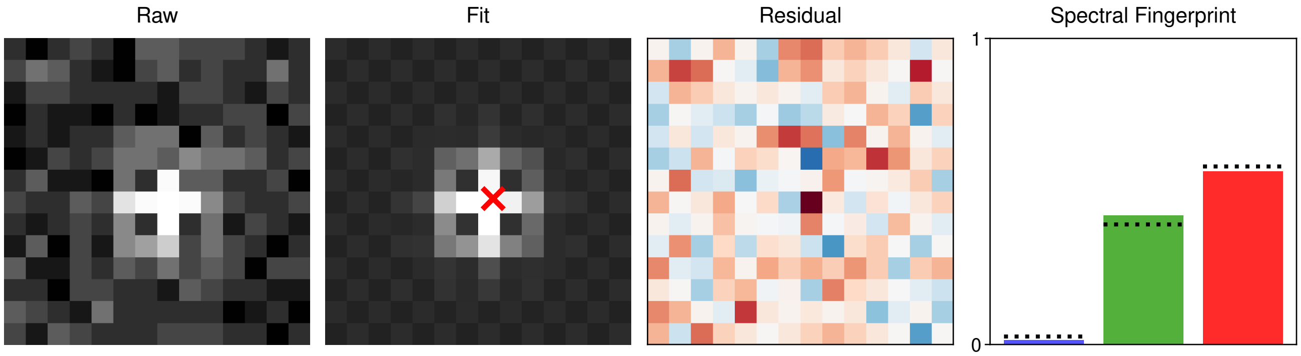


**Fig. S7. An example of the strategy of fitting the raw image directly on an ATTO 565 molecule (1000 photons, 10 background photons).**

We performed simulations using camera calibration values drawn from a Ximea MC050CG-SY sCMOS sensor (camera characterisation details in table S2), a NA of 1.49, a pixel size of 69 nm, a uniform background of 10 photons per pixel, and a Gaussian pre-smoothing kernel with $\sigma$ = 1.5 pixels applied before fitting.

Optical throughput was modelled using the measured spectral response of a Semrock dichroic mirror (Di03-R405/488/561/635) and notch filter (NF03-405/488/561/635E). Three dyes spanning the visible spectrum were evaluated—ATTO 488, ATTO 565, and ATTO 647N—and thus the input spectra for these simulations were these fluorophore spectra multiplied by the filter

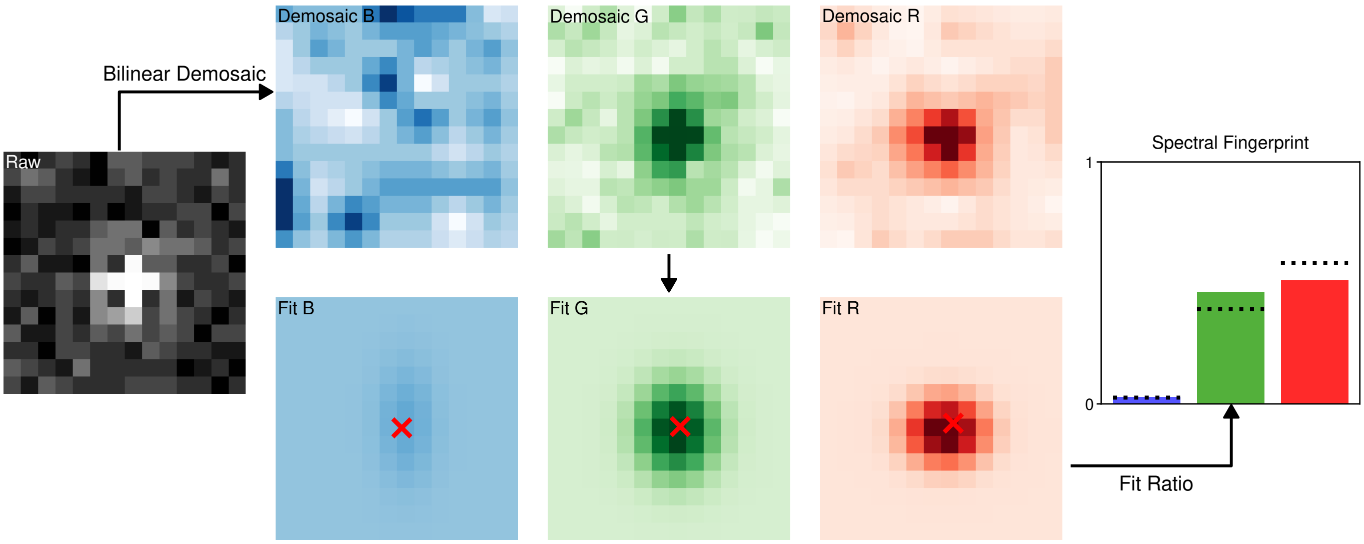


**Fig. S8.** **An example of the strategy of using bilinear demosaicing, and fitting the demosaiced images on an ATTO 565 molecule (1000 photons, 10 background photons).**

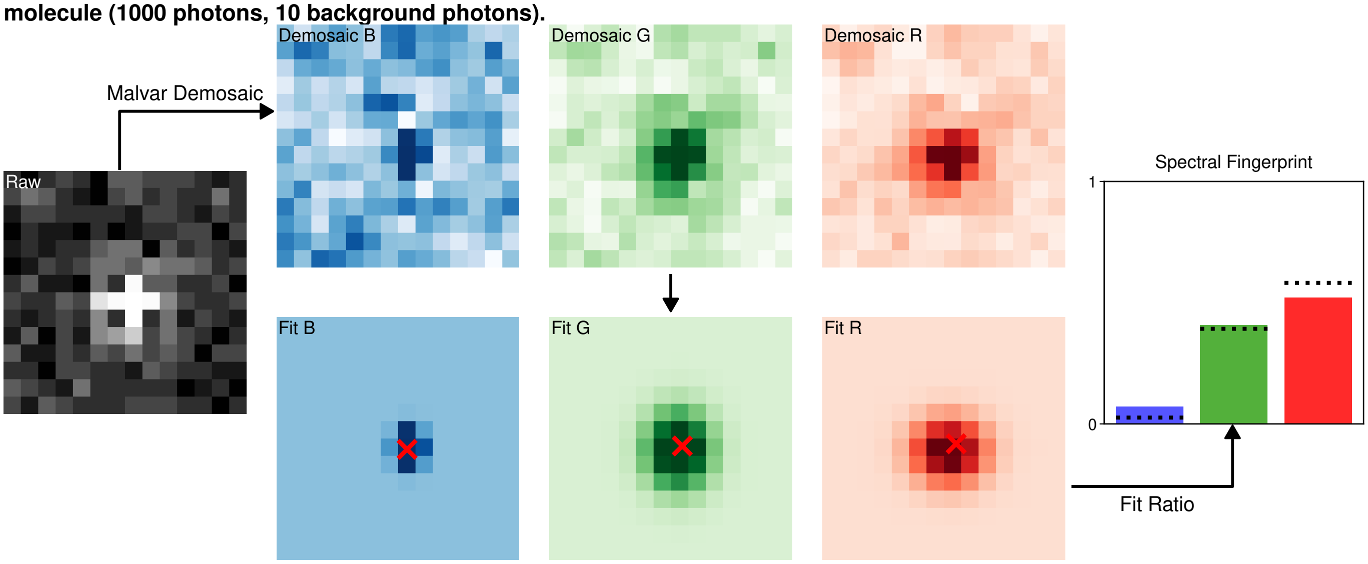


**Fig. S9.** **An example of the strategy of using Malvar demosaicing, and fitting the demosaiced images on an ATTO 565 molecule (1000 photons, 10 background photons).**

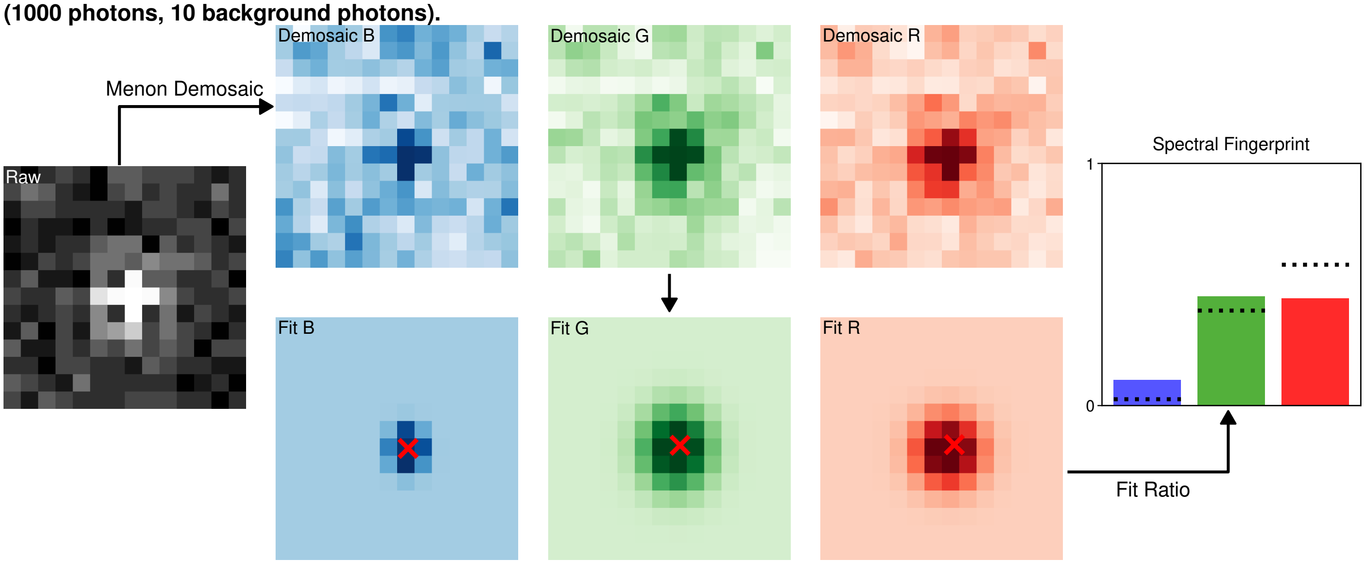


**Fig. S10.** **An example of the strategy of using Menon demosaicing, and fitting the demosaiced images on an ATTO 565 molecule (1000 photons, 10 background photons).**

transmission spectra.

For each condition, a two-dimensional Gaussian point-spread function (for full details of image simulation, see section S1) was placed at a position drawn uniformly within the central pixel of a simulated sensor patch and either fitted by the direct fitting algorithm (see section S3) or by the demosaic and fit strategy outlined above. This process was repeated 10,000 times (bootstraps) per photon level to obtain statistical estimates of spatial precision $\sigma_{xy} = \sqrt{(\sigma^2_{\epsilon_x} + \sigma^2_{\epsilon_y})/2}$, and colour precision ($\sigma_{\text{colour}}$, the standard deviation of the Euclidean distance in pixel QE from the known spectral fingerprint of the dye). Simulations were performed across 200 photon levels spaced logarithmically between 500 and 50,000 photons.

Spatial precision and colour precision for the different fitting strategies are shown in Fig. S11. As is clear by inspection, the S$^3$M strategy of directly fitting the raw data with the model described in section S3 far outperforms any of the demosaicing algorithms. This is true across all dyes, with precision values per dye shown in Fig. S12. This, by inspection of the relative pixel QEs returned by different fitting strategies (Fig. S13–S15), appears to be due to the significantly larger number of poor fits the bilinear demosaicing strategy leads to, and the systemic bias that Malvar and Menon demosaicing introduce into the results.

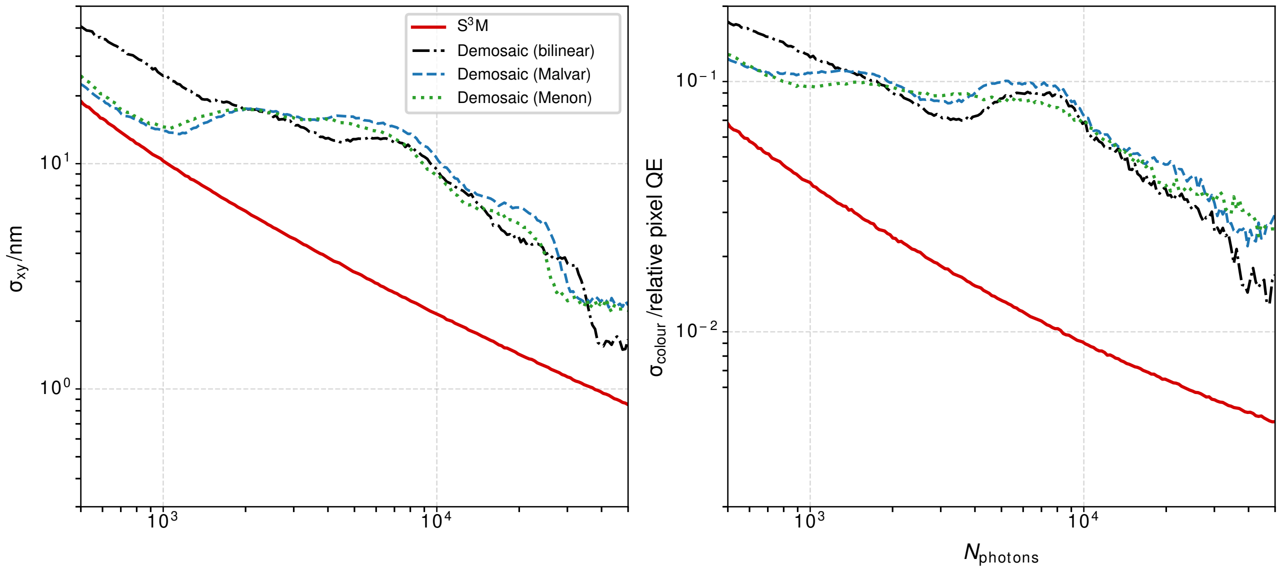


**Fig. S11. Fitting raw data *vs.* demosaicing.** Each demosaicing algorithm tested performs worse than the S$^3$M strategy across a range of photons.

As an aside, the S$^3$M fitting strategy also outperforms demosaicing in terms of efficiency. Timing the fitting strategy of fitting one image directly *versus* three demosaiced images shows clearly that S$^3$M is a more efficient strategy (Fig. S16).

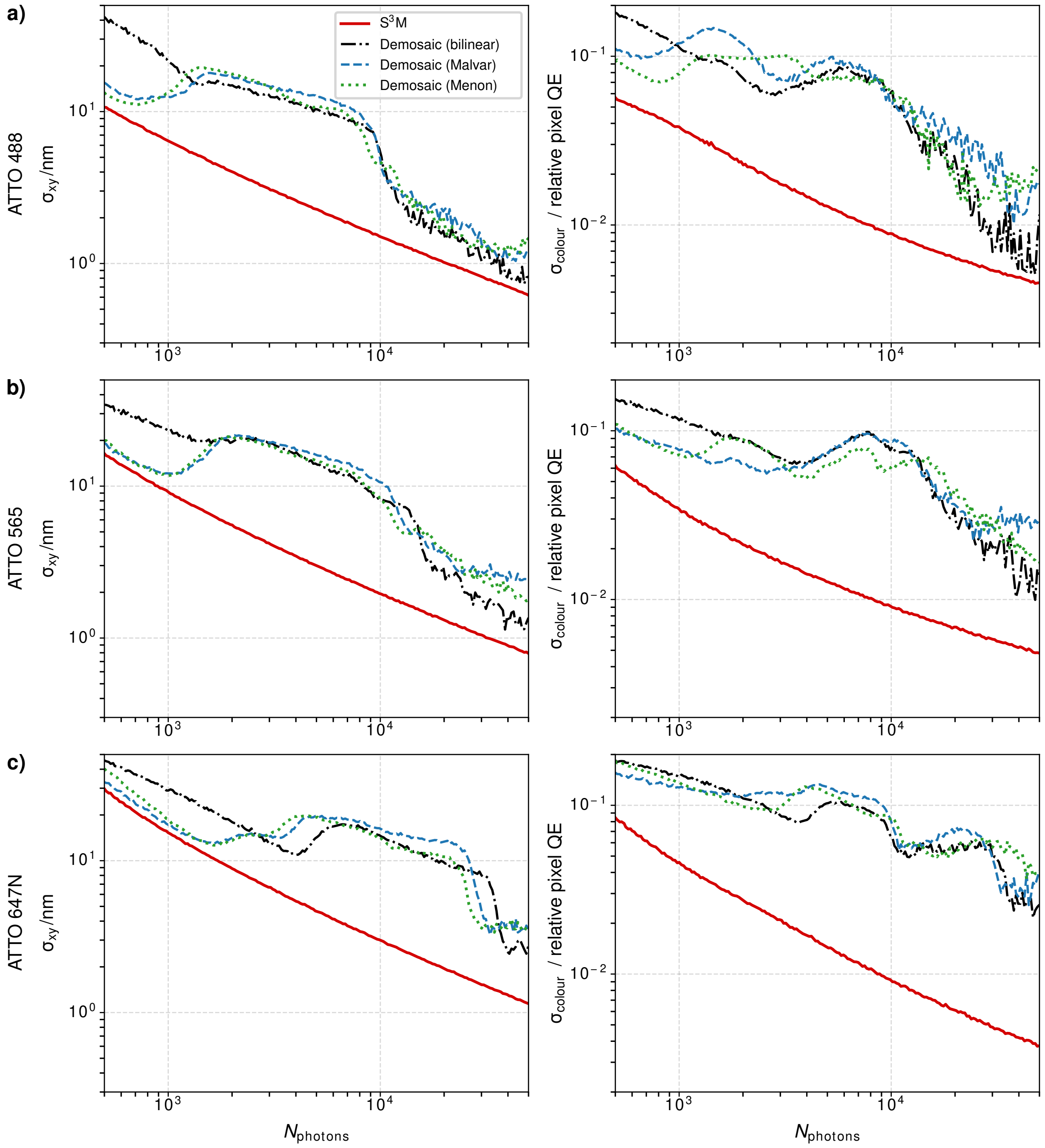


**Fig. S12. Fitting raw data *vs.* demosaicing. a)** Localisation and colour precision for ATTO 488. **b)** Localisation and colour precision for ATTO 565. **c)** Localisation and colour precision for ATTO 647N.

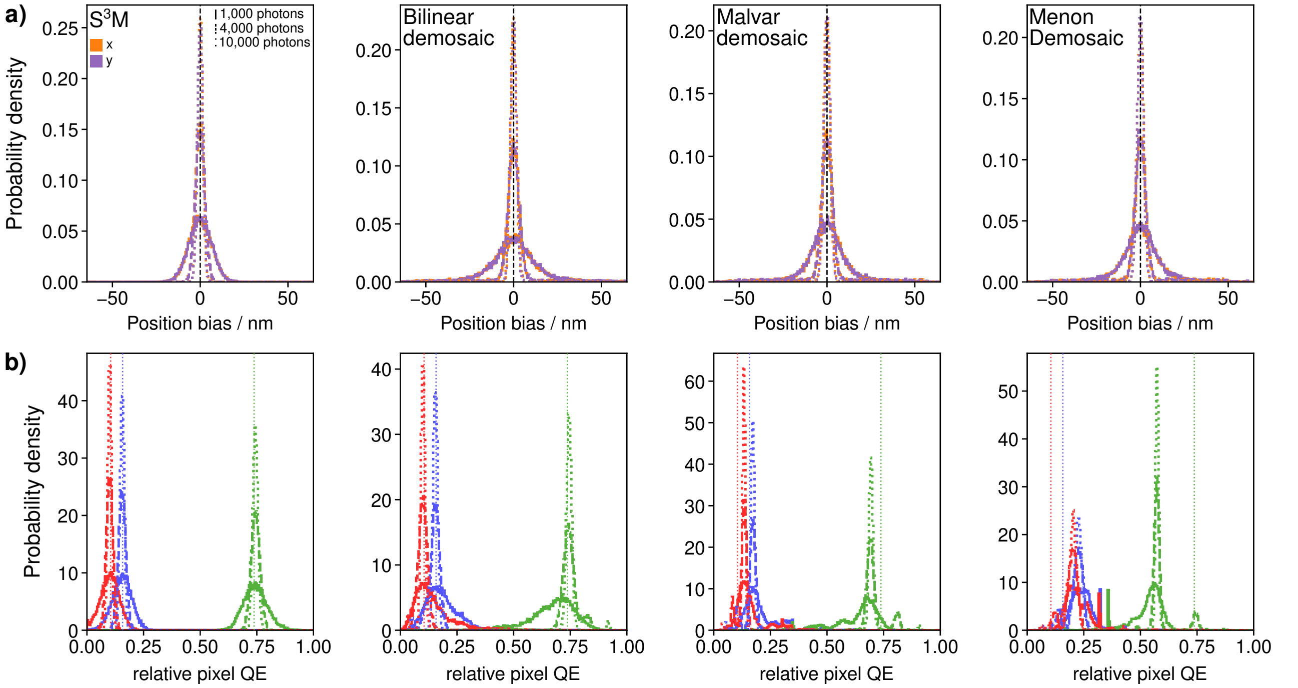


**Fig. S13. Localisation precision bias histograms and relative pixel QE results for different fitting strategies on ATTO 488. a)** Localisation bias for x and y at three photon values. **b)** Relative pixel QE values extracted by different fitting strategies at three different photon values.

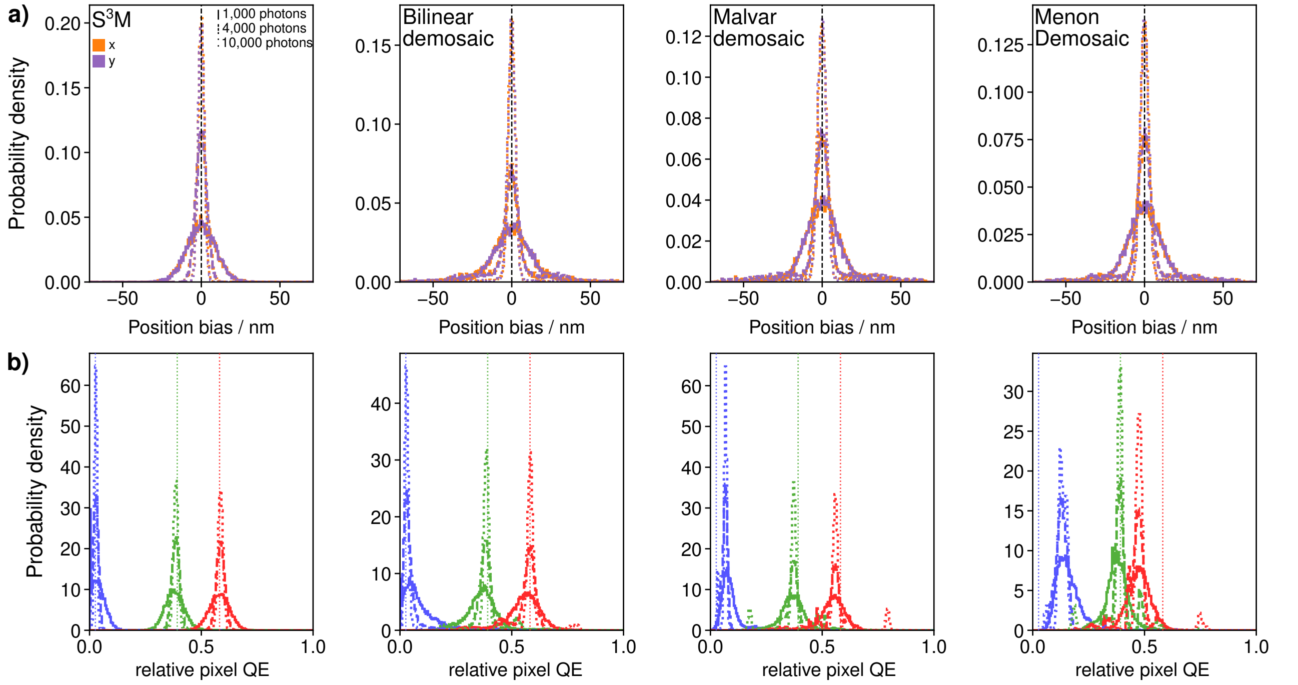


**Fig. S14. Localisation precision bias histograms and relative pixel QE results for different fitting strategies on ATTO 565. a)** Localisation bias for x and y at three photon values. **b)** Relative pixel QE values extracted by different fitting strategies at three different photon values.

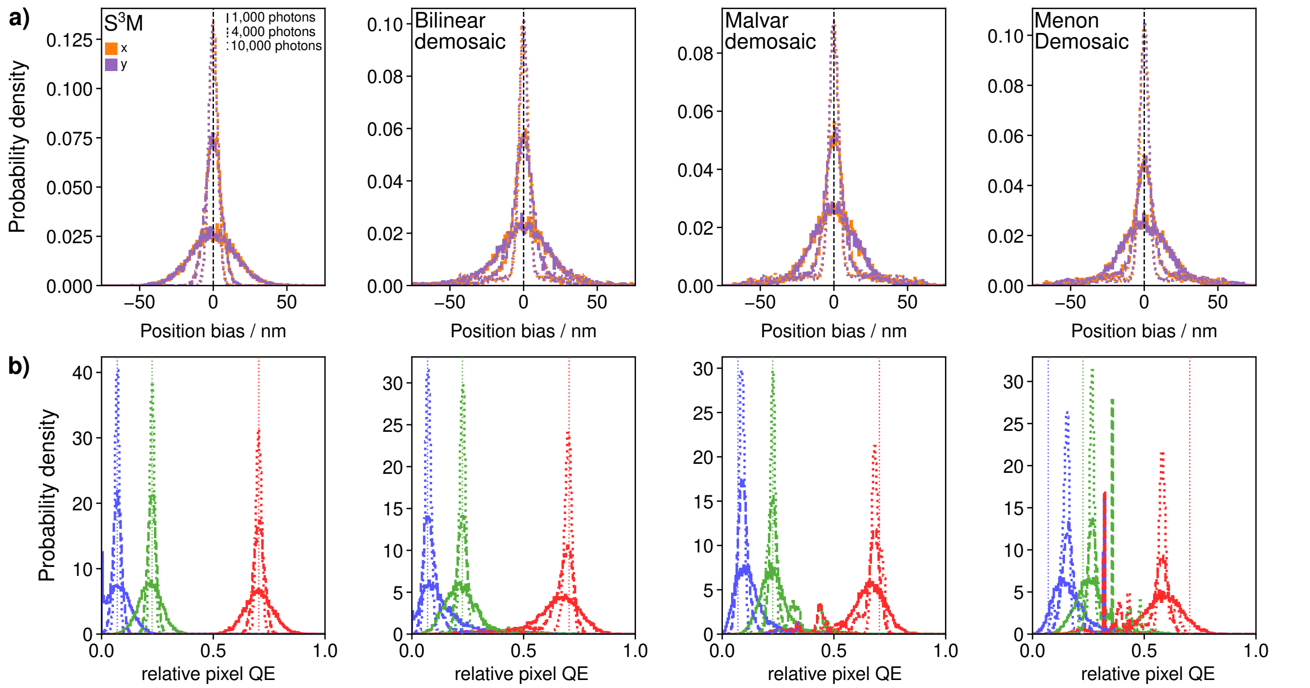


**Fig. S15. Localisation precision bias histograms and relative pixel QE results for different fitting strategies on ATTO 647N. a)** Localisation bias for x and y at three photon values. **b)** Relative pixel QE values extracted by different fitting strategies at three different photon values.

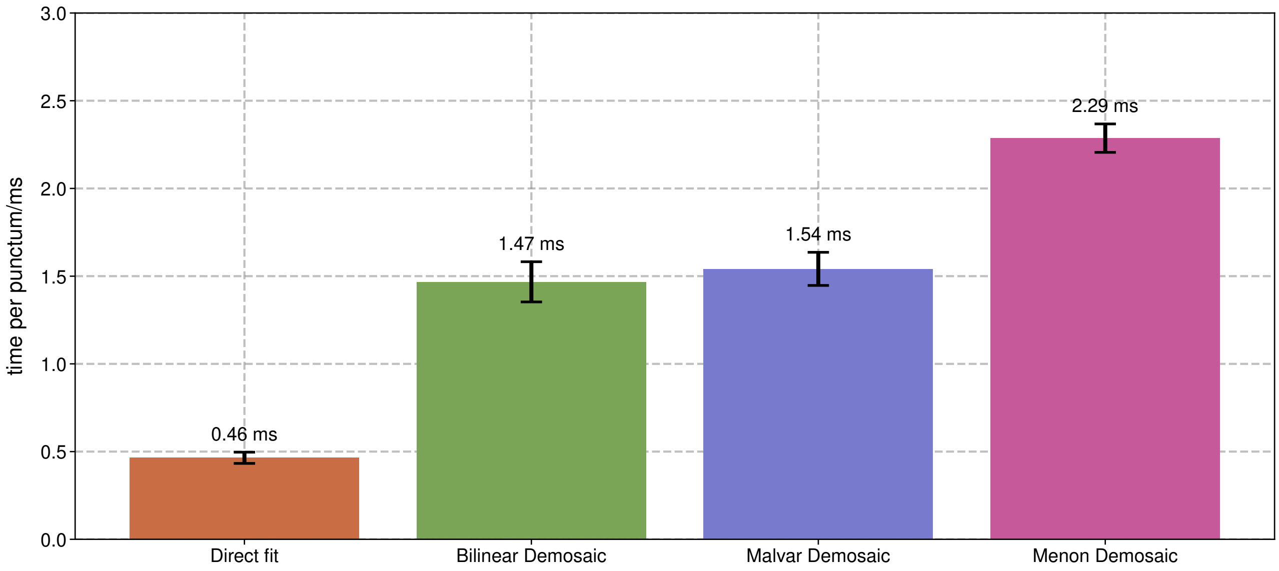


**Fig. S16. Timing, per single punctum with no parallelisation applied, of fitting strategies.** Fitting algorithms were run 10,000 times.

#### Supplementary Note S5: Thorlabs CS505CU Single-Molecule Data

Below (Fig. S17) we provide exemplar photobleaching traces of three ATTO 647N dyes imaged with the Thorlabs CS505CU. Additionally, we provide zoomed-in PSFs of 25 additional molecules below in Fig. S18.

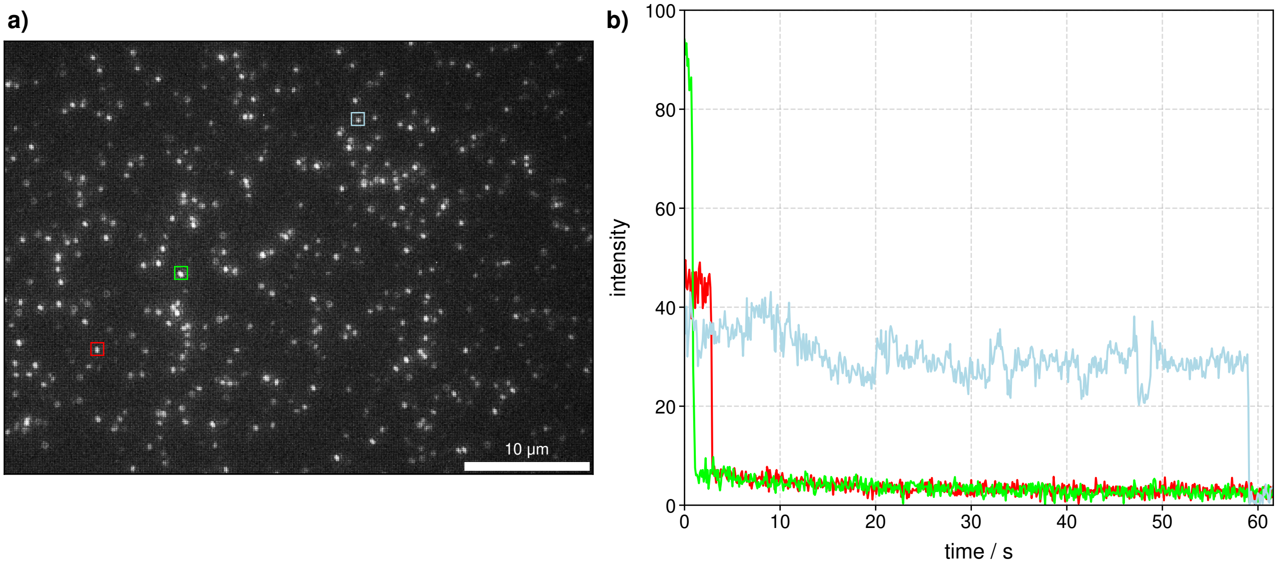


**Fig. S17. ATTO 647N. a)** A single FOV of 100 pM ATTO 647N molecules in 1 % PVA. **b)** Single ATTO 647N molecules show characteristic single-step photobleaching traces expected for single-molecules.

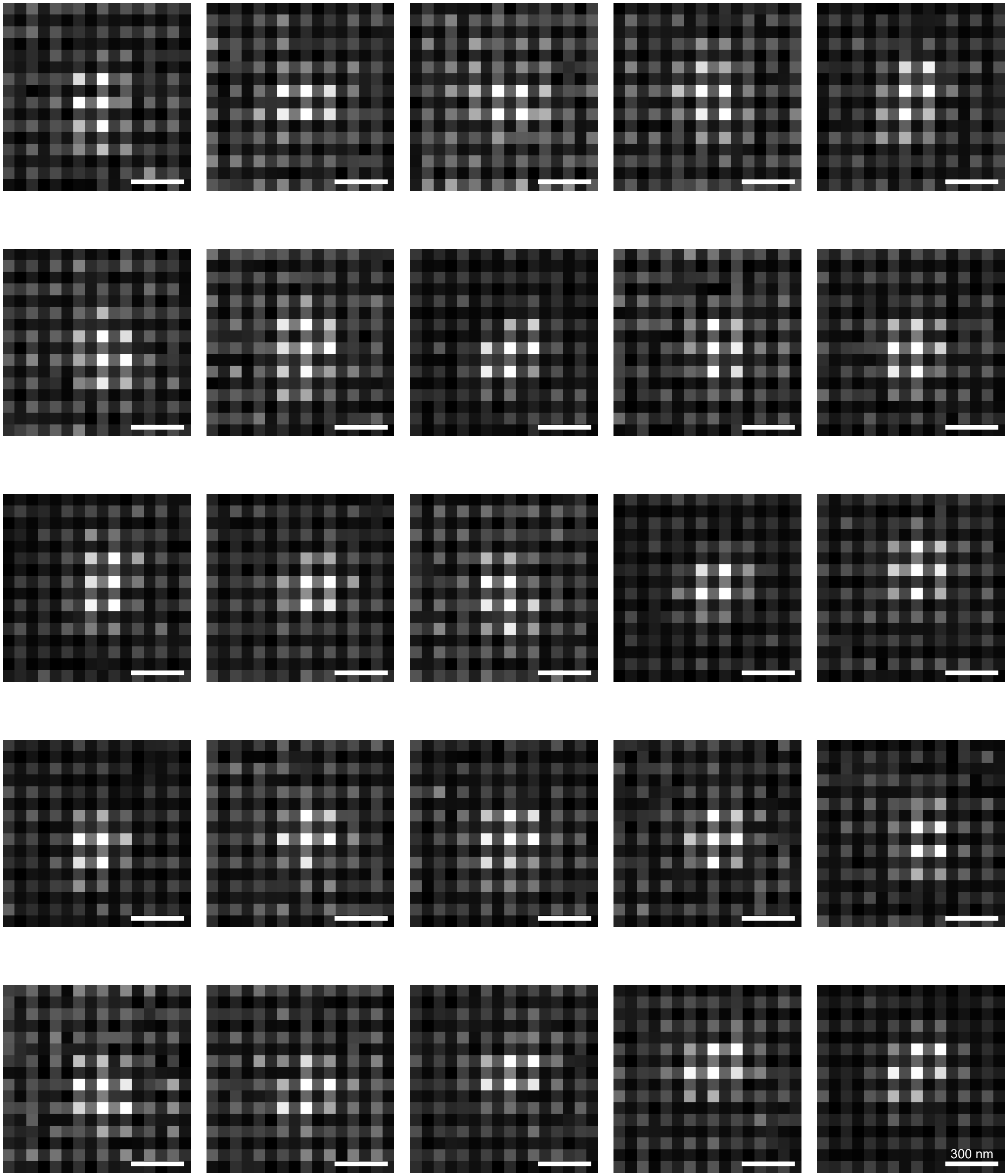


**Fig. S18. ATTO 647N PSFs.** 25 PSFs of ATTO 647N molecules in 1 % PVA. Scalebar is 300 nm.

#### Supplementary Note S6: ZWO ASI 585MC Single-Molecule Data

Below (Fig. S19) we provide exemplar photobleaching traces of three ATTO 647N dyes imaged with the ZWO ASI 585MC. Additionally, we provide zoomed-in PSFs of 25 additional molecules below in Fig. S20.

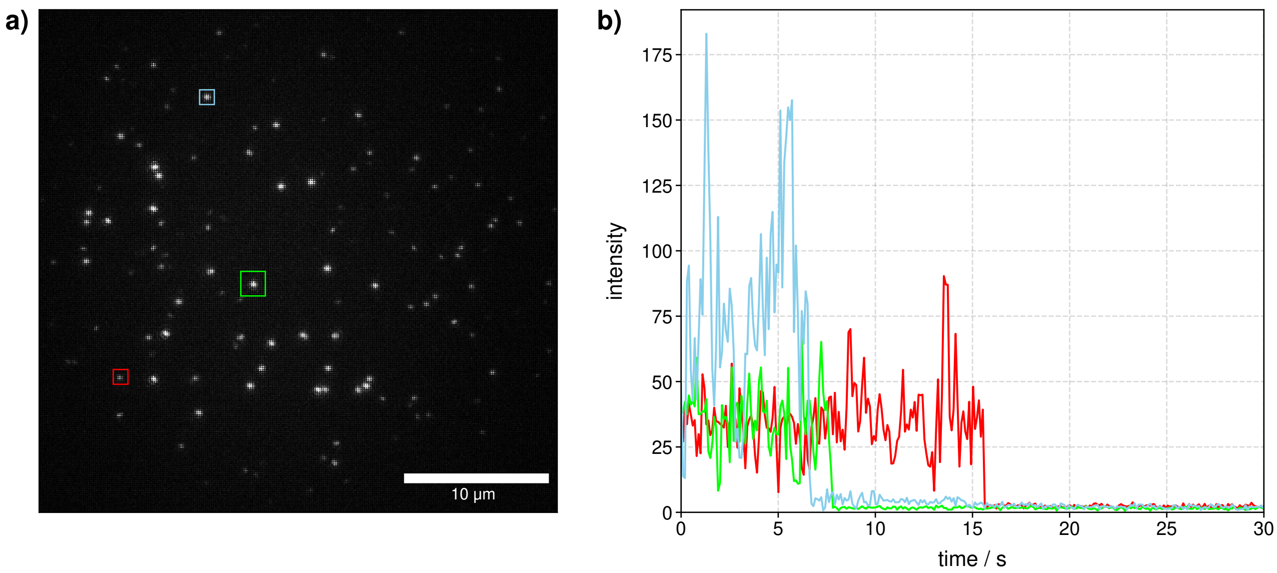


**Fig. S19. ATTO 647N. a)** A single FOV of 100 pM ATTO 647N molecules in 1 % PVA. **b)** Single ATTO 647N molecules show characteristic single-step photobleaching traces expected for single-molecules.

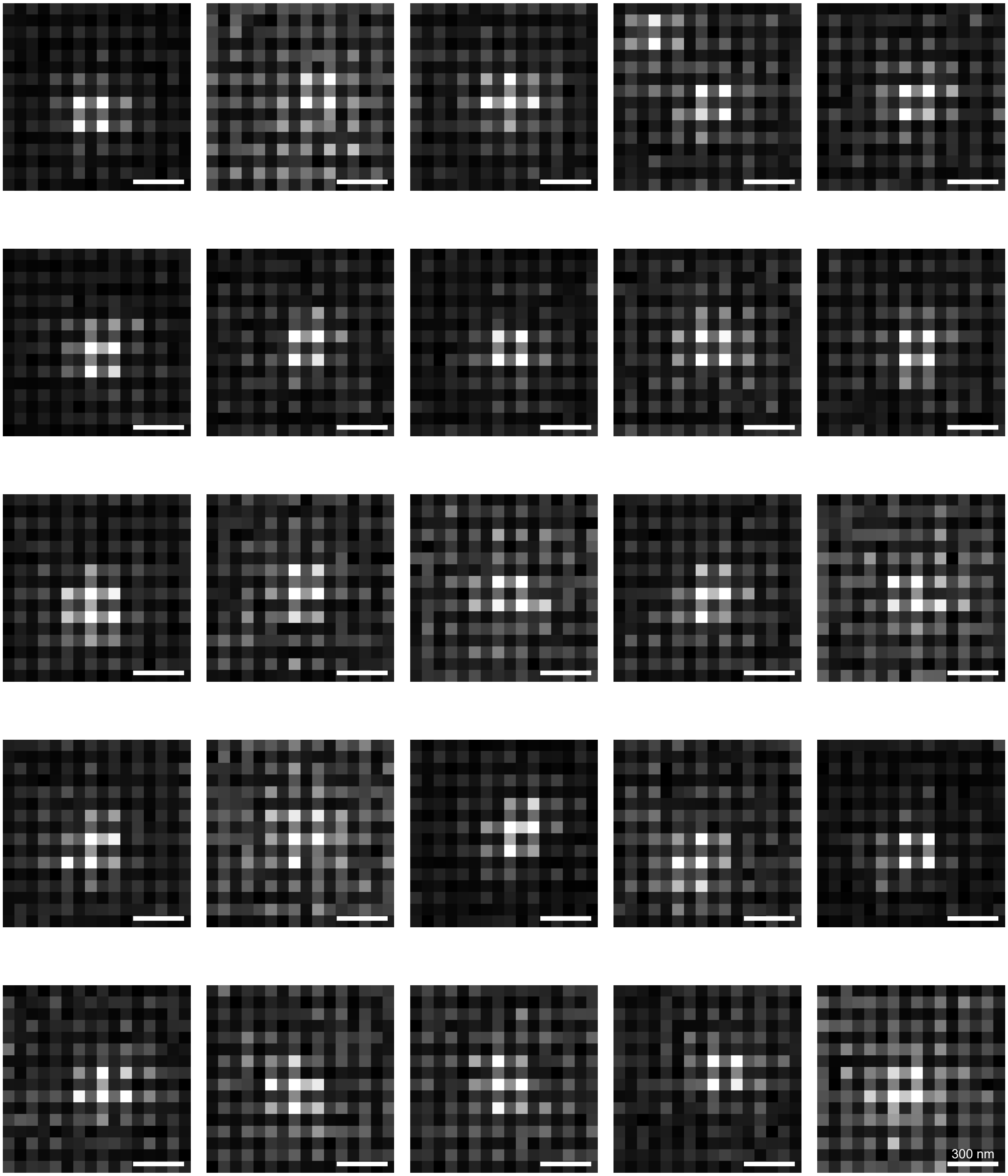


**Fig. S20. ATTO 647N PSFs.** 25 PSFs of ATTO 647N molecules in 1 % PVA. Scalebar is 300 nm.

#### Supplementary Note S7: Single-step photobleaching of single dyes

Below (Fig. S21–S32) we provide exemplar photobleaching traces of every dye imaged using the Ximea MC050CG-SY.

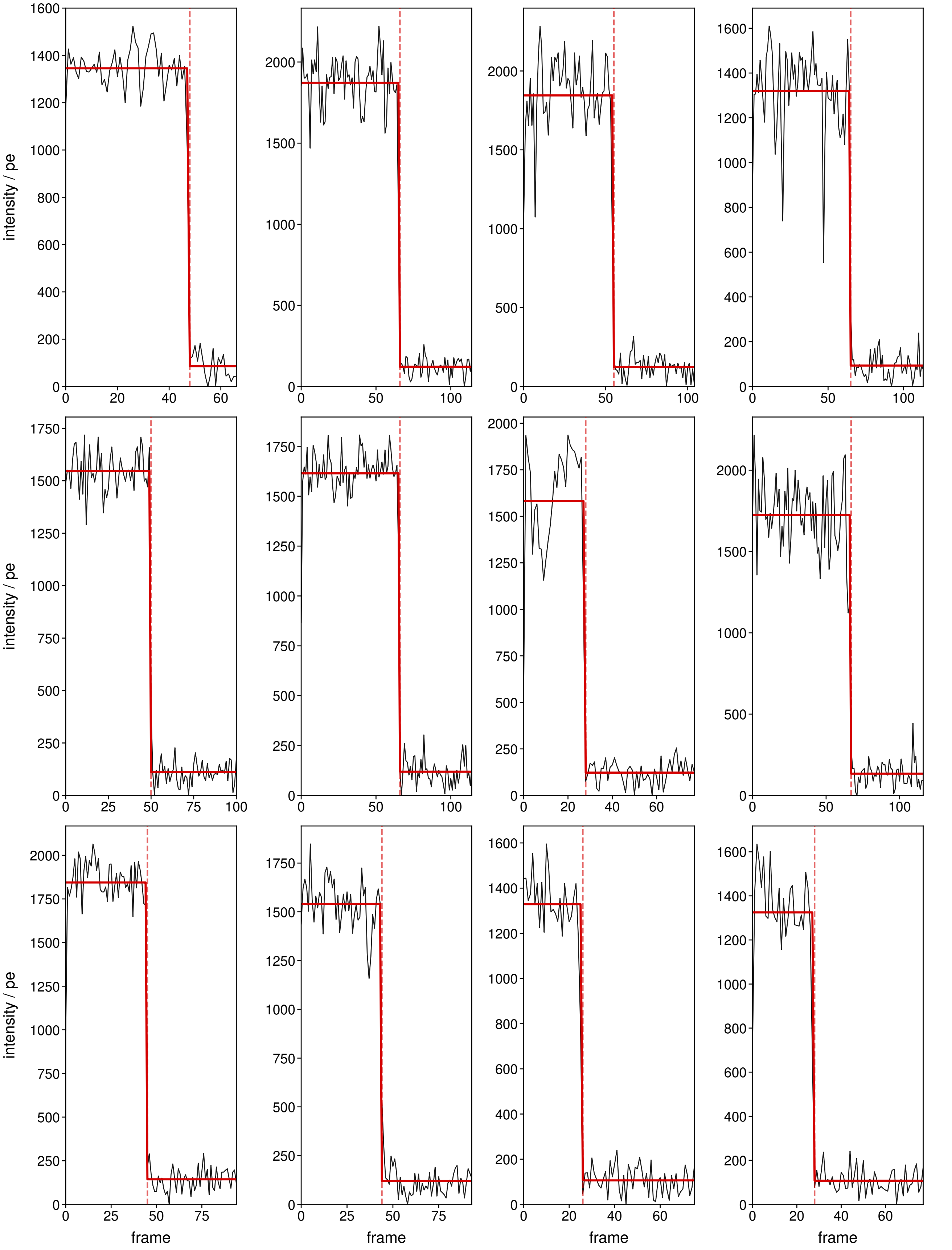


**Fig. S21. ATTO 488.**

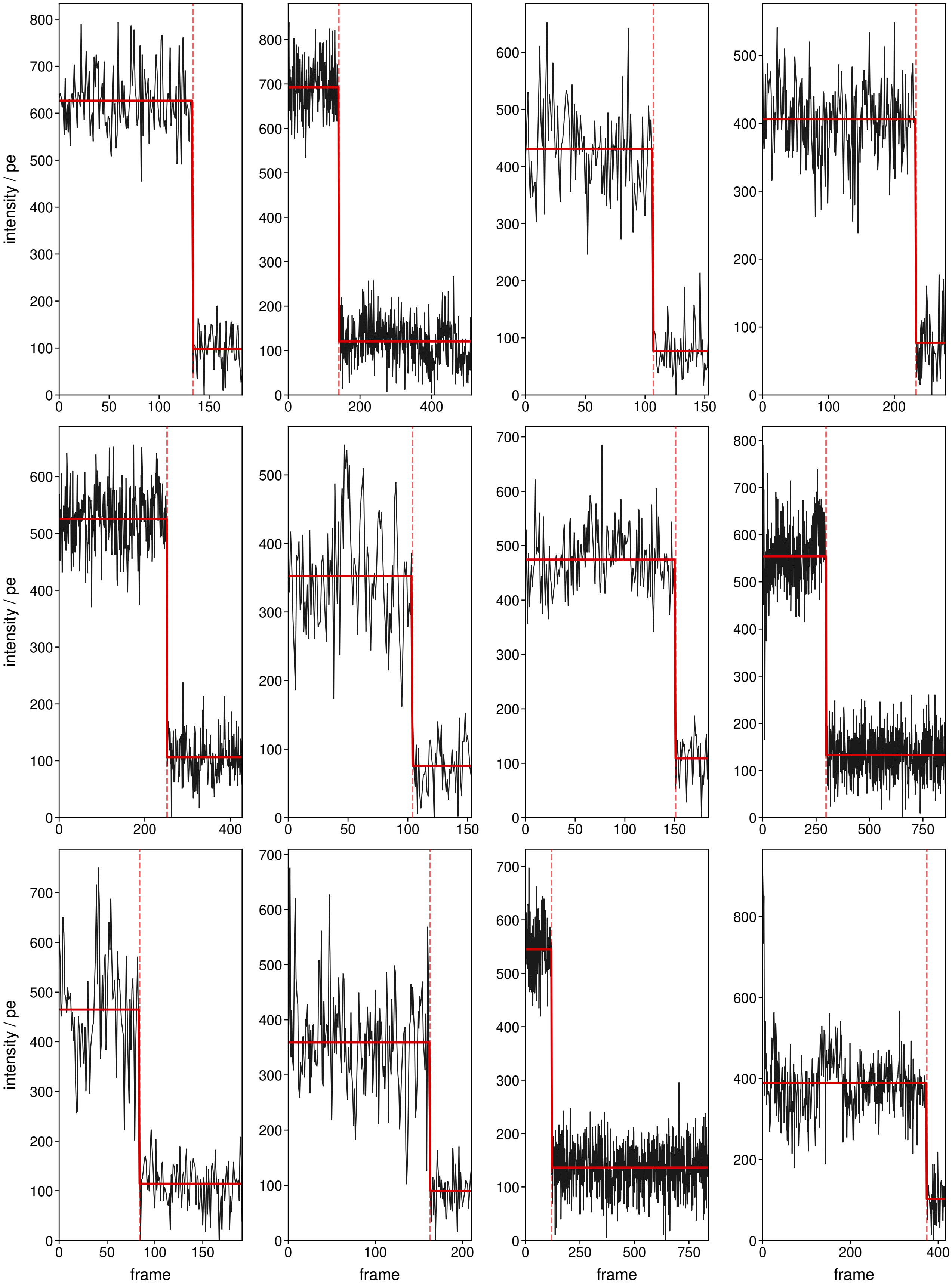


**Fig. S22. ATTO 514.**

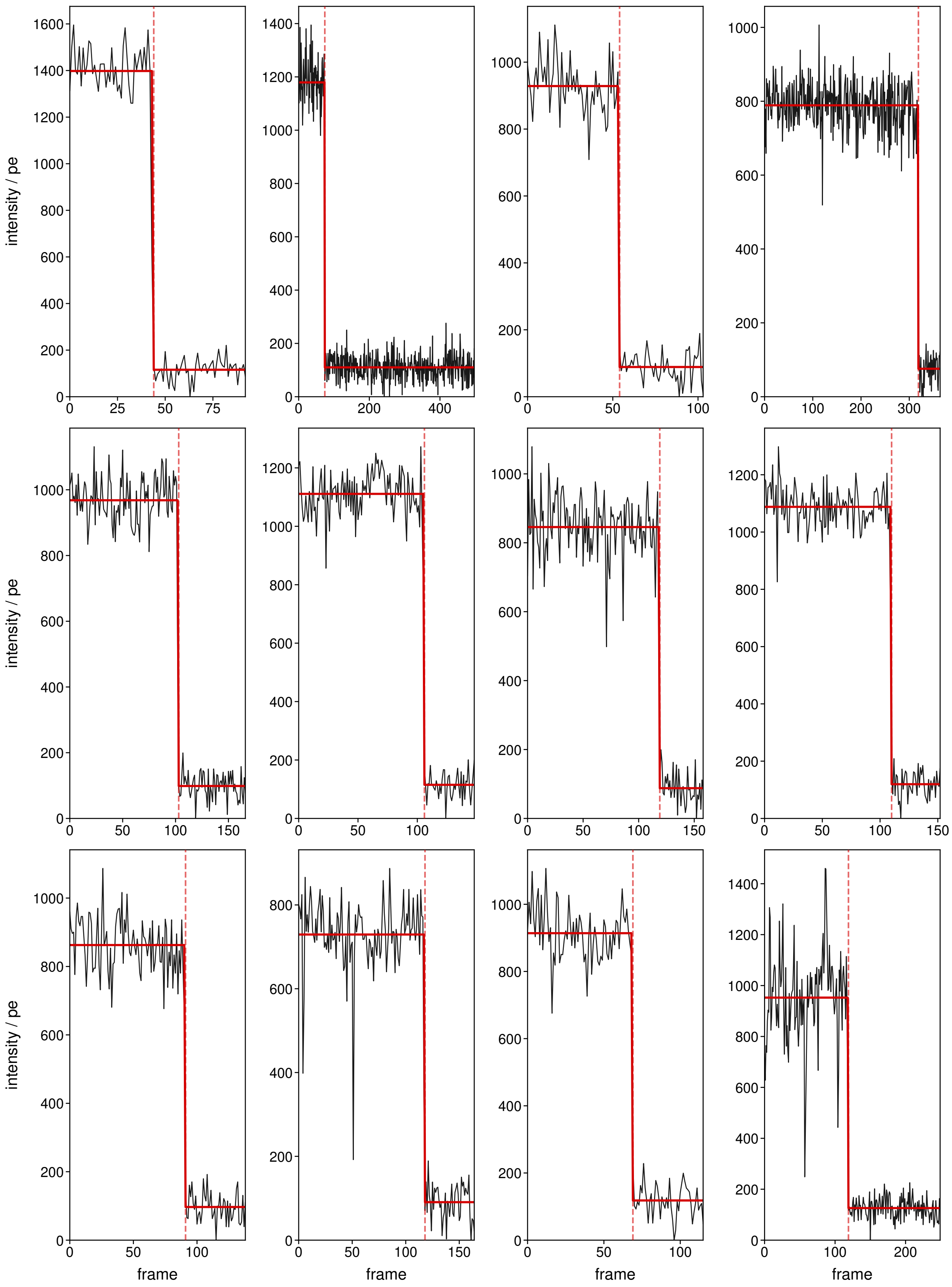


**Fig. S23. ATTO 520.**

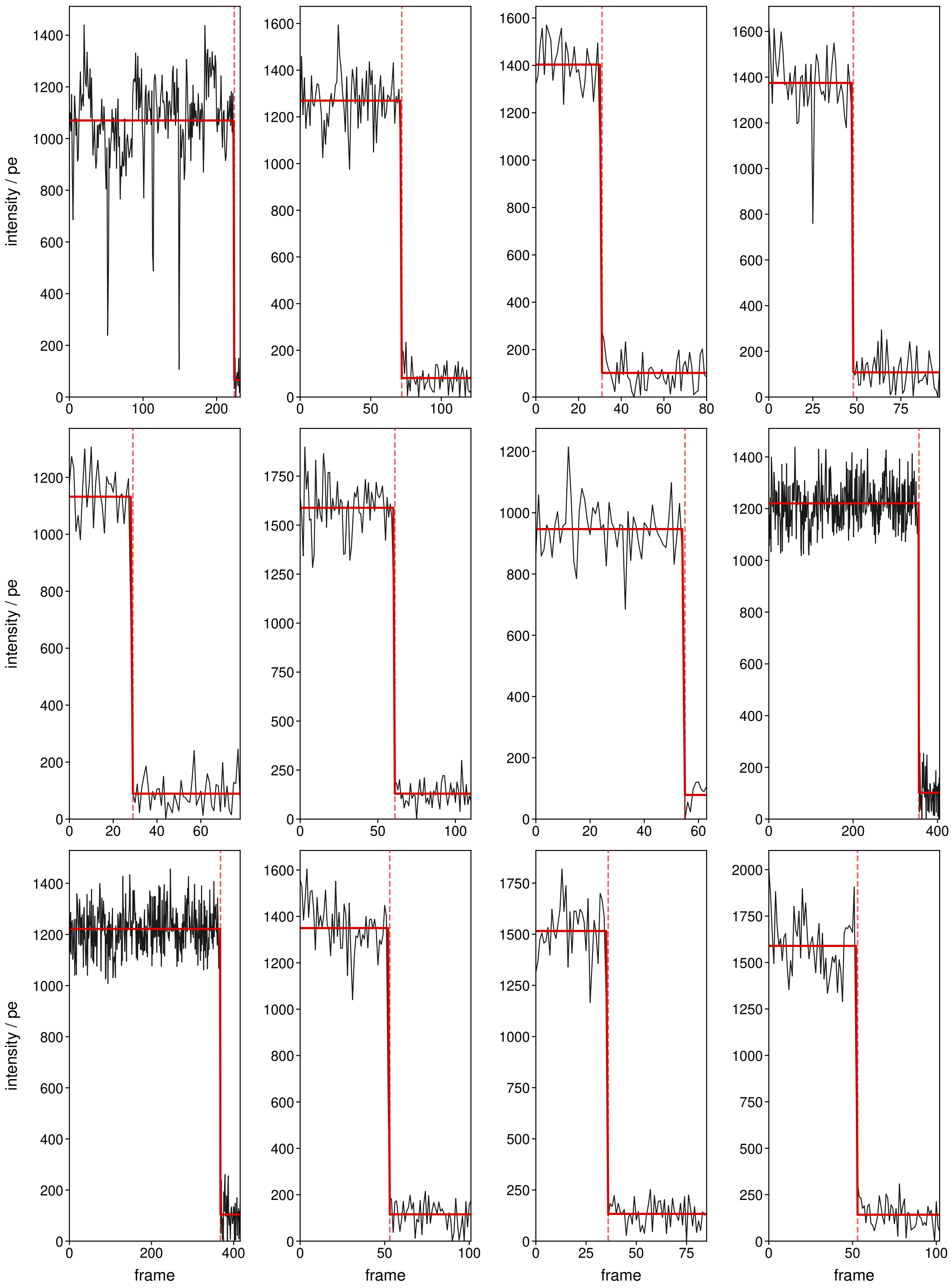


**Fig. S24. ATTO Rho6G.**

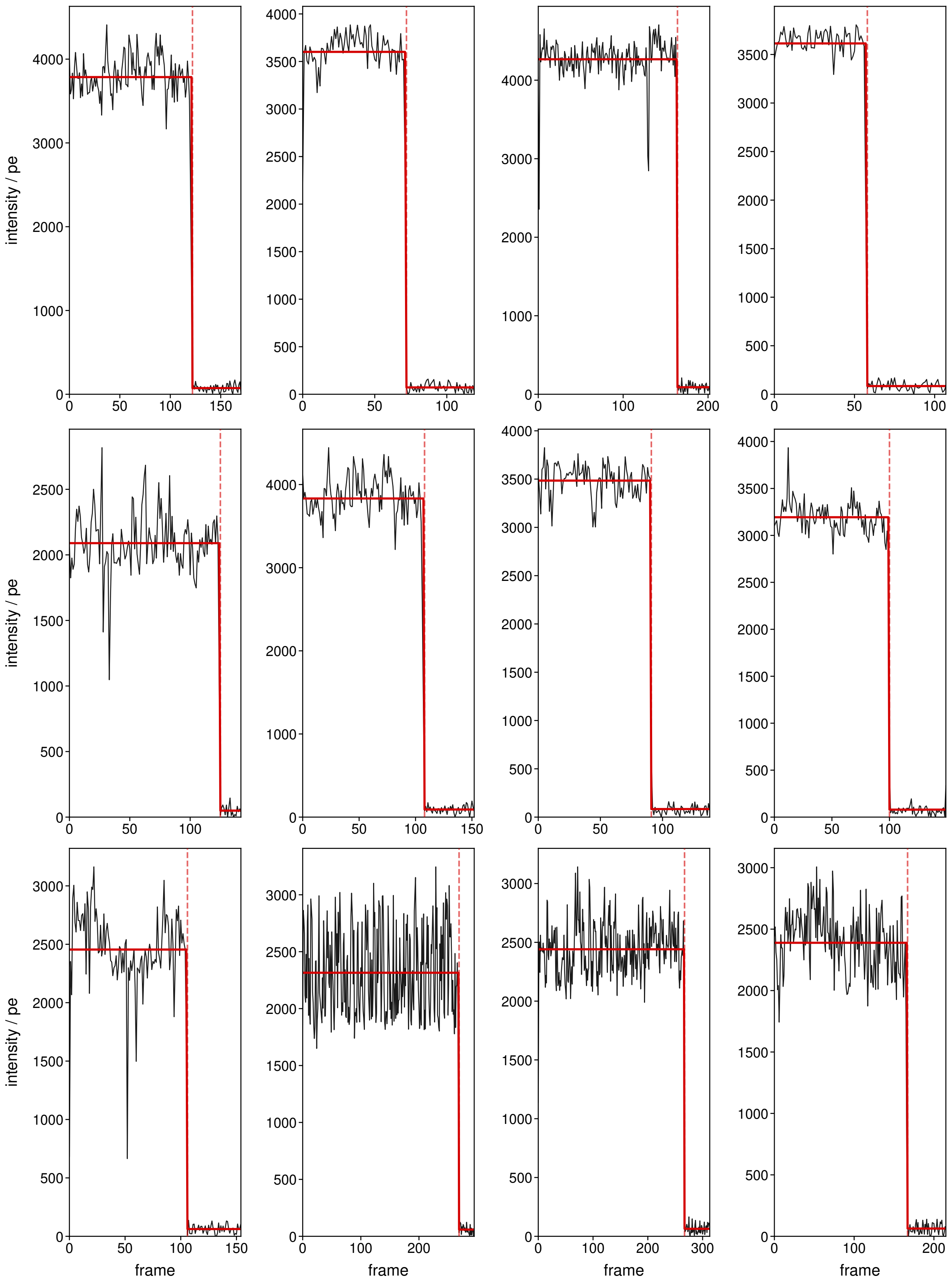

**Fig. S25. ATTO 565.**

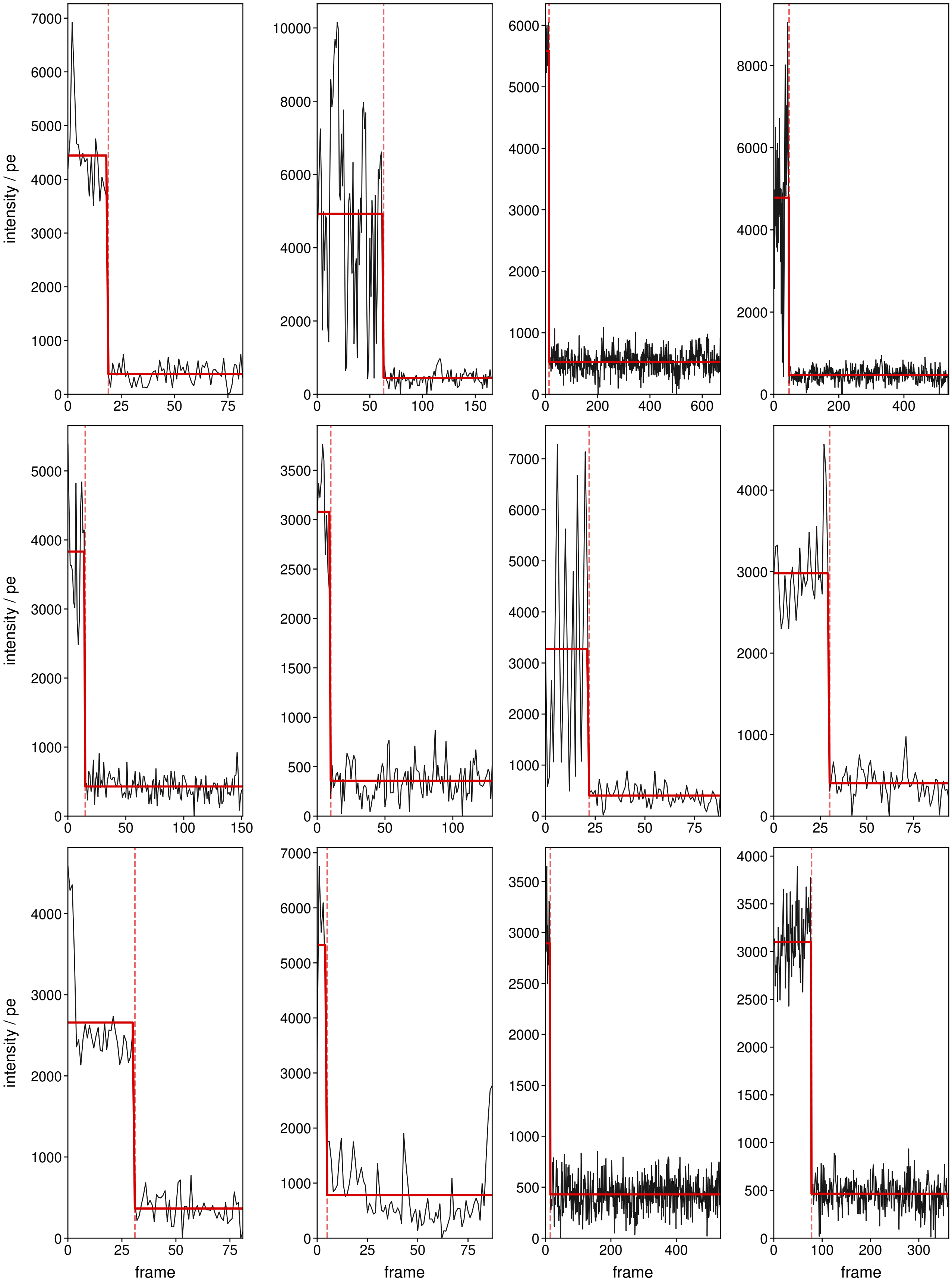


**Fig. S26. ATTO 594.**

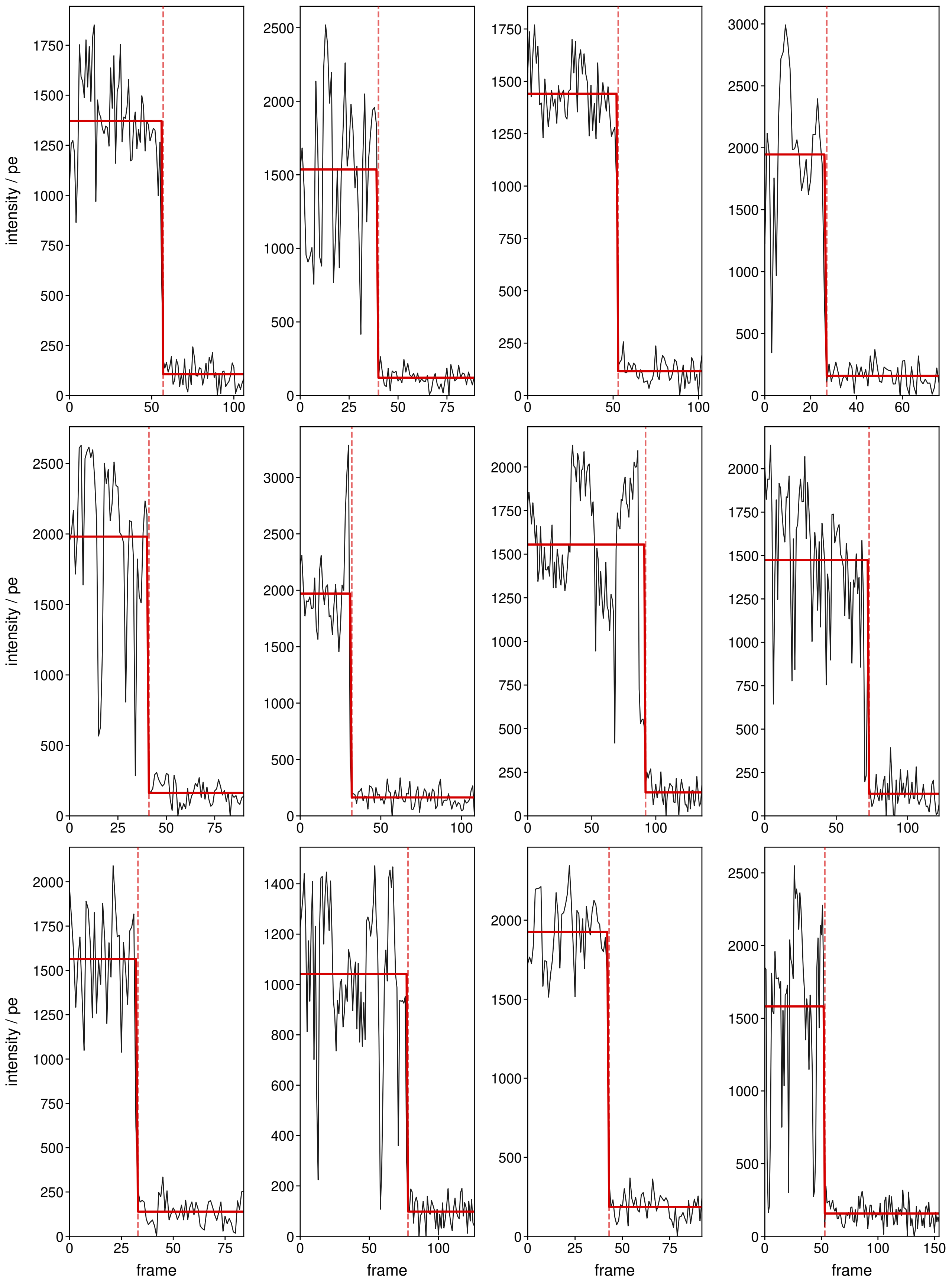


**Fig. S27. ATTO 620.**

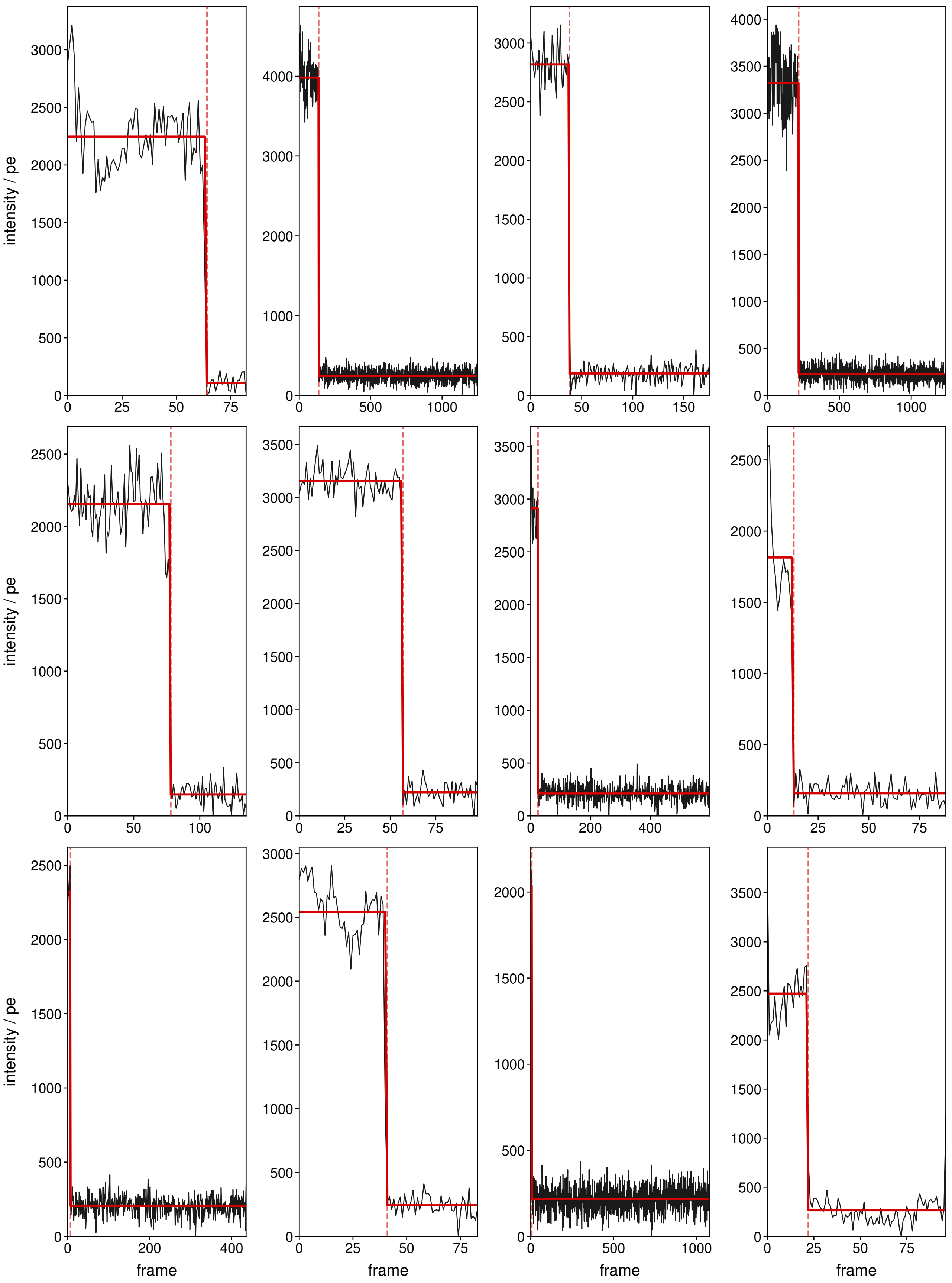


**Fig. S28. ATTO 633.**

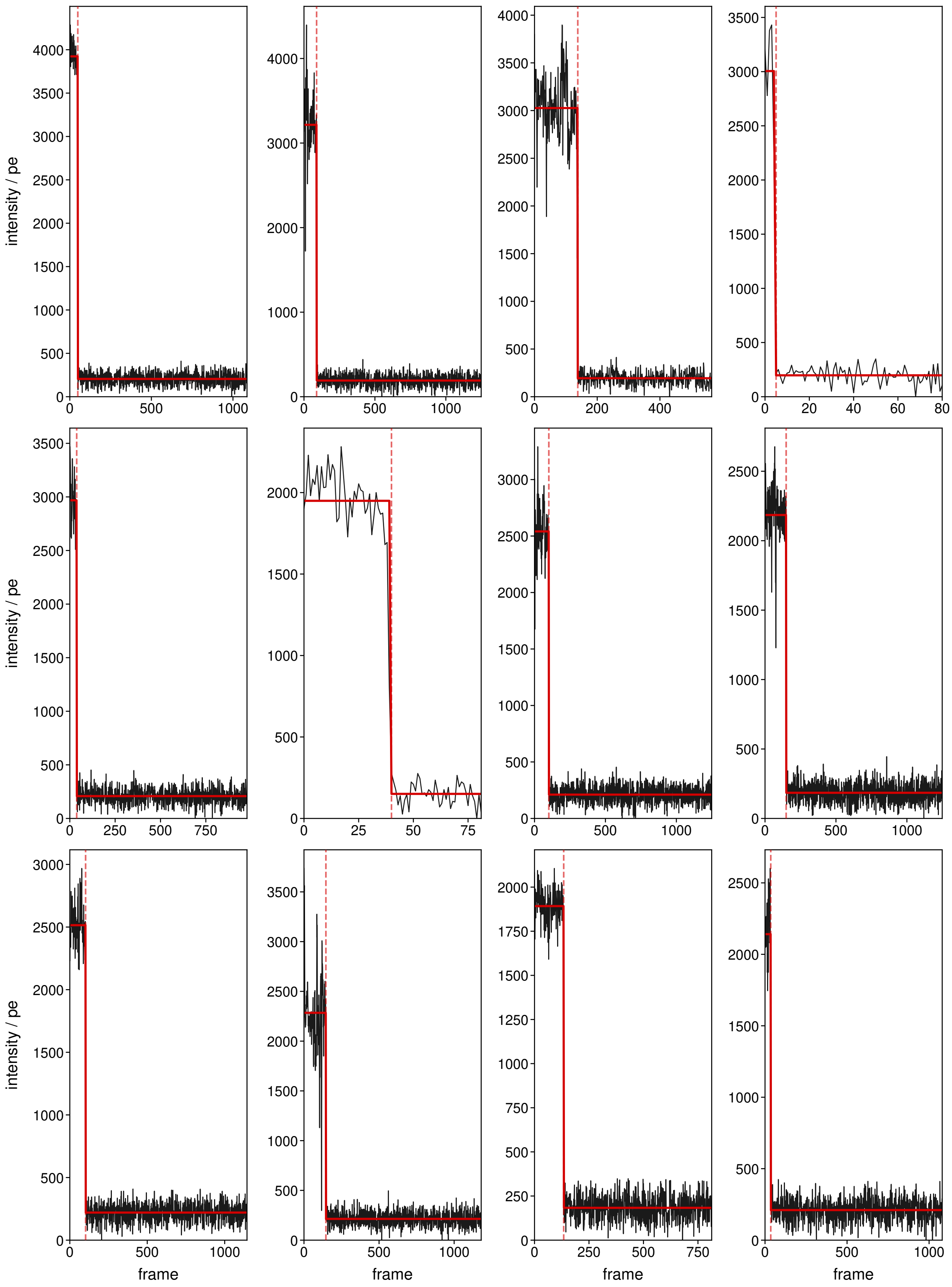


**Fig. S29. ATTO 647N.**

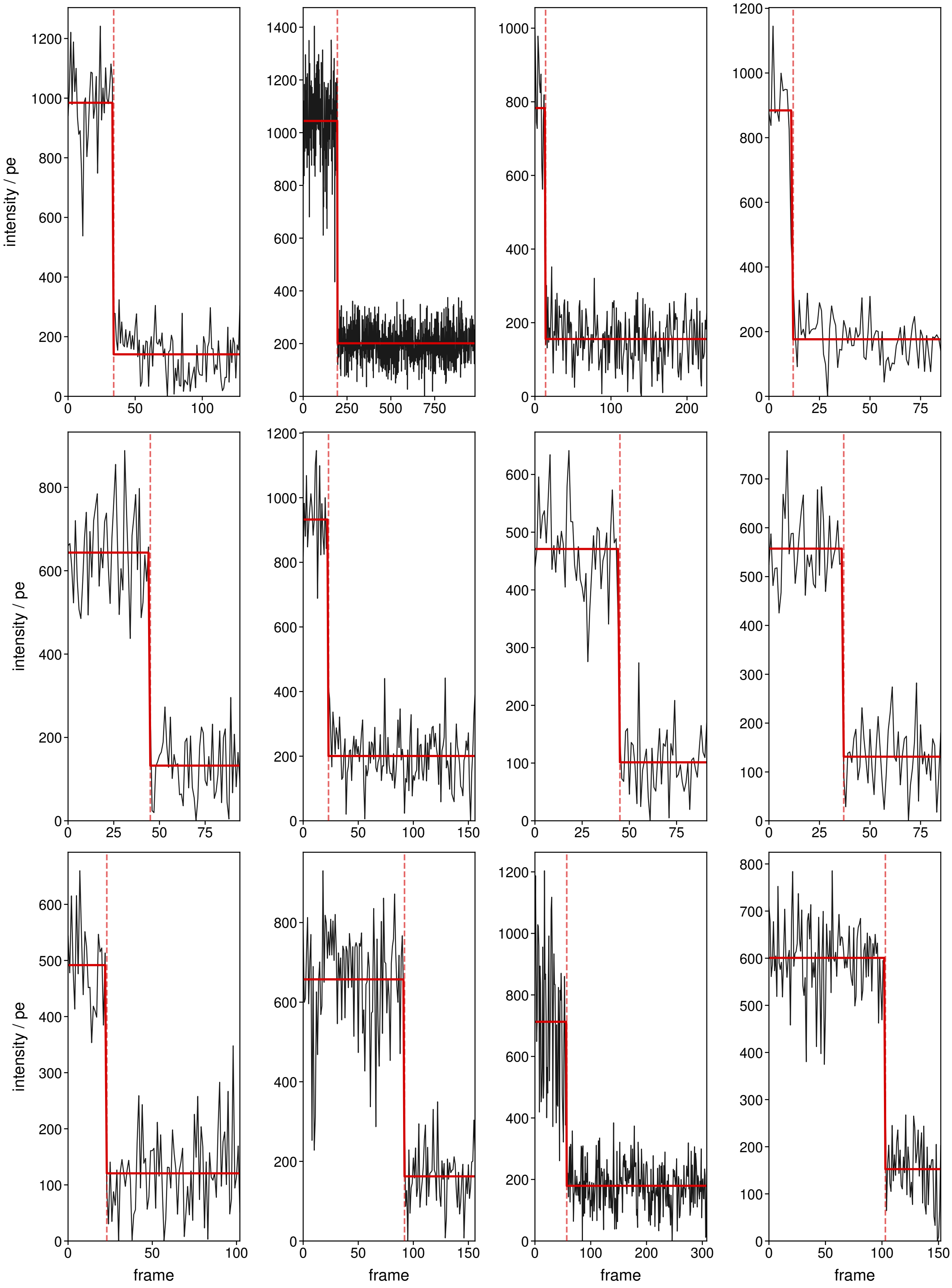


**Fig. S30. ATTO 655.**

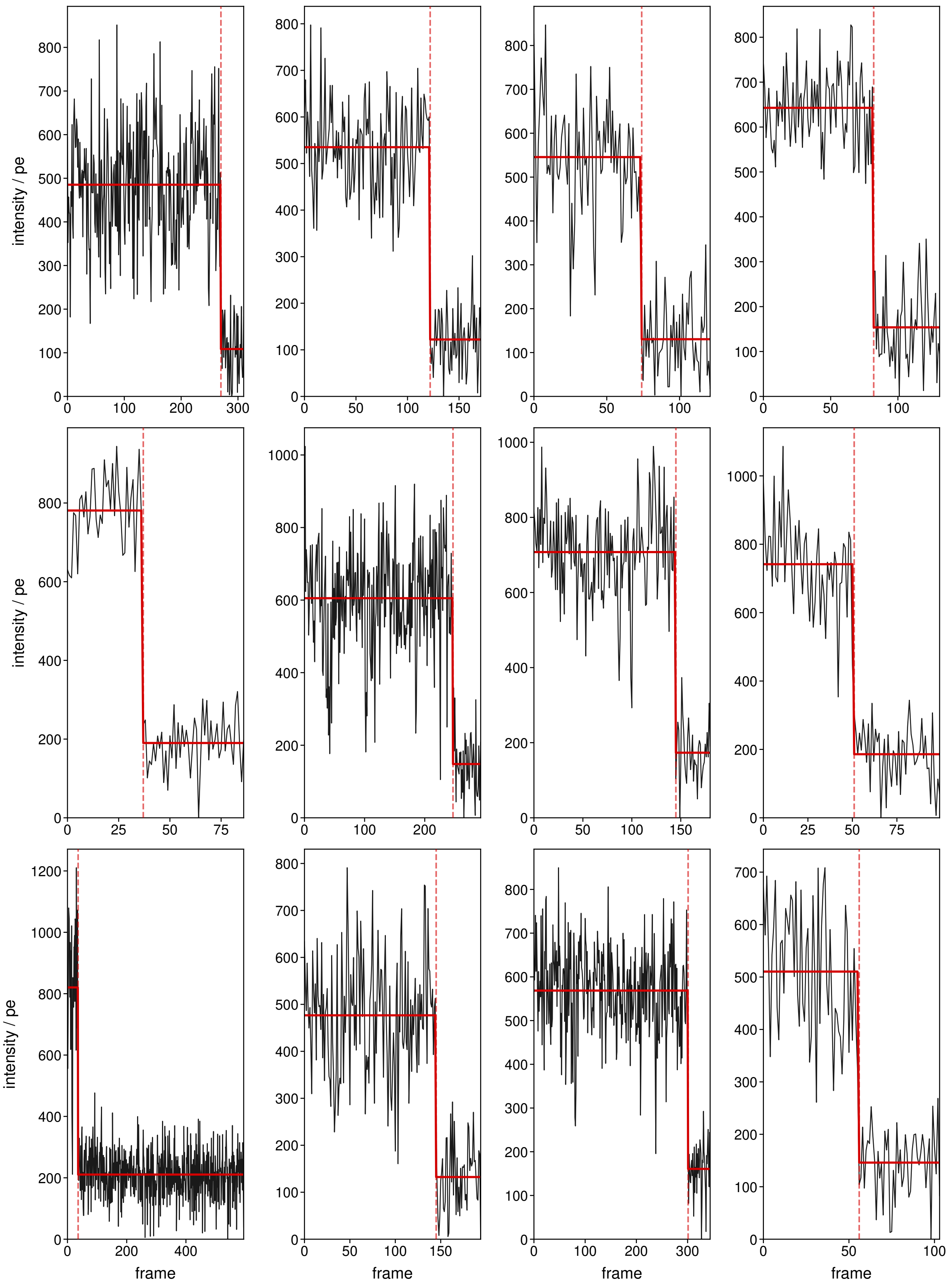


**Fig. S31. ATTO 700.**

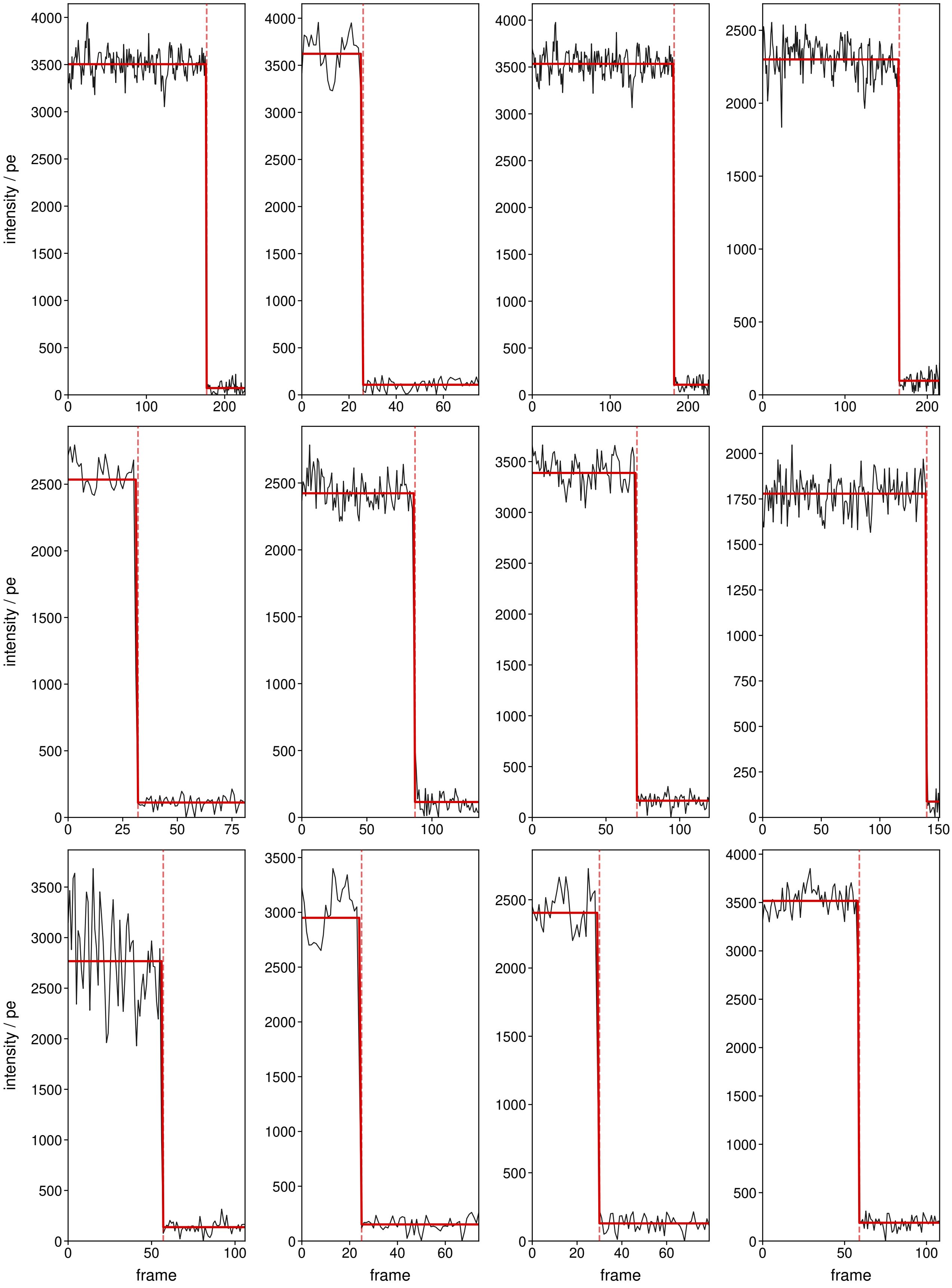


**Fig. S32. LD 655.**

#### Supplementary Note S8: Single-molecule spectral fingerprint *vs.* predicted value

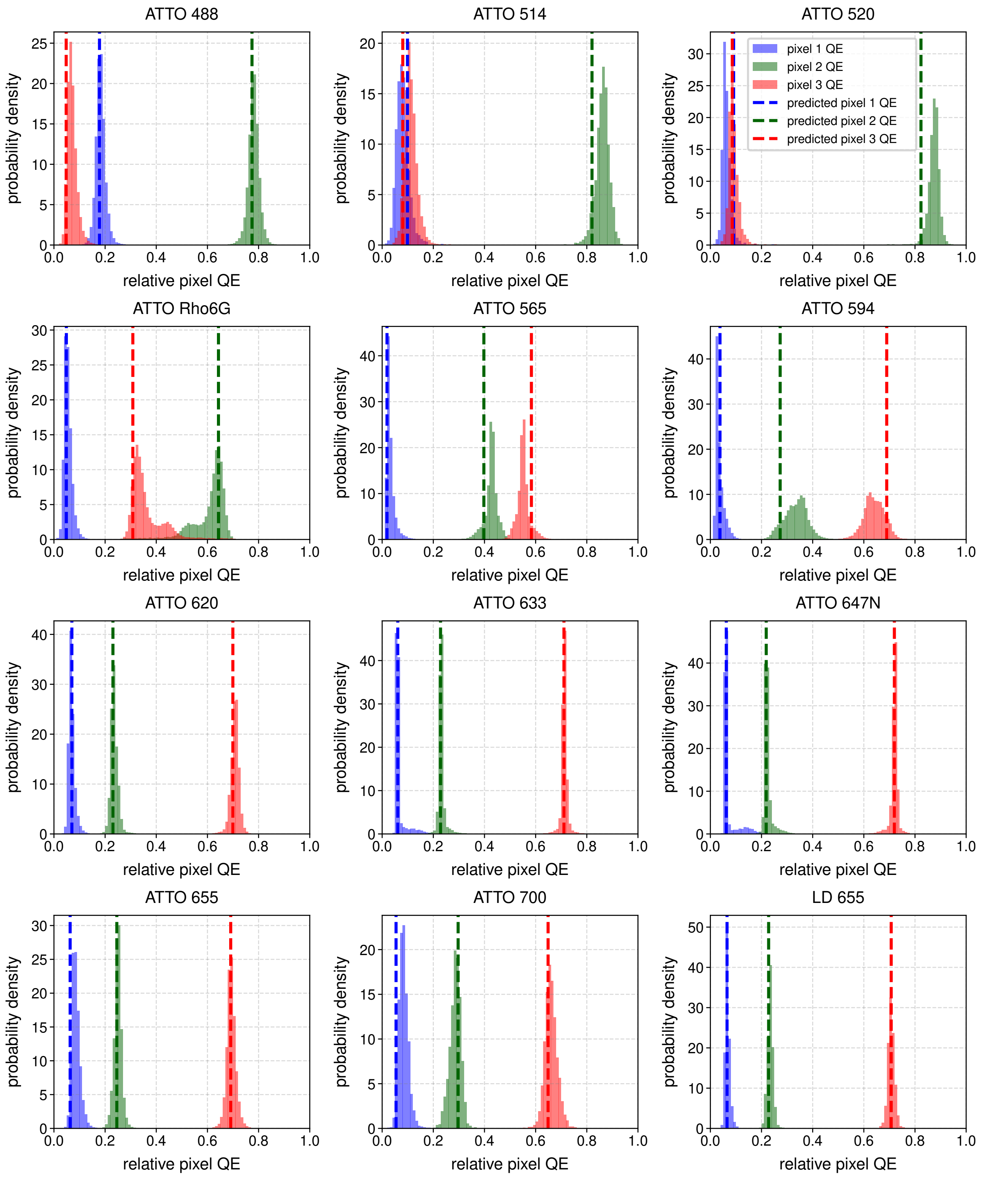


**Fig. S33. Single-molecule spectral fingerprint versus predicted value.** Histograms of the relative QE of each pixel per molecule for the 12 dyes in SI Fig. S21–S32. Predicted values are derived from the available spectra of each dye, the optical filters used, and the pixel dependent QE (see Section S1).

#### Supplementary Note S9: Effect of Background on S$^3$M

Biological imaging often includes autofluorescence, scattering and other sources of background.(10) We therefore performed simulations to determine the effect of the Signal-to-Background ratio, SBR, on the extraction of spectral signature and on localisation precision. Here we define SBR as

$$\mathrm{SBR} = (I_{\mathrm{sig}} + I_{\mathrm{bkg}})/I_{\mathrm{bkg}}. \tag{S20}$$

Thus for a given number of photons $I_{\mathrm{sig}}$ given out by a puncta, each pixel has a background value added based on determining the PSF area ($3\sigma_{\mathrm{PSF}}$), and dividing $I_{\mathrm{bkg}}$ by this area. Explicitly, this means that the per-pixel number of background photons is

$$I_{\mathrm{bkg,per\ pixel}} = I_{\mathrm{sig}}/((\mathrm{SBR}-1)\times 3\sigma_{\mathrm{PSF}}^2). \tag{S21}$$

To give a practical handle on this, at an $I_{\mathrm{sig}}$ of 500 and an SBR of 1.1 for an ATTO 488 emitter, this results in 126.9 background photons per pixel. For the same SBR and an ATTO 647N, this results in 75.3 photons per pixel.

We performed simulations using camera calibration values drawn from a Ximea MC050CG-SY sCMOS sensor (camera characterisation details in table S2), a NA of 1.49, a pixel size of 69 nm, 25 varying background levels from SBR=1.1 to SBR=100, and a Gaussian pre-smoothing kernel with $\sigma$ = 1.5 pixels applied before fitting. For each condition, a two-dimensional Gaussian point-spread function (for full details of image simulation, see S1) was placed at a position drawn uniformly within the central pixel of a simulated sensor patch and either fitted by Levenberg-Marquardt optimisation (for full details of fitting see S3). This process was repeated 10,000 times (bootstraps) per photon level to obtain statistical estimates of spatial precision ($\sigma_{\mathrm{xy}}$, the root-mean-square localisation error averaged over x and y) and pixel QE precision ($\sigma_{\mathrm{pixel\ QE}}$, the standard deviation of the Euclidean distance in pixel QE space from the known spectral fingerprint of the dye). Three dyes spanning the visible spectrum were evaluated—ATTO 488, ATTO 565, and ATTO 647N—across 200 photon levels spaced logarithmically between 500 and 50,000 photons.

We highlight the effect of SBR is similar to that of decreasing photon level in Fig. S34. At high background level or at low photon values, the distribution of fit results is considerably broader than at high SBR, high photon values. Crucially however the effect of background is the same as that of decreasing the number of photons received by the detector (*i.e.* a broadening of the spectral fingerprint). An increasing background is accounted for by the analysis routine and does not introduce any bias in the spectral fingerprint estimation. The effect of SBR and number of photons impinging on the detector for the three dyes simulated is fully shown on the localisation precision in Fig. S35, and on the spectral fingerprint estimation precision in Fig. S36.

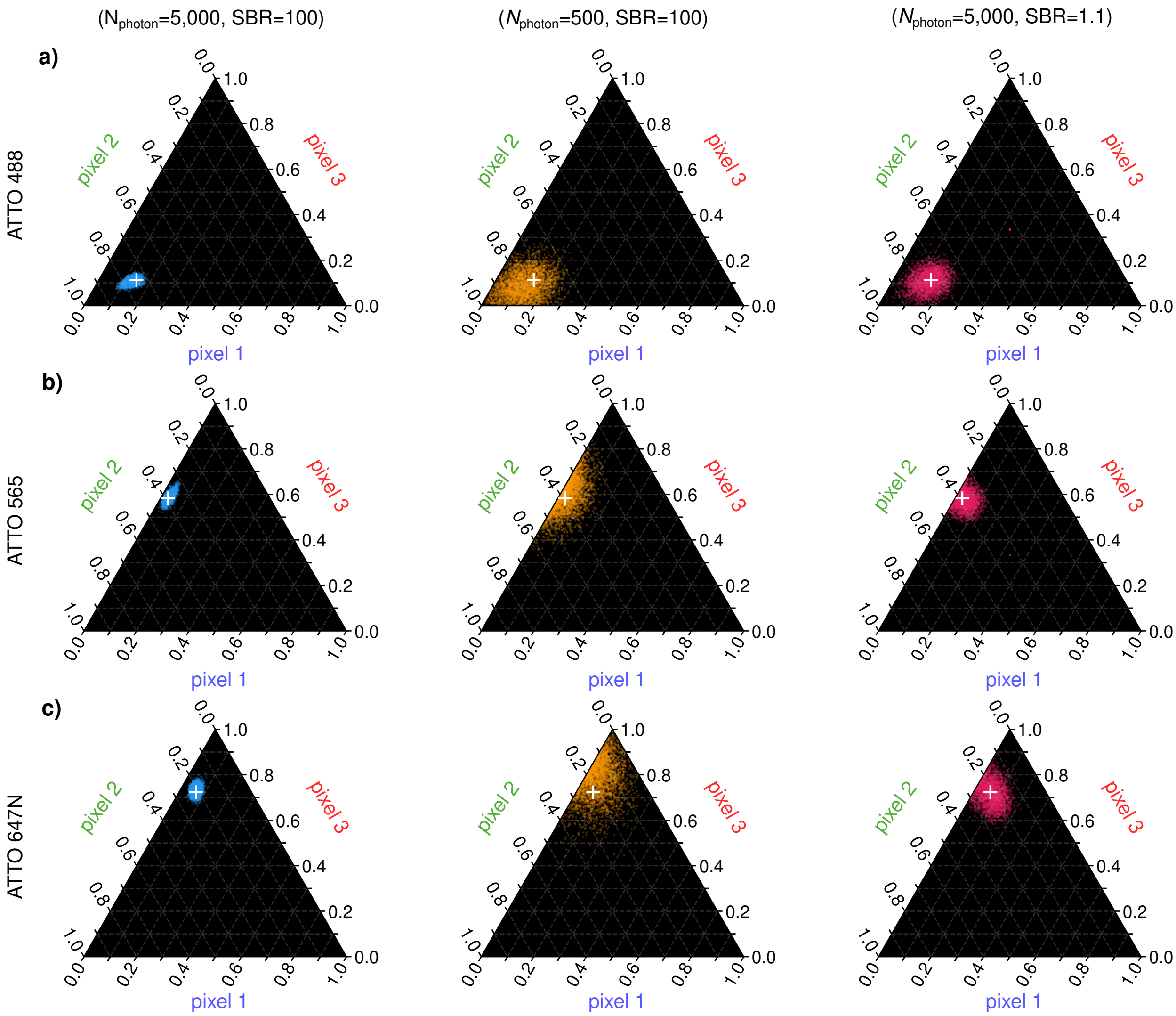


**Fig. S34. The effect of SBR on the extraction of spectral fingerprints. a)** The effect of changing signal (middle panel) and SBR (rightmost panel) on the spectral fingerprint of ATTO 488 molecules. **b)** The effect of changing signal (middle panel) and SBR (rightmost panel) on the spectral fingerprint of ATTO 565 molecules. **c)** The effect of changing signal (middle panel) and SBR (rightmost panel) on the spectral fingerprint of ATTO 647N molecules.

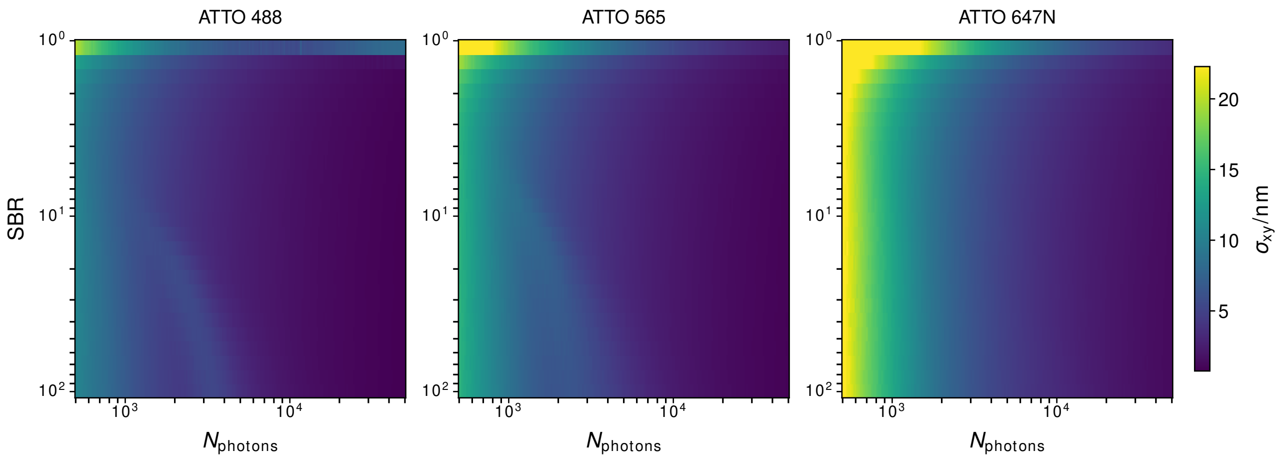


**Fig. S35. Influence of SBR and $N_{\mathrm{photon}}$ on localisation precision.**

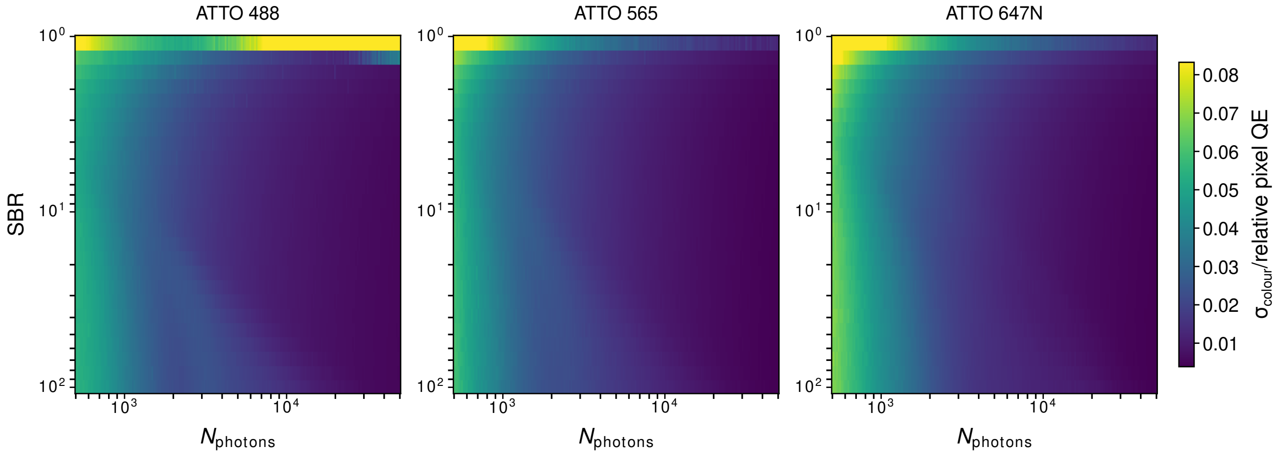


**Fig. S36. Influence of SBR and $N_{\mathrm{photon}}$ on spectral fingerprint estimation precision.**

#### Supplementary Note S10: Limitations of S$^3$M

The limitations of S$^3$M can be placed in two categories: computational time of the analysis, and physical restrictions of the detector. We note that the physical restrictions of the detector comprise the majority of the limitations. The first limitation is the read noise of commercially available Bayer patterned detectors. Generally, these industrially patterned CMOS detectors have a higher read noise (ranging from 0.85–2.49 RMS $e^-$) (S18) than commonly used sCMOS detectors such as the Kinetix (Teledyne Photometrics) which has a read noise of 0.7 RMS $e^-$. As a result, even without the inevitable losses from the Bayer pattern, these industrial CMOS detectors will have a lower SNR than most conventional sCMOS detectors.

A more subtle limitation is in the effective pixel size of the image needed for S$^3$M. S$^3$M requires the PSF of the molecule to be adequately sampled across the pattern of the detector (see a simulation of the effect of pixel size on localisation and spectral fingerprint precision, Fig. S37), meaning more pixels must be used to fit the spectral fingerprint and location simultaneously. This partially reduces SNR by spreading photons over more pixels, as well as trivially reducing the effective area that can be used for imaging.

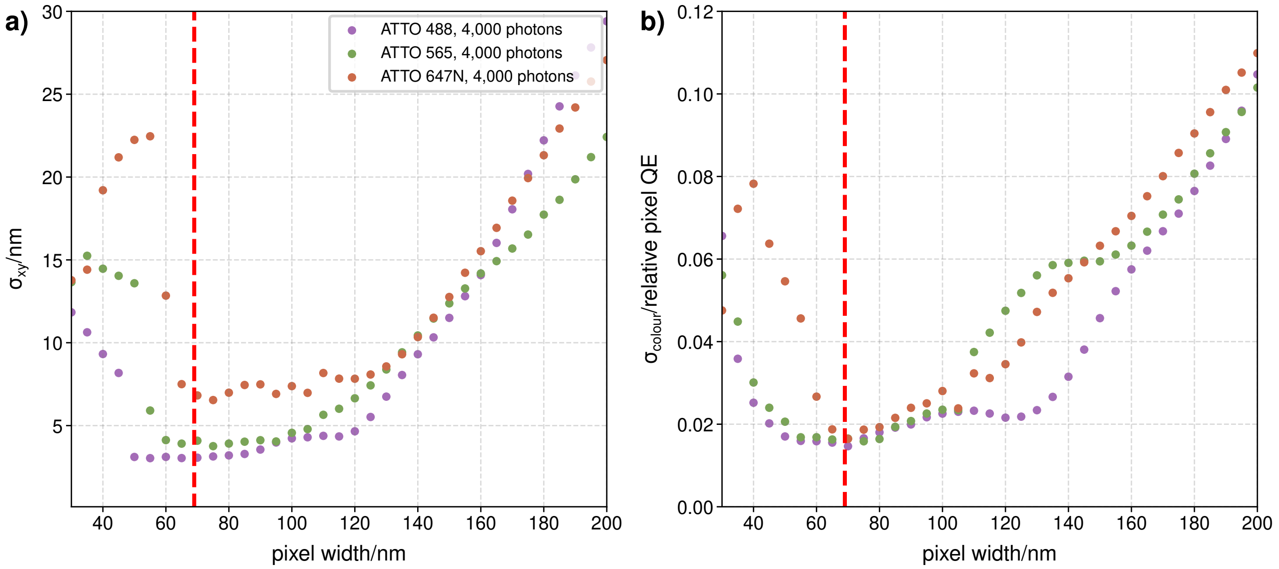


**Fig. S37. Effect of pixel size on S$^3$M precision. a)** Effect of pixel width on localisation precision for three fluorophores emitting 4,000 photons. **b)** Effect of pixel width on spectral fingerprint precision for three fluorophores emitting 4,000 photons.

Moreover, because each PSF includes a spectral fingerprint, the ability to accurately identify molecules is reduced when PSFs overlap, reducing the overall density of molecules that can be used with S$^3$M. We have quantified this by simulating 1,000 512×512 images at varying emitter densities, with the camera parameters of the Ximea camera (table S2). We did this for three dyes (ATTO 488, ATTO 565, and ATTO 647N) at three different photon levels (1,000 photons per punctum, 4,000 photons per punctum, and 10,000 photons per punctum). Exemplar simulations are shown in Figures S38–S40.

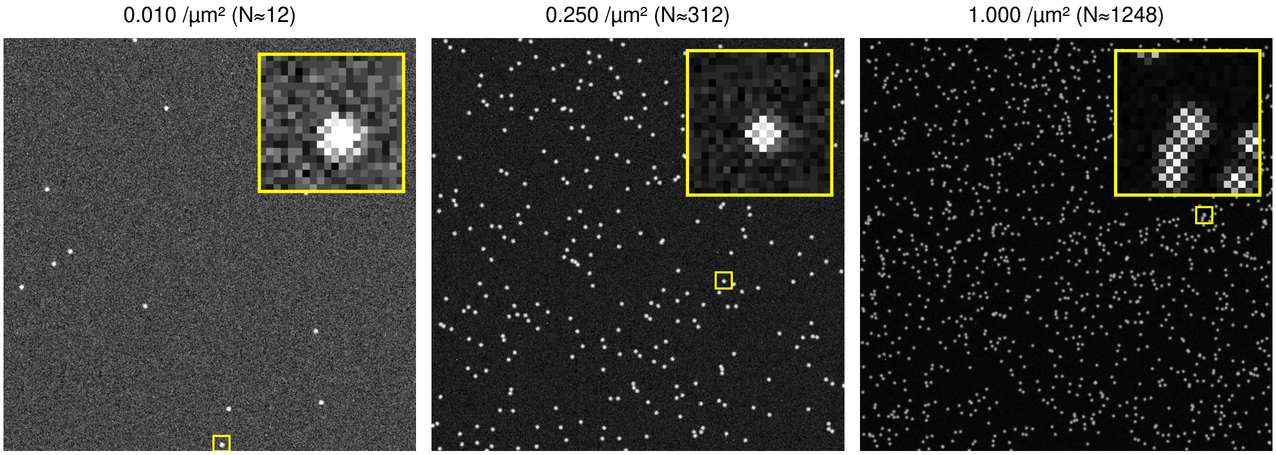


**Fig. S38. Exemplar simulated ATTO 488 molecules, emitting 4,000 photons, at varying densities.**

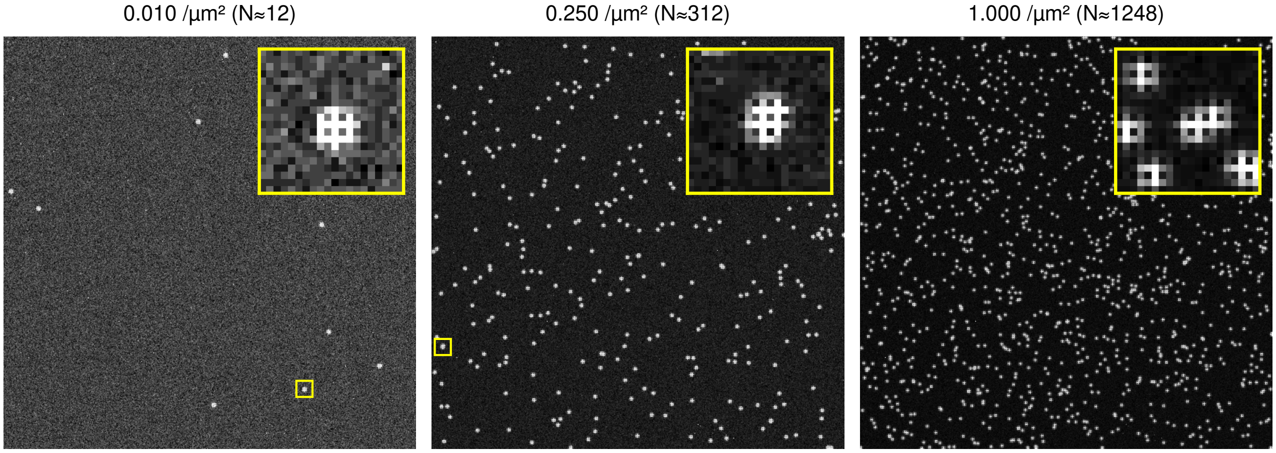


**Fig. S39. Exemplar simulated ATTO 565 molecules, emitting 4,000 photons, at varying densities.**

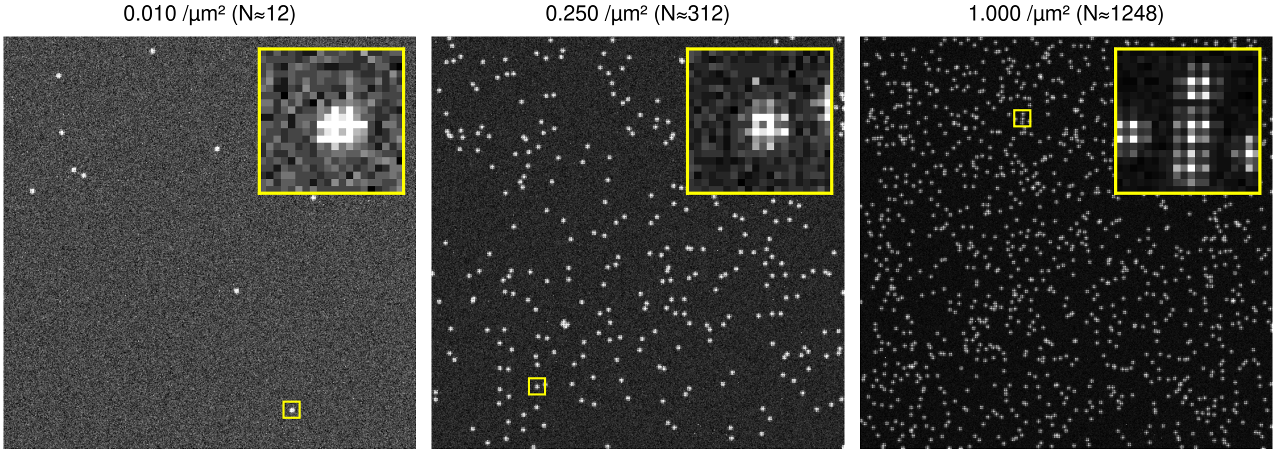


**Fig. S40. Exemplar simulated ATTO 647N molecules, emitting 4,000 photons, at varying densities.**

We compared the Bayer patterned detector and a detector with the same pixel size but the Prime 95B's QE at each pixel (patterned vs unpatterned). We then quantified how many of these puncta were satisfactorily fit (fit returning a ground truth position within the PSF width) as a function of emitter density. These data are shown in Fig. S41. We plot the Jaccard index,

$$\frac{\mathrm{TP}}{\mathrm{TP}+\mathrm{FP}+\mathrm{FN}}$$

where TP stands for true positive, FP for false positive, and FN for false negative. We also plot the positive predictive value (PPV),

$$\frac{\mathrm{TP}}{\mathrm{TP}+\mathrm{FP}}$$

and the Recall

$$\frac{\mathrm{TP}}{\mathrm{TP}+\mathrm{FN}}.$$

Each of these parameters quantifies how well S$^3$M responds to emitter density. As can be seen by inspection of these parameters, the patterning of the detector means S$^3$M is outperformed at high densities by unpatterned detectors, limiting the density range of molecules. However, the PPV for all dyes at all photon ranges is above 90% even at high densities, and the Recall above 60% below a $\rho_{\mathrm{loc}}$ of 0.4/µm$^{-2}$. These are within the practical ranges of single-molecule and super-resolution experiments,(11) and thus this means S$^3$M is useable for practical experiments. Importantly, the effect of density is only on how many puncta are successfully found and analysed—the spectral fingerprints recovered from the analyses (Fig. S42) show no density-related effects.

Additionally, the Bayer pattern does not respond linearly with emission wavelength, unlike spectral measurements with a diffraction grating(12) or a prism.(13) As seen in Fig. 1a, there are regions where small changes in the wavelength result in very different relative QE values. As a result, chromophores near these variable regions have very broad spectral fingerprints, as seen in Fig. 3f with Qdot 585, which despite a similar spectral width has a much broader spectral fingerprint than Qdot 655 (Fig. S43).

The most obvious drawback of S$^3$M is that due to the spectral sensing enabled by the separate pixel QEs, inevitably this leads to a loss of photoelectrons generated by the detector, and thus a decrease in localisation precision. To give an example of the loss in photoelectrons generated by the separate and lower pixel QEs, for ATTO 488, 565 and 647N molecules the effective QE of the Ximea MC050CG-SY is 0.30, 0.24, and 0.16 respectively. To compare to the Prime95B, the effective QEs of the same dyes are 0.79, 0.77, and 0.67. Thus the differing pixel QEs introduce a 2.63–4.19$\times$ loss in photoelectrons generated, corresponding to 40–50% drop in SNR. To compare achievable localisation precision between an S$^3$M-compatible camera and a typical single-molecule camera, we performed 10,000-repeat Monte Carlo bootstrap simulations at each of 20 photon levels logarithmically spaced between 200 and 20,000 detected photons for three dyes covering the visible emission spectrum (ATTO 488, ATTO 565, ATTO 647N), at a fixed background of 10 photons/pixel and NA = 1.49. We did this for a simulation using the Ximea camera gain, read noise, QE *etc.*, and for a simulation of a Prime95B camera. Both cameras were simulated with an identical 69 nm pixel size. The Ximea simulation used measured camera parameters (Table S2); the Prime95B was instead modelled from its manufacturer-specified quantum-efficiency curve, gain (2 e$^-$/ADU) and read noise (1.6 e$^-$ RMS). For each bootstrap repeat, a synthetic emitter was generated with Poisson-distributed photon and background shot noise and camera-appropriate read noise, then localised by weighted least-squares Gaussian fitting, with localisation precision ($\sigma_{\mathrm{xy}}$) reported as the resulting positional scatter relative to the known ground-truth position.

Results from these simulations are shown in Fig. S44. As expected, the Prime95B has superior localisation precision for all dyes tested. The vast majority of this is explained by the improved QE—Fig. S44b shows that the factor of improvement is close to the absolute QE difference calculated for the dyes in question.

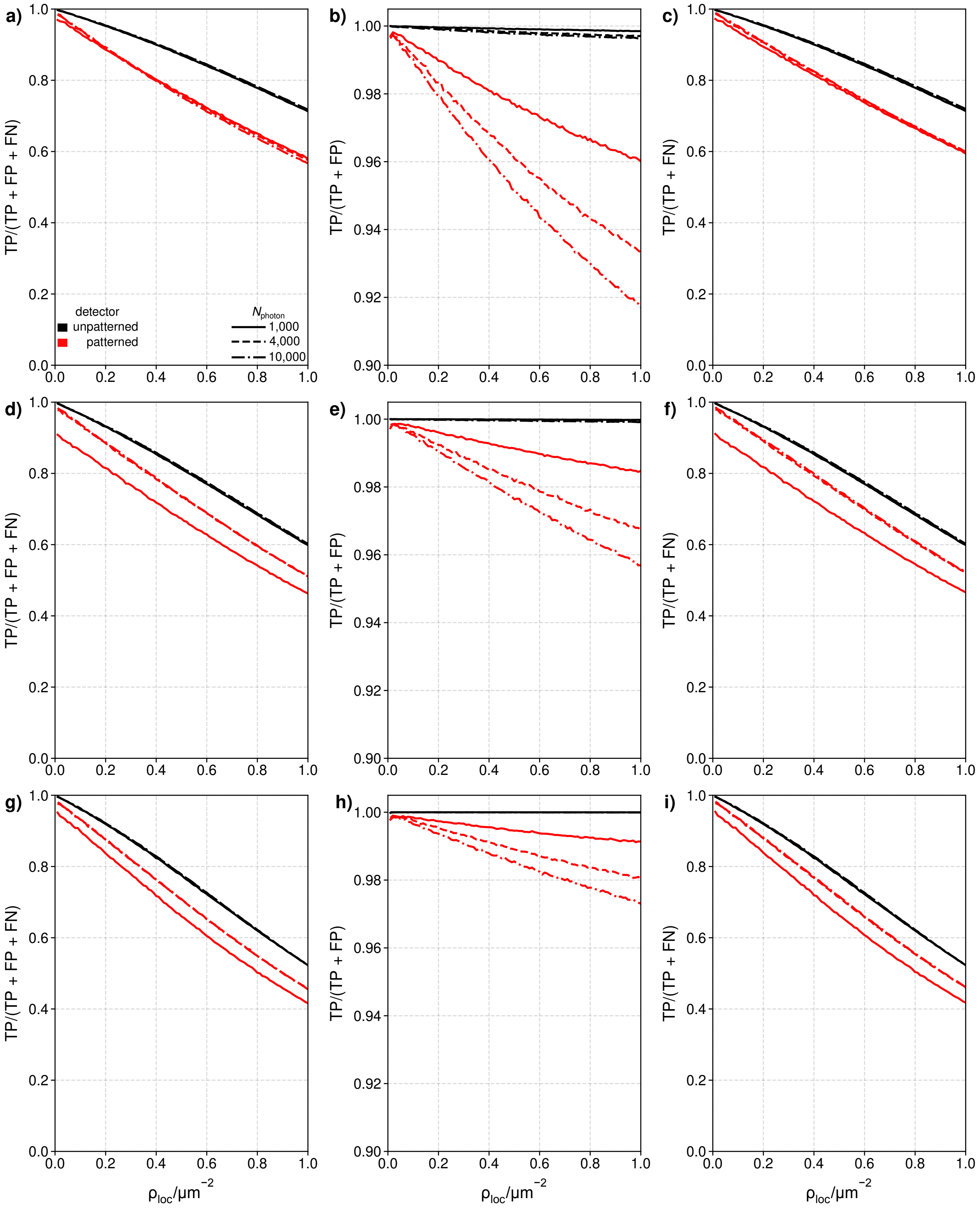


**Fig. S41. The effect of emitter density.** **a)** Jaccard index, **b)** PPV, and **c)** Recall for ATTO 488 at a range of photon values. **d)** Jaccard index, **e)** PPV, and **f)** Recall for ATTO 565 at a range of photon values. **g)** Jaccard index, **h)** PPV, and **i)** Recall for ATTO 647N at a range of photon values.

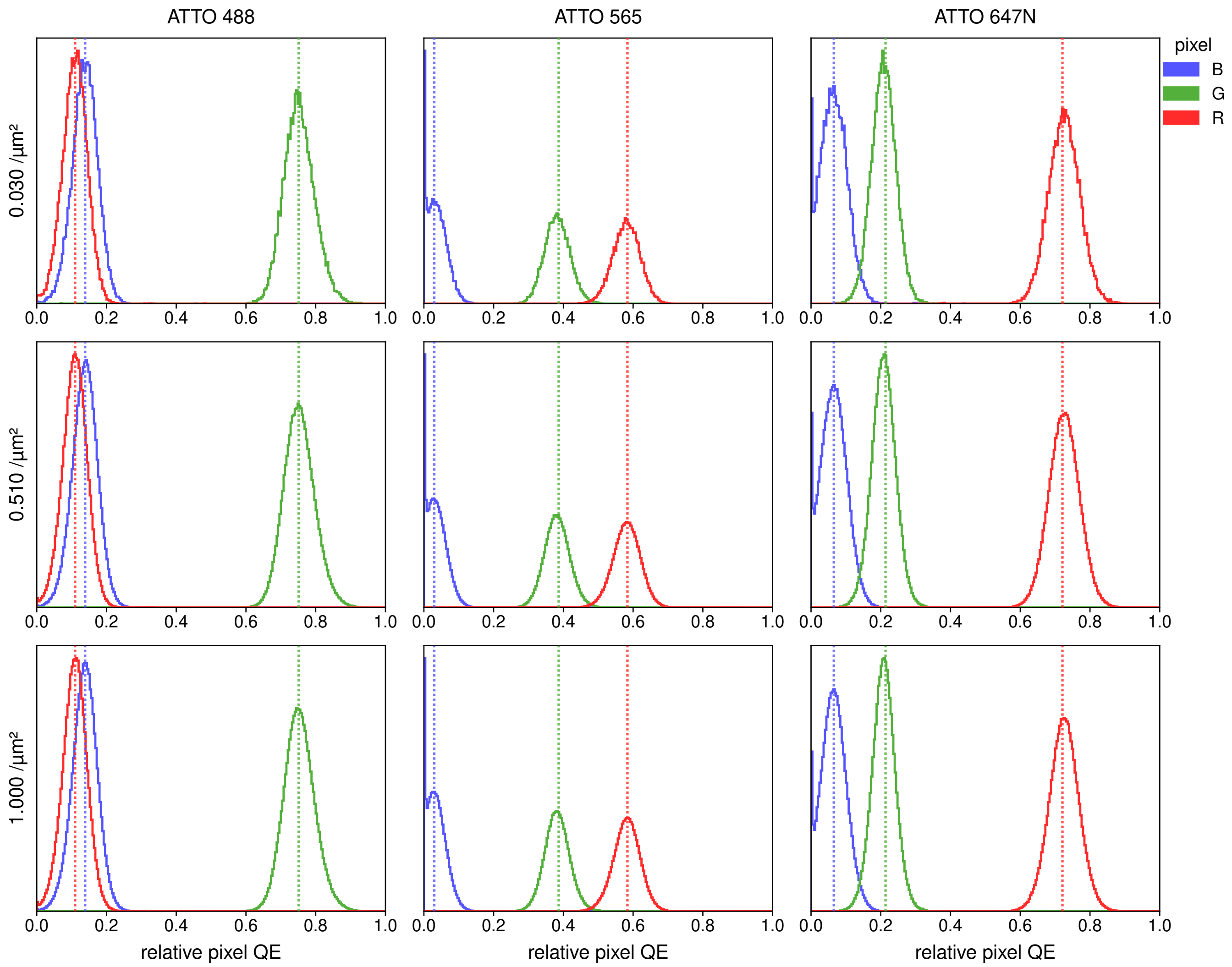


**Fig. S42. The effect of emitter density on spectral fingerprint.** Histograms of spectral fingerprints for ATTO 488, ATTO 565 and ATTO 647N at 1,000 photons show that density does not affect the spectral fingerprint.

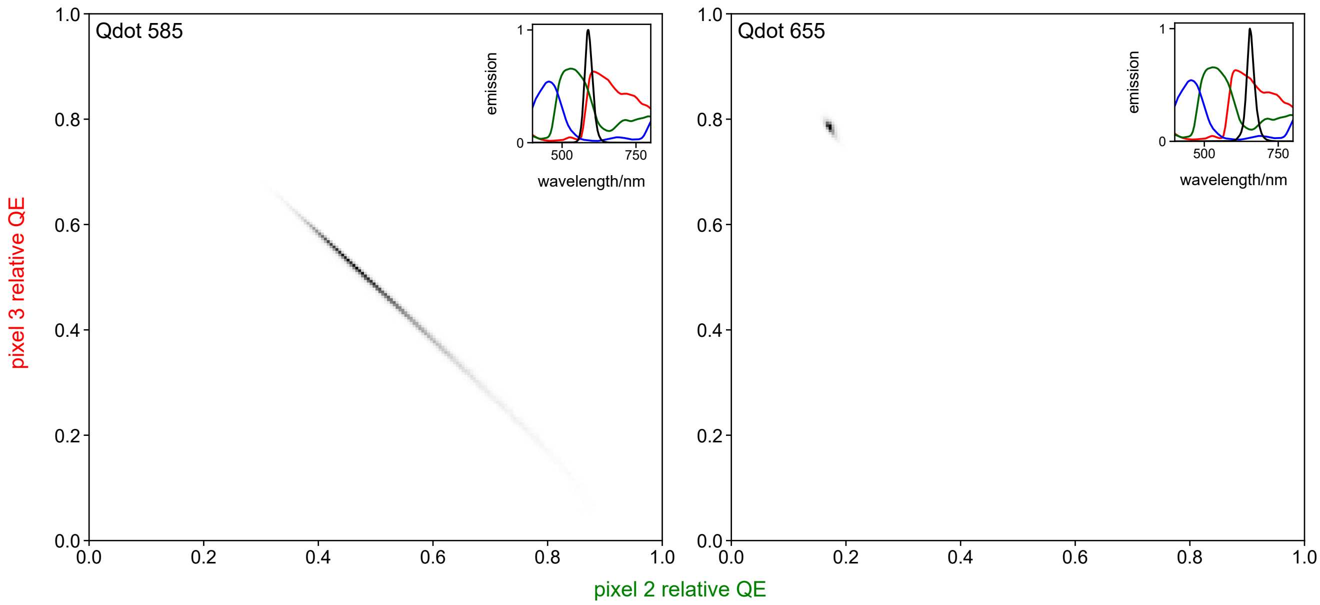


**Fig. S43. Spectral fingerprint width as a function of spectra.** 2D histograms of the mean pixel 2 and pixel 3 relative QEs for Qdot 585 and Qdot 655. Insets show their emission spectra overlaid with the spectral response of the Bayer pattern.

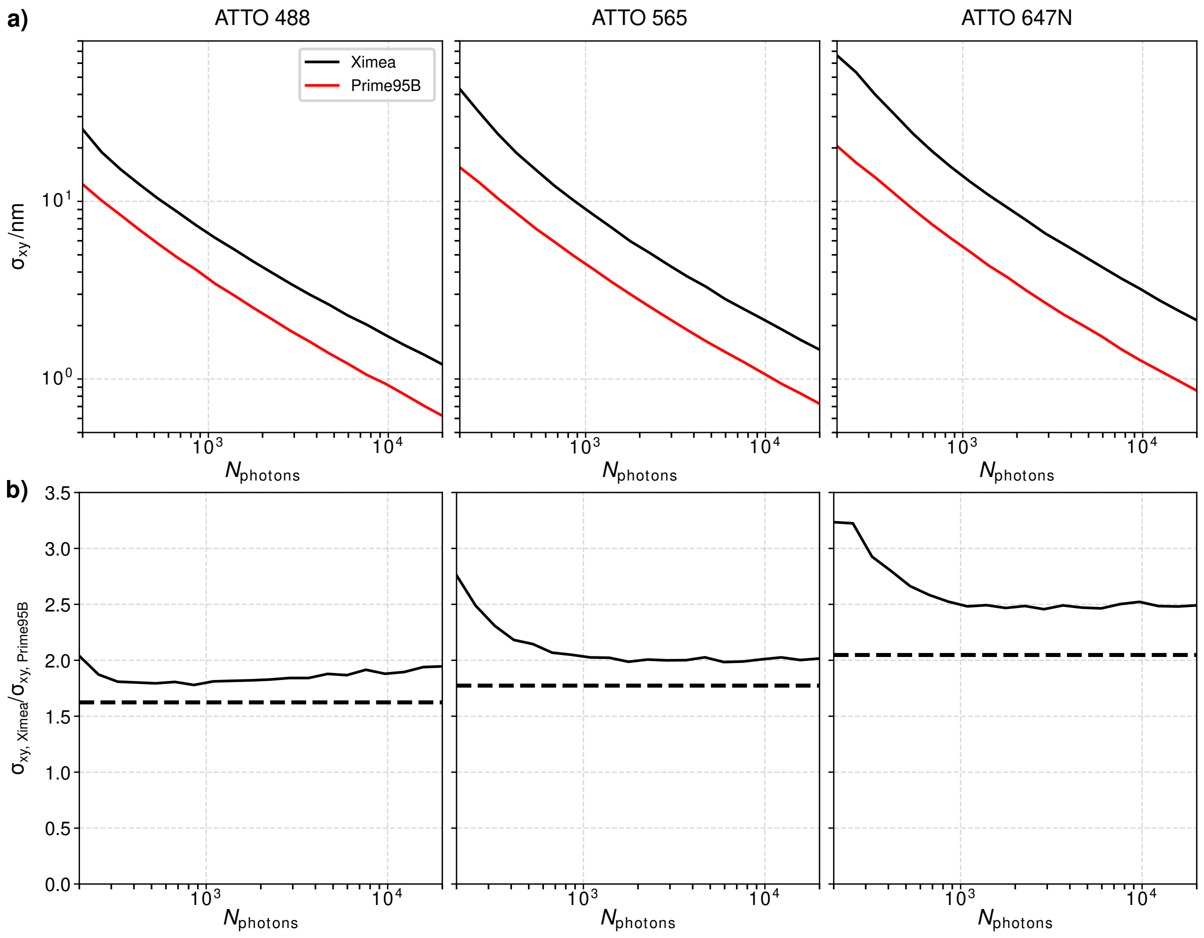


**Fig. S44. Direct comparison between Prime95B and Ximea detector. a)** Localisation precision, $\sigma_{\mathrm{xy}}$, for the Prime95B and the Ximea camera for the dyes simulated here. **b)** A comparison of the improvement in localisation precision between the Prime95B and the Ximea detector. The dashed black line is the improvement expected from QE differences alone.

Finally, we compare $S^3M$ to a number of spectral-sensing single-molecule techniques and compare their reported performance (table S1). Perhaps the largest drawback is the lack of direct spectral sensing, which ultimately results in a worse spectral precision when compared to methods that use a dispersive element. However, despite a moderate loss in photons, $S^3M$ exceeds in usable detector area, demonstrated colours per detector, and localisation precision, all while requiring low experimental complexity.

| Method | $\eta_{loc}$ in % [a] | Usable FOV per Detector (%) | Demonstrated Colours per Detector | $\sigma_{loc}$ per 2,000 photons (nm)[b] | $\sigma_{\lambda}$ per 2,000 photons (nm)[b] | Direct Spectral Sensing? | Experimental Complexity[c] |
|---|---|---|---|---|---|---|---|
| $S^3M$ | 31–58[d] | 100 | 6 | 10 | 10–40[e] | No | Low |
| sPAINT/SR-STORM(12, 13) | ≈40 | ≈33 | 4 | ≈18 | ≈4 | Yes | Moderate |
| PSF Engineering(14, 15) | 87 | 100 | 4 | $> 5$[f] | None | No | High |
| SDsSMLM(16) | ≈47 | ≈50 | 2 | ≈10 | $< 1$ | Yes | High |
| Channel Splitting | ∼95% | 25–50%[g] | 2–4[g] | ≈5[h](17) | None | No | Low |
| Vortex FLFM (18) | ≈95 | ≈33 | 4 | 10–20 | 2 | Yes | High |

**Table S1.** Comparison between $S^3M$ and other spectral sensing single-molecule techniques.

a Percentage of light that can be used for localisation of the total light collected.

b 2,000 photons incident on the detector.

c Experimental complexity includes day-to-day alignment, calibration of spectral sensing elements, and colour registration. d Based on simulated values of a range of single-molecule dyes with and without a Bayer mask.

e Based on forward model of Nile Red, value is wavelength dependent. See S17 for a detailed explanation.

f Based on a simulation of 5,000–10,000 photons.

g This depends on whether channels are split onto the same (decreases usable FOV but increases colours per detector) or multiple detectors (decreases colours per detector but increases usable FOV).

h Assumed to be equivalent to a typical sCMOS detector.

#### Supplementary Note S11: Effect of Motion Blur on Extraction of Spectral Signature

When performing single-molecule or single-particle tracking (SPT), motion blur is always inherently present.(19) This blur will become larger at longer detector exposure times and/or faster molecular motion, and may impact the localisation precision in SPT data. Unlike in techniques such as sPAINT,(12) the spatial and spectral estimations cannot be decoupled in $S^3M$. Thus, we anticipate that, at high motion blur, this will degrade both the precision of estimating the molecular position and spectral signature. The effect of motion blur on the $S^3M$ PSF is shown in Fig. S45.

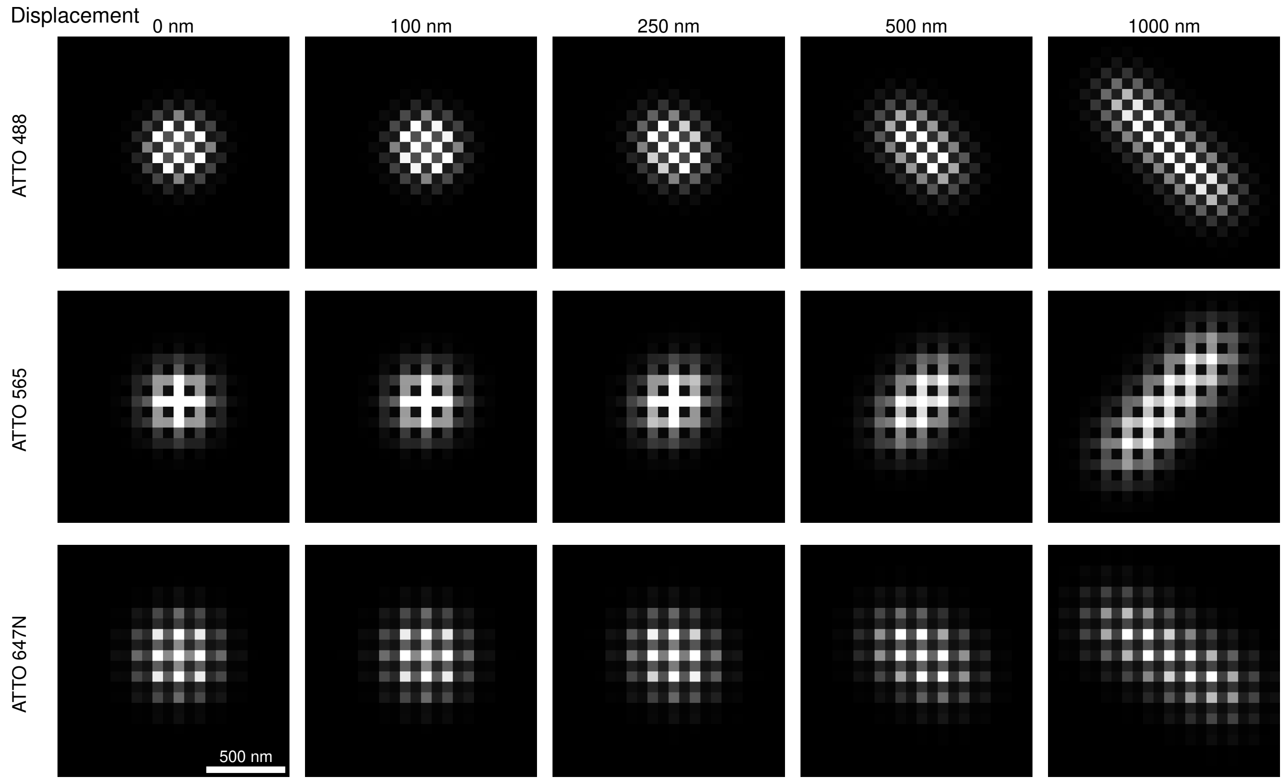


**Fig. S45. The effect of motion blur on the $S^3M$ PSF.** Exemplar simulations (10,000 photons per molecule, no noise) on a range of displacements for 3 dyes.

We thus performed simulations at a range of displacements, corresponding to a range of diffusion coefficients at the 50 ms camera exposure time we used in experiment, for a range of photon values. Motion blur was modelled as a molecule moving with constant speed $v$ in a uniformly random azimuthal direction throughout the detector exposure time $T = 50\,\text{ms}$. The time-integrated PSF was approximated by summing the Gaussian intensity distributions at 51 equally-spaced positions along the trajectory, yielding a per-frame displacement $d = vT$. Simulations were performed for three fluorophores (ATTO 488, ATTO 565, and ATTO 647N) across 50 velocities linearly spaced from 0 to $10{,}000\,\text{nm}\,\text{s}^{-1}$ (displacements 0–1,000 nm), and over 100 photon levels geometrically spaced between 500 and 20,000 detected photons, with 10,000 bootstrap realisations per condition and 5 background photoelectrons per pixel. 500 photons was chosen as the low end of the simulations as we typically reject localisation fits below 500 photons. Camera noise was modelled using the measured gain, offset, variance, read-noise, and relative quantum efficiency maps of the Ximea camera. Because motion blur elongates the PSF asymmetrically, we used elliptical Gaussian fitting (recovering independent $\sigma_x$, $\sigma_y$, and orientation $\theta$ per localisation) to directly compare to the analysis of the SPT data. From each set of localisations we extracted two metrics: the lateral localisation precision $\sigma_{xy} = \sqrt{(\sigma^2_{\epsilon_x} + \sigma^2_{\epsilon_y})/2}$, and the spectral signature precision $\sigma_{\text{pixel QE}}$. We then analysed these data to determine how these metrics degraded as a function of motion blur. Exemplar fits for each dye as a function of increasing motion blur are shown in Figures S46–S48.

As can be seen in Fig. S49 and S50 respectively, the localisation precision does degrade as a function of increasing motion blur. At high levels of motion blur, the localisation precision may degrade by an order of magnitude, as one would expect. This corresponds to diffusion coefficients in considerable excess of what we measured in Fig. 4 however. The typical range of diffusion coefficients measured in this paper would give degradations in localisation precision of a more manageable $\sim 3\times$. For the photon ranges of the dyes in Fig. 4 (median photons per frame $\pm$ median absolute deviation $770 \pm 274$), this leads to localisation precision on the order of 10–40 nm. As Fig. S50 shows, the $\sigma_{\text{pixel QE}}$ is less sensitive to motion blur, changing by a factor of $\sim 2$ across the displacement range simulated here. This leads to pixel QE precision on the order of 0.1–0.2 per frame in the diffusion data shown in Fig. 4.

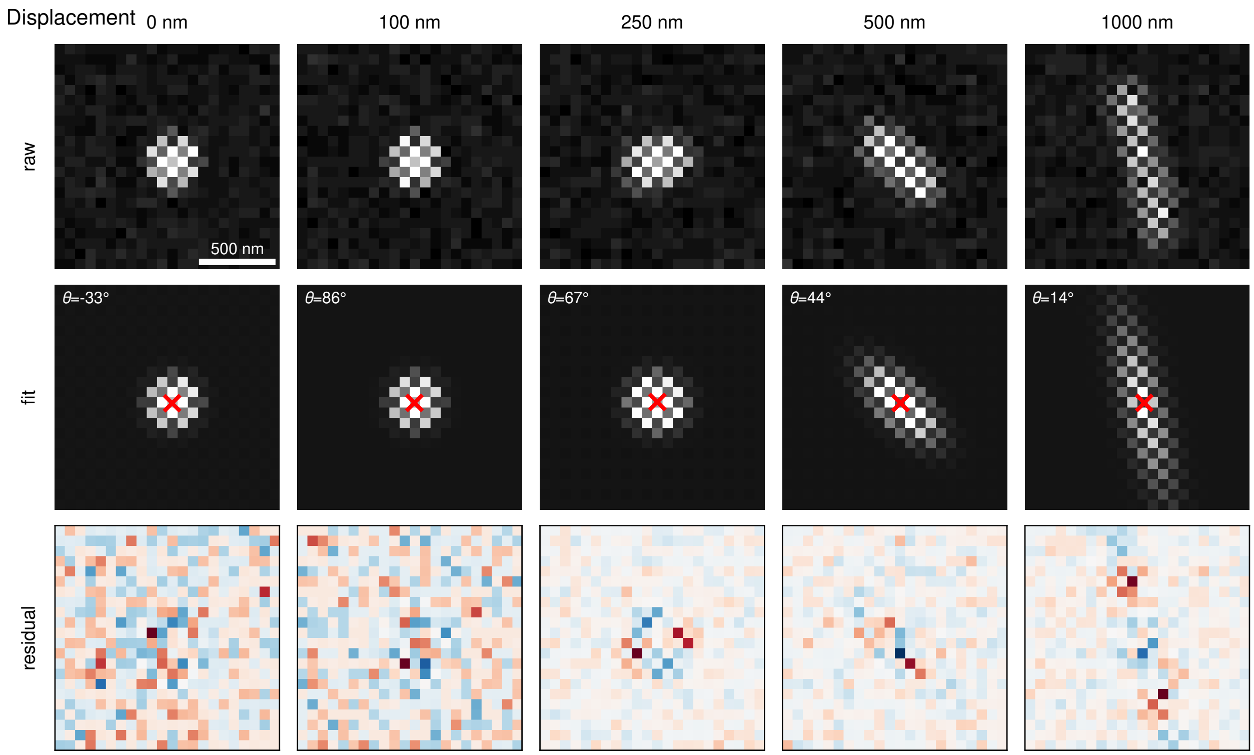


**Fig. S46. The effect of motion blur on the S$^3$M fitting.** Exemplar fits (4,000 photons per molecule, 5 background photons) on a range of displacements for ATTO 488.

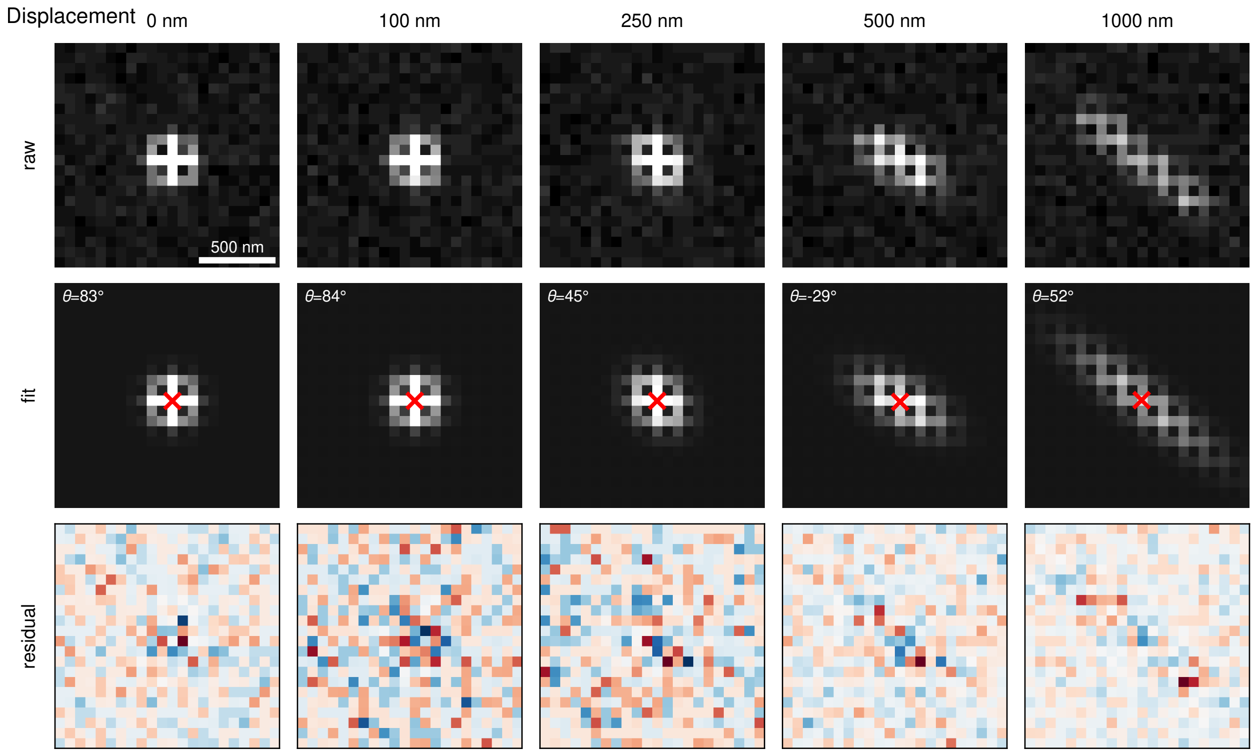


**Fig. S47. The effect of motion blur on the S$^3$M fitting.** Exemplar fits (4,000 photons per molecule, 5 background photons) on a range of displacements for ATTO 565.

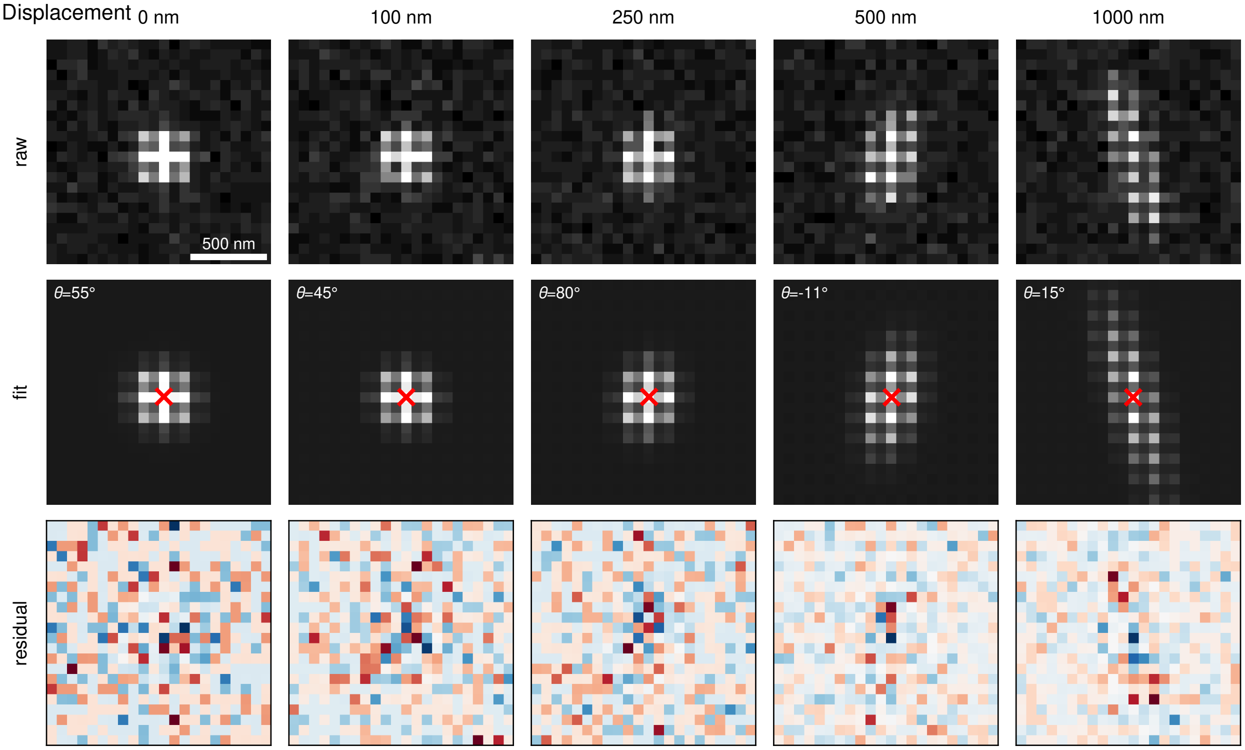


**Fig. S48. The effect of motion blur on the S$^3$M fitting.** Exemplar fits (4,000 photons per molecule, 5 background photons) on a range of displacements for ATTO 647N.

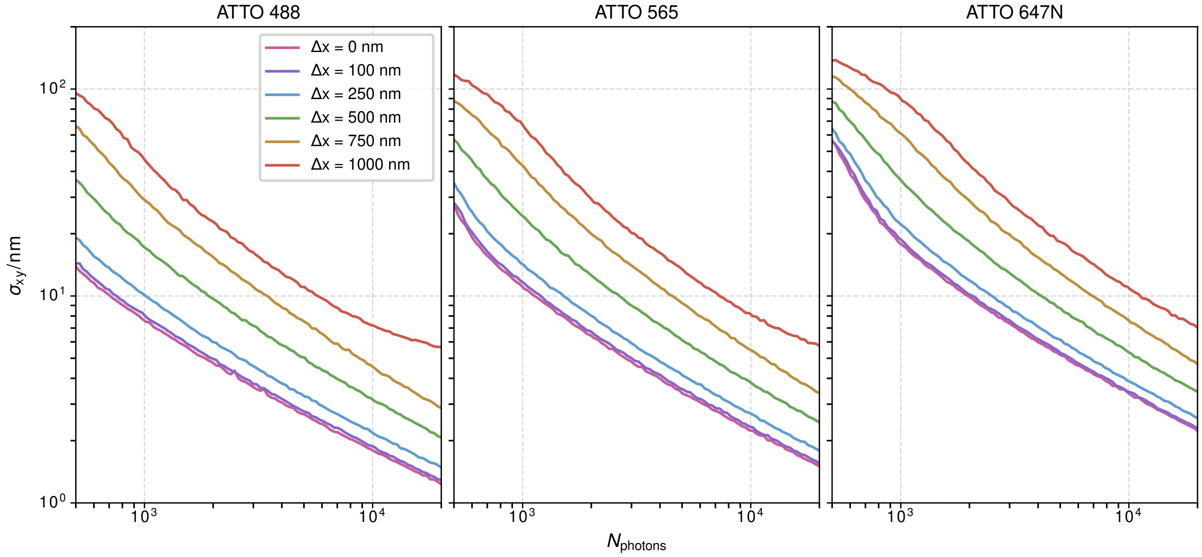


**Fig. S49. The effect of motion blur on localisation precision in S$^3$M.** 1D displacements in a single frame correspond to diffusion coefficients, assuming a 50 ms exposure time, of 0, 0.03, 0.15, 0.60, 1.43 and 2.5 μm$^2$/s respectively.

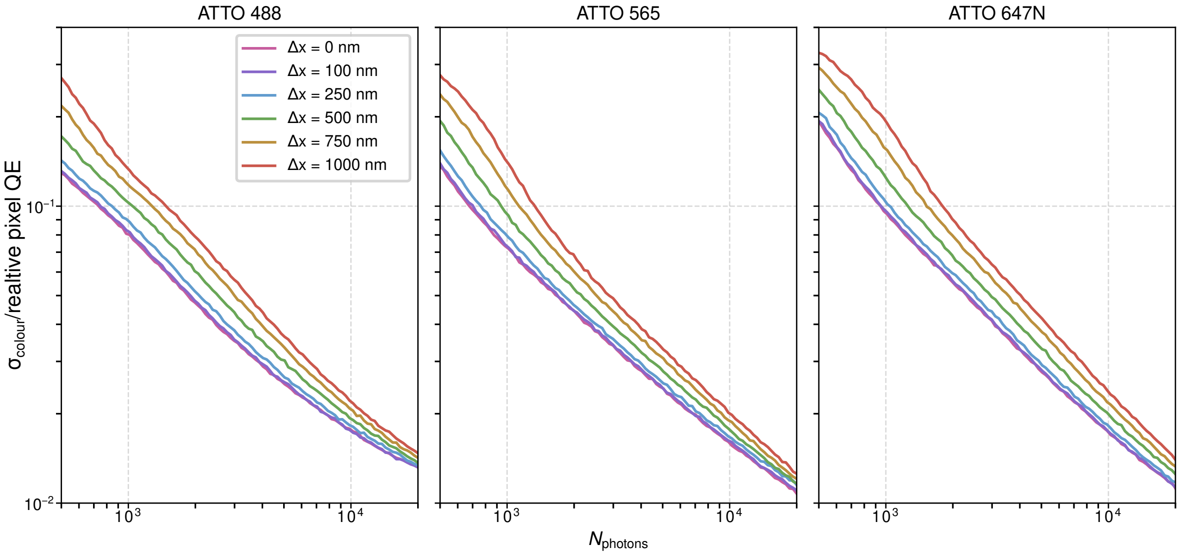


**Fig. S50. The effect of motion blur on pixel QE precision in S$^3$M.** 1D displacements in a single frame correspond to diffusion coefficients, assuming a 50 ms exposure time, of 0, 0.03, 0.15, 0.60, 1.43 and 2.5 μm$^2$/s respectively.

#### Supplementary Note S12: Effect of Z Defocus on Extraction of Spectral Signature

The distance of an emitter from the focal plane of the objective will impact the shape of the PSF. In order to test the effect of this on the localisation precision and the accurate extraction of spectral signature, we imaged Tetraspeck™ beads, embedded in 1% PVA, at a range of Z positions relative to the objective focal plane (see Table S6 for power density and filters). Specifically, this involved imaging for 100 frames per Z plane, and fitting each of these frames, with a circular 2D Gaussian (see section S3) to determine X, Y, and pixel QE positions per emitter. In total 87 beads were used for this analysis. Exemplar emitters are shown in Fig. S51. We then used these data to determine, per emitter, the localisation precision—comparing individual localisation to the mean of the 100 localisations at each Z plane. These results are shown in Fig. S52, and show that the precision of both the position estimation and pixel QE estimation degrade as the emitter is at larger distance from the focal plane, as expected.(20) In the case of localisation precision, this is a 4.5-fold degradation over 1 μm. In the case of pixel QE (*i.e.* spectral signature estimation), this is a 2-fold degradation.

We also highlight that Z defocus causes a bias in the estimation of spectral signature—the different pixel QE estimations at changing focus are shown in Fig. S53–S56. At relatively small defocus values of ± 200 nm, there is little effect (Fig. S53 and S54). At larger defocus values (Fig. S55 and S56) the estimate significantly differs from the value at z=0 nm. Thus at large values of defocus, there is a likelihood that an emitter would be misclassified (*i.e.* the spectral signature value would overlap with that of another emitter). At practical ranges of defocus (±200 nm) that are typically accessed in TIRF microscopy, and in SMLM, this has little effect on the spectral signature. S$^3$M is thus robust to 400 nm of defocus.

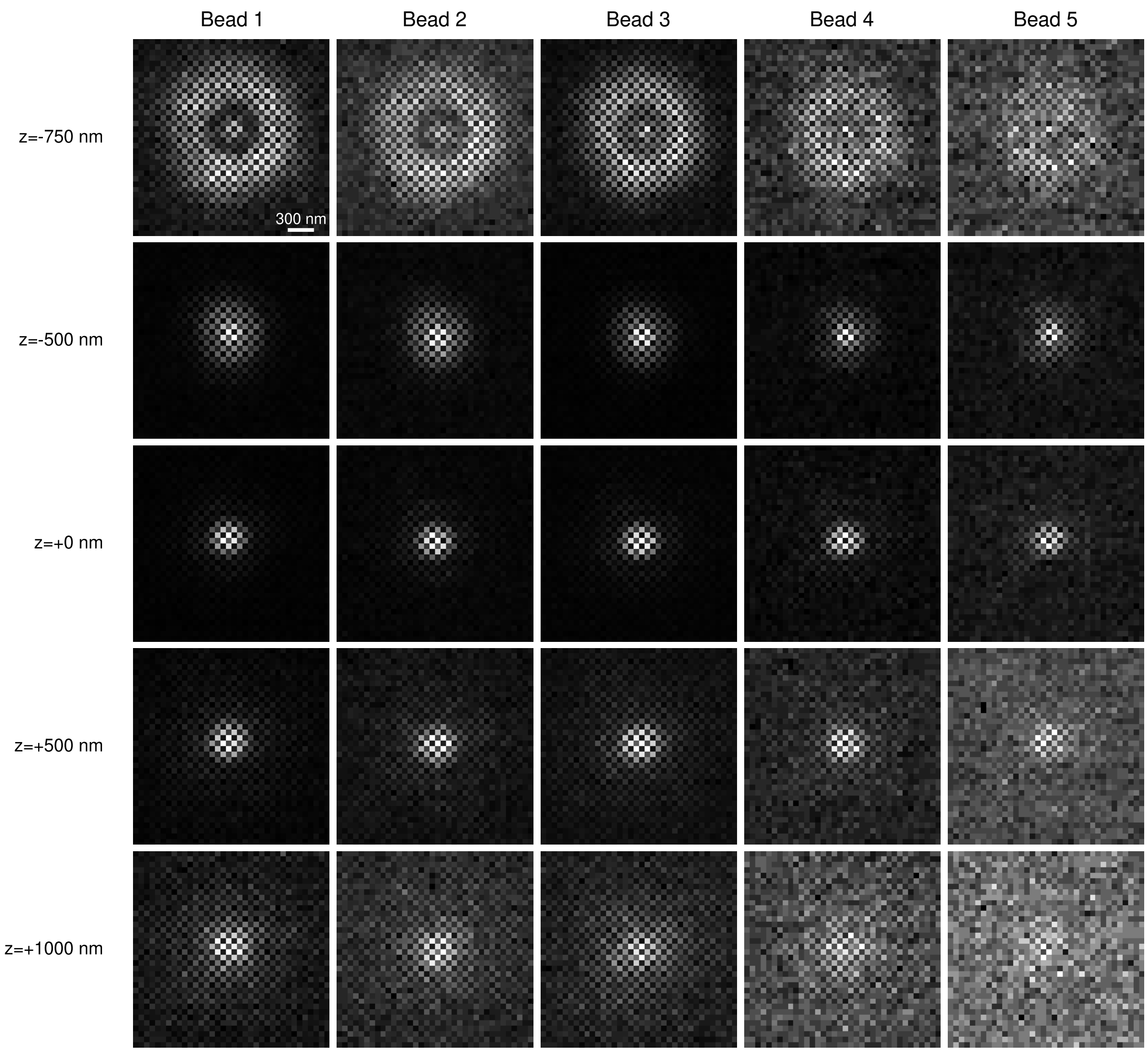


**Fig. S51. Exemplar emitters at varying levels of focus.** Five Tetraspeck™ beads imaged at varying distances from the focal plane, showing the effect of defocus on the resultant spatial pattern.

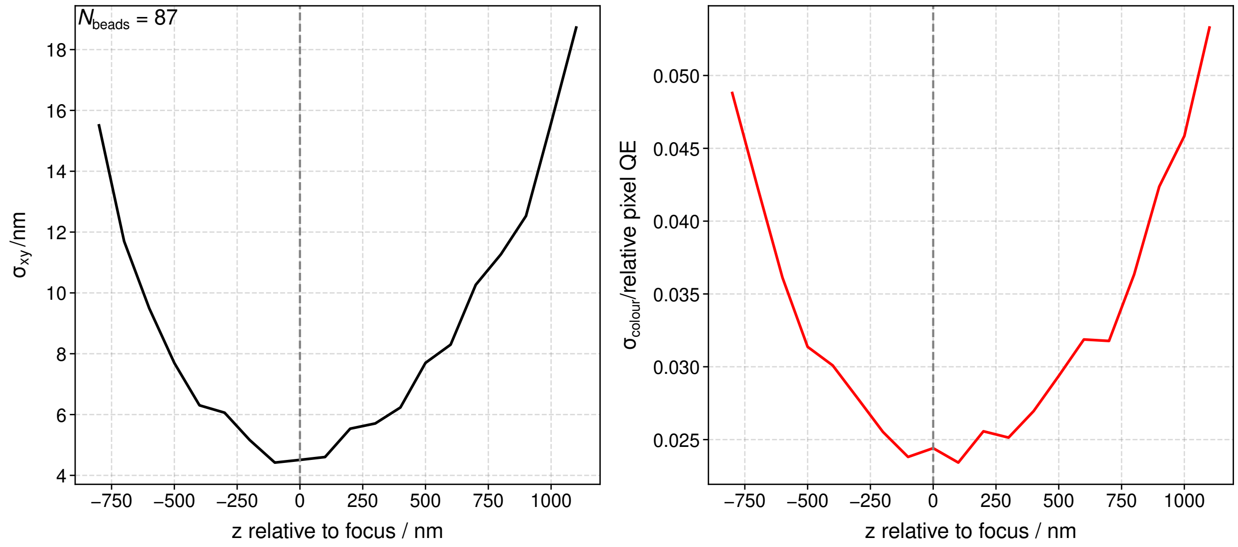


**Fig. S52. The effect of Z defocus on localisation and spectral signature precision.**

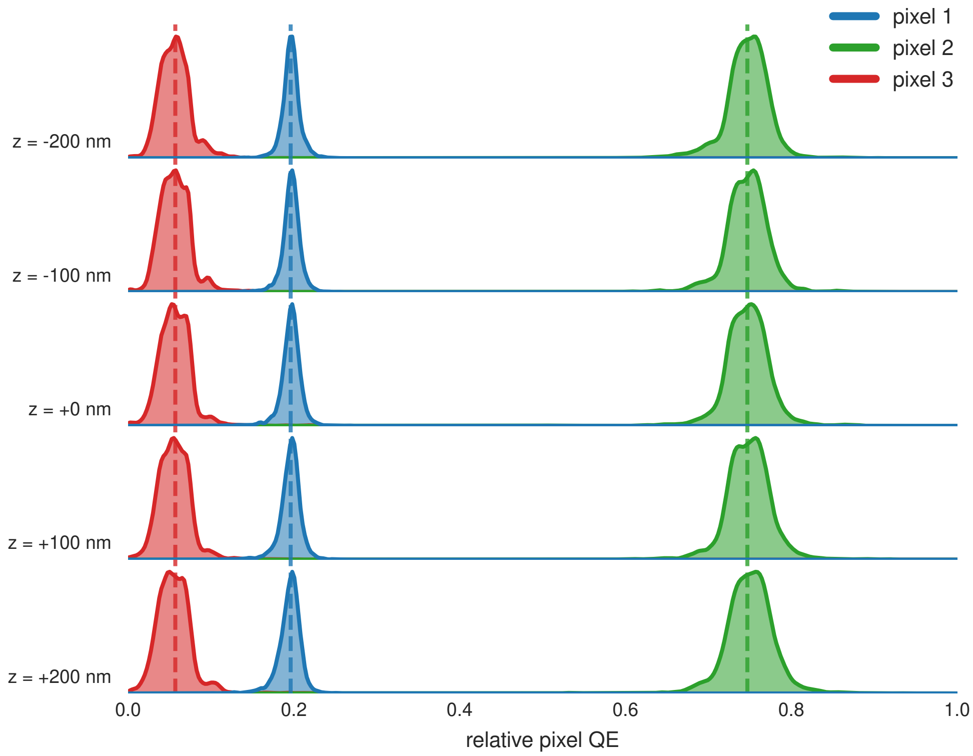


**Fig. S53. The effect of Z defocus on spectral signature bias.** Over a 400 nm defocus range, the spectral signature values remain stable.

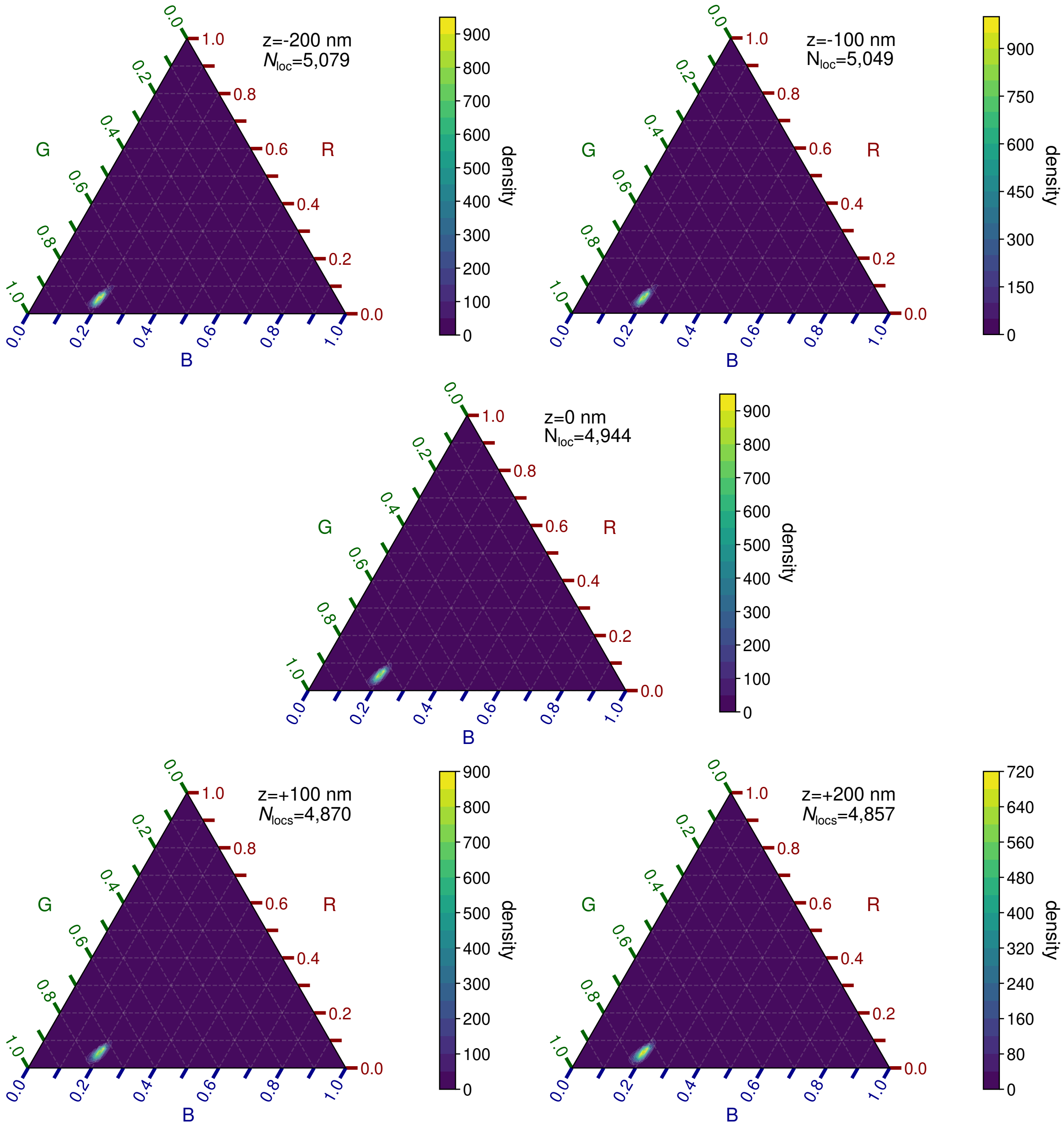


**Fig. S54. Ternary plots demonstrating effect of Z defocus on spectral signature bias.** Over a 400 nm defocus range, the spectral signature values remain stable.

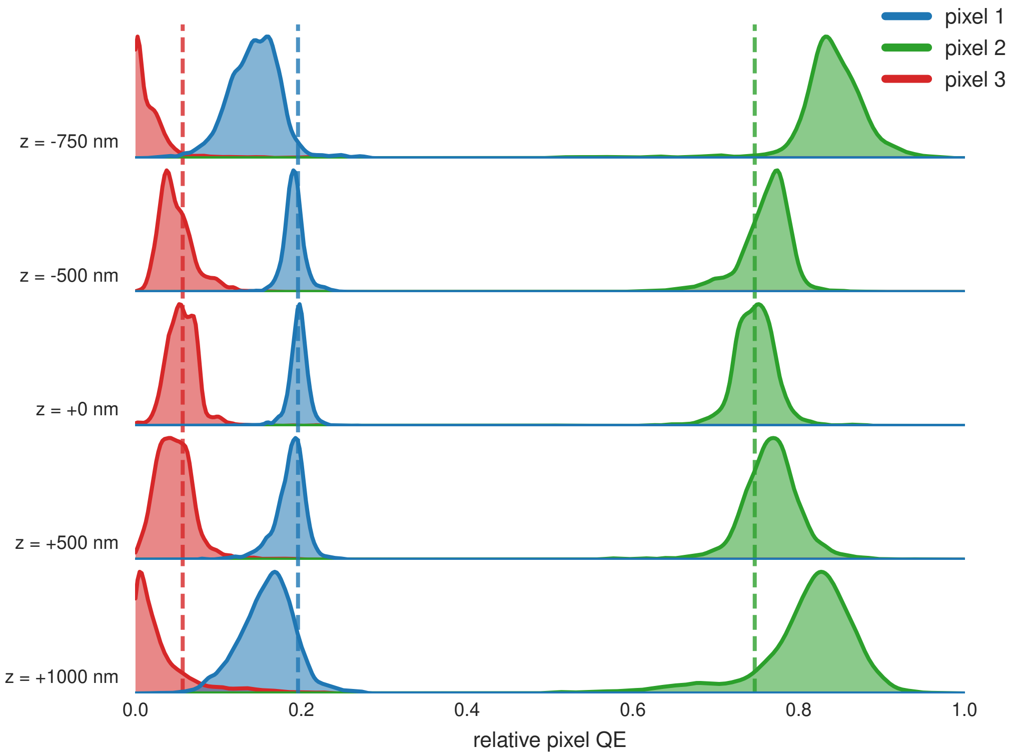


**Fig. S55. The effect of Z defocus on spectral signature bias.** As the z position of the emitter moves increasingly from the focal plane, the estimates of the pixel QEs move increasingly from their values at Z=0.

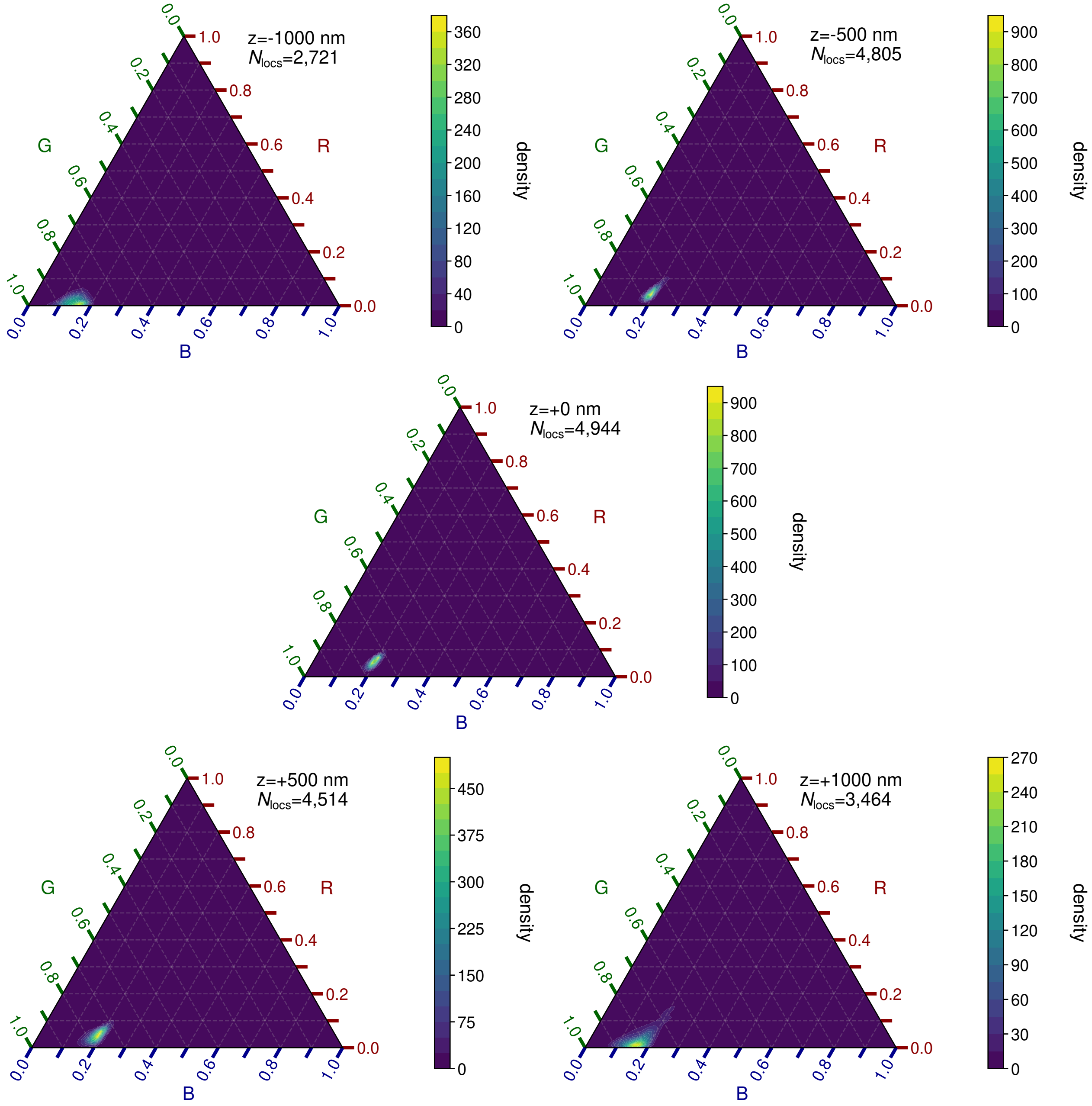


**Fig. S56. Ternary plots demonstrating effect of Z defocus on spectral signature bias.** As the z position of the emitter moves increasingly from the focal plane, the estimates of the pixel QEs move increasingly from their values at Z=0.

#### Supplementary Note S13: dSTORM on HeLa cells

**Methods.** HeLa cells were cultured at 37 °C and 5 % $CO_2$ in DMEM supplemented with 10 % FBS, 1 % penicillin/streptomycin, and 1 % glutamine. The day prior to fixation, cells were seeded onto argon plasma cleaned #1.5 glass µ-Slides (Ibidi, 80827). On the day of imaging, cells were washed 3x with PBS. Cells were fixed and permeabilised in Cytoskeleton Buffer Sucrose (10 mM MES, 138 mM KCl, 3 mM $MgCl_2$, 2 mM EGTA, 4.5 % sucrose) supplemented with 4 % formaldehyde and 0.2 % Triton X-100 at room temperature for 6 minutes before being moved back to 37 °C and 5 % $CO_2$. Cells were then washed three times with PBS + 0.1 % Tween 20. This follows the protocol as described in Daly *et al.*(11) Gold nanoparticles (60 nm Gold Colloid, BBI Solutions, EM.GC60/7) were diluted to a concentration of 2.5 % v/v in PBS before being deposited onto the cells for 20 minutes. Excess gold nanoparticle solution was removed and cells were washed three times with PBS + 0.1 % Tween 20, before incubating in a blocking solution of PBS + 5 % BSA for 60 minutes at room temperature. A solution of NanoTag-Antibody conjugate was prepared by incubating a solution of 1 µM Alexa Fluor 647 NanoTag (FluoTag®-X2 anti-Mouse Ig kappa light chain, NanoTag Biotechnologies, N1202-AF647-S) with 50 µg/mL Anti-alpha Tubulin antibody (DM1A, Abcam, ab7291) in PBS for 20 minutes at room temperature with constant shaking, before being diluted to a final concentration of 2.5 µg/mL NanoTag-Antibody conjugate in PBS + 0.1 % Triton X-100 + 1 % BSA. The blocking solution was removed, and cells were washed three times with PBS + 0.1 % Tween 20. Cells were then incubated in NanoTag-Antibody conjugate solution at room temperature for 60 minutes. Excess NanoTag-Antibody conjugate solution was removed, and cells were washed six times with PBS prior to imaging. Cells were imaged with HILO illumination in a dSTORM buffer consisting of 50 mM Tris-HCl, 10 mM NaCl, 10 % glucose, 10 mM MEA, 84 µg/mL catalase, and 0.2 mg/mL GLOX adjusted to pH 8.

**Results.** Fig. S57a & b shows $S^3$M's compatibility with dSTORM imaging. Here, we imaged the $\alpha$-tubulin network of a HeLa cell using the blinking of Alexa Fluor 647. As expected, we see the thin tubular network and highlight that cuts through these fibrillar structures (Fig. S57b) show the super-resolved nature of the data, Fig. S57c. This data shows widths of the microtubule network of 70–100 nm, well below the Abbe diffraction limit of ∼225 nm for Alexa Fluor 647. Equally, the distances between the two microtubules would have been difficult to see without super-resolved data. More quantitatively, the Fourier ring correlation (FRC)(21) shows a resolution of 86 nm±4 nm (see Fig. S58).

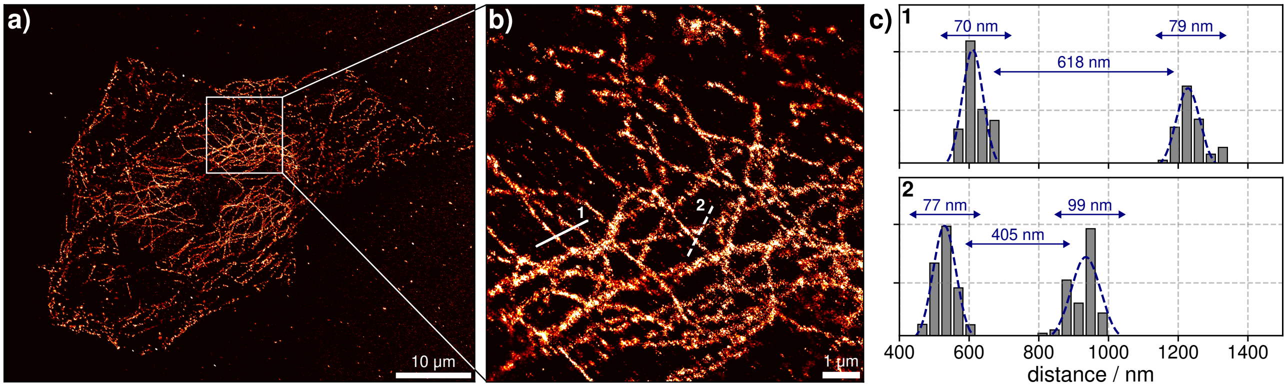


**Fig. S57. dSTORM imaging with $S^3$M. a)** dSTORM image of $\alpha$-tubulin network in a HeLa cell labelled with AlexaFluor 647. **b)** Zoom-in of the $\alpha$-tubulin network. **c)** Line plots demonstrating the resolution of individual tubulin below the diffraction limit highlighted in **b**

The FRC curve was calculated using a Python port of the MATLAB code written by Nieuwenhuizen *et al.*(21) Error bounds are ±1 standard deviation across 20 independent random half-splits.

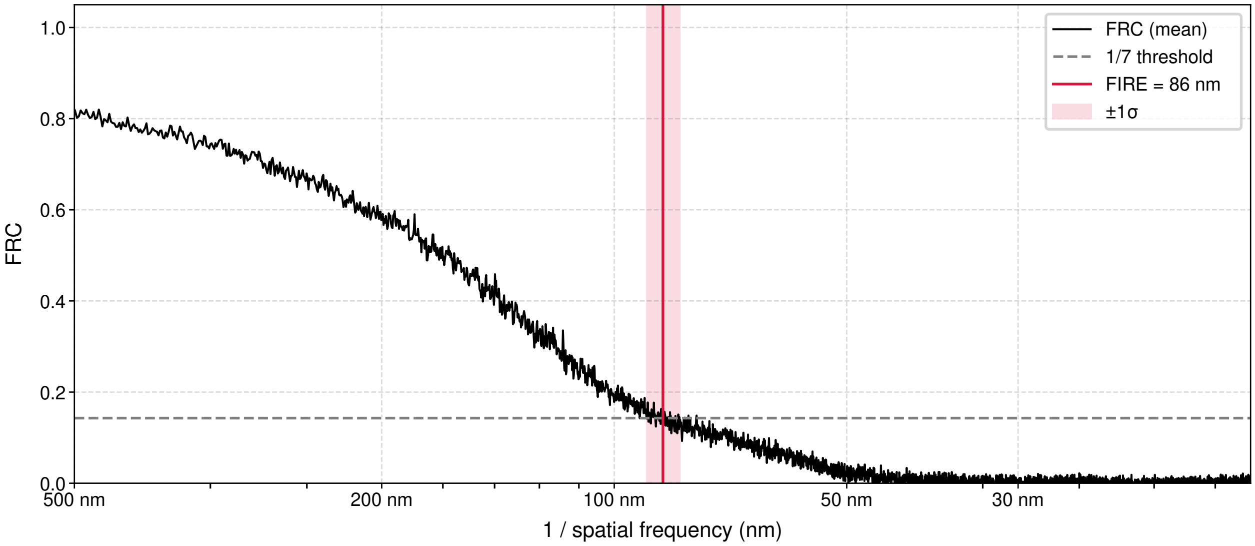


**Fig. S58.** FRC curve for the dSTORM experiment, showing the 86 nm resolution achieved.

#### Supplementary Note S14: Raw Data and Fits of ATTO 655 and Cy3B DNA-PAINT on BSC-1

Here we show some of the data processing of the ATTO 655 (Fig. S59) and Cy3B (Fig. S60) localisations from the DNA-PAINT on BSC-1. For each class of localisation we show 5 instances of the raw data, the weighting for the fits, the reconstructed PSF and localisation, the residual of the fit, and the extracted spectral fingerprint.

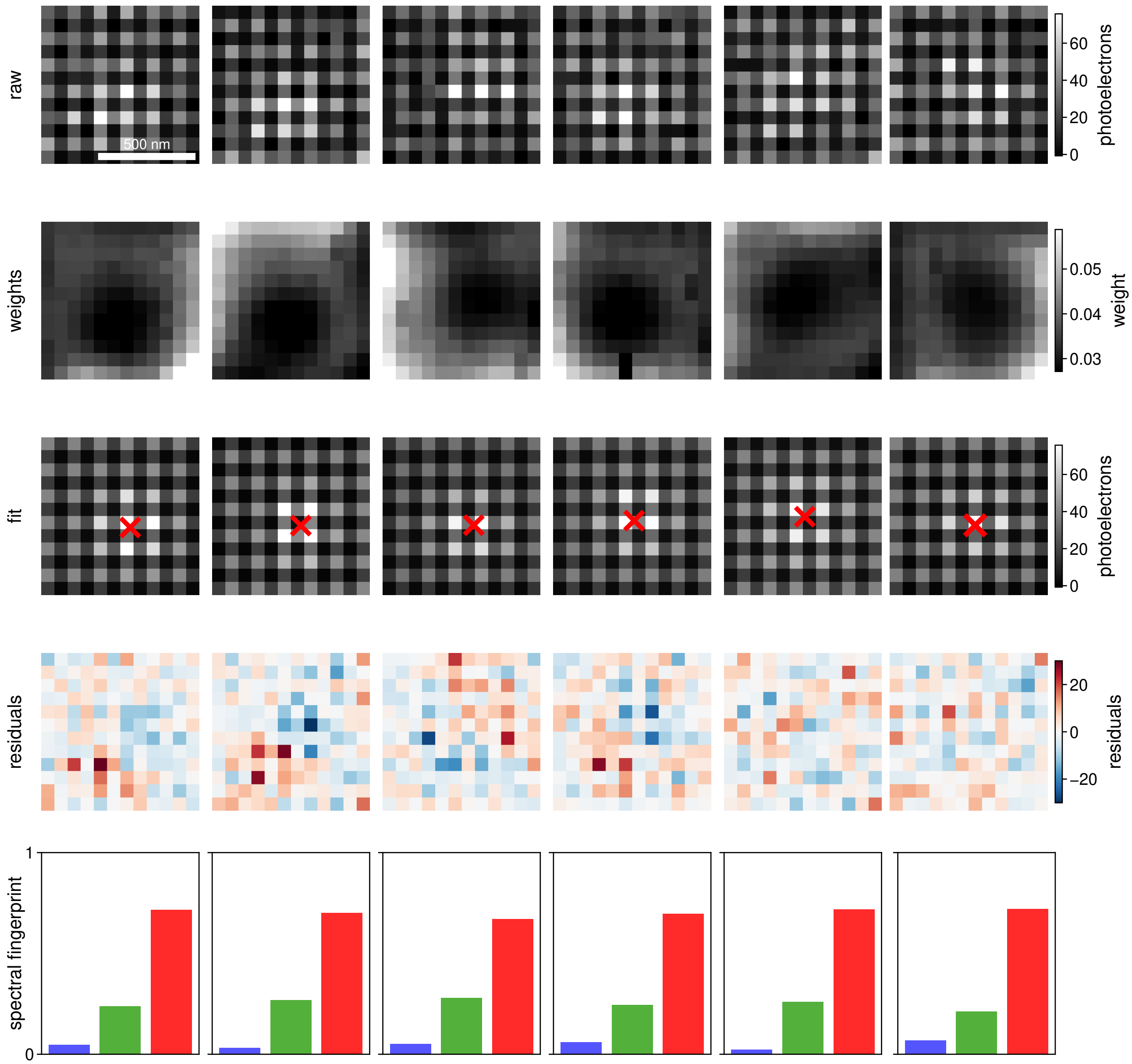


**Fig. S59.** Fit localisations of ATTO 655 from the DNA-PAINT on BSC-1 cells

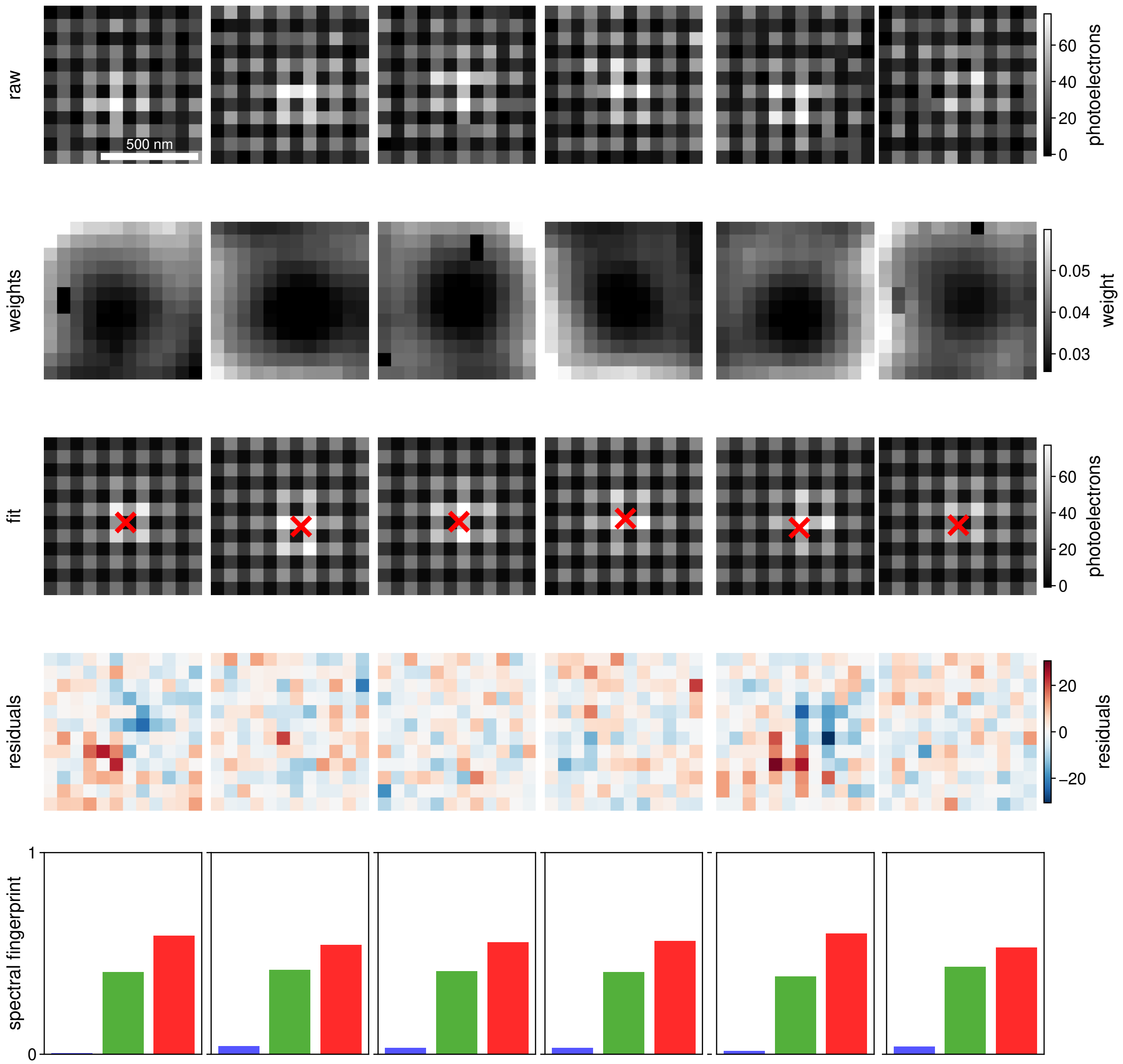


**Fig. S60.** Fit localisations of Cy3B from the DNA-PAINT on BSC-1 cells

#### Supplementary Note S15: ATTO 594 and Cy3B DNA-PAINT on BSC-1 Cells

Here we highlight that S$^3$M enables multiplexed super-resolution imaging using spectrally similar dyes, performing two-colour DNA-PAINT using Cy3B and ATTO 594 imager strands simultaneously. These dyes have a 50 nm peak-to-peak difference, and significant spectral overlap. Nevertheless, S$^3$M is able to spectrally separate localisations in one shot (Fig. S61).

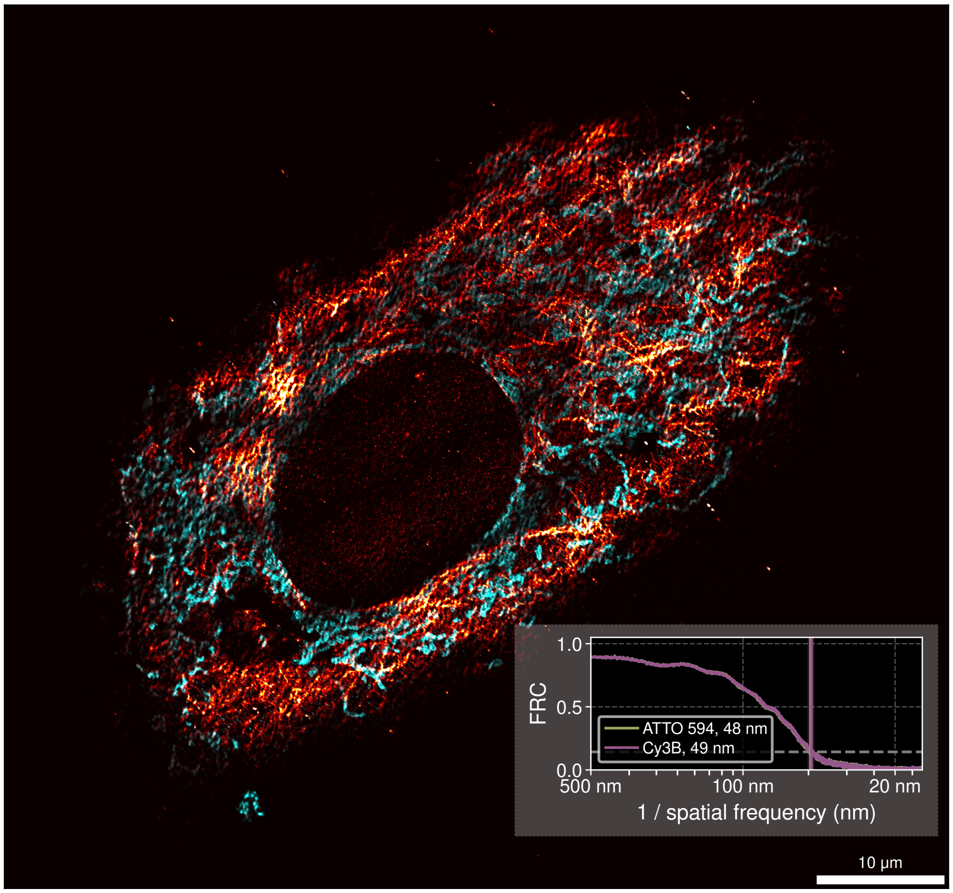


**Fig. S61. DNA-PAINT imaging with spectrally similar dyes using S$^3$M.** Two-colour DNA-PAINT image of ATTO 594 labelled vimentin (orange) and Cy3B labelled mitochondria (cyan) in a BSC-1 cell. Inset shows FRC curve, highlighting the 50 nm resolution achieved in both channels. The methods are the same as for Fig. 5, but with an ATTO 594 labelled strand instead of ATTO 655.

#### Supplementary Note S16: Alternative Pattern Simulations

One of the key principles of S$^3$M is that the underlying spatial pattern of pixels can be arbitrary, allowing it to be used beyond the standard Bayer mosaic (RGGB 2×2 repeating unit). To demonstrate this, we simulated three different dyes across the visible spectrum (ATTO 488, ATTO 565, and ATTO 647N) on six different spatial pixel arrangements: the Bayer-pattern, a GGBR pattern, an RGBR pattern, an RRGB pattern, the Fujifilm X-Trans sensor, and an entirely random arrangement of RGB pixels (Fig. S62). Each simulation uses an NA of 1.49, a pixel size of 69 nm, a uniform background of 5 photons per pixel, and has the equivalent pixel QE, gain, read noise and variance of the Bayer-patterned Ximea MC050CG-SY sensor (Table S2). For each condition, a two-dimensional PSF was placed at a position drawn uniformly within the central pixel unit cell (*i.e.* the RGGB repeating unit, in the case of a Bayer pattern) of the sensor and fit. This process is repeated 100,000 times to obtain the spatial and colour precision across 200 photon levels, logarithmically spaced between 500 and 50,000 photons. In each spatial arrangement, molecules can be localised and their spectral fingerprint fit, and show photon dependent precisions (Fig. S63). Across all tested arrangements, we show that S$^3$M still functions, and does not require a specific pixel arrangement—*i.e.* does not require the Bayer-pattern. The pixels can retain the same RGB ratios and be rearranged (GGBR), the colour balance can be altered (RGBR and RRGB), the repeating unit can be large (6×6 for the X-Trans), or there can be no underlying pattern at all (Random). As long as the pixel arrangement is known *a priori*, S$^3$M can function with that detector.

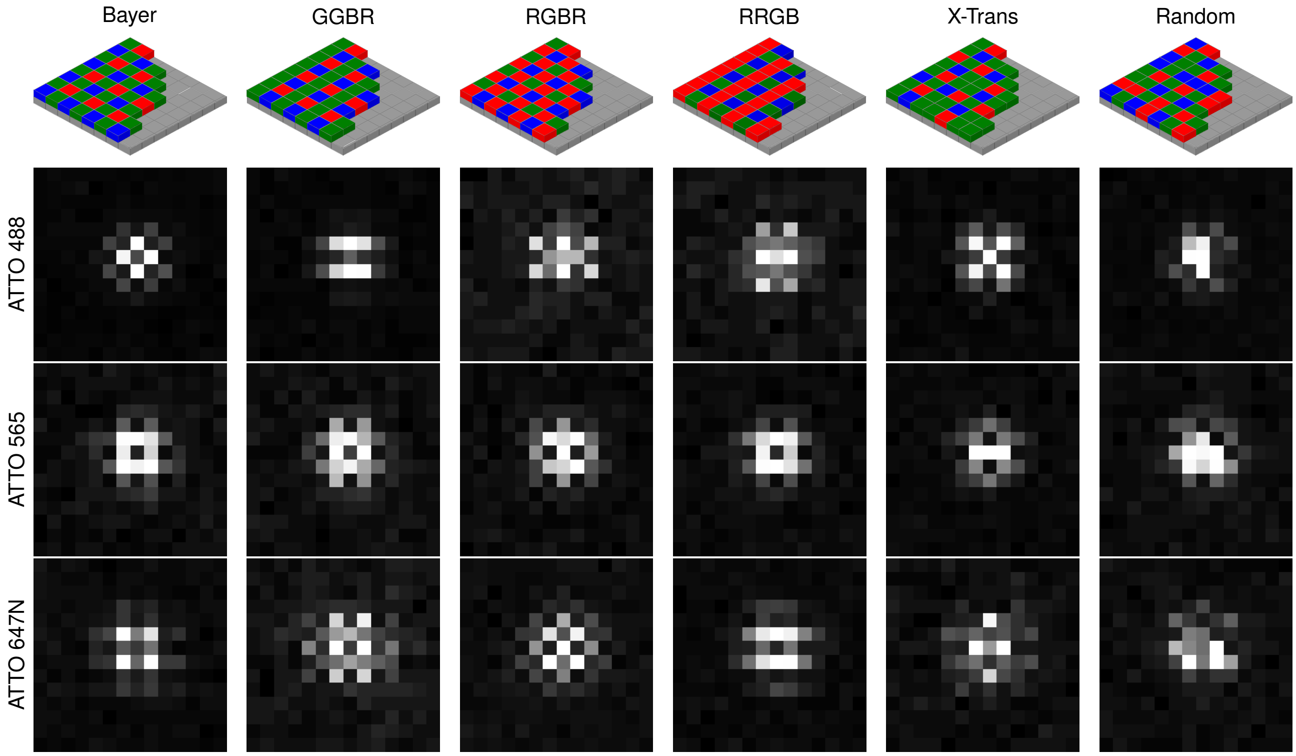


**Fig. S62. Simulated Spatial Patterns.** Schematic diagram of each spatial arrangement of pixels. Below are the resulting PSFs of ATTO 488, ATTO 565, and ATTO 647N on each respective detector.

Another key principle of S$^3$M is that the underlying spectral response of pixels can be arbitrary, allowing it to be used beyond the standard RGB Bayer mosaic. To demonstrate this, we simulated three different dyes across the visible spectrum (ATTO 488, ATTO 565, and ATTO 647N) on six different spectral pixel patterns: the Bayer-pattern, a Cyan-Yellow-Magenta CYYM pattern, an RGB-Emerald pattern, an RGB-White pattern, a Phasor of the visual wavelength range, and an RGB-near IR of RGB pixels (Fig. S64). Each simulation uses an NA of 1.49, a pixel size of 69 nm, and a uniform background of 5 photons per pixel, and has pixel QEs shown in Fig. S64b. The remaining camera parameters (gain, read noise and variance) are the same as the Bayer-patterned Ximea MC050CG-SY sensor (Table S2). Simulations were performed and PSFs were fit as described before, this time for the new spectral patterns (Fig. S66). Similar to changing the pixel spatial arrangements (Fig. S63), S$^3$M is able to localise molecules and determine their spectral fingerprint. Together, these simulations show the generality of S$^3$M; so long as the pattern and QEs are known, the technique can localise and quantify a spectral fingerprint of individual molecules. Generally, we find that among all the pixel arrangements and QE possibilities, that the Bayer pattern performs well for localisation and spectral fingerprinting. That, combined with the wide availability of Bayer-patterned detectors, makes it an ideal starting point for most applications.

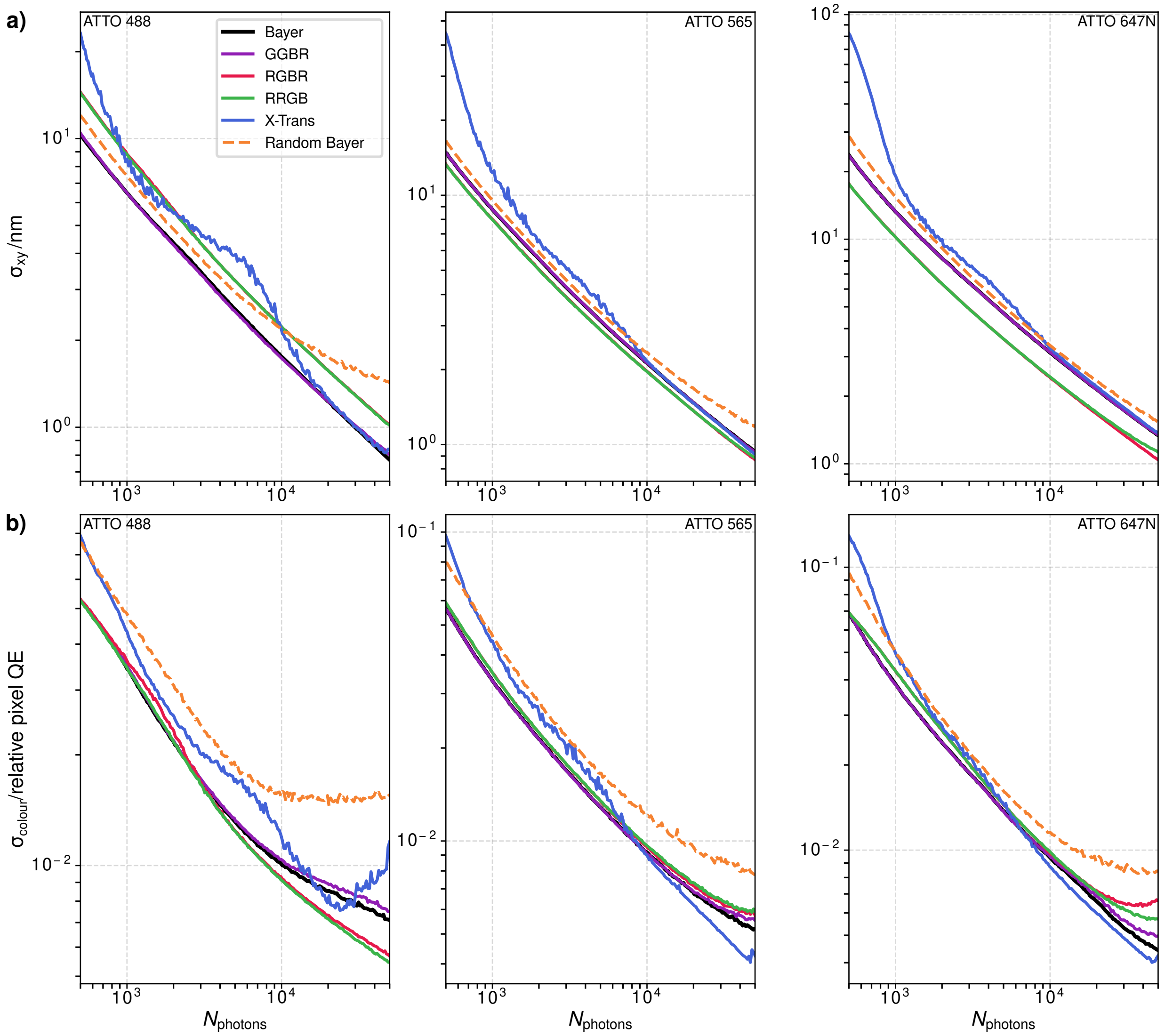


**Fig. S63. Precision of Spatial Patterns. a)** The localisation precision at a range of photon values for each pixel arrangement. **b)** The colour precision at a range of photon values for each pixel arrangement. From left to right, the dyes are ATTO 488, ATTO 565, and ATTO 647N.

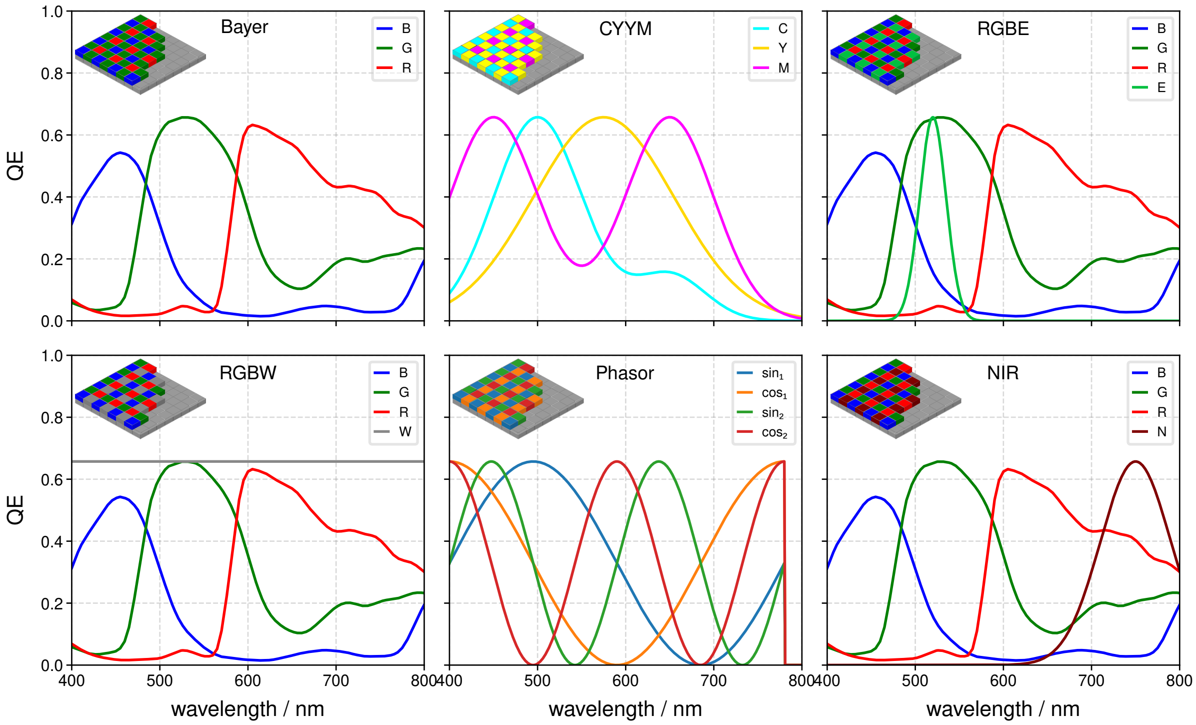


**Fig. S64. Arbitrary spectral filters.** The simulated spectra used in simulations, with inset diagrams showing schematics of the pixel arrangements.

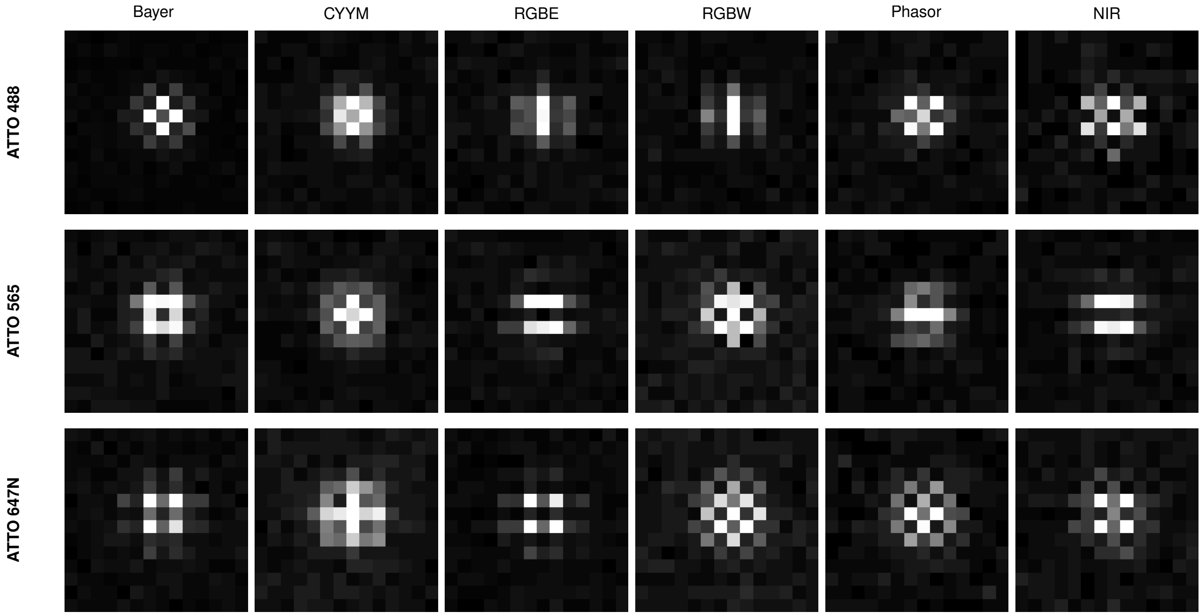


**Fig. S65. Exemplar PSFs with different spectral QEs.** Exemplar PSFs of simulated ATTO 488, ATTO 565, and ATTO 647N dyes with different spectral QE patterns.

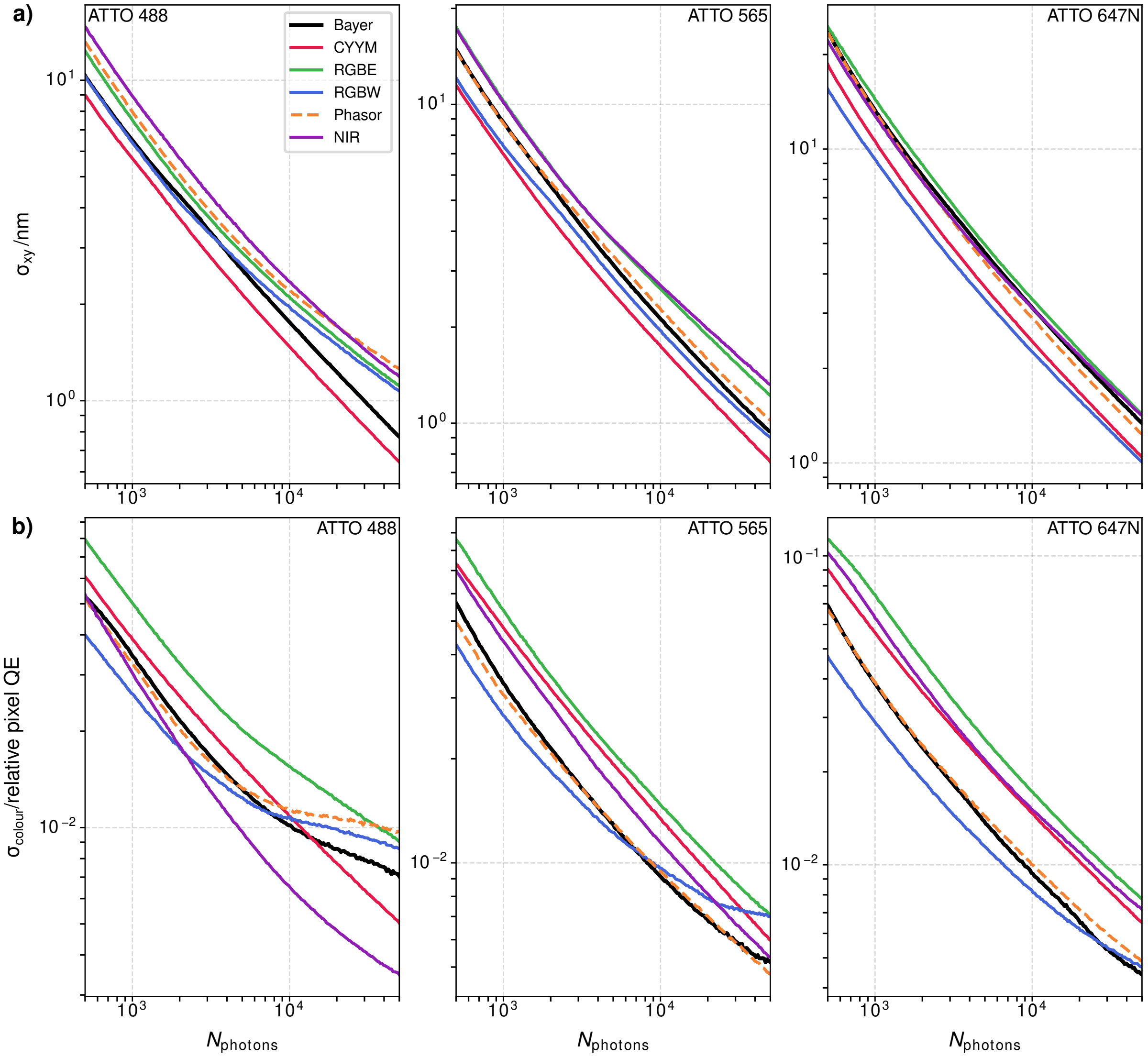


**Fig. S66. Precision of different spectral QEs. a)** The localisation precision at a range of photon values for each spectral pixel pattern. **b)** The colour precision at a range of photon values for each pixel arrangement. From left to right, the dyes are ATTO 488, ATTO 565, and ATTO 647N.

#### Supplementary Note S17: Nile Red/NR4A Forward Model

The Nile Red/NR4A forward model consists of three stages. First, the spectrum of Nile Red, converted to the transition dipole moment representation,(22) was fit with a skew Gaussian model of

$$\tilde{I}(E; A, \mu, \sigma, \alpha) \;=\; \frac{A}{\sigma\sqrt{2\pi}} \exp\left(-\frac{(E-\mu)^2}{2\sigma^2}\right) \cdot \left[1+\mathrm{erf}\left(\alpha\,\frac{E-\mu}{\sigma}\right)\right], \tag{S22}$$

where $E$ is the photon energy, $A$ is the amplitude, $\mu$ the peak energy, $\sigma$ the Gaussian width, and $\alpha$ a skewness parameter. This fit is shown in Fig. S67. For the forward model, we assume that the skewness and width of the spectra do not change and fix these at $\sigma = 0.1630$ eV and $\alpha = -1.565$ respectively. Thus the sole parameter of the forward model is to predict what $\mu$ would lead to the observed pixel QE fraction values from a PSF fit. An example of how these values change with changing mean wavelength is shown in Fig. S68.

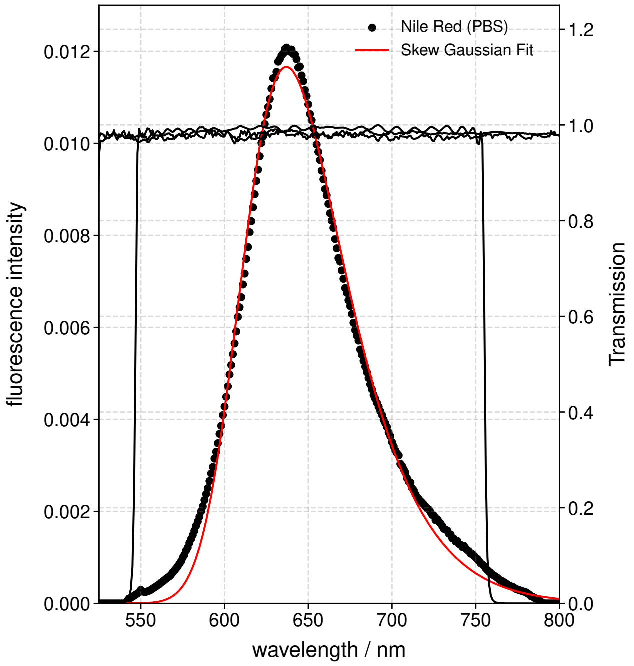


**Fig. S67.** An example of the fit to the fluorescence spectrum of Nile Red, showing also the typical bandpass filter and longpass filter used in the experiment.

Typically, the raw localisations were not fit directly to this model. Instead, to improve the stability of the fit, the data was pixellated on a grid (here 50 nm×50 nm pixels) and the localisations grouped based on their location on the grid. If more than 5 localisations were present in a pixel, then these localisations were averaged *via* an inverse-parameter error weighted mean, then these pixel-level results were fit to produce the results shown in Fig. 5. This analysis pipeline is shown schematically in Fig. S69.

We tested the two stages of the forward model by simulating 10,000 different puncta for a range of Nile Red peak wavelengths, from low (200) to high (10,000) photons per punctum—taking these ranges from our experimental dataset. We tested three background levels; 20, 45 and 70, again based on experimental data. As can be seen in Fig. S70, at the level of individual localisations the spectral estimation precision is $\sim$10 nm at above 1,000 photons for Nile Red wavelengths <600 nm, and increases as the Nile Red shifts more to the red. In Fig. S71, we tested the bias on this estimation—at low photon values the bias at mean emission wavelengths >580 nm is relatively large, only decreasing to reasonable (<20 nm) levels above 10,000 photons. However, we do not typically fit the raw localisations, instead grouping them into pixels to reduce the effect of bias, increase the effective photon numbers for each estimation, and to reduce fit time.

We thus tested the pixellation and localisation grouping element of the forward model workflow. Here, we took simulated images at a range of dye photon and background values, and fit them. These fits were then used to generate “pixels” by randomly sampling $N$ (where $N$ ranged from 1 to 50) localisations from the same underlying wavelength distribution. We then fit these inverse-error weighted “pixels” and examined if this grouping process improved the underlying fit. The precision and bias are plotted *versus* $N_{\mathrm{photons}}$ per pixel in Fig. S72. As can be seen, this significantly improves the spectral precision and

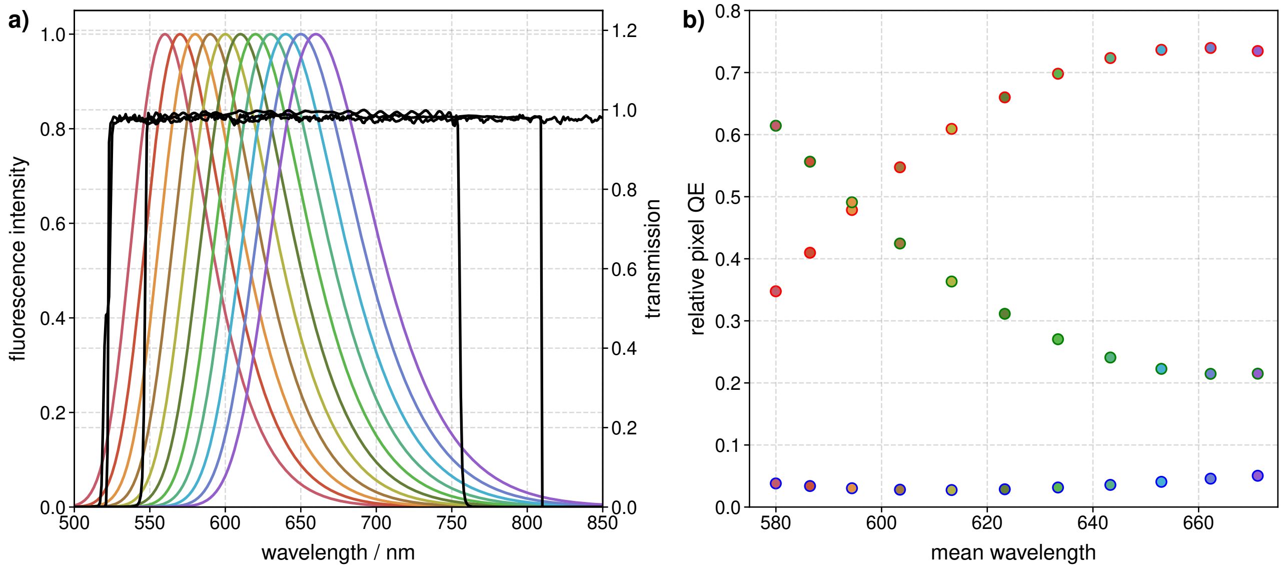


**Fig. S68.** An example of a shifting Nile Red/NR4A emission spectrum, and the corresponding effect that would have on the relative pixel QEs in the $S^3M$ analysis pipeline.

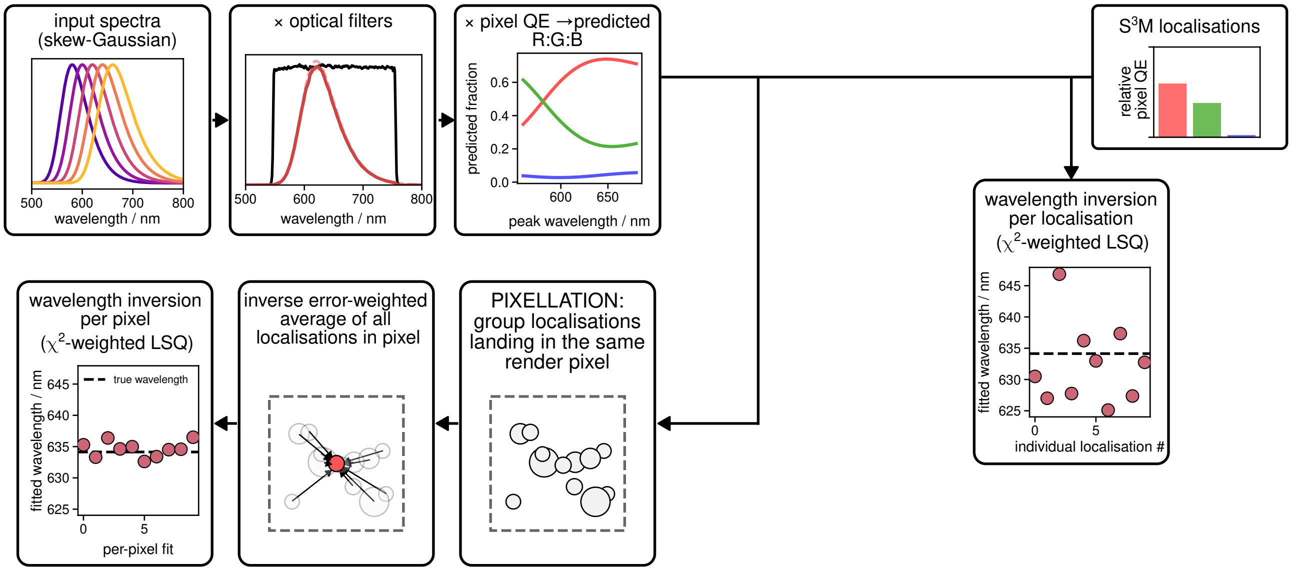


**Fig. S69.** A schematic of the analysis pipeline of Nile Red and NR4A data.

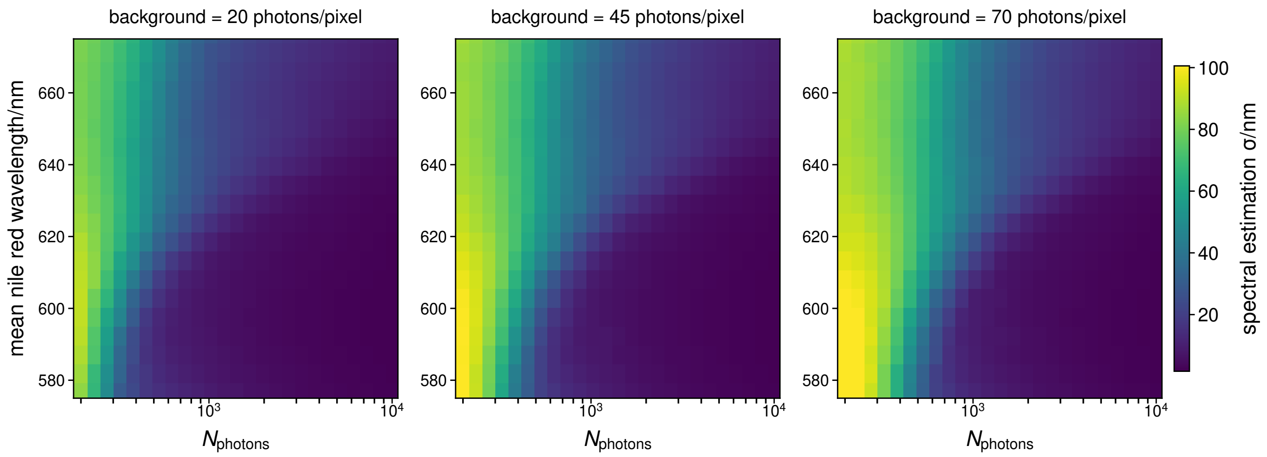


**Fig. S70. Validation of the Nile Red Forward Model.** Precision of the spectral estimate at a range of different background values.

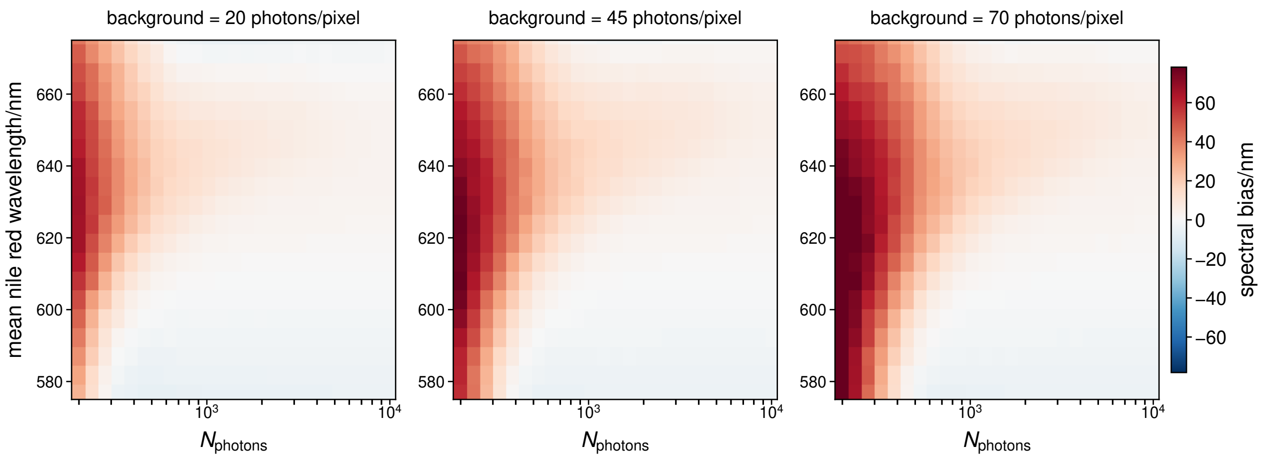


**Fig. S71. Validation of the Nile Red Forward Model.** Bias of the spectral estimate at a range of different background values.

bias, with the practical experimental photon ranges ($10^4$–$10^7$) and spectral ranges (600–640 nm) showing precisions of <2 nm and biases <|1| nm.

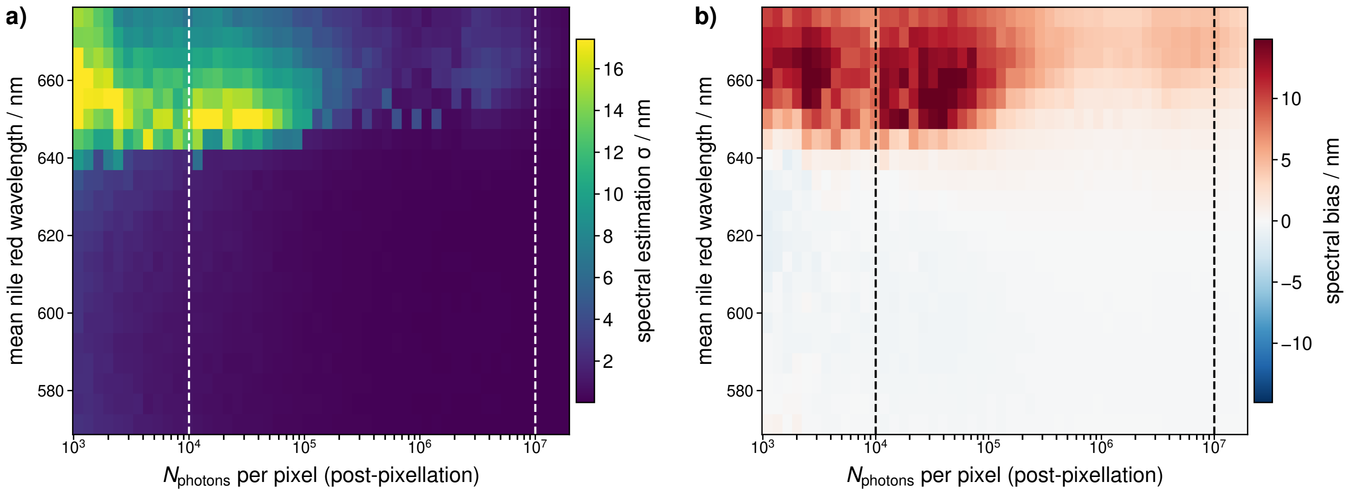


**Fig. S72. Validation of the Nile Red Forward Model including pixellation. a)** Precision of the spectral estimate as a function of photons per pixel, with the practical experimental ranges marked as dashed lines. **b)** Bias of the spectral estimate as a function of photons per pixel, with the practical experimental ranges marked as dashed lines.

#### Supplementary Note S18: Camera Calibration

The cameras used in this work were calibrated using the method described in Huang *et al.*,(17) but with a slight change to the experimental procedure. The code used to accomplish this was based on the code developed by Bruggeman.(23) In brief, the experimental procedure for the calibration of a "standard" (s)CMOS camera is to record full sensor images at 6+ different intensities (including dark frames) with, in our experience, at least 20,000 frames per intensity level. For the CMOS cameras used in this work (Thorlabs CS505CU, Ximea MC050CG-SY, ZWO ASI 585MC), with their additional spatially-patterned detector arrays, in order to determine the gain and read-noise per pixel, filters were placed in front of the white light source used to ensure 7 different intensities across the camera's sensitivity for each different pixel channel. Specifically, for the "blue" pixels a 10 nm bandpass filter centred at 473 nm (Semrock FF01-473/10-25) was used, for the "green" pixels a 44 nm bandpass filter centred at 520 nm (Semrock FF01-520/44-25) was used, and for the "red" pixels a 40 nm bandpass filter centred at 692 nm (Semrock FF01-692/40-25) was used. This set of images was then used to determine the pixel-dependent gain and read noise. These are summarised for the cameras used below in Table S2. The results of the full-sensor calibration were uploaded to Zenodo.(24) We note that, owing to the pixels' equal sensitivity in the NIR, in principle all pixels could be calibrated at once with an NIR bandpass filter or NIR light source.

| | Thorlabs CS505CU | Ximea MC050CG-SY | ZWO ASI 585MC |
|---|---|---|---|
| Median Offset ± MAD (ADU) | 100.4 ± 0.1 | 32.4 ± 0.2 | 242.5 ± 0.2 |
| Median Gain ± MAD (ADU/$e^-$) | 0.47 ± 0.01 | 0.414 ± 0.004 | 3.59 ± 0.02 |
| Median Read noise ± MAD ($e^-$) | 1.91 ± 0.06 | 2.34 ± 0.06 | 0.73 ± 0.05 |
| Read noise RMS ($e^-$) | 1.97 | 2.49 | 0.85 |

**Table S2.** Camera calibration results for the multiple CMOS cameras used in this work. MAD stands for median absolute deviation.

#### Supplementary Note S19: Timeline of commercially available Bayer detectors

A number of commercially available Bayer-patterned detectors released since 2018 have reached sufficient quality as to enable single-molecule detection, which we highlight in Fig. 1c. The specific technical data relating to the detectors plotted in Fig. 1c is shown in Table S3. The groups of Ozcan, Tinnefeld and Acuna highlighted that Bayer detectors presented a challenge for single-molecule detection in low-NA point-of-care setups,(25) with single-molecule sensitivity being achieved recently by Loretan *et al.* (26) The low NA objectives and specific detectors used precluded sensitivity to dyes across the 500–750 nm spectral regime, with only dyes close to the peak QE of ~550 nm (ATTO 542, Cy3B) demonstrated as detectable. We have compiled, from the manufacturers, data on peak QE and read noise of a variety of commercially available Bayer-patterned sensors (Tables S3 and S4). These data, plotted in Figure S73, highlight that these detectors have only recently (post-2018) reached the quality level to enable simultaneous single-molecule detection and the extraction of spectral signatures.

| Detector | Read noise RMS ($e^-$) | Peak QE | Filter Pattern | Pixel Size/µm | Release Year | Specification Source |
|---|---|---|---|---|---|---|
| Thorlabs CS505CU | 1.97 | 0.65 | Bayer | 3.45 × 3.45 | 2018 | Table S2 & Manufacturer |
| Thorlabs CS165CU | 4.00 | 0.65 | Bayer | 3.45 × 3.45 | 2020 | Manufacturer |
| Thorlabs CS126CU | 2.5 | 0.65 | Bayer | 3.45 × 3.45 | 2020 | Manufacturer |
| Ximea MC050CG-SY | 2.49 | 0.67 | Bayer | 3.45 × 3.45 | 2019 | Table S2 & Manufacturer |
| ZWO ASI 183MC | 1.60 | 0.84 | Bayer | 2.4 × 2.4 | 2021 | Manufacturer |
| ZWO ASI 585MC | 0.70 | 0.91 | Bayer | 2.9 × 2.9 | 2022 | Table S2 & Manufacturer |
| ZWO ASI 676MC | 0.56 | 0.83 | Bayer | 2.0 × 2.0 | 2024 | Manufacturer |

**Table S3.** Technical data of commercially available Bayer detectors plotted in Fig. 1c.

| Detector | Read noise RMS ($e^-$) | Peak QE | Filter Pattern | Pixel Size/μm | Release Year | Specification Source |
|---|---|---|---|---|---|---|
| Ximea MQ013CG-E2-S7 | 27.70 | 0.47 | Bayer | 5.30 × 5.30 | 2010 | Manufacturer |
| Thorlabs DCC1645C | 25.00 | 0.45 | Bayer | 3.60 × 3.60 | 2012 | Manufacturer |
| Thorlabs DCC1240C | 30.00 | 0.45 | Bayer | 5.30 × 5.30 | 2012 | Manufacturer |
| Ximea MX022MG-CM-X2G2 | 13.34 | 0.48 | Bayer | 5.50 × 5.50 | 2012 | Manufacturer |
| Ximea MQ042CG-CM-S7 | 16.14 | 0.48 | Bayer | 5.50 × 5.50 | 2012 | Manufacturer |
| Ximea MQ003CG-CM | 20.00 | 0.42 | Bayer | 7.40 × 7.40 | 2013 | Manufacturer |
| Ximea MQ020CG-E2 | 7.00 | 0.47 | Bayer | 4.50 × 4.50 | 2013 | Manufacturer |
| Ximea MQ013CG-E2 | 25.00 | 0.47 | Bayer | 5.30 × 5.30 | 2013 | Manufacturer |
| Ximea MQ013CG-ON | 10.00 | 0.48 | Bayer | 4.80 × 4.80 | 2013 | Manufacturer |
| Ximea MT200CG-CM | 8.00 | 0.55 | Bayer | 6.40 × 6.40 | 2013 | Manufacturer |
| Ximea MT023CG-SY | 7.36 | 0.70 | Bayer | 5.86 × 5.86 | 2014 | Manufacturer |
| Ximea MQ013CG-ON-S7 | 8.99 | 0.48 | Bayer | 4.80 × 4.80 | 2014 | Manufacturer |
| Ximea MQ022CG-CM-S7 | 15.95 | 0.48 | Bayer | 5.50 × 5.50 | 2016 | Manufacturer |

**Table S4.** Technical data of a selection of commercially available Bayer detectors available pre-2018.

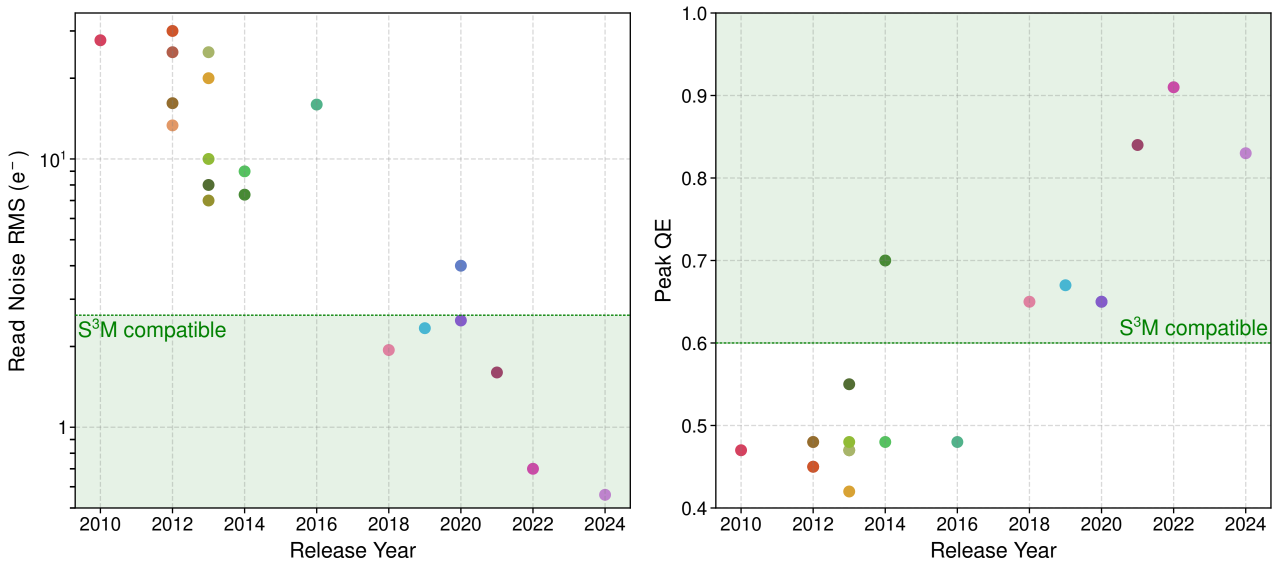


**Fig. S73. a)** Read noise RMS and **b)** Peak QE *vs.* detector release year, highlighting that it is post-2018 when a range of cameras began to be made available that could achieve single-molecule detection and the simultaneous extraction of spectral information. Points are colour-coded to highlight the same camera's read noise and peak QE. Shaded green areas highlight QE and read noise ranges compatible with S$^3$M.

#### Supplementary Note S20: Additional *S. aureus* imaging

Below we show additional information on the *S. aureus* imaging from Fig. 5f–h. This includes demonstrations that extracted wavelength is independent of detected photons (Fig. S74), examples of 5 processed localisations from raw data (Fig. S75), and a technical replicate taken on the ZWO ASI 585MC (Fig. S76) demonstrating local areas of high and low mean wavelengths.

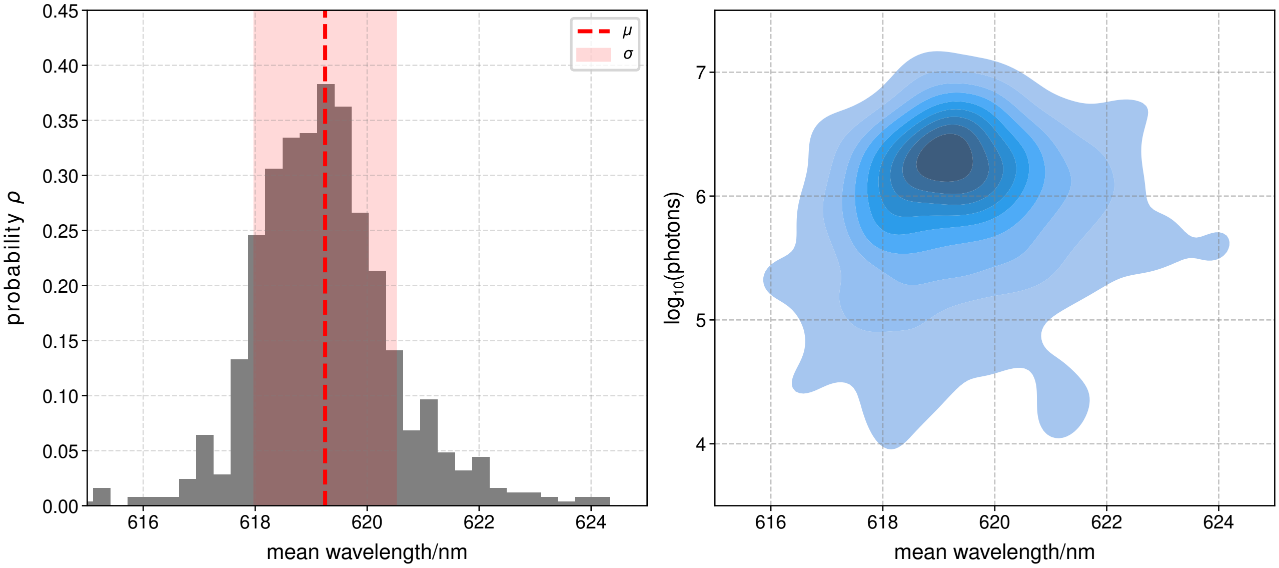


**Fig. S74. *S. aureus* imaging shows no correlation between number of photons per analysed pixel and mean wavelength. a)** The mean wavelength histogram is highly clustered at 620 nm with a $\sigma = \pm 1$ nm. **b)** KDE plot highlights no correlation between the number of photons per analysis.

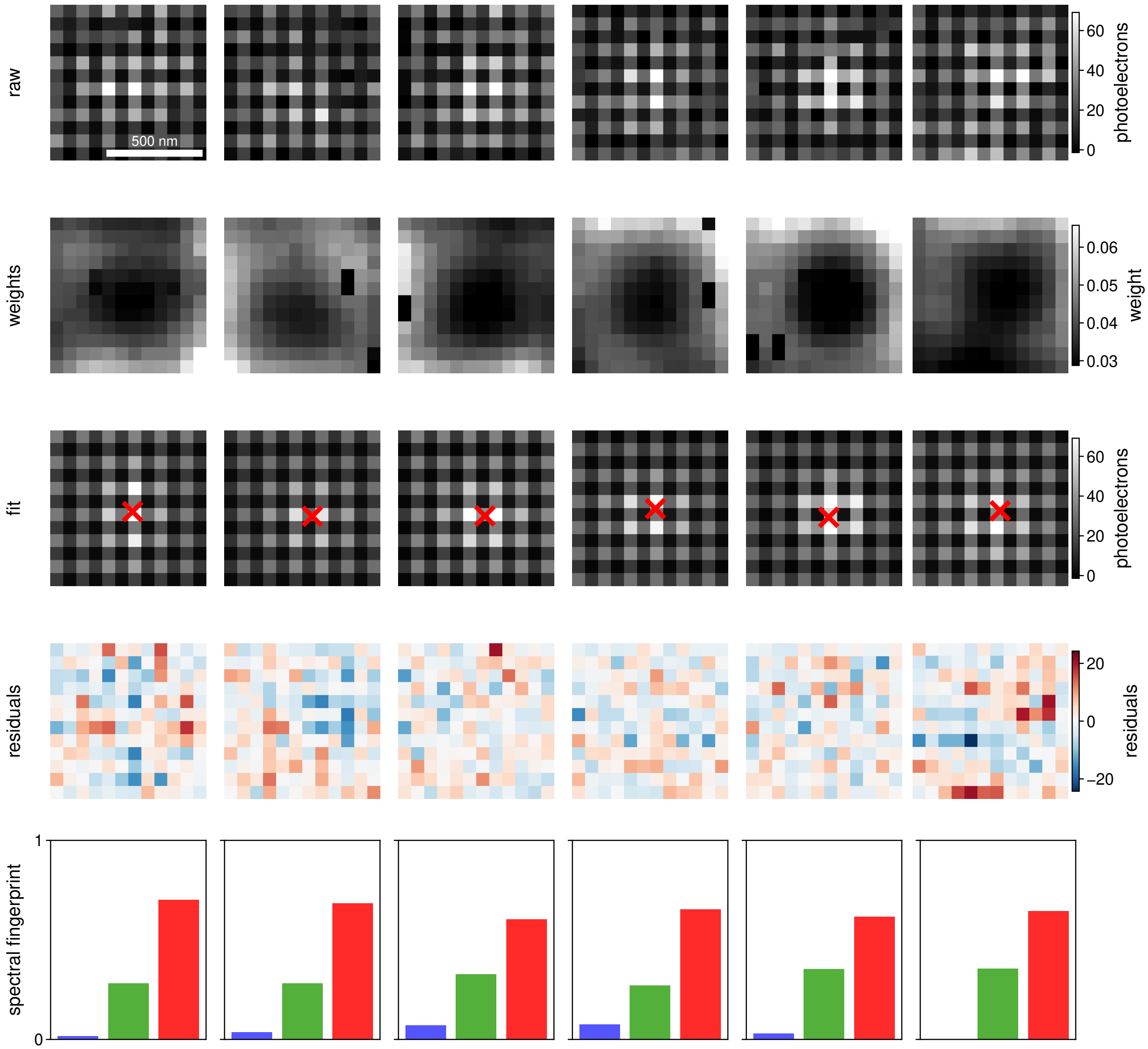


**Fig. S75. Fitting NR4A localisations from *S. aureus*.** Raw localisations, weights, reconstructed PSFs with localisations, residuals, and extracted spectral fingerprints of 5 different molecules.

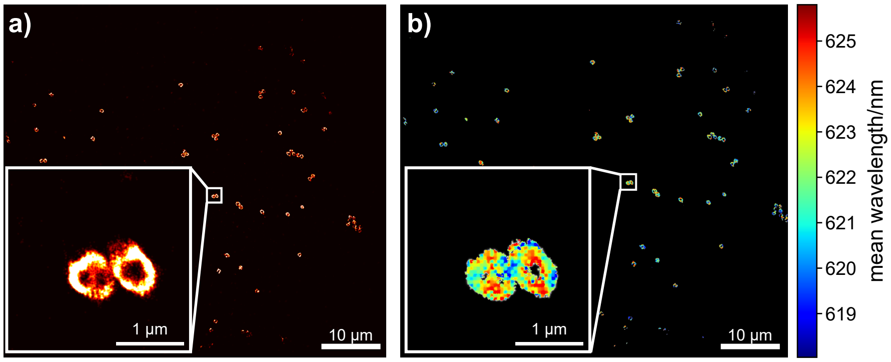


**Fig. S76. *S. aureus* imaging done on the ZWO ASI 585MC. a)** Super-resolution image, inset shows two select cells. **b)** Spectrally-resolved image, inset shows the same select cells from **a)**.

#### Supplementary Note S21: DNA Sequences

| Strand | Sequence |
|---|---|
| J7b | Cy5-CCCTAGCAAGCCGCTGCTACGG |
| J7h | Cy3-CCGTAGCAGCGCGAGCGGTGGG |
| J7r | biotin-HEG-HEG-CCCACCGCTCGGCTCAACTGGG |
| J7x | CCCAGTTGAGCGCTTGCTAGGG |

**Table S5.** DNA sequences used in this work. HEG = Hexaethylene glycol.

#### Supplementary Note S22: Calibrating Precision

Localisation precision and colour precision were measured experimentally with a sample of TetraSpeck™ beads immobilised in PVA. Images were taken at a variety of excitation wavelengths and powers to sample a range of photon count rates and colour ratios per frame (See Imaging Conditions in Table S3). After performing puncta fitting and drift correction,(27) the trajectory of individual beads was linked together by their positional coordinates using an HDBSCAN algorithm(28) so their behaviour can be tracked over time. Here, the definition of the precision is the standard deviation of the fitted positions of a bead from the mean position of the bead.(12) For each individual bead, the precision is calculated as

$$\sigma_{\text{xy}} = \sqrt{\frac{\sum(X_N - \bar{X})^2}{N}} \tag{S23}$$

where $X_N$ is the measured position at some observation $N$ and $\bar{X}$ is the average position. For plotting the precision curve in Figure 2d, each point in black consists of the precision in localisation of a single bead, plotted against the average number of photons detected per frame for that bead. For colour information, each individual bead generates three colour values $A$ and therefore three measured precisions—one for each pixel type $p$.

$$\sigma_{\text{colour}} = \sqrt{\frac{\sum(A_{N,p} - \bar{A}_p)^2}{N}} \tag{S24}$$

For plotting the precision curve in Figure 2d, each point consists of the precision of a single colour for an individual bead, plotted against the average number of detected photons of that colour. Due to the layout of the Bayer mask, green pixels are twice as common as red and blue, leading to a $\sqrt{2}$ reduction in variation compared to the other colour types. For this reason, each precision point for the green pixel is multiplied by a factor of $\sqrt{2}$.

#### Supplementary Note S23: Quantifying Dye Pair Accuracy

Here we show graphically the processing of quantifying dye pair misidentification earlier described in the Methods section, using the ATTO 565 and ATTO 633 data as an example. First, the cumulative data of all emitted photons from each molecule is recorded. To establish the mean pixel QE values for each of the underlying components, we fit data sets with high cumulative photons that show a clear bimodal distribution to a two-component GMM (S77a).

Because the mean pixel QE of each component should not vary as a function of photons, only the weighting of each population and standard deviation of pixel QE, we use the mean values of the high photo data set to fit the lower photon data sets. With the means fixed to a set value, only the new weights and standard deviations of each population are fit with a GMM (S77b). Finally, using Bayes' selection rule, we sort the data set with the GMM to predict which molecules belong to which component. Once these predictions are made, they are used as the ground truth for a bootstrapping process to identify how many times they can be correctly assigned (S77c). The data set is bootstrapped 10,000 times, and the misidentification rate is recorded. This process is repeated for a range of cumulative photon values from 300 photons to 250,000 photons.

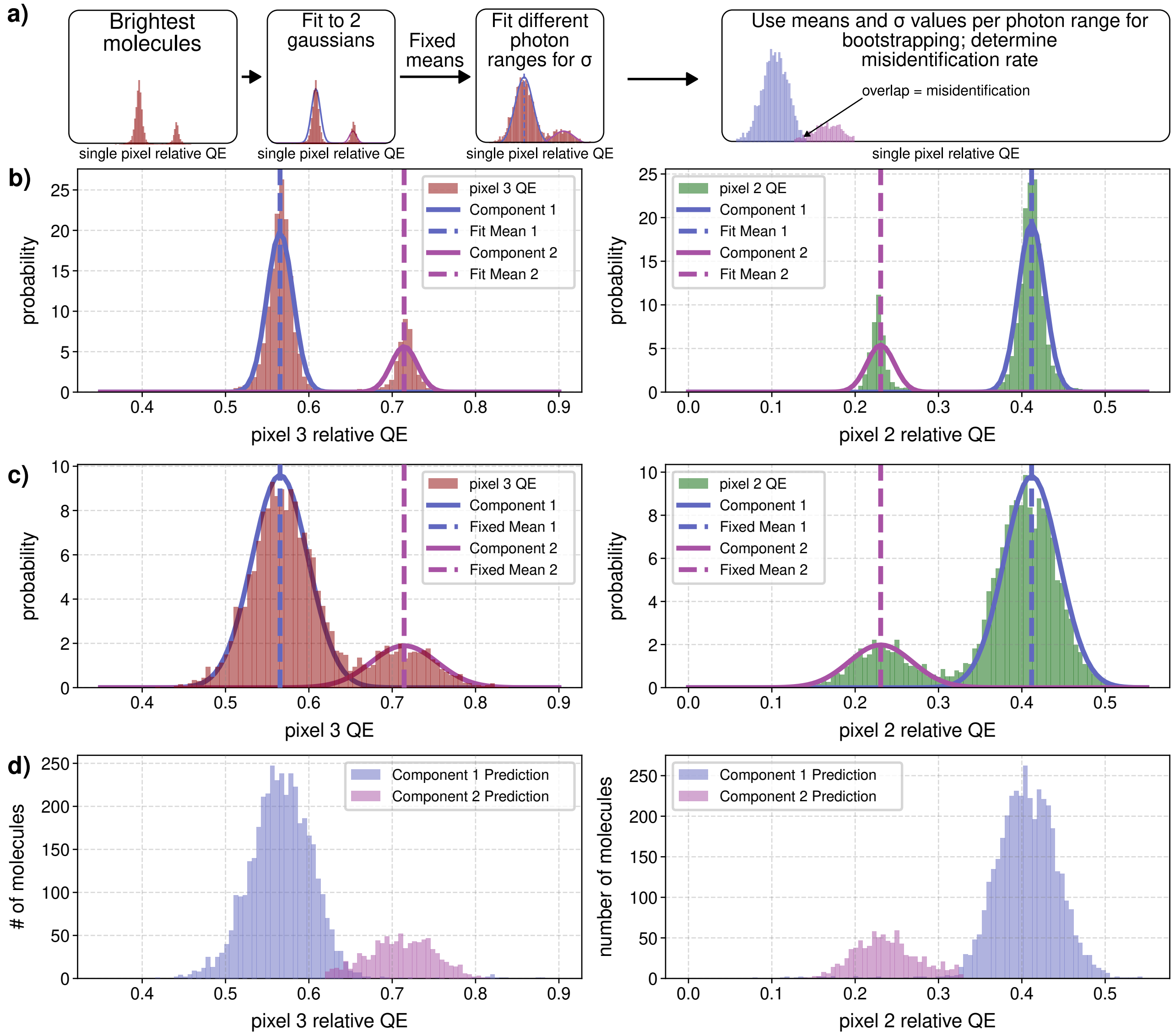


**Fig. S77. Workflow of quantifying dye mixtures. a)** High photon (here >20,000 photons) data of a mixture of ATTO 565 and ATTO 633 showing clear separation of populations is fit with a GMM to establish the mean pixel QE values for each component. **b)** Lower photon (here 1,760 – 2,508) data is fit to two populations using a GMM where the means are fixed to the value derived from the high photon data. **c)** Lower photon data sorted by population, these act as the ground truth for bootstrapping.

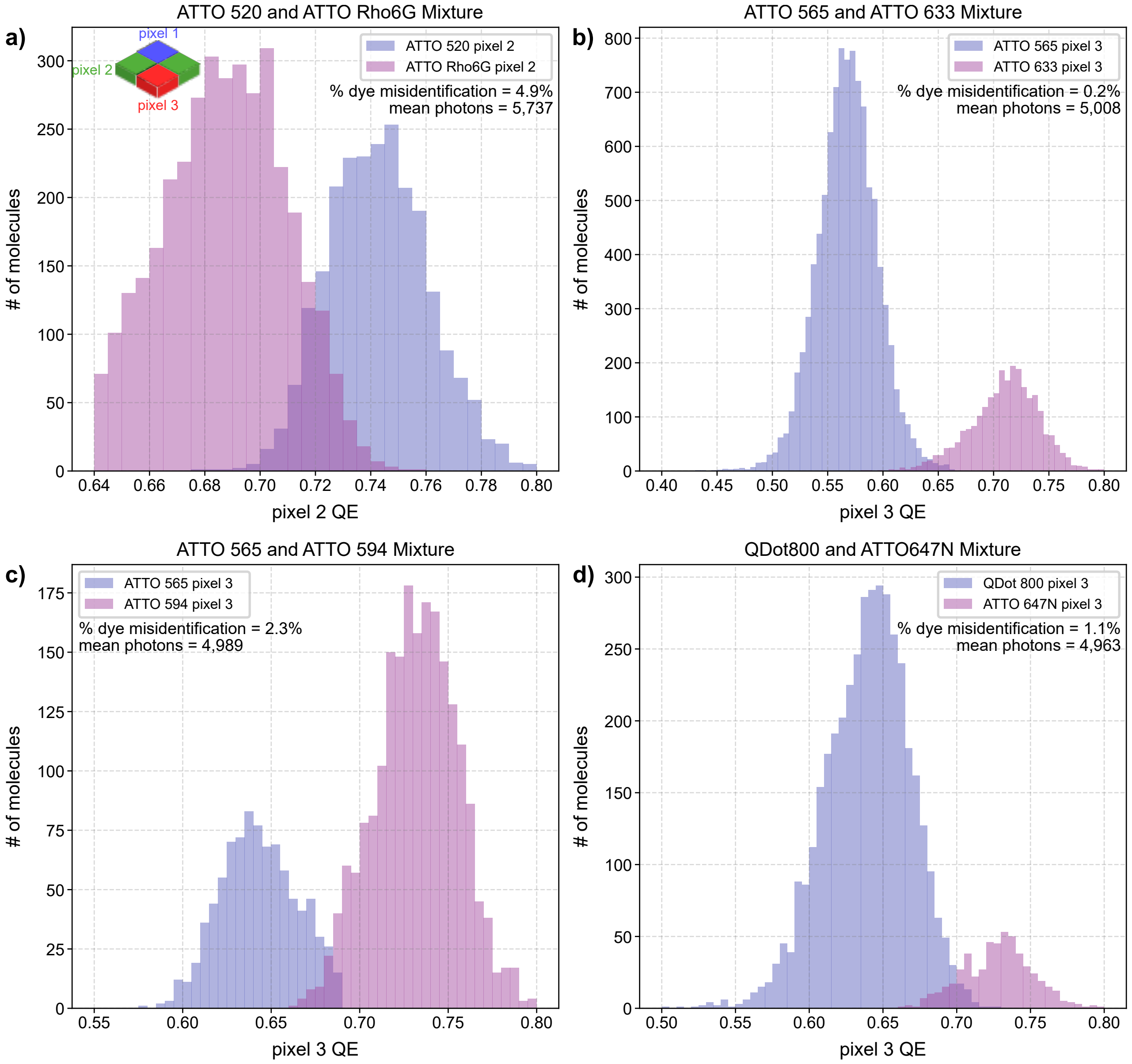


**Fig. S78. Predicted populations of experimental dye mixtures a)** ATTO 520 and ATTO Rho6G. **b)** ATTO 565 and ATTO 633. **c)** ATTO 565 and ATTO 594. **d)** ATTO 647N and Qdot 800

#### Supplementary Note S24: Quantum Dot Multiplexing

The accuracy for assigning the identity of each QD in a mixture can be estimated by generating a synthetic data set of quantum dots of known identity. To create this synthetic data set, we randomly draw 4,500 relative QE values from real data sets of each spectrally distinct quantum dot measured separately for a total of 27,000 data points (Fig. S79a). This synthetic data set is analysed with a Gaussian mixture model like before, but now has a ground truth assignment we can compare to (a confusion matrix Fig. 3e). In a mixture of even proportions, we expect an average true positive rate of greater than 94 %. In the worst case, Qdot 605 has the lowest true positive rate of 88 % due to its spectral proximity to Qdot 585, Qdot 705, and Qdot 800. In the best case scenario, the highly spectrally distinct Qdot 655 was always correctly assigned in our synthetic data set.

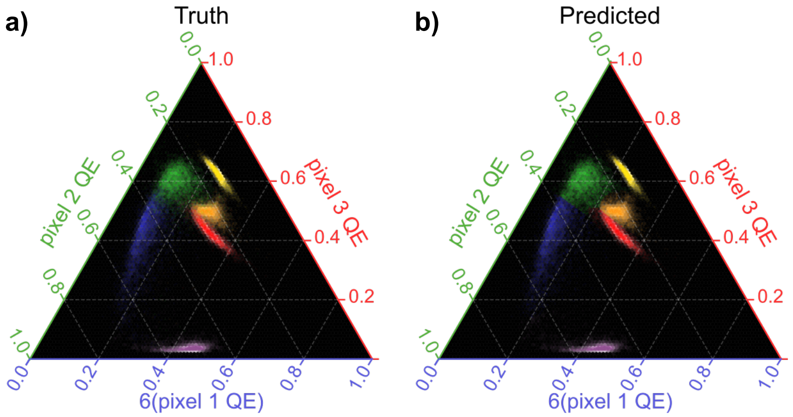


**Fig. S79.** **a)** Ground truth of synthetic data set derived from measurements of each species individually. **b)** Predicted identities of the synthetic data set (Qdot 525 in violet, Qdot 585 in blue, Qdot 605 in green, Qdot 655 in yellow, Qdot 705 in orange, and Qdot 800 in red).

#### Supplementary Note S25: FRET Efficiency Calculation

Because donor and acceptor information are measured within the same PSF in S$^3$M, measuring the FRET efficiency $E_{\mathrm{FRET}}$ requires prior knowledge of the relative QE of the donor and acceptor alone on the detector ($A_{\mathrm{Donor},p}$ and $A_{\mathrm{Acceptor},p}$). The relative QE of the detected signal $A_{\mathrm{data},p}$ will be a mixture of $A_{\mathrm{Donor},p}$ and $A_{\mathrm{Acceptor},p}$ weighted by some mixing factor $\alpha$.

$$A_{\mathrm{data},p} = \alpha A_{\mathrm{Acceptor},p} + (1-\alpha) A_{\mathrm{Donor},p} \tag{S25}$$

Using the three relative QE measurements, a linear equation can be solved for the mixing factor $\alpha$. This can then be converted to the apparent FRET efficiency $E_{\mathrm{App}}$ by weighting the mixing factor by the overall detection efficiency of the donor and acceptor $\eta_{\mathrm{Donor}}$ and $\eta_{\mathrm{Acceptor}}$ with the quantum yields of the donor and acceptor $\phi_{\mathrm{Donor}}$ and $\phi_{\mathrm{Acceptor}}$. The overall detection efficiency is the product of the efficiency for a photon of a particular wavelength to pass through our optical filters, as well as the QE of the detector (See Section S1).

$$E_{\mathrm{App}} = \frac{\alpha/\eta_{\mathrm{Acceptor}}\phi_{\mathrm{Acceptor}}}{\alpha/\eta_{\mathrm{Acceptor}}\phi_{\mathrm{Acceptor}} + (1-\alpha)/\eta_{\mathrm{Donor}}\phi_{\mathrm{Donor}}} \tag{S26}$$

While the quantum yields are not measured directly here, the reported quantum yields of Cy3 and Cy5 on duplexed DNA are 0.16 and 0.29 respectively.(29, 30) Using the spectra of the optical elements in our setup, and the spectra of the dyes Cy3 and Cy5, we can estimate $E_{\mathrm{App}}$ for the Holliday Junctions. We see a bimodal distribution of apparent FRET efficiencies with fitted means of 0.21 and 0.51, similar to the values seen in prior studies on the same Holliday Junction.(31, 32)

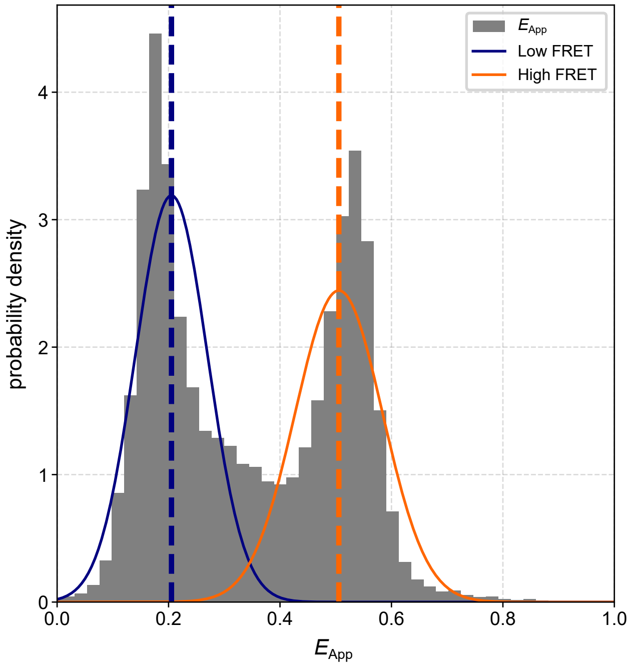


**Fig. S80.** Apparent FRET efficiencies of Holliday Junctions.

#### Supplementary Note S26: Resolving Spectral Shifts

We can define the minimum spectral fingerprint shift by determining the width of the relative pixel QE distributions of the two populations, effectively the precision of the relative pixel QE defined earlier as $\sigma_{\mathrm{colour}}$. We can provide a theoretical spectral fingerprint sensing curve as a function of the number of photons detected using the fit precision curve in Fig. 2d. If the populations are Gaussian and of similar widths, we can estimate the resolution needed as $2 \times \sigma_{\mathrm{colour}}$.
In the case of the FRET experiment, the average number of photons detected per frame from a molecule is 825, resulting in a $\sigma_{\mathrm{pixel2}}$ of 0.073 and 0.077 and $\sigma_{\mathrm{pixel3}}$ of 0.079 and 0.079 for the two Gaussian low/high FRET populations. In this case, the $\sigma_{\mathrm{pixel2}}$ and $\sigma_{\mathrm{pixel3}}$ of these populations are highly similar, so we will instead use an average sigma. These two populations can be separated so long as they are separated by a factor of 2 sigma, in this case 0.150 and 0.158.

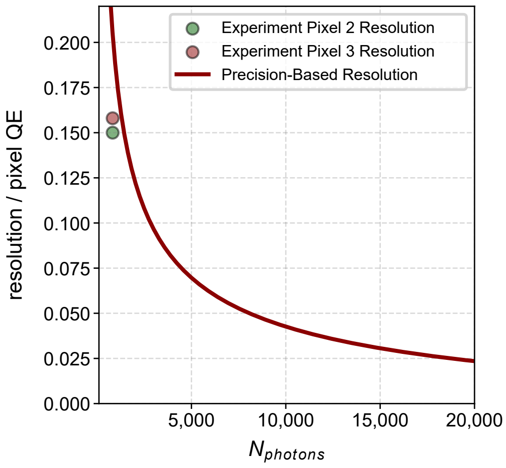


**Fig. S81.** Estimated resolution in spectral shifts based on precision values (solid red line). Experimentally derived resolutions of each pixel type are shown as scattered points.

Both of these values are below the observed change in the mean relative pixel QE values observed for the High-FRET and Low-FRET populations of 0.19 and 0.18 for pixel 2 and pixel 3 respectively.

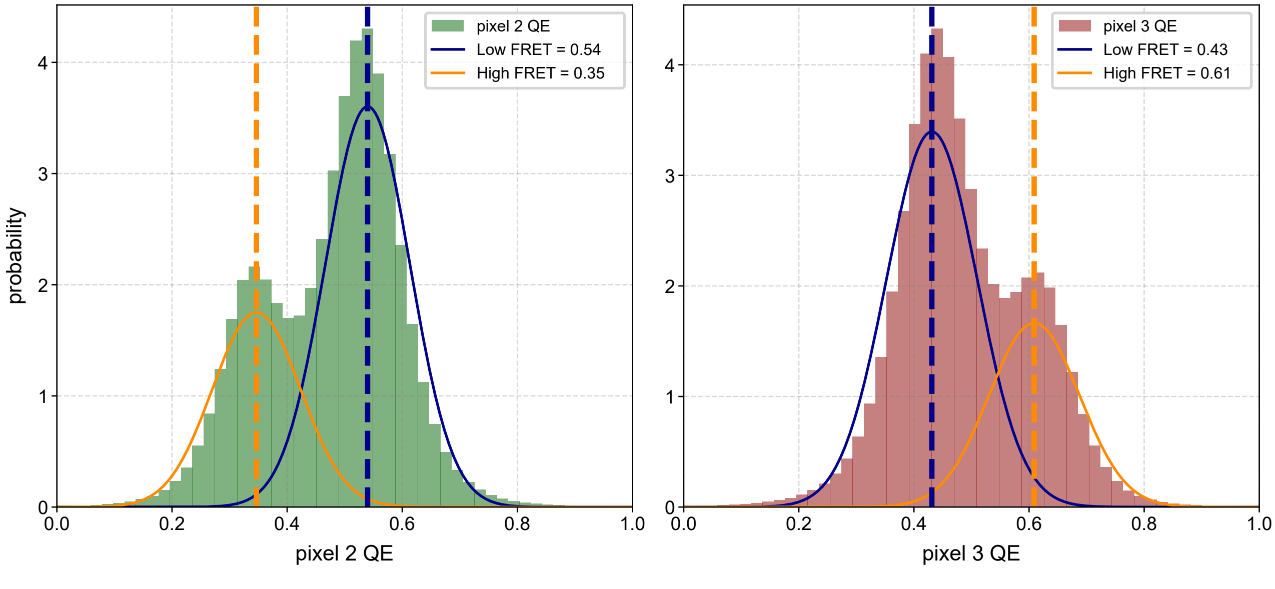


**Fig. S82.** Relative pixel QE values per frame from experimental FRET data of Holliday Junctions. Low-FRET and High-FRET populations are fit with a two Gaussian model.

Additionally, the number of samples of each population is important in determining if a change in relative pixel QE is due to noise of our measurement or genuine change in state. Assuming a false-positive rate of 0.025, we can express the change in relative pixel QE we wish to resolve $\Delta$relative pixel QE as a function of the experimental variation of each state $\sigma_{\mathrm{pixel\ n}}$ as well as the minimum number of samples per state $N_{\mathrm{frames}}$.(33)

$$\Delta \text{relative pixel QE} = 2\left(\frac{\sigma_{\text{Low FRET}}}{\sqrt{N_{\text{frames}}}} + \frac{\sigma_{\text{High FRET}}}{\sqrt{N_{\text{frames}}}}\right) \tag{S27}$$

Using this equation and the experimentally derived values, we can determine that we only need an $N_{frames} = 4$ to accurately sense a change in relative pixel QE. In principle, longer trajectories can sense even smaller shifts in relative pixel QE. In the Holliday Junction experiments, the median number of frames per changepoint are 43 and 47 for the low-FRET and high-FRET states, resulting in sensitivities to spectral fingerprint below 0.05.

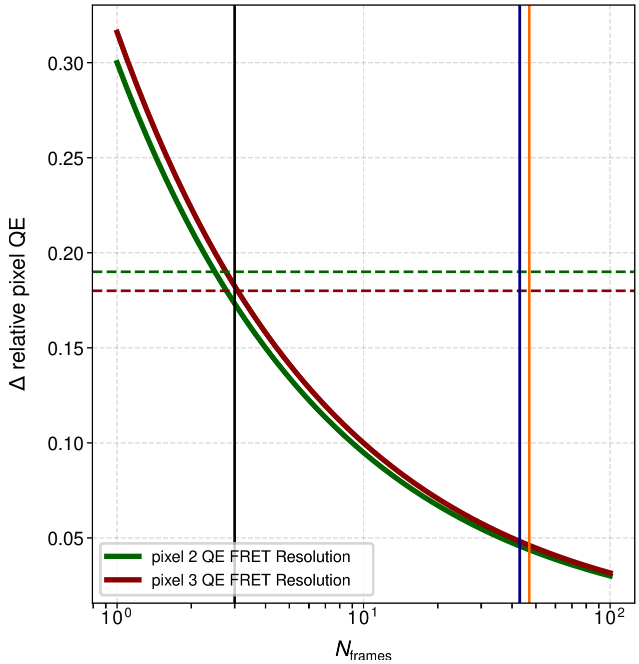


**Fig. S83.** Resolution of relative pixel QE change as a function of frames detected in each state. Horizontal green and red dashed lines indicate the difference in relative pixel QE for pixels 2 and 3 for the experimentally observed FRET states. The black vertical line indicates the minimum number of frames needed to detect the experimentally observed change in relative pixel QE, the blue and orange vertical lines indicate the median number of frames for the low-FRET and high-FRET changepoints.

#### Supplementary Note S27: Example Localisations of Holliday Junctions

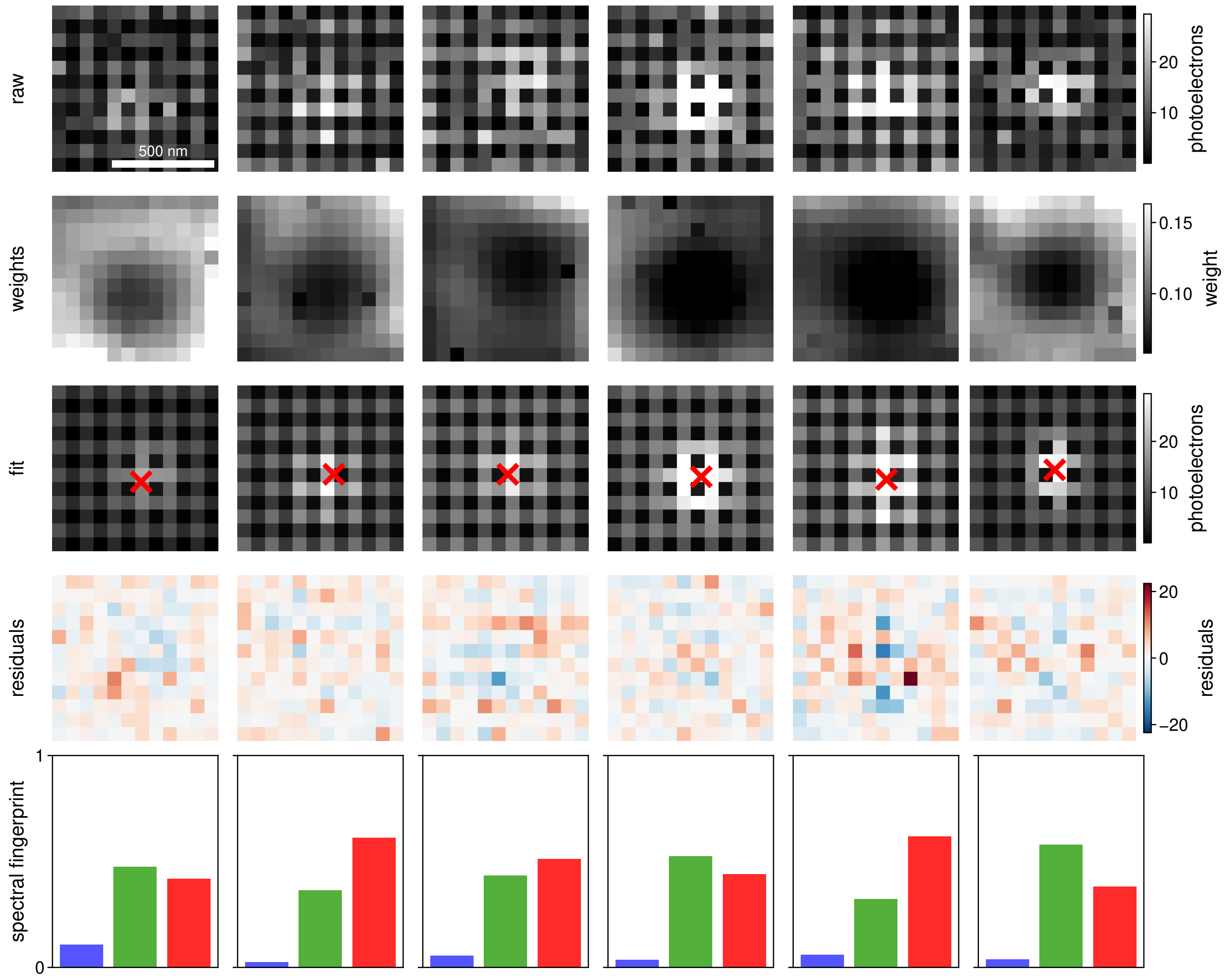


**Fig. S84. Fitting FRET localisations of Holliday Junction data.** Raw localisations, weights, reconstructed PSFs with localisations, residuals, and extracted spectral fingerprints of 5 different molecules.

#### Supplementary Note S28: Example Localisations of $\alpha$-Synuclein

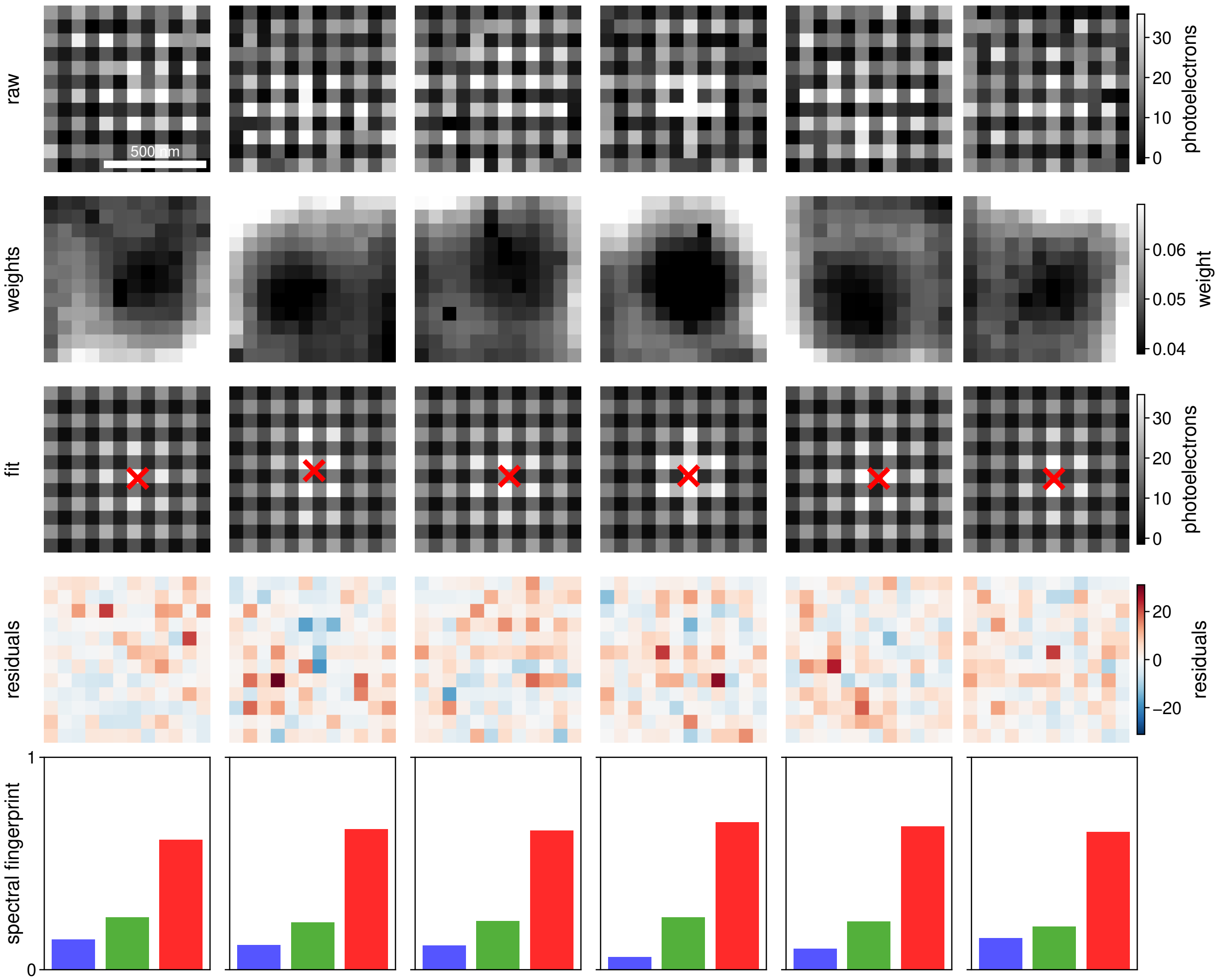


**Fig. S85. Fitting Nile Red localisations of $\alpha$-Synuclein data.** Raw localisations, weights, reconstructed PSFs with localisations, residuals, and extracted spectral fingerprints of 5 different molecules.

#### Supplementary Note S29: Fourier Ring Correlation for *S. aureus*

The FRC curve was calculated using a Python port of the MATLAB code written by Nieuwenhuizen *et al.*(21) Error bounds are $\pm 1$ standard deviation across 20 independent random half-splits.

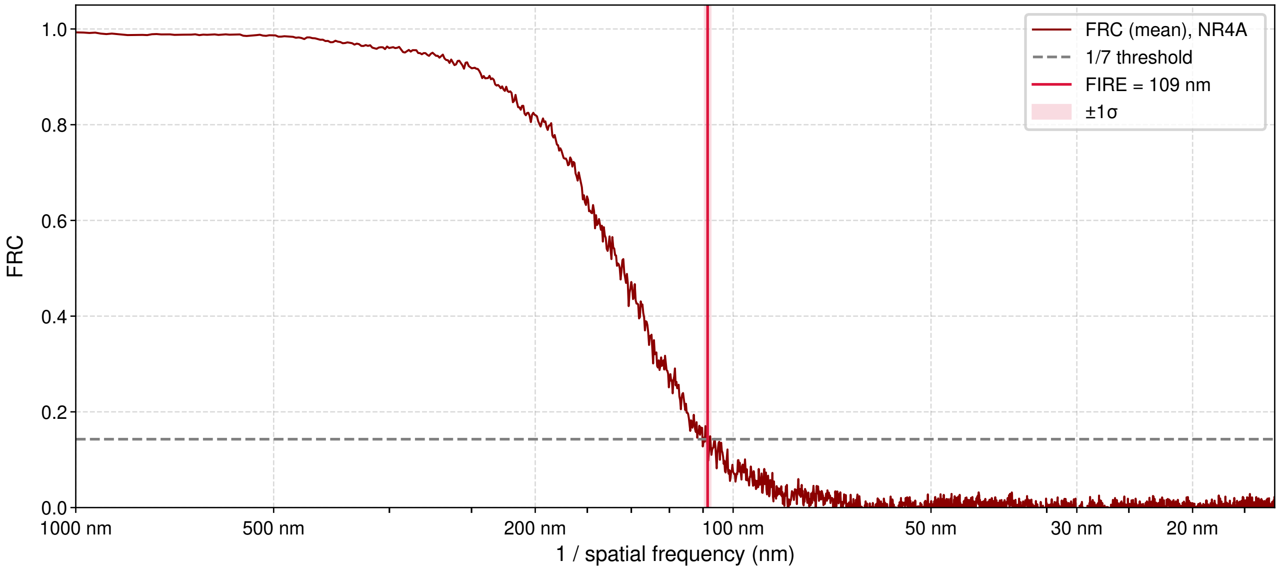


**Fig. S86.** FRC curve for the NR4A PAINT experiment, showing the 109 nm resolution achieved.

#### Supplementary Note S30: Nile Red and NR4A Characterisation

Both Nile Red and its derivative NR4A show solvatochromic properties that provide information on the local environment. Here we demonstrate that for both dyes on a bulk level, the emission is highly dependent on both the presence of proton donors as well as the solvent polarity. By using a mixture of acetonitrile and water, we can tune the proton donation rate of our mixture while maintaining polarity.(34) As we increase the ratio of water to acetonitrile, the proton donation ability increases, and the emission is red-shifted. Additionally, we find that by changing the overall polarity of the solvent, we find a similar red-shift in the emission of both Nile Red and NR4A. However, in the case of NR4A, we were unable to measure the fluorescence in very low polarity solvents due to poor solubility, but literature suggests in very low polarity solvents like dioxane the peak emission can be as low as ∼580 nm.(35)

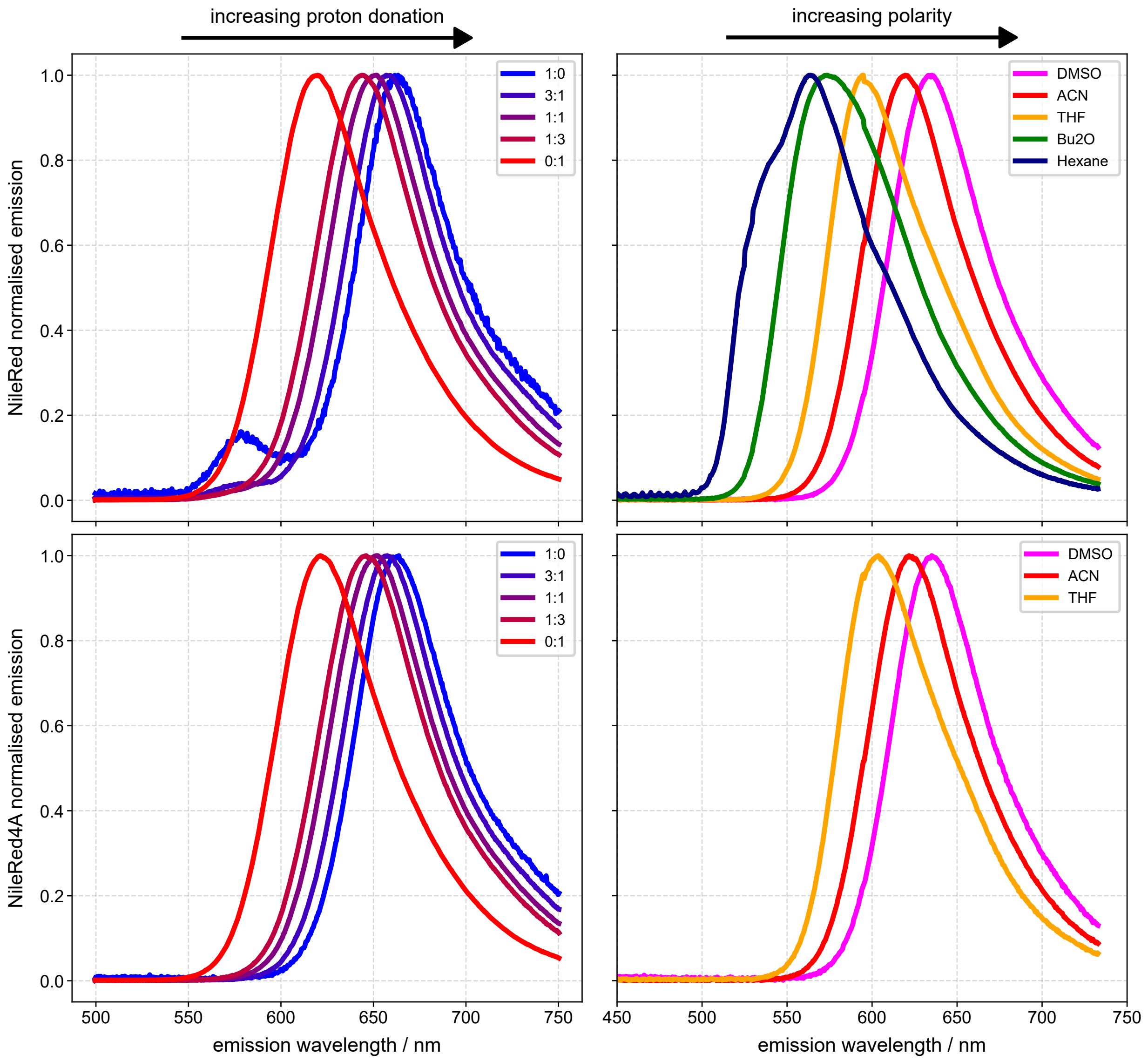


**Fig. S87.** Emission spectra of Nile Red (top row) and NR4A (bottom row) in different solvent conditions. Left shows dependence on proton donation, legends include the ratio of water:acetonitrile. Right shows dependence on polarity, with different solvents shown in the legend. DMSO = Dimethyl sulfoxide, ACN = Acetonitrile, THF = Tetrahydrofuran, Bu2O = Di-butyl ether.

#### Supplementary Note S31: Shift-Invariance

Because each pixel type has different spectral sensitivity in $S^3M$, the detected signal of a molecule is not just a function of the PSF width and where on the pixel it is centred, but the underlying spectra and the pixel type. $S^3M$ fits both position and spectral fingerprint jointly, allowing it to accurately determine both parameters, without any bias relating to which pixel type it is centred on. To demonstrate the shift-invariance of the extracted parameters, we simulated PSFs of three fluorophores across the visible spectrum (ATTO 488, ATTO 565, and ATTO 647N), each time changing where the molecule is centred. In each simulation, the molecule emits 1,000 photons and has an average background per pixel of 10 photons. The spectral fingerprint and position are then fit for that simulated PSF. This process is repeated 10,000 times for each position, sampling 25 points across the 2×2 Bayer pattern. For each of the dyes simulated, we show an exemplar 5×5 grid of positions across the Bayer pattern where the molecule is centred to highlight the changing PSF pattern as a function of molecular position (Fig. S88–S90). We also show the fit spectral fingerprint (Fig. S91–S93), and the positional bias of each fit localisation (Fig. S94–S96). In each instance, we find little bias in the fit spectral fingerprint and position across the pattern—each fit is centred on the ground truth, and the distribution of fit values does not vary significantly as a function of position.

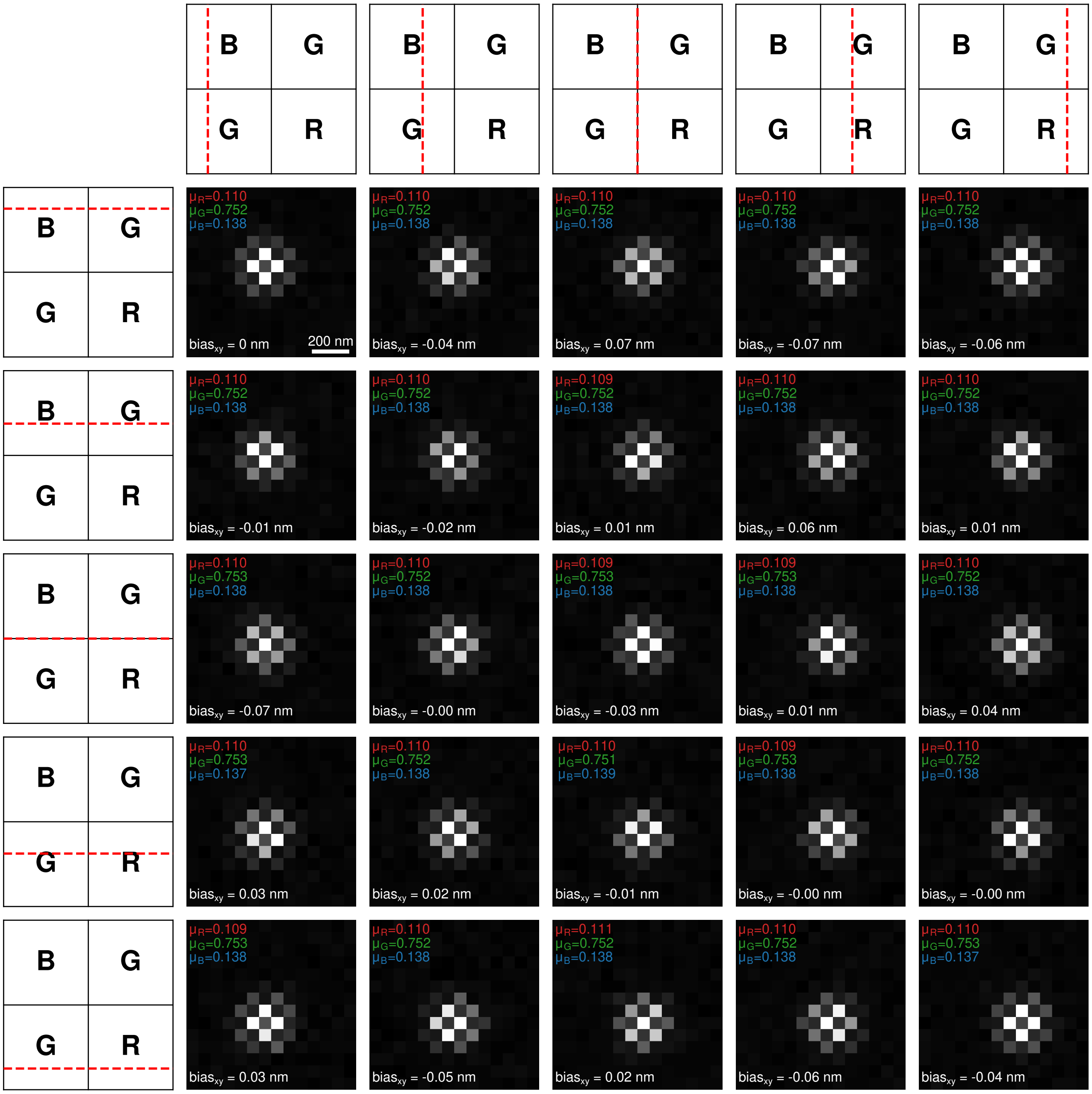


**Fig. S88.** Simulated PSFs (4,000 photons, 4 background photons) of ATTO 488 as a function of relative position on the Bayer pattern. μ and bias values are from 1,000 photon fits of Fig. S91 and Fig. S94.

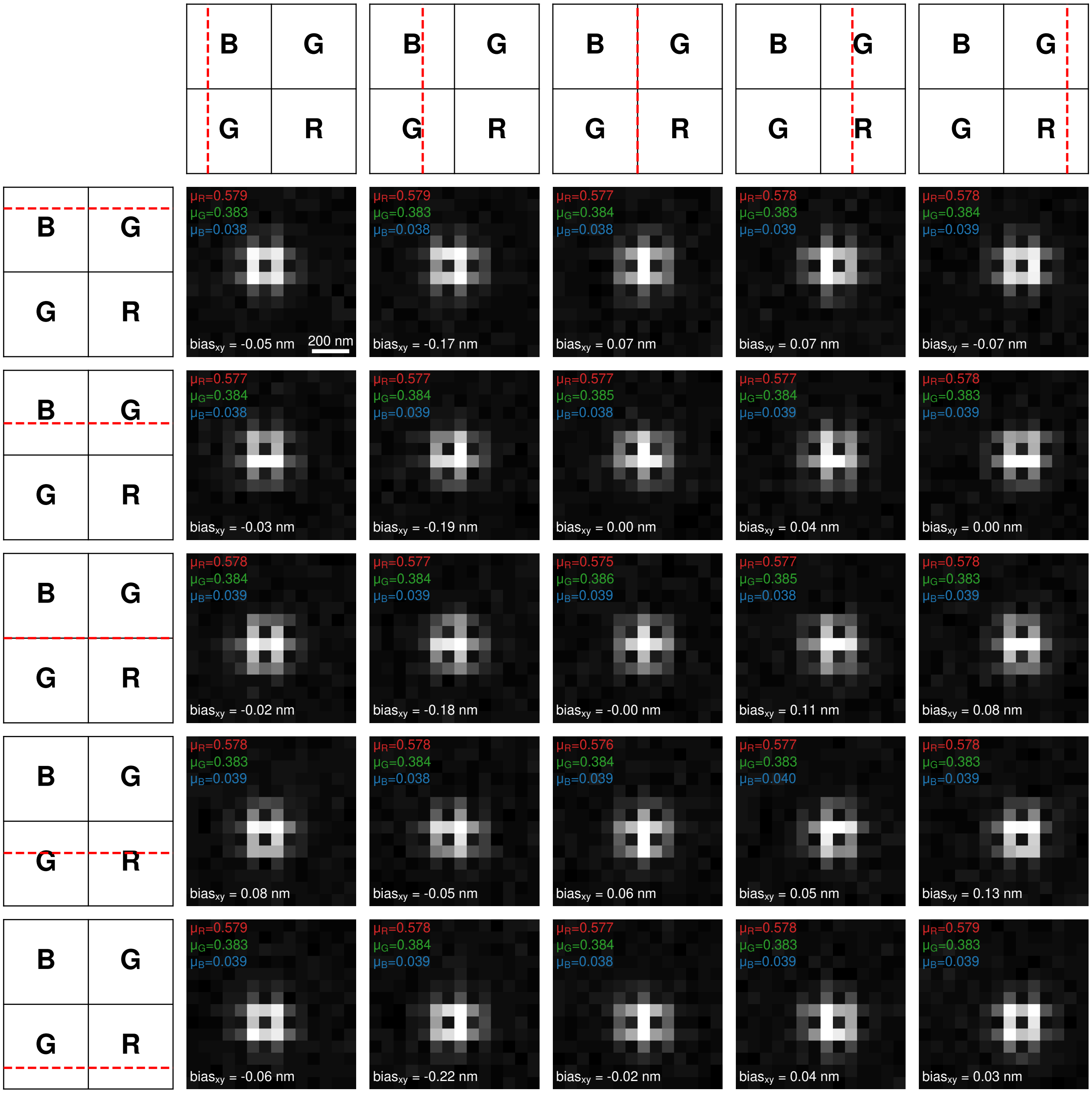


**Fig. S89.** Simulated PSFs (4,000 photons, 4 background photons) of ATTO 565 as a function of relative position on the Bayer pattern. μ and bias values are from 1,000 photon fits of Fig. S92 and Fig. S95.

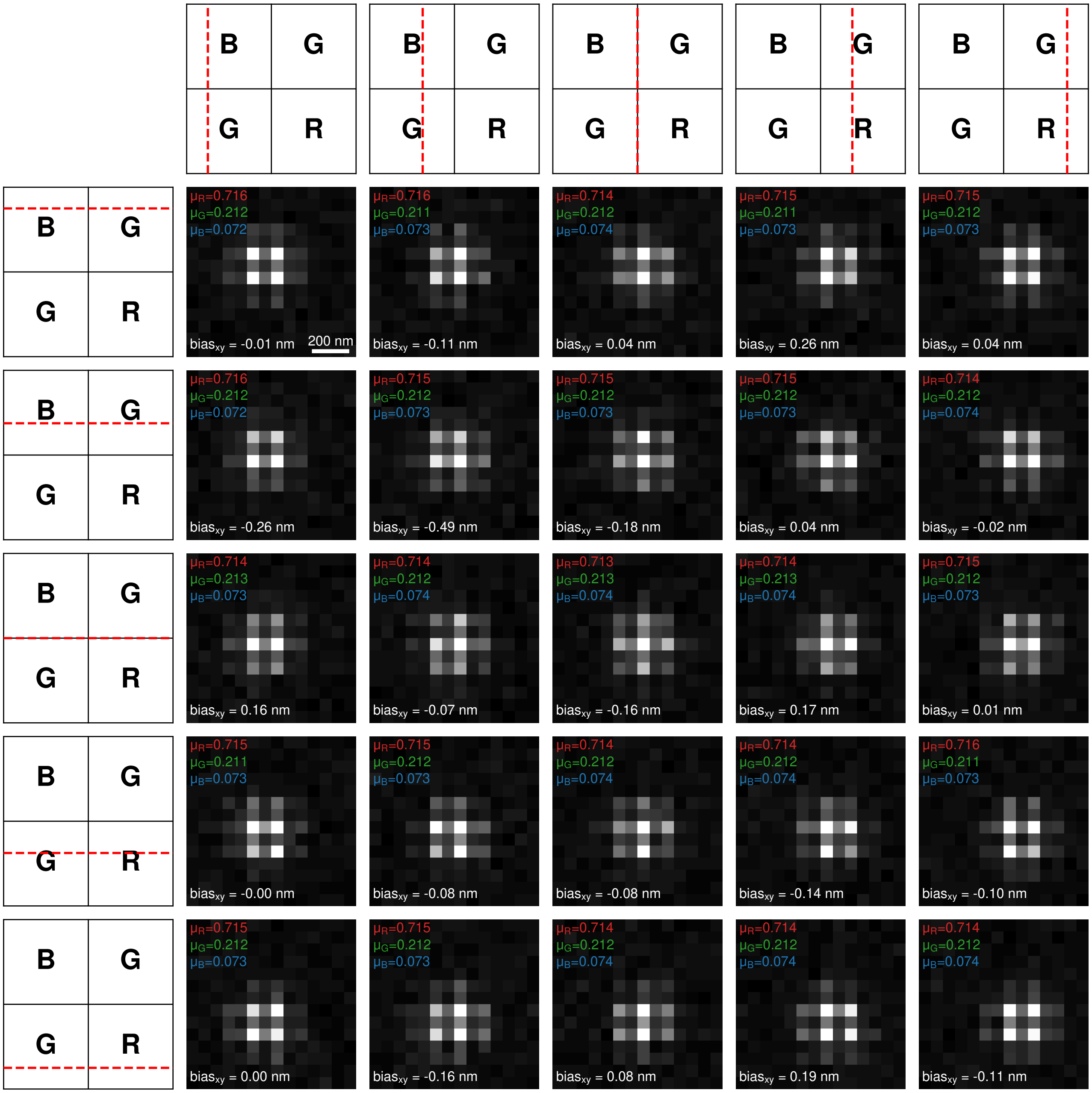


**Fig. S90.** Simulated PSFs (4,000 photons, 4 background photons) of ATTO 647N as a function of relative position on the Bayer pattern. $\mu$ and bias values are from 1,000 photon fits of Fig. S93 and Fig. S96.

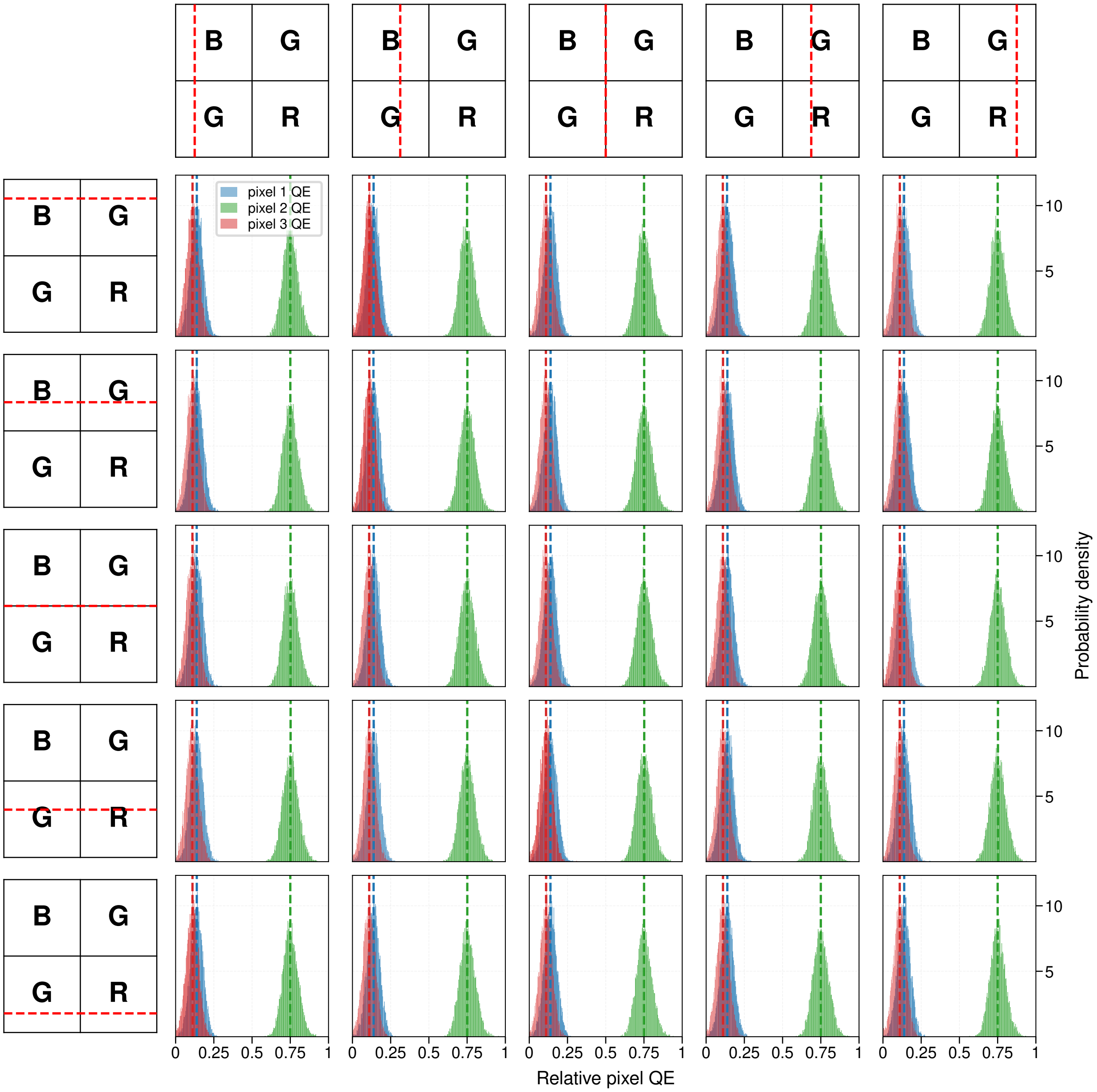


**Fig. S91.** Fit spectral fingerprints of ATTO 488 as a function of relative position on the Bayer pattern. The ground truths of each pixel QE are shown as dashed vertical lines. Here each localisation was simulated as 1000 photons, with 10 background photons.

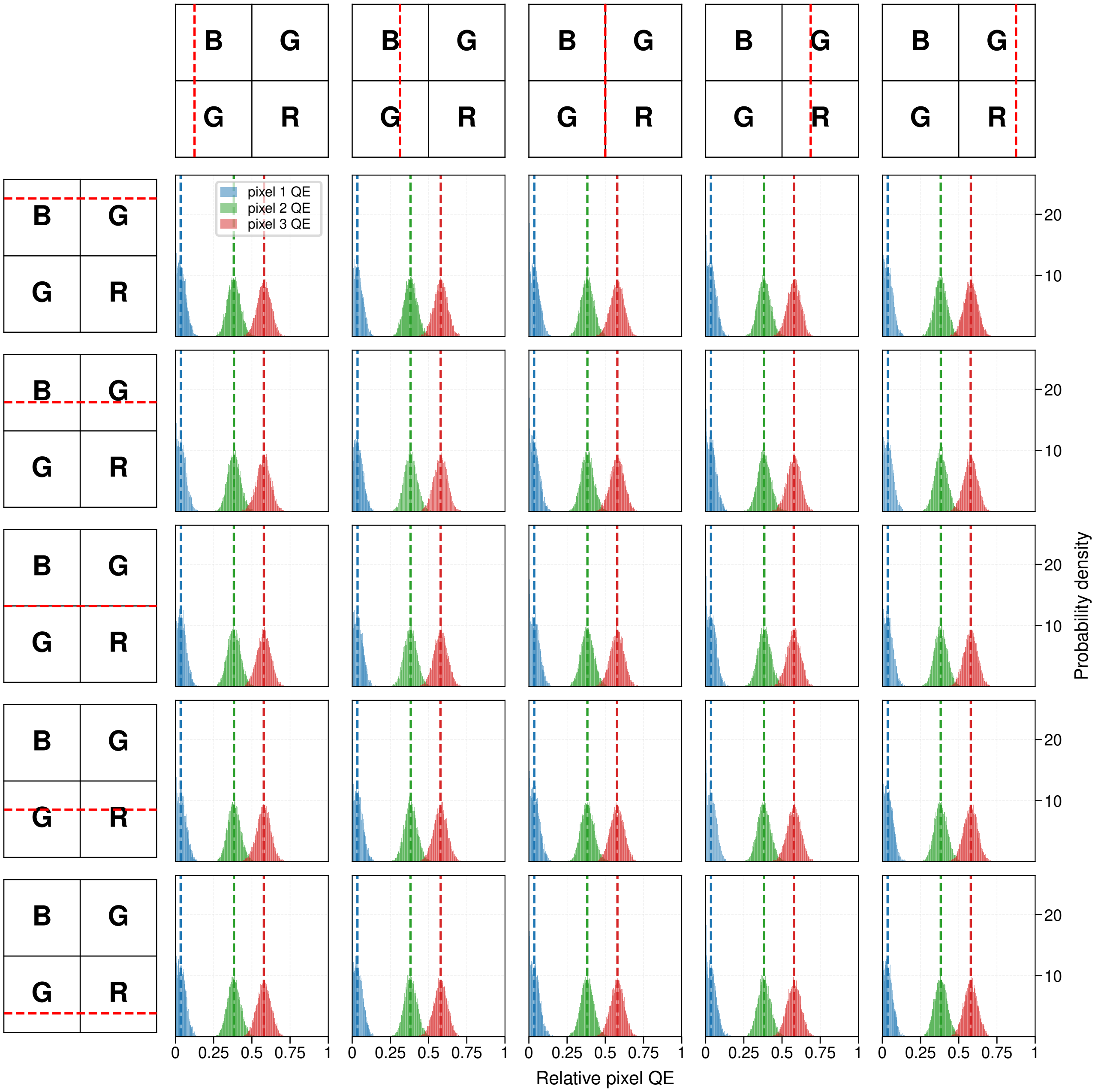


**Fig. S92.** Fit spectral fingerprints of ATTO 565 as a function of relative position on the Bayer pattern. The ground truths of each pixel QE are shown as dashed vertical lines. Here each localisation was simulated as 1000 photons, with 10 background photons.

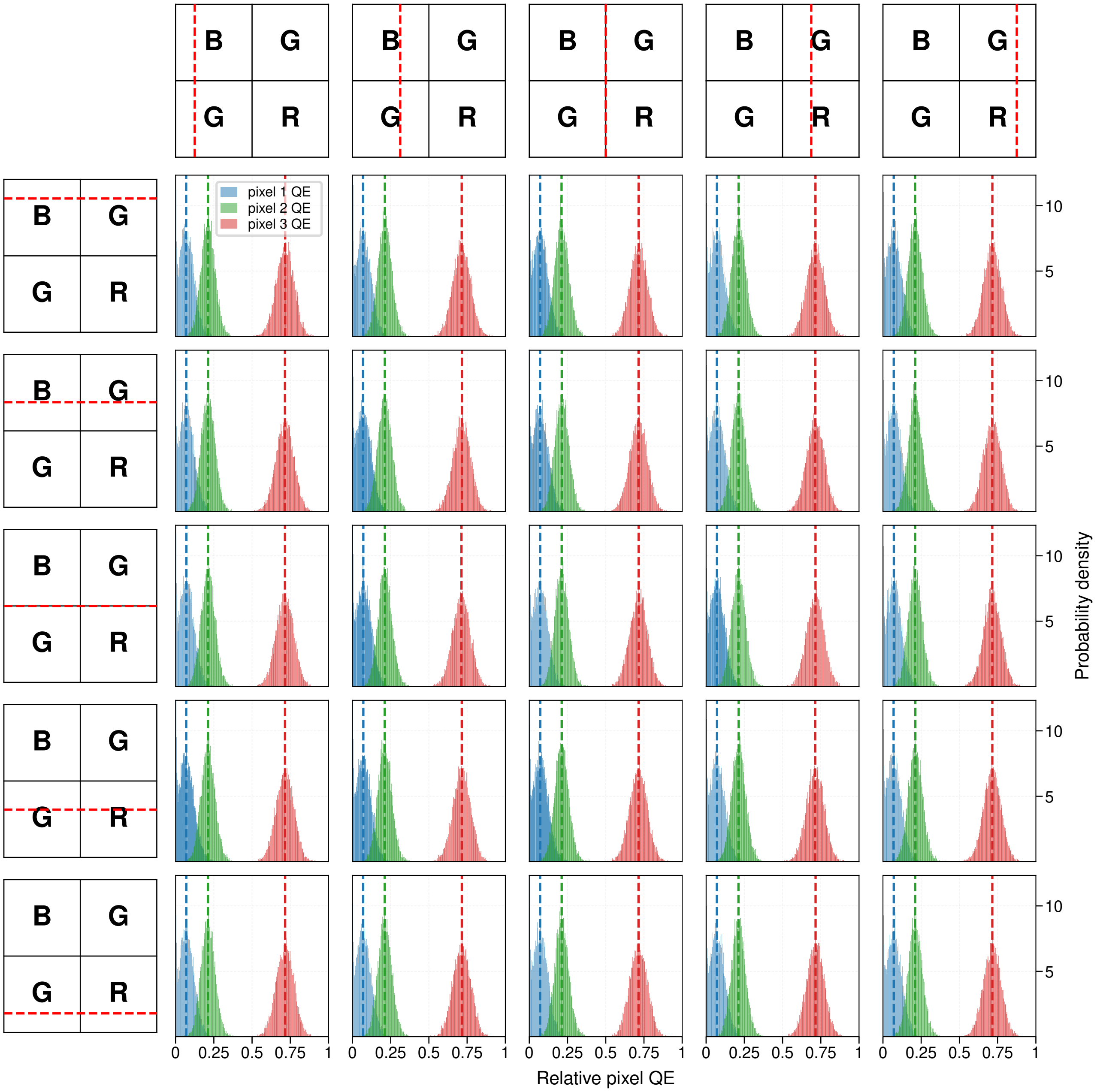


**Fig. S93.** Fit spectral fingerprints of ATTO 647N as a function of relative position on the Bayer pattern. The ground truths of each pixel QE are shown as dashed vertical lines. Here each localisation was simulated as 1000 photons, with 10 background photons.

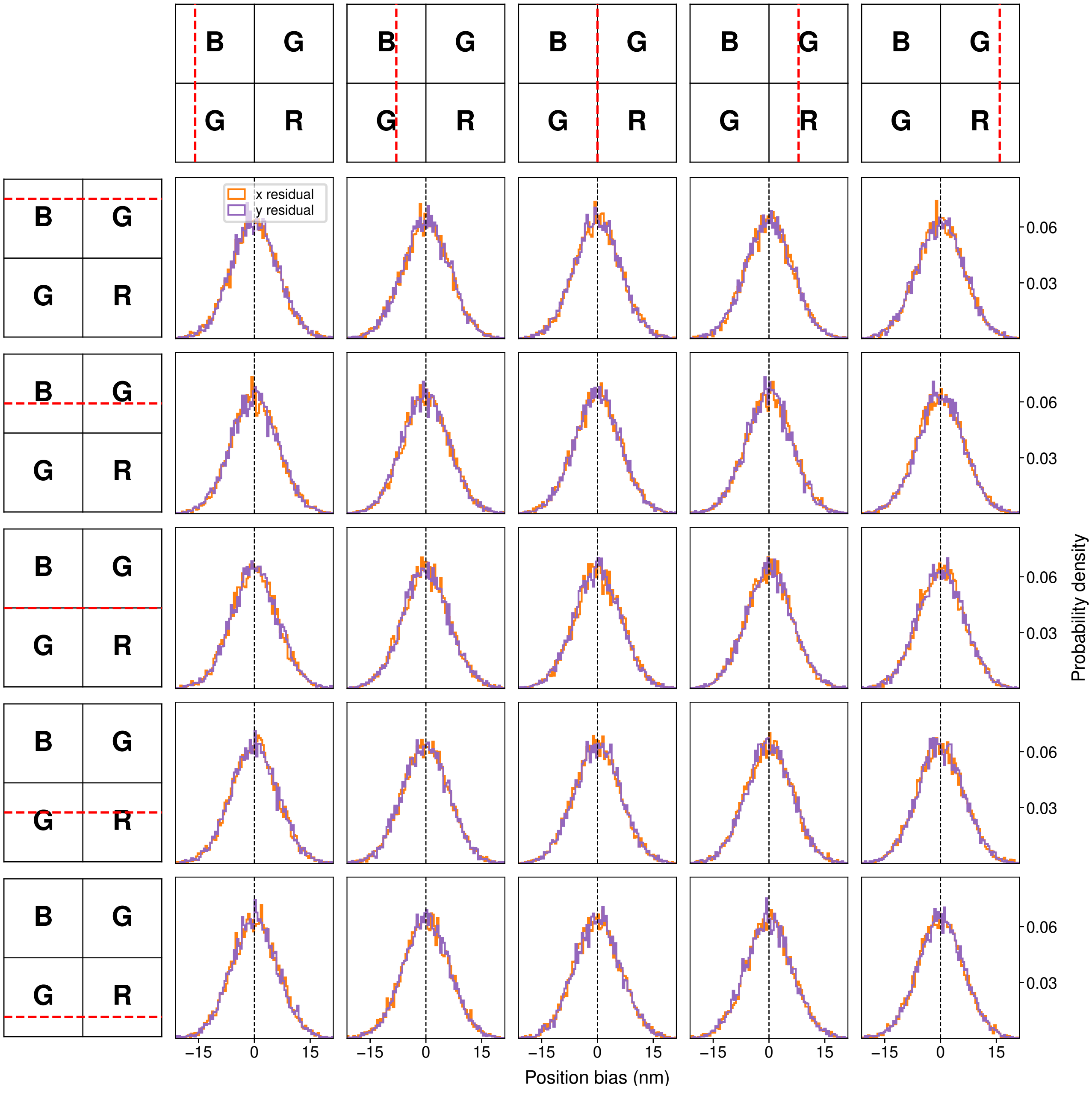


**Fig. S94.** Residual of the fit position in x and y of ATTO 488 as a function of relative position on the Bayer pattern. The true unbiased position is shown as a dashed vertical line. Here each localisation was simulated as 1000 photons, with 10 background photons.

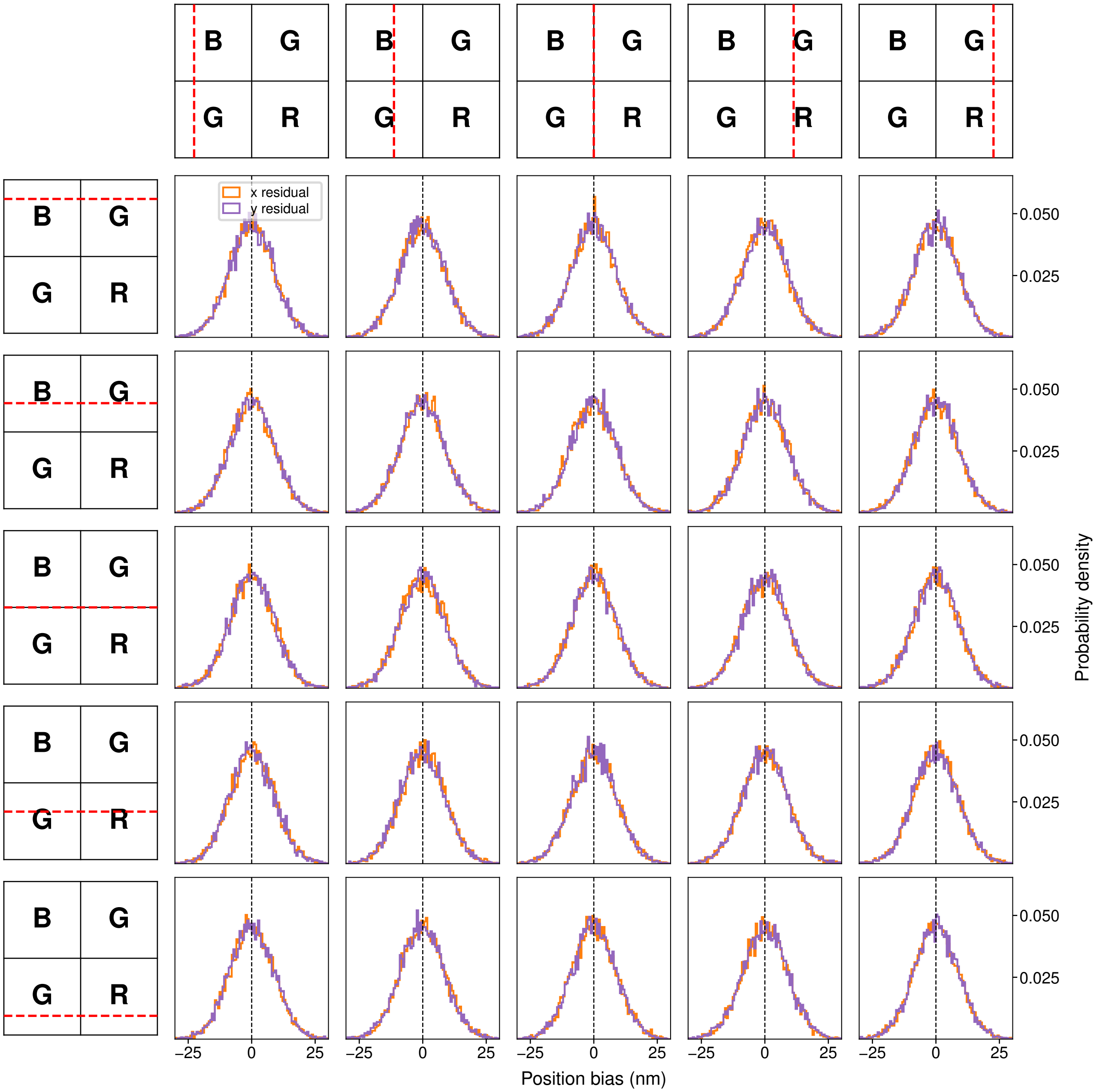


**Fig. S95.** Residual of the fit position in x and y of ATTO 565 as a function of relative position on the Bayer pattern. The true unbiased position is shown as a dashed vertical line. Here each localisation was simulated as 1000 photons, with 10 background photons.

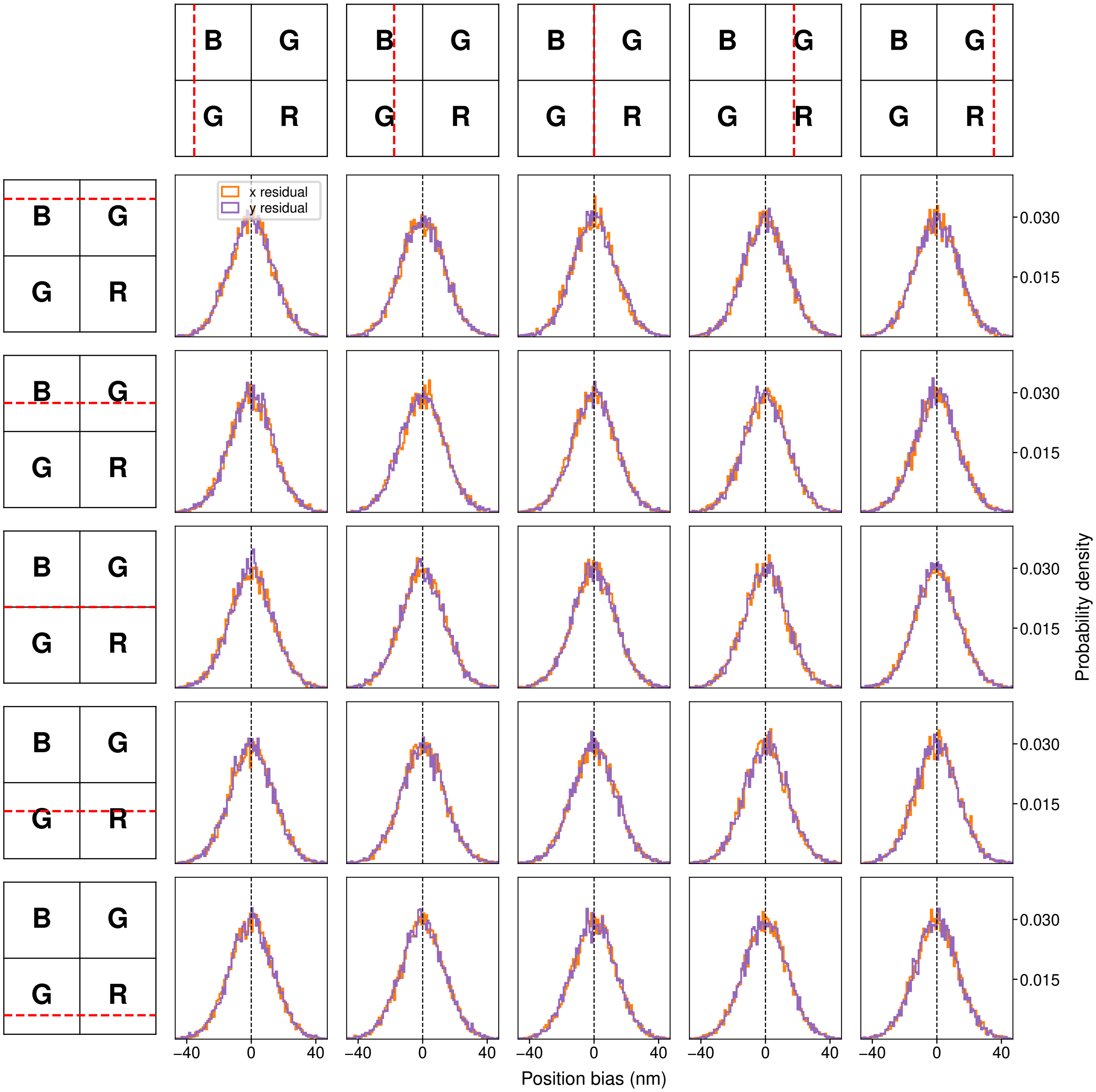


**Fig. S96.** Residual of the fit position in x and y of ATTO 647N as a function of relative position on the Bayer pattern. The true unbiased position is shown as a dashed vertical line. Here each localisation was simulated as 1000 photons, with 10 background photons.

#### Supplementary Note S32: Zoom-Ins of Experimental PSFs

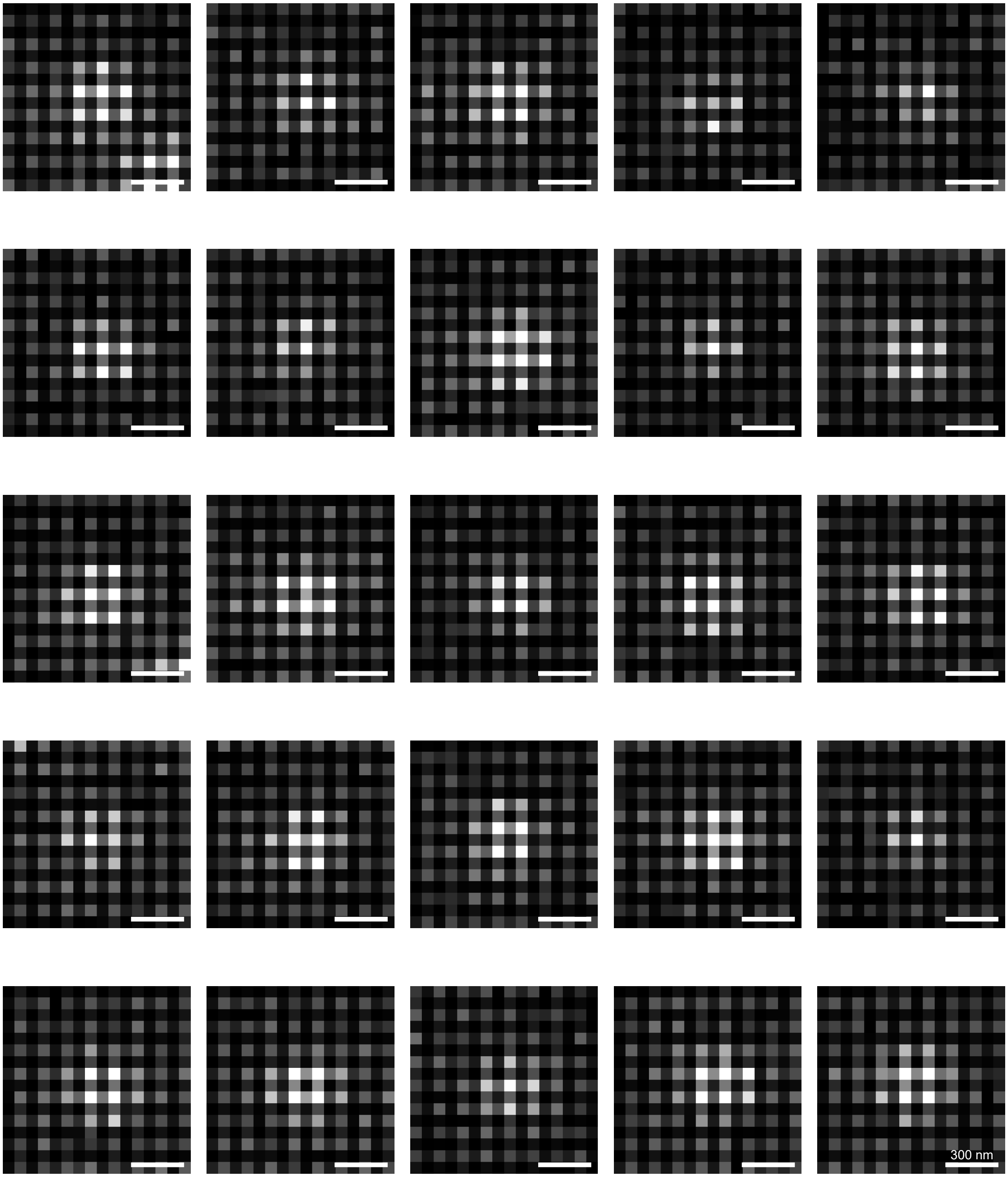


**Fig. S97.** Zoom-Ins of ATTO 633 molecules. Scale bar is 300 nm.

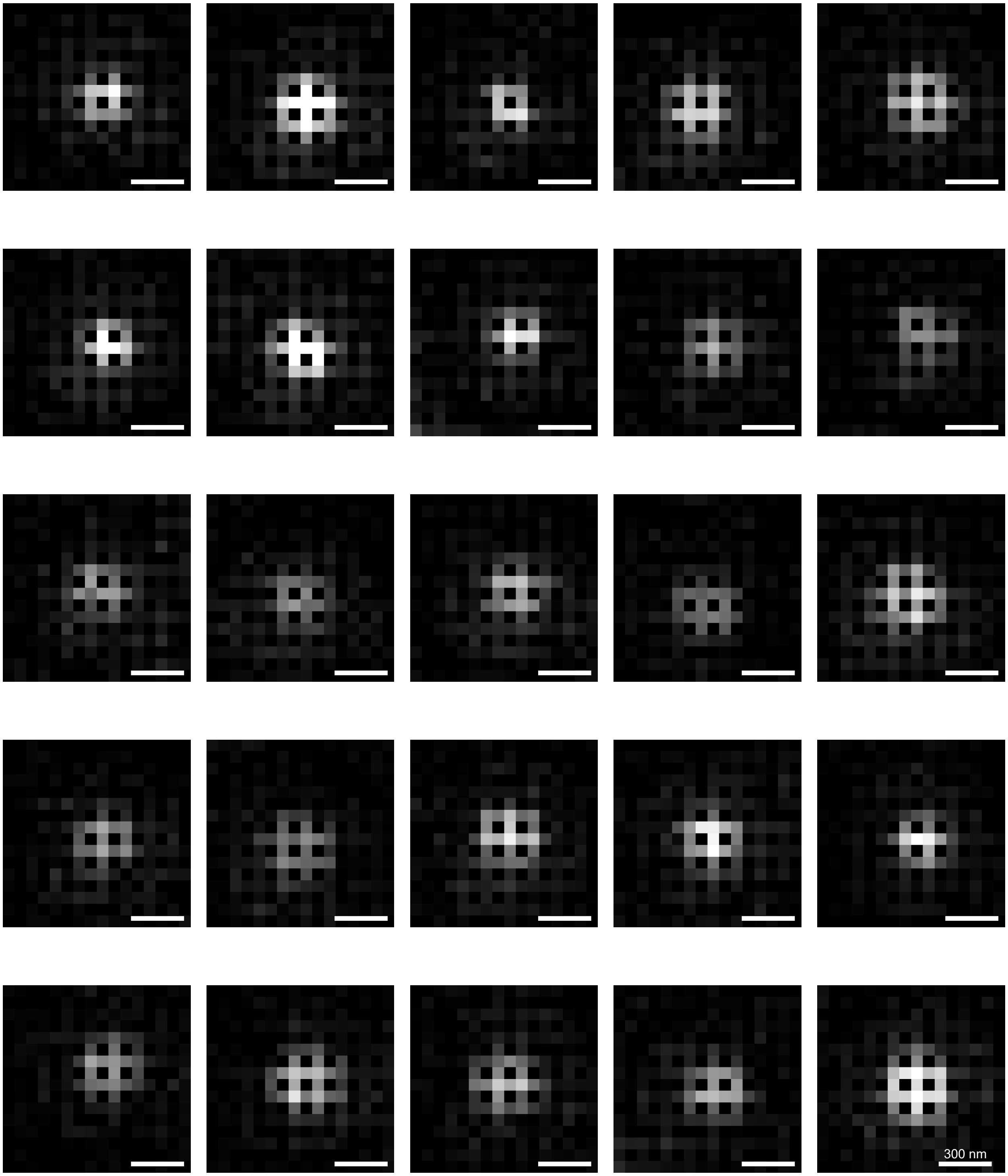


**Fig. S98.** Zoom-Ins of ATTO 565 molecules. Scale bar is 300 nm.

#### Supplementary Note S33: Imaging Conditions

| Sample | $\lambda_{exc}$/ nm (power $\rho$ in kW/cm$^2$) | Dichroic Mirrors | Emission Filters | $T_{exposure}$/ ms ($N_{frames}$/FOV) |
|---|---|---|---|---|
| HeLa Cell dSTORM (AF647) | 638 (1.18) | Di03-R405/488/561/635-t1 (Semrock) | BLP01-635R-25 (Semrock) | 40 (50,000) |
| MASSIVE Cells DNA-PAINT (Cy3B, ATTO 655) | 561 (0.14), 638 (0.17) | Di03-R405/488/561/635-t1 (Semrock) | NF03-405/488/561/635E-25 (Semrock), BSP01-785R (Semrock) | 75 (60,000) |
| $\alpha$-Synuclein aggregates PAINT (Nile Red) | 515 (0.38) | Di03-R514-t1 (Semrock) | FF01-515-LP (Semrock), FF01-650/200-25 (Semrock) | 50 (50,000) |
| PAINT imaging of *Staphylococcus aureus* (Nile Red 4A) | 515 (0.32) | Di03-R514-t1 (Semrock) | FF01-515-LP (Semrock), FF01-650/200-25 (Semrock) | 50 (50,000) |
| Holliday Junction FRET (Cy3, Cy5)* | 515 (0.57) | Di03-R514-t1 (Semrock) | FF01-515-LP (Semrock), FF01-650/200-25 (Semrock) | 10 (2,500) |
| 0.1 µm TetraSpeck Beads | 488 ($3.5\times10^{-4}$–$3.7\times10^{-2}$) 561 ($7.5\times10^{-4}$–$4.3\times10^{-2}$) 638 ($3.9\times10^{-3}$–$3.0\times10^{-2}$) | Di03-R405/488/561/635-t1 (Semrock) | NF03-405/488/561/635E-25 (Semrock), BSP01-785R (Semrock), BLP01-488R-25 (Semrock) | 50 (200) |
| Single-Molecule Tracking (AF488, AF555, JF646) | 488 (0.08), 561 (0.04), 638 (0.18) | Di03-R405/488/561/635-t1 (Semrock) | NF03-405/488/561/635E-25 (Semrock), BSP01-785R (Semrock), BLP01-488R-25 (Semrock) | 50 (1,000) |
| Z-Defocus Experiments | 488 (0.008) | Di03-R405/488/561/635-t1 (Semrock) | BLP01-488R-25 (Semrock), FF01-520/44-25 (Semrock) | 50 (100 per Z) |

**Table S6.** Imaging conditions for super-resolution and single-molecule spectroscopy experiments. * ZWO ASI 585MC was used for this experiment.

| Sample | $\lambda_{exc}$/ nm (power $\rho$ in kW/cm$^2$) | Dichroic Mirrors | Emission Filters | $T_{exposure}$/ ms ($N_{frames}$/FOV) |
|---|---|---|---|---|
| ATTO 488 | 488 (0.16) | Di03-R405/488/561/635-t1 (Semrock) | BLP01-488R-25 (Semrock), FF01-520/44-25 (Semrock) | 100 (1,000) |
| ATTO 514 | 515 (0.29) | Di03-R514-t1 (Semrock) | FF01-540/80 (Semrock), FF01-515-LP (Semrock) | 100 (1,000) |
| ATTO 520 | 515 (0.29) | Di03-R514-t1 (Semrock) | FF01-540/80 (Semrock), FF01-515-LP (Semrock) | 100 (1,000) |
| ATTO Rho6G | 515 (0.29) | Di03-R514-t1 (Semrock) | FF01-515-LP (Semrock) | 100 (1,000) |
| ATTO 565 | 561 (0.18) | Di03-R405/488/561/635-t1 (Semrock) | BLP02-561R-25 (Semrock), FF01-582/64-25 (Semrock) | 100 (1,250) |
| ATTO 594 | 561 (0.38) | Di03-R488/561-t1 (Semrock) | NF03-405/488/561/635E-25 (Semrock), BSP01-785R (Semrock) | 100 (1,250) |
| ATTO 620 | 638 (0.08) | Di03-R405/488/561/635-t1 (Semrock) | BLP01-635R-25 (Semrock) | 100 (1,250) |
| ATTO 633 | 638 (0.42) | Di03-R405/488/561/635-t1 (Semrock) | NF03-405/488/561/635E-25 (Semrock), BSP01-785R (Semrock) | 100 (1,250) |
| ATTO 647N | 638 (0.42) | Di03-R405/488/561/635-t1 (Semrock) | NF03-405/488/561/635E-25 (Semrock), BSP01-785R (Semrock) | 100 (1,250) |
| ATTO 655 | 638 (0.42) | Di03-R405/488/561/635-t1 (Semrock) | NF03-405/488/561/635E-25 (Semrock), BSP01-785R (Semrock) | 100 (1,000) |
| ATTO 700 | 638 (0.49) | Di03-R405/488/561/635-t1 (Semrock) | NF03-405/488/561/635E-25 (Semrock), BSP01-785R (Semrock) | 100 (1,000) |
| LD 655 | 638 (0.22) | Di03-R405/488/561/635-t1 (Semrock) | BLP01-635R-25 (Semrock) | 100 (750) |

**Table S7.** Imaging conditions for single-molecule, single-dye measurements.

| Sample | $\lambda_{exc}$/ nm (power $\rho$ in kW/cm$^2$) | Dichroic Mirrors | Emission Filters | $T_{exposure}$/ ms ($N_{frames}$/FOV) |
|---|---|---|---|---|
| ATTO 520 + ATTO Rho6G | 515 (0.03) | Di03-R514-t1 (Semrock) | FF01-515-LP (Semrock) | 100 (600) |
| ATTO 565 + ATTO 594 | 561 (0.06) | Di03-R488/561-t1 (Semrock) | BLP01-568R (Semrock) | 100 (2,500) |
| ATTO 647N + Qdot 800 | 638 (0.08) | Di03-R405/488/561/635-t1 (Semrock) | BLP01-635R-25 (Semrock) | 100 (650) |
| ATTO 565 + ATTO 633 | 561 (0.03), 638 (0.14) | Di03-R405/488/561/635-t1 (Semrock) | NF03-405/488/561/635E-25 (Semrock), BSP01-785R (Semrock) | 100 (1,200) |
| Qdot 525, Qdot 585, Qdot 605, Qdot 655, Qdot 705, Qdot 800 (individuals and mixtures) | 515 (0.09) | Di03-R514-t1 (Semrock) | FF01-515-LP (Semrock) | 100 (500) |

**Table S8.** Imaging conditions for single-molecule mixture experiments.

#### Supplementary Note S34: Optical path for experiments

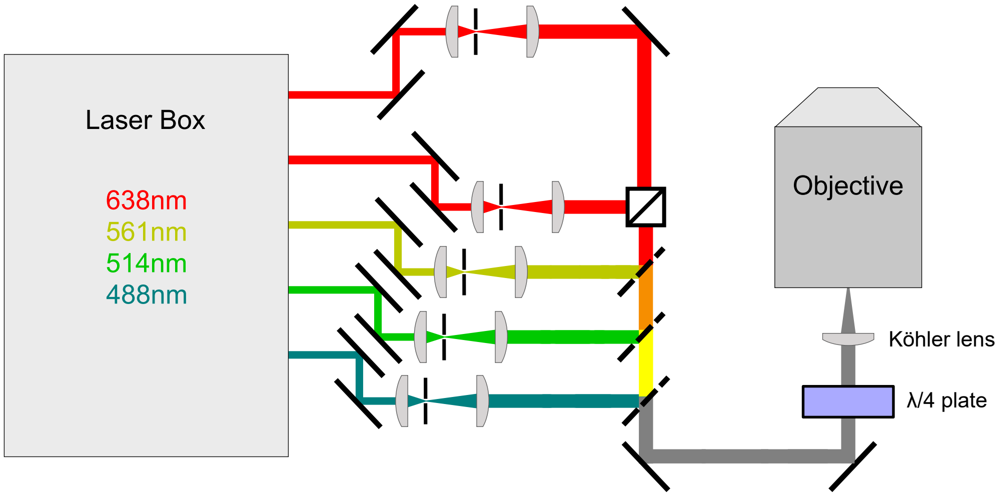


**Fig. S99.** Optical setup for experiments.
